# Macrocyclic phage display for identification of selective protease substrates

**DOI:** 10.1101/2025.03.13.643185

**Authors:** Franco F. Faucher, Scott Lovell, Matilde Bertolini, Kristýna Blažková, Emily D. Cosco, Matthew Bogyo, Marta Barniol-Xicota

## Abstract

Traditional methods for identifying selective protease substrates have primarily relied on synthetic libraries of linear peptides, which offer limited sequence and structural diversity. Here, we present an approach that leverages phage display technology to screen large libraries of chemically modified cyclic peptides, enabling the identification of highly selective substrates for a protease of interest. Our method uses a reactive chemical linker to cyclize peptides on the phage surface, while simultaneously incorporating an affinity tag and a fluorescent reporter. The affinity tag enables capture of the phage library and subsequent release of phages expressing optimal substrates upon incubation with a protease of interest. The addition of a turn-on fluorescent reporter allows direct quantification of cleavage efficiency throughout each selection round. The resulting identified substrates can then be chemically synthesized, optimized and validated using recombinant enzymes and cells. We demonstrate the utility of this approach using Fibroblast Activation Protein alpha (FAPα) and the related proline-specific protease, dipeptidyl peptidase-4 (DPP4), as targets. Phage selection and subsequent optimization identified substrates with selectivity for each target that have the potential to serve as valuable tools for applications in basic biology and fluorescence image-guided surgery (FIGS). Overall, our strategy provides a rapid and unbiased platform for effectively discovering highly selective, non-natural protease substrates, overcoming key limitations of existing methods.

## Introduction

Proteases play crucial roles in diverse aspects of normal cellular functions as well as in many pathological conditions, such as cancer. As a result, they are valuable targets for therapeutic and diagnostic agents, which are often designed based on substrate specificity of the target protease. Therefore, optimal substrates serve as a core component of inhibitors, contrast agents, and pro-drugs.^1–8^ For most applications, highly selective substrates are required to distinguish a target protease from homologous enzymes with similar active site architectures.^9^ This is particularly important for contrast agents, where off-target activation can lead to false-positive signals and increased background noise. Similarly, for therapeutic applications, lack of selectivity can result in toxicity and poor clinical outcomes.^5,10,11^

Given the importance of proteases as therapeutic targets, several methods have been developed to identify peptide substrates, including mass spectrometry-based approaches, combinatorial screens, and display libraries.^12,13^ Although mass spectrometry enables the discovery of natural peptide substrates for uncharacterized proteases, these methods are often labor-intensive and technically complex.^14–18^ Combinatorial peptide libraries provide a more accessible approach and enable the incorporation of non-canonical amino acids, which can enhance substrate selectivity. However, these libraries require extensive synthesis prior to screening, making them resource-intensive and the complexity of sequences is limited by the need to chemically synthesize individual or small pools of substrates in parallel.^19–21^ Alternatively, genetically encoded display technologies including phage display, mRNA display and DNA encoded libraries (DELs), allow efficient screening of highly diverse peptide pools (10^10^ to 10^13^).^21–25^ These technologies can be applied for the identification of substrates, yet most current applications still primarily focus on linear substrates, which are conformationally flexible and prone to off-target proteolysis. Consequently, these methods often yield non-selective substrates, which require significant medicinal chemistry efforts to improve selectivity. Hence, there remains an unmet need for a rapid screening method capable of surveying highly diverse non-linear peptides to identify selective substrates with constrained, non-canonical conformations.

Human fibroblast activation protein alpha (FAPα) and dipeptidyl peptidase-4 (DDP4) are structurally and functionally related serine proteases with therapeutic relevance in cancer and diabetes, respectively.^26,27^ FAPα and DPP4 share 68% sequence homology and belong to the S9 protease family, which includes dipeptidyl peptidase 8 (DPP8), dipeptidyl peptidase 9 (DPP9) and prolyl oligopeptidase (PREP).^28,29^ Because the S9 family of proteases share substrate specificity, preferring proline at the N-terminal position of the scissile bond, it remains challenging to identify selective peptide substrates for individual family members. To date, the best examples of FAPα and DPP4 selective substrates and inhibitors are based on small molecule inhibitor scaffolds.^30^ FAPα is overexpressed in 90% of carcinomas and multiple other cancers, where it is predominantly found in cancer associated fibroblasts (CAFs).^26^ High FAPα expression correlates with poor prognosis, making it a promising target for both therapeutic and imaging applications.^31–35^ Notably, ^18^F-labeled FAPα inhibitors have demonstrated superior specificity and signal intensity compared to the gold-standard ^18^F-Flourodeoxyglucose (FDG) radiotracer.^36^ However, despite the development of potent FAPα inhibitors, such as the proline-containing small molecule UAMC1110, these compounds suffer from cross-reactivity with DPP4, DPP9, and PREP, limiting their clinical use.^37–39^ Currently, to address these limitations in selectivity, there is growing interest in exploring alternative scaffolds, including cyclic peptides as substrates and inhibitors.^40^ The best characterized FAPα homolog is DPP4, which is a validated drug target for type 2 diabetes.^41^ Recent studies have shown that FDA-approved DPP4 inhibitors also inhibit bacterial DPP4 homologs, altering the gut microbiome and potentially leading to adverse effects.^11^ This cross-reactivity between DPP4 homologs is attributed to the small size of current DPP4 inhibitors, which provides limited selectivity. Consequently, developing non-traditional DPP4 scaffolds with improved selectivity profiles, may enable development of therapeutics with improved outcomes and reduced side effects.^42^

Here, we present a phage display-based method for the rapid identification of highly selective substrates from large pools of non-natural cyclic peptides using native, untagged proteases as targets. Our approach employs a two-step biorthogonal modification chemistry on the surface of the phage to generate macrocyclic peptide libraries featuring a protease recognition site containing a fluorogenic reporter and affinity tag.^43–45^ The covalent cyclization of peptides on the phage surface increases rigidity and enhances both selectivity and proteolytic stability of the resulting substrates.^46,47^ We have developed a direct-to-biology strategy for rapid hit validation and optimization of the selectivity and affinity of the most promising phage-selected peptides. To demonstrate the utility of our approach, we used a library comprising over one billion cyclic peptides to simultaneously screen for both FAPα and DPP4 substrates using a single proline-based linker for peptide modification on-phage. Using this approach, we were able to develop both a FAPα-selective macrocyclic peptide substrate, designed for use as a protease-activated imaging probe for fluorescence image-guided surgery (FIGS)^4,8^ and a highly selective DPP4 substrate. These results highlight the robustness of our approach for screening homologous proteases with similar active sites and substrate preferences. Overall, this method provides a rapid way to identify highly selective cyclic peptide substrates for any target protease of interest.

## Results

### Biorthogonal chemical linker design for the substrate-phage display platform

We used an fdg-3p0ss phage vector, with a disulfide-free pIII protein, encoding for a library of peptides with the AACX_7_CG general sequence.^48^ To generate unnatural macrocyclic structures we chemically modified the displayed linear library using an exogenous linker (**Figure 1a**). This linker was designed to be biorthogonal with the phage and to introduce multiple functionalities that aid in identifying selective peptide substrates for our target proteases. Specifically, the linker incorporates a proline residue, to drive binding to the active site of DPP4 and FAPα, which are P1-proline specific proteases. Adjacent to the proline, we placed a 7-amino-4-carbamoylmethylcoumarin (ACC) to facilitate real-time monitoring of library enrichment by measurement of fluorescent signals produced in the released phage. Traditionally, library selection is monitored through colony titration, a labor-intensive and time-consuming process. However, the ACC moiety enables direct fluorescence-based detection upon cleavage of the proline-ACC amide bond, providing a faster and more reproducible alternative. Additionally, the linker features a terminal biotin to immobilize the phage library onto neutravidin magnetic beads. This facilitates removal of unmodified linear peptides during the first step of enrichment. The chemical modification of the linear peptide library is performed on-phage, using a previously reported 2-step process.^43–45^ First, the displayed linear peptides are cyclized using 5-dichloropentane-2,4-dione (DPD), which reacts with the two fixed-position cysteines, and introduces a di-ketone motif. In the second step, the di-ketone undergoes a biorthogonal Knorr-pyrazole cyclization with the terminal hydrazine of the protease substrate linker to form the final library of macrocyclic protease substrates (**Figure 1b**). A key advantage of our biorthogonal linker is its tunability and accessibility. It can be synthesized using solid phase chemistry, and by modifying its active site-directing amino acids, it can be tailored to any proteases of interest. Before initiating the first panning round, we assessed the substrate scope of our linker by testing it directly against a panel of S9 proteases (**Figure S1**). As expected, FAPα and DPP4 cleaved the linker in a dose-dependent manner, with DPP4 exhibiting greater efficiency. In contrast, PREP and DPP9 showed no detectable activity.

**Figure 1.**
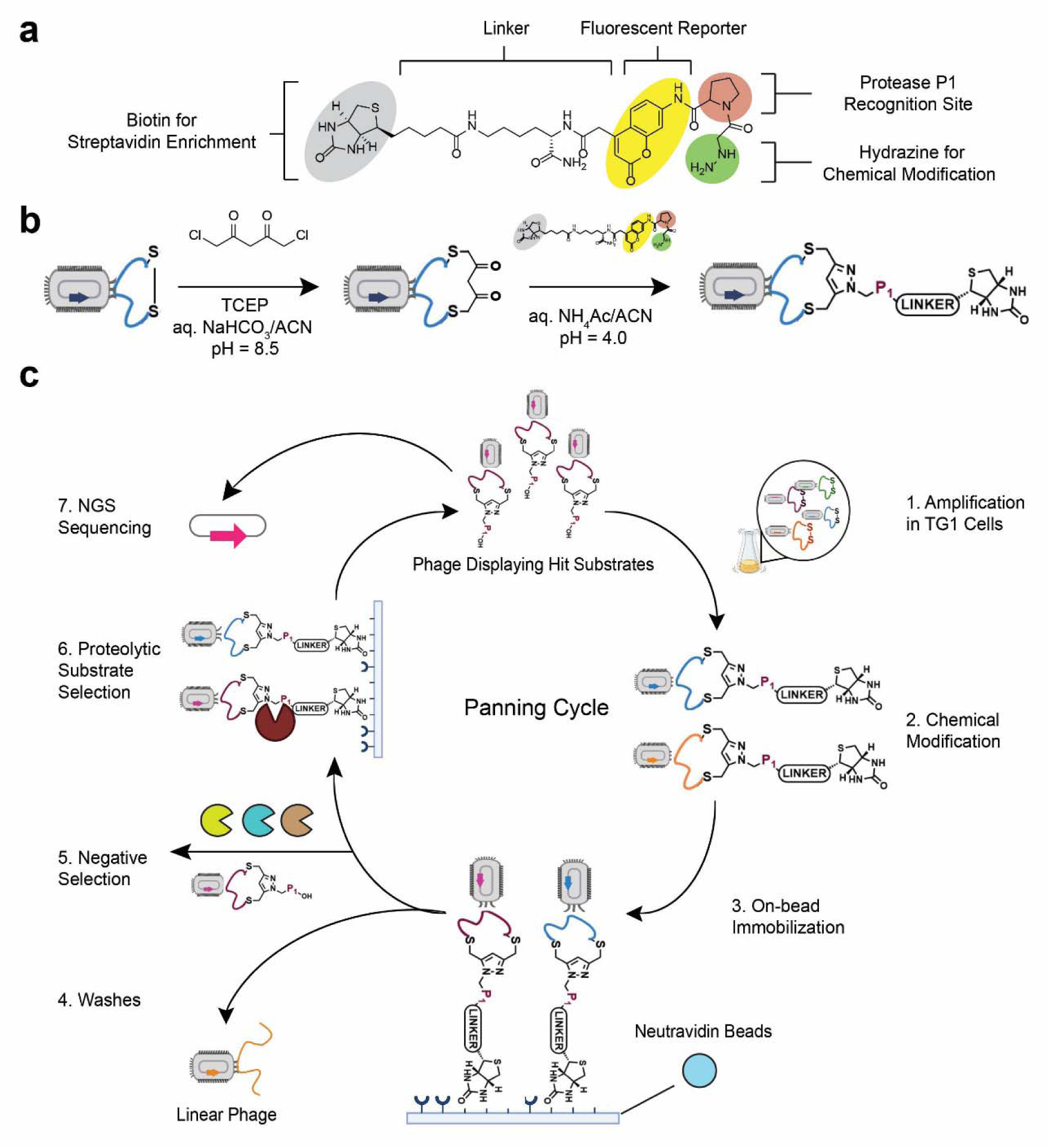
Phage display platform for direct substrate identification. **a.** Structure of the biorthogonal linker used to generate macrocyclic peptide protease substrates that includes: (1) a protease recognition site, proline is used as it is the preferred P1 of FAPα and DPP4 (2) a biotin affinity handle for isolation of phage prior to protease digestion and (3) a fluorescent reporter for direct monitoring of substrate cleavage during selection. **b.** Two-step modification of the cysteines displayed on the pIII coat protein of the phage: first, covalent cyclization with 1,5-dichloro-2,4-pentanedione (DPD), followed by the addition of the biorthogonal hydrazine linker to form a pyrazole. **c.** Schematic representation of the phage panning cycle starting from the expansion of native libraries and finishing with next generation sequencing (NGS) of the selected phage library after 5 panning rounds. A negative selection or counter-selection step with a mix of off-target proteases was performed before addition of the target protease in round 5. Round 5 of panning was performed in triplicates, which were sequenced (NGS). See **Supplementary Methods** for full panning protocol.

### Substrate-phage display identifies selective macrocyclic substrates for FAPα and DPP4

To identify selective peptide substrates for FAPα and DPP4, we subjected our unnatural library to five rounds of panning (**Figure 1c**). Prior to selection, we optimized key aspects of the panning protocol, including library immobilization, proteolytic selection, and washing conditions (**Figure S2 and SI Methods**). Once these conditions were established, we amplified the phage library in TG1 cells and subsequently modified it with our biorthogonal linker, via the two-step cyclization, yielding the final macrocyclic substrate library. Unreacted DPD was removed after the first step via Zeba column purification to prevent side reactions. The lack of toxicity of the chemical modification was confirmed by phage titration, which showed no significant changes before and after modification. On average, the modification efficiency on the pIII coat protein was 95%, as determined by biotin pulse-chase experiments (**Figure S3**).^45^ The freshly modified phage library was immobilized on neutravidin magnetic beads and submitted to nine washes with decreasing Tween concentrations, to remove unbound phage. Titration confirmed complete removal, as no colonies were detected in the final wash. Proteolytic selection was then performed by incubating the immobilized library with either FAPα, DPP4, or a no-protease control for 2 h at 37°C. Cleaved macrocycles, along with their corresponding phages, were released into the supernatant and collected for subsequent panning rounds. To favor the enrichment of higher-affinity and more efficiently cleaved substrates, we increased the stringency of the selection conditions for each round, by using decreasing concentrations of protease (**Figure 2c**). In the fifth and final selection round, a counter-selection step was introduced to eliminate non-selective sequences. Specifically, prior to adding the protease of interest, we incubated the immobilized phage library with a panel of homologous S9 proteases (**Figure 2c**) to remove substrates susceptible to off-target cleavage. After each panning round, we monitored protease-mediated selection to confirm successful library enrichment. In the first round, enrichment was assessed using traditional plaque-forming unit (PFU) titration (**Figure S4**). For subsequent rounds, we implemented a real-time fluorescence tracking strategy using the ACC fluorogenic reporter in our linker, which becomes fluorescent upon proteolysis. By comparing fluorescence intensity between protease-treated and control libraries, we could track substrate cleavage and, therefore, enrichment. A neutravidin bead-only control was also included to account for background fluorescence. We observed enrichment of the FAPα-treated library after each round. Importantly, even after the counter-selection step, fluorescence intensity in round five exceeded that of all previous rounds, indicating efficient selection of FAPα-specific substrates (**Figure 2a**). Similarly, the DPP4-treated library exhibited enrichment after rounds 2, 3, and 5, with the highest fluorescence signal detected in round five (**Figure 2b**).

**Figure 2.**
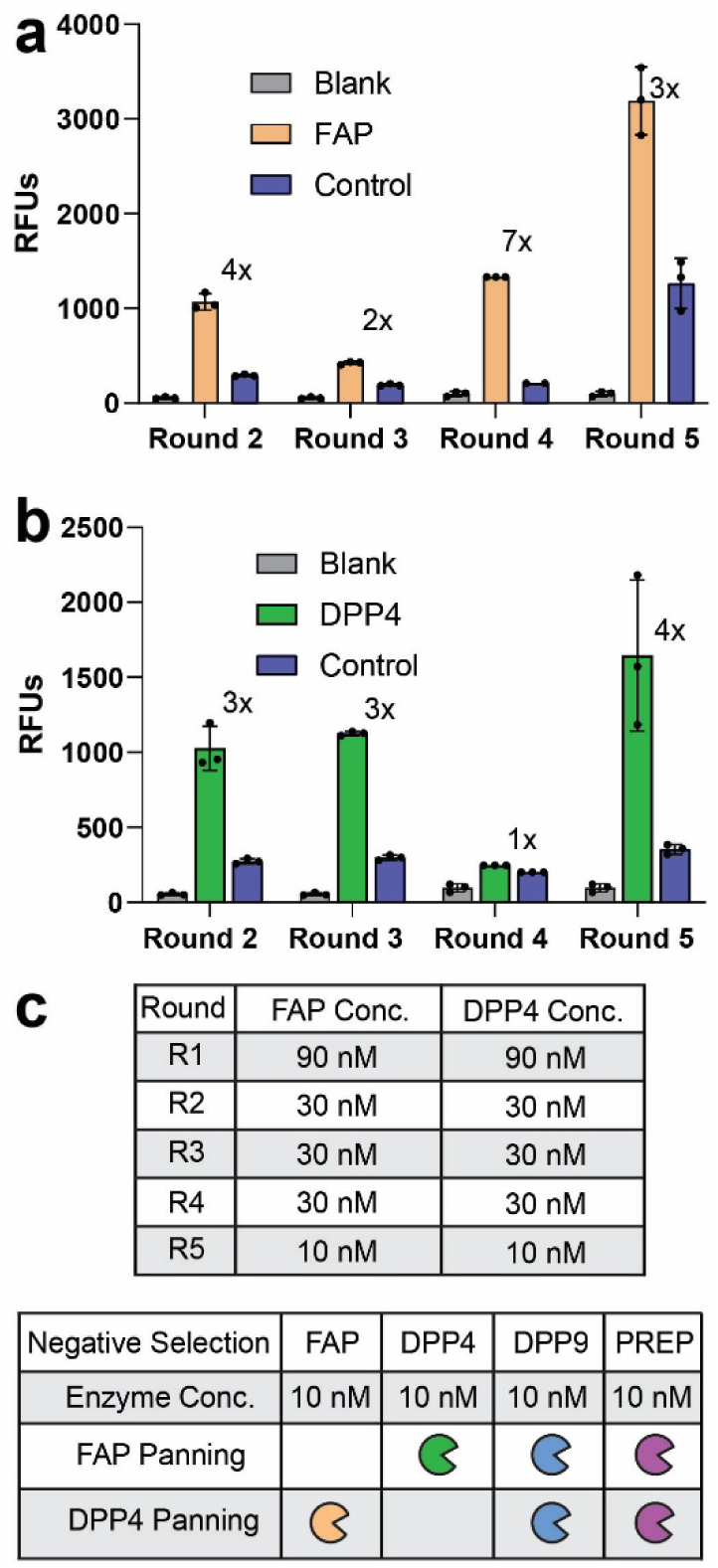
Protease concentration used for selection and library enrichment after each round of panning. **a. b.** Library enrichment after each panning cycle for FAPα (**a**) and DPP4 (**b**). The fold enrichment of protease vs control (no-protease treated library) is indicated. Enrichment was determined in triplicates by monitoring free ACC fluorescence after proteolytic cleavage. Shown as relative fluorescence units (RFUs). Error bars represent the standard deviation. Note that round 1 enrichment was monitored via traditional tittering and is available in **Figure S4**. **c.** Protease concentrations used per panning round and for negative selection. Off-target proteases were pre-mixed before their addition for negative selection.

After the five selection rounds, the resulting enriched phage libraries were amplified in TG1 *E. coli*, plasmid DNA was extracted, and next-generation sequencing (NGS) was performed to identify consensus substrate sequences. To select FAPα and DPP4-specific cleavable sequences from our NGS data, we developed a rapid hit identification and substrate validation workflow. First, we analyzed the NGS data using an RStudio pipeline that retrieves unique sequences in the library selected against our target protease, by comparing it to the control library output. This pipeline translates the amplified variable pIII region codons into amino acids and performs a differential enrichment analysis using a negative binomial model. It also implements quality control parameters tailored to our library, filtering out peptides with sizes deviating from the expected 13 amino acids, mutations in fixed positions, or sequences containing more than two cysteines (see **SI Methods**). Using this pipeline, we chose peptide sequences with the desired fold-enrichment relative to the control, ensuring a false discovery rate below 0.05 (**Figures S5 and S6**). Peptides were then clustered using the GibbsCluster 2.0 algorithm, and the top-scoring hits from each cluster were designated as representative sequences for the consensus motifs.^49^ In the FAPα-primed library, peptide hits were classified in 6 unique clusters, based on sequence similarity. A total of 33 FAPα-specific peptides, covering all clusters, were selected for synthesis. Of note, several clusters showed clear patterns such as the SLRINSL motif in cluster 1 (**Figure 3c**). For the DPP4 panning, peptide hits were classified into seven clusters, and 39 sequences were synthesized, including two non-selective sequences as negative controls (**Figure S7**). When testing phage-display hits, the purification step often represents a major bottleneck. To circumvent this, we implemented a direct-to-biology approach by testing crude peptides in a fluorescent enzymatic assay. This assay leverages the fluorescent properties of ACC within the substrate structure, which remains optically silent when the substrate is intact. However, upon protease cleavage at the proline site, the ACC is excised and fluoresces at 445 nm after excitation with 345 nm light (**Figure 3a**). This mechanism provides a straightforward readout of peptide activity, enabling rapid evaluation of their potential as selective substrates. To achieve a reliable readout, we optimized substrate preparation with a two-step synthesis. First, linear peptides were synthesized via solid-phase chemistry and cyclized using DPD. Next, the cyclized peptides were reacted with the linker to form the final macrocyclic protease substrates, which contained both proline and the ACC fluorophore, ready for direct testing (**Figure 3b** and **Supplementary Methods**). In all cases, in the crude mixture, the desired macrocyclic peptide was the predominant product, with trace amounts of impurities, such as unreacted linker (see **Supplementary Spectra**).

**Figure 3.**
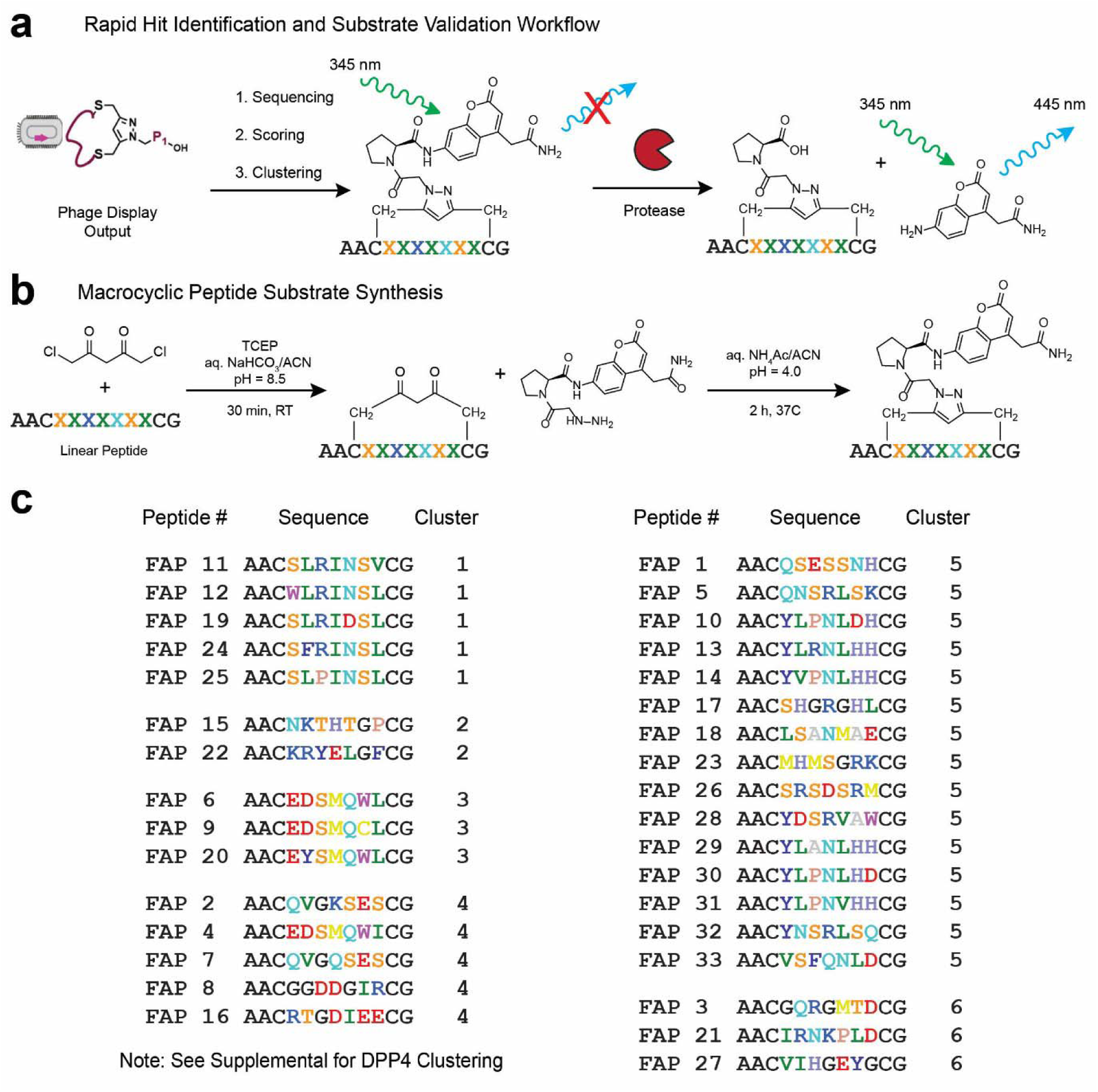
Macrocyclic peptide substrate identification and synthesis. **a.** Schematic representation of the substrate validation process. Once the selected phage library is sequenced and the output NGS data is analyzed, the peptide hits are selected, synthesized and tested as crude substrates to identify promising leads. **b.** Two-step synthesis of macrocyclic peptide substrates from linear peptides. See **Supporting Information** for detailed synthetic methods. **c.** Top-hit peptide sequences synthesized for FAPα. These sequences were selected from a larger number of peptide hits, which were classified in 6 clusters using GibbsCluster.^49^ Numbers represent the enrichment order calculated from the pipeline. Constant amino acids at the peptides’ N and C-terminal are indicated in black. Variable amino acids (X) are colored using the RasMol coloring scheme, except glycine (G) that is colored black for visual purposes.

### Validation and structural optimization of the non-canonical FAPα-selective substrates

To evaluate the potential of the 33 selected peptide hits as FAPα substrates, we tested them using the previously described fluorogenic assay (**Figure S8**). Of the 33 crude peptides, 30 showed fluorescence after incubation with FAPα, indicating cleavage by the protease. We selected the 7 peptides with the highest fluorescence values and purified them for further analysis. The purified macrocycles were tested against FAPα and against a panel of S9 proteases including DPP4, DPP9, and PREP, to assess their selectivity (**Figure 4a**). All tested peptides were cleaved by FAPα, with FAP-24 and FAP-33 showing the highest specificity. FAP-33 exhibited 10, 18, and 3-fold selectivity over DPP4, DPP9, and PREP, respectively while FAP-24 displayed 9, 13, and 6-fold selectivity over these same proteases. To confirm that activity and selectivity arose from the phage-selected peptide sequences, we prepared scrambled versions of FAP-33 and FAP-24 (FAP-33s and FAP-24s). These scrambled variants showed a loss of activity and diminished selectivity, validating the sequence specificity of the identified selective peptides (**Figures 4b and 4c**). To improve the activity and selectivity of our top hit substrate we performed medicinal chemistry efforts to introduce simple structural modifications. First, we examined the importance of the amino acids in the fixed positions outside of the macrocycle by synthesizing truncated versions (**Figure 4b and 4c**). From these variants, macrocyclic substrate FAP-33.4, was not synthetically accessible due to the poor solubility of its linear precursor. Interestingly, the truncation of one N-terminal Ala reduced FAPα activity and selectivity, however eliminating both N-terminal fixed Ala residues enhanced FAPα activity and retained or improved selectivity (**Figure 4b and 4c**). From the truncated analogs, we selected the most promising substrate FAP-33.2 for further optimization. An alanine scan of FAP-33.2 identified serine in position 2, and leucine in position 6, as potential modification points, as their replacement with Ala did not reduce FAPα activity. In fact, the modified FAP-33.2A6, with an alanine at position 6, was a more optimal substrate than the parent FAP-33.2 while also retaining selectivity. All other positions were found to be essential, with their substitution resulting in a loss of activity (**Figure S9**). We also tested alkylation of the N-terminus of FAP-33.2 by introducing acetyl or terminal alkyne groups, both of which led to a loss of activity.

**Figure 4.**
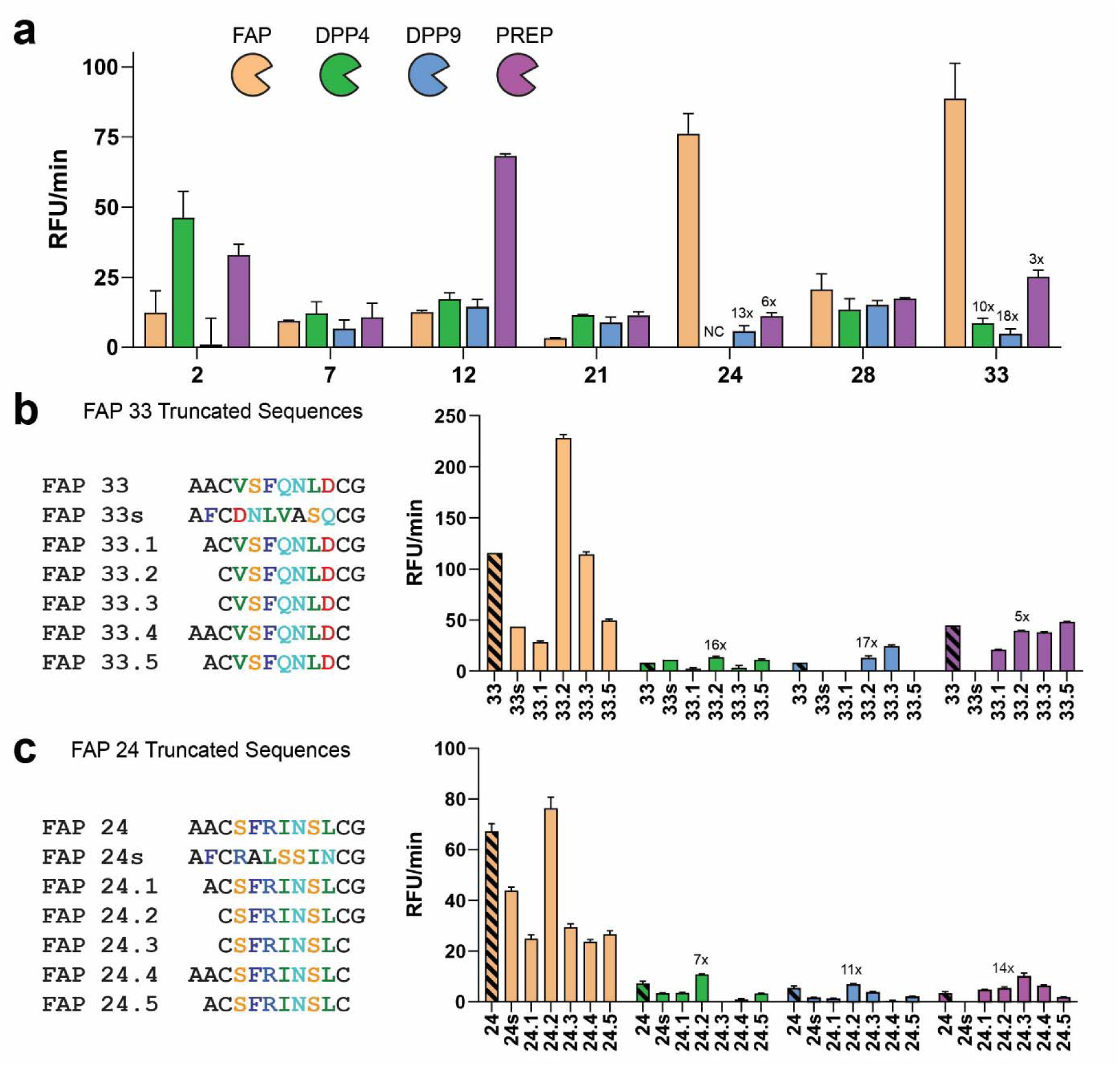
FAPα-selective fluorogenic macrocyclic substrates. **a**. Selectivity data for purified FAPα peptides. Peptides selected after crude testing were purified and screened at 200 µM at 37°C against a panel proteases: 10 nM FAPα (orange) and its homologs 2 nM DPP4 (green), 6 nM DPP9 (blue) and 6 nM PREP (purple). The selectivity for FAPα vs the off-target protease is shown over the bar for selected substrates (NC: not calculated, due to undetected fluorescence). **b. and c.** Structure-activity series for peptide 33 (**b**) and 24 (**c**) including random sequence scrambling and truncation of N and C-terminal residues. The activity of each substrate for FAPα and each of the 3 off-target proteases is plotted as RFU/min. Protease concentrations are 10 nM FAPα (orange), 2 nM DPP4 (green), 6 nM DPP9 (blue) and 3 nM PREP (purple). The original reference peptides are represented in stripes and the selectivity for FAPα vs the off-target protease is shown for select substrates. Error bars represent the standard deviation.

We next screened several unnatural amino acids (UAA) at non-essential positions 2 and 6, and at critical position 3. Replacing phenylalanine in position 3 with its aliphatic equivalent abolished activity, confirming the need for an aromatic ring in that position. However, even when selecting UAAs containing 6-membered aromatic rings able to engage in π-π stacking interactions, all modifications resulted in a substantial reduction of activity, revealing a tight structure-activity relationship at position 3 (**Figure S10**). At position 2, most modifications caused a reduction of activity, except for replacing the original isopropyl chain with a methoxymethyl chain, which retained activity. On the contrary, position 6 had greater tolerance for structural diversity, and introducing 2,4-diaminobutyric acid (Dap) yielded FAP-33.28, a substrate with activity comparable to the commercial substrate z-Gly-Pro (**Figure S11**).

### Validation and structural optimization of non-canonical DPP4 substrates

Using the same phage library used for FAPα to screen DPP4, we identified 61 peptide hits and selected the top 37 sequences for synthesis, along with two negative controls. Of note, less than 2% of the identified DPP4 peptide hits were found in the FAPα panning, highlighting the specificity of our approach. To evaluate the identified DPP4 substrates, we directly assessed selectivity by testing the crude macrocycles against DPP4, alongside the homologous proteases FAPα and DPP9. We selected the substrates with a favorable selectivity index for DPP4 and fluorescence levels exceeding those of the negative controls, to further confirm their potential as DPP4 selective substrates (**Figure S12**). We purified and retested the eight top candidates, and found that only peptide DPP4-10 retained selectivity. DPP4-10 displayed a 20-, 4-, and 3-fold preference for DPP4 over FAPα, DPP9, and PREP, respectively (**Figure 5a**). We synthesized three randomly scrambled analogs. Surprisingly, one of them, DPP4-10s, exhibited greater activity and improved selectivity for DPP4 than the original peptide. The two additional scrambled analogs were not as active or as selective as DPP4-10s (**Figure 5b**). This relative minimal impact of scrambling may be due to the fact that the DPP4-10 peptide contains two alanines, two histidines, two serines, and two amide-containing residues (Asn and Gln). This overall low sequence diversity might enable the preservation of structures which display amino acids in similar ways despite scrambling. To further probe the structure–activity relationship of DPP4-10s, we synthesized and tested truncated variants with deletions in the conserved region. All truncations led to a loss of both activity and selectivity (**Figure 5c**), confirming the sequence-dependent nature of DPP4-10s activity.

**Figure 5.**
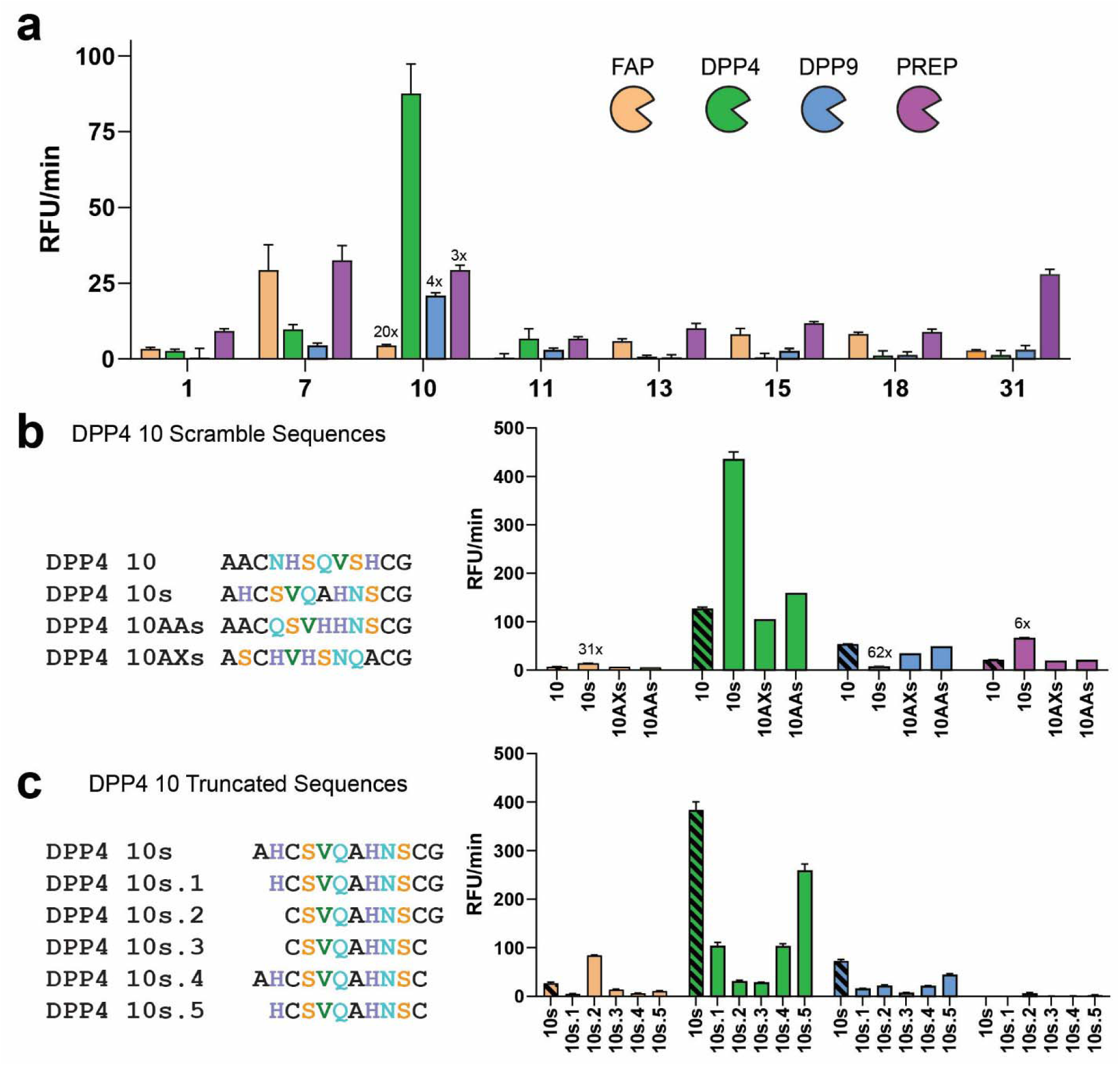
DPP4-selective macrocyclic peptides identified using chemically modified phage display. All substrates were tested at 200 µM at 37°C in triplicates against 2 nM DPP4 (green) and its homologs: 10 nM FAP (orange), 6 nM DPP9 (blue) and 3 nM PREP (purple). The activity of each substrate for DPP4 and the three off-target proteases is plotted as RFU/min. **a**. Selectivity data for purified DPP4 peptides. Peptides selected after crude testing (**Figure S12**) were purified and screened against a panel proteases. **b**. Structure-activity analysis of peptide DPP4-10 scrambled analogs, generated by randomizing the amino acid sequence. **c**. Structure-activity analysis of DPP4-10s analogs with sequential elimination of N- and C-terminal residues.

### Phage display-identified peptides are selective substrates for FAPα or DPP4 in cells

After confirming that our FAPα- and DPP4-selective macrocyclic peptides could discriminate among homologous proteases, we evaluated their performance in a complex cellular context. To model cancer conditions relevant for FAPα-targeted fluorescent contrast agents, we used HEK293 Tet-Off cells with inducible FAPα expression. Initially, we validated our experimental setup by benchmarking the commercial FAPα and DPP4 substrates zGP-AMC and GP-AMC in the presence of the reversible covalent inhibitors UAMC1110 (FAPα) and VbP (DPP4) (**Figure 6** and **Supporting Information**). We confirmed that FAPα activity was detected only in induced cells, no signal observed in wild-type or doxycycline-treated (Tet-Off) HEK293 cells (**Figure 6**). Of note, we observed that the zGP-AMC substrate was processed in the presence of both inhibitors suggesting it can be cleaved by proteases other than FAPα and DPP4 (**Figure 6a, S13**). DPP activity corresponding to DPP4 as well as other DPP-like proteases was detected in all cell lines (**Figure 6b**). Using this assay, we assessed the selectivity of our optimized macrocyclic substrates for FAPα and DPP4. Because FAP-33.28 proved challenging to synthesize, we used FAP-33.2A6 and FAP-33.2, for the next steps. FAP-33.2A6 was selectively cleaved in FAP+ cells, with hydrolysis suppressed by the FAPα inhibitor UAMC1100 but not by the DPP4 inhibitor VbP, confirming its FAPα specificity even in a complex cellular environment (**Figure 6c**). Surprisingly, its analog FAP-33.2 showed no activity in cells, likely due to poor solubility (**Figure S14**). FAP-24.2 was a FAPα substrate in cells, though less selective than FAP-33.2A6, as pre-treatment with VbP partially reduced its fluorescence signal, suggesting some processing by DPP4 (**Figure 6d**). This off-target proteolysis likely explained the faster hydrolysis of FAP-24.2 compared to FAP-33.2A6 in cells, a result that contrasted with findings from purified enzymes. Importantly, both substrates did not show any processing by other proteases in HEK293 cells, demonstrating their dependence on FAPα for efficient cleavage (**Figure 6d**). To test DPP4-10s, we spiked cells with recombinant DPP4 to increase the detection window (**Figure S13**). This substrate was selectively cleaved by DPP4, remaining resistant to overexpressed FAPα and other cellular proteases (**Figure 6e**). Hence, we validated our phage display approach as an effective method for identifying macrocyclic peptide substrates that are both active and highly selective for their protease targets, even in complex cellular environments. Our optimized cyclic peptide substrates demonstrated superior selectivity compared to commercial substrates, underscoring the potential of our platform for developing probes with the high levels of specificity required for use as imaging and therapy agents.

**Figure 6.**
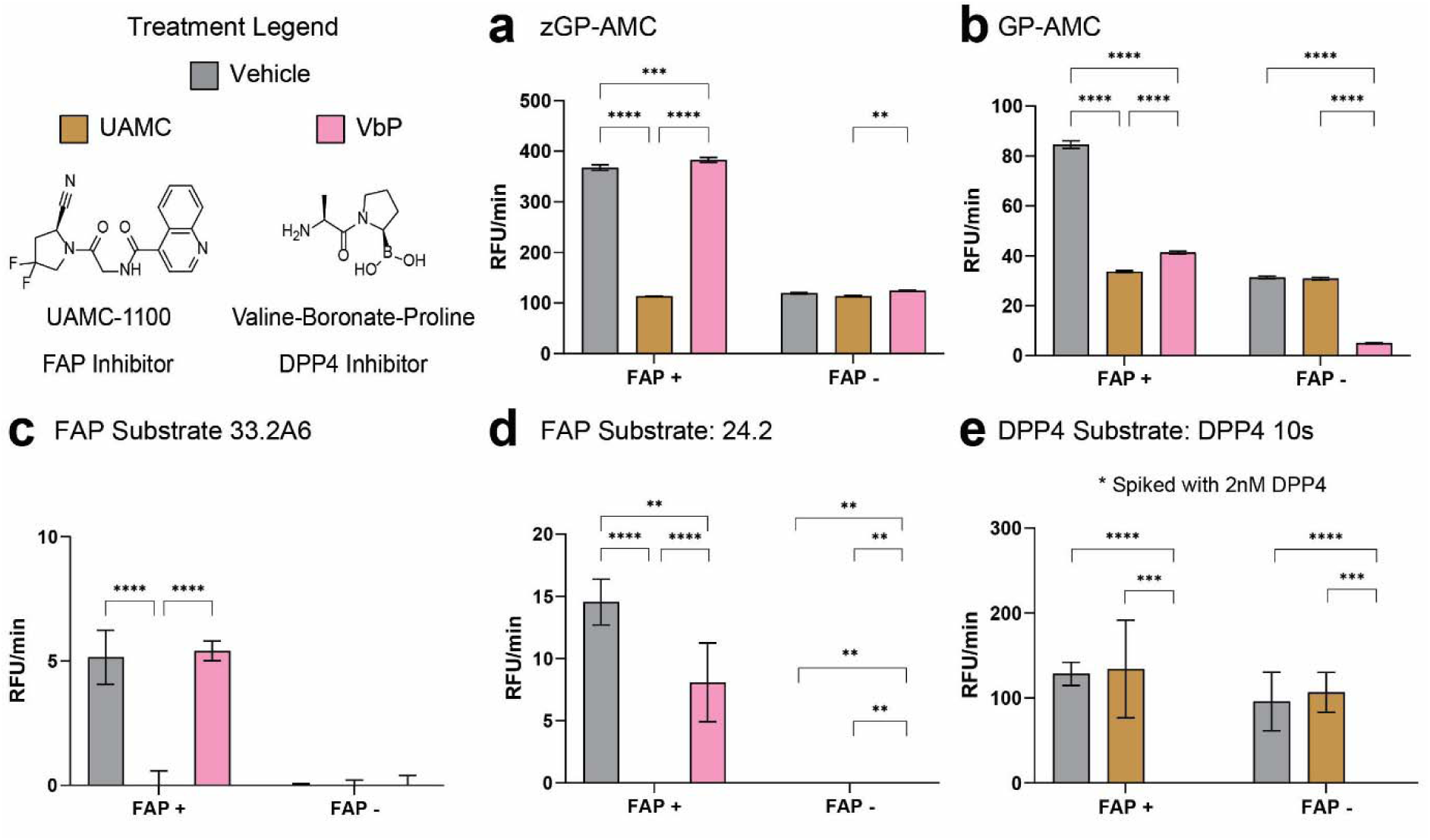
FAPα and DPP4 substrates identified via chemically modified phage display exhibit target-specific activity in cells. HEK293 cells expressing FAPα (FAP+), HEK293 cells with doxycycline-supressed FAPα expression (FAP-) and wild type HEK293 cells (**Figure S13**) were used to assess *in vitro* substrate cleavage. In all experiments, prior to 200 µM substrate addition, cells were treated with either DMSO (vehicle) or 1 µM covalent inhibitor. Substrates used were: **a.** zGP-AMC, **b**. GP-AMC, **c.** FAP-33.2A6, **d.** FAP-24.2 and **e.** DPP4-10s. All experiments were performed at 37°C in triplicates, the mean slope in RFUs per minute is plotted where the error bars show one standard deviation. A two-way ANOVA was performed with Tukey’s (**a-d**) or Sidak’s (**e**) multiple comparisons test. P values represent: **<.005, **<.0005,****<.0001. **a.** Fluorescence of zGP-AMC in FAP+ cells was diminished upon UAMC treatment, but not with VbP, indicating FAPα processing. Notably, this substrate produced residual fluorescence that could not be inhibited, in all three cell lines, suggesting the involvement of proteases other than FAPα and DPP4 with shared substrate specificity. **b**. DPP4 activity profile screened using GP-AMC. FAP+ cells show DPP-like activity, which partially corresponds to overexpressed FAPα, as shown by partial inhibition by UAMC. All cell lines showed DPP activity, some corresponding to DPP4, as confirmed by reduction of this activity by VbP. **c.** FAP33.2A6 selectively detects FAPα activity in cells with no observed off-targets **d**. FAP24 detects FAPα activity in cells, being partially hydrolysed by other proteases **e.** DPP4-10s is a selective substrate for DPP4, even under conditions of FAPα overexpression. To test DPP4-10s, cells were supplemented with recombinant DPP4 (2 nM) due to the low basal DPP4 activity in these cells.

## Discussion

Identification of protease substrates has traditionally relied on libraries of linear peptides or small-molecule fragment screens. Here, we introduce an alternative strategy that employs chemically modified phage display to generate macrocyclic peptide substrates distinct from those identified through conventional methods. Macrocyclic peptides, due to their conformational constraints, offer reduced entropic penalties upon binding, enhancing target engagement.^33,34^ However, constraining the backbone of peptides requires screening of exceptionally large pools of potential substrates since small changes in the rigid backbone often have large impacts on target binding. To overcome this significant challenge for rigid cyclic peptide substrates, we chose to use phage display methods which enable screening of billions of chemically diverse peptides. Using an established 2-step linker chemistry, we were able to introduce both a core substrate recognition sequence as well as a tags for purification and quantification of the cleaved peptides. The resulting strategy enables direct screening using small quantities of native (i.e. untagged) proteases. Furthermore, the ease of chemical synthesis of the linker enables the approach to be tailored to any protease of interest by building in an initial substrate recognition sequence into the linker. Ultimately, this approach could be used to select for cyclic peptide substrates that are selective for relevant biological samples that contain any protease activity or mixture of activities.

To validate our approach, we selected FAPα and DPP4, two clinically relevant and homologous serine proteases. Our method successfully identified non-canonical macrocyclic peptide substrates that were efficiently cleaved by the chosen target protease. To generate the non-canonical substrate library, we cyclized the displayed peptides on-phage using a biorthogonal chemical linker. Importantly, despite the inherent activity of our substrate linker for DPP4 and FAPα (**Figure S1a**), we were able to tune selectivity through selection pressure in multiple rounds of panning. In addition, we used the same macrocyclic peptide substrate library for both DPP4 and FAPα even though one (FAPα) is an endopeptidase and the other (DPP4) is an exopeptidase. These differences in substrate preferences between the two enzymes highlight our method’s ability to accommodate proteases with distinct positional specificities, demonstrating its broad applicability.

Our approach also enabled the discovery of substrates capable of distinguishing a specific protease among several homologs from the same family (**Figures 4 and 5**). This suggests that macrocyclization enhances selectivity through both active site and exosite interactions. In addition to achieving selectivity within homologous proteases, it is crucial that a substrate resists proteolysis by other endogenous proteases. To address this, we tested the selectivity of our optimized peptide substrates using a cell-based assay in which many off-target proteases likely exist. We found that the commercially available ‘selective’ substrates, zGP-AMC and GP-AMC, produced signals in HEK cells even when target proteases were chemically inhibited (**Figure 6a and 6b**). In contrast, our macrocyclic substrates showed no processing by off-target proteases. Furthermore, our optimized DPP4 substrate, DPP4-10s, remained completely resistant to off-target cleavage, even in conditions where its target homolog, FAPα, was overexpressed, underscoring the high selectivity achieved through our method (**Figure 6e**). Our results demonstrate that our phage display approach effectively yields highly selective macrocyclic peptide substrates for a protease of interest.

The design of the peptide library is fundamental for the success of any display method. The library composition dictates key structural and chemical properties of the encoded peptides, including macrocycle size, strain, and chemical diversity. We employed a library with the structure AACX₇CG, where the first alanine serves as the N-terminus and X represents one of the 19 proteinogenic amino acids, excluding cysteine. This macrocyclic design was selected to balance diversity and structural constraint. The seven variable positions produce a theoretical diversity exceeding 1 billion sequences, while the nine-membered ring provides conformational rigidity while maintaining moderate flexibility compared to smaller cyclic peptides. While this design enabled efficient cleavage of the phage-displayed peptides by both FAPα and DPP4 (**Figure 2a and 2b**), the identified substrate hits exhibited varying levels of activity, often with slow cleavage rates, highlighting opportunities for structural optimization. For FAPα substrates, we found that the most effective optimization strategy involved removing the N-terminal alanine residues (**Figures 4b and 4c**). This suggests that our initial library structure was suboptimal for FAPα and that screening a truncated library, such as CX₇CG, could yield alternative peptide sequences with enhanced activity. Despite these potential limitations, the chosen macrocycle size and strain parameters may have been critical in conferring high selectivity to our peptides, which outperformed the commercial substrate zGP-AMC (**Figure 6**).

Similar to FAPα substrates, DPP4 substrates also exhibited slow cleavage kinetics; however, any modifications to the best hit, DPP4-10s, resulted in reduced activity. An explanation could be that the replacement of an alanine in a fixed position with a histidine in DPP4-10s, might have enabled an optimal interaction with the target (**Figure 5**). This finding suggests that by keeping the N-terminal alanine constant we may have excluded sequences with stronger target engagement. Future library designs incorporating variability at the N-terminal position could improve both activity and selectivity of the initial hits. While further optimization, such as the incorporation of non-canonical amino acids, could enhance the properties of lead compounds, refining the initial library design could yield substrates with improved catalytic efficiency and selectivity. A next-generation library might include multiple macrocycle sizes or introduce variability at currently fixed positions to expand the chemical and structural diversity of potential hits. Taken together, our results demonstrate that our phage display method is a powerful tool for rapidly identifying selective cyclic peptide substrates.

An important objective when developing our approach was to identify neo-substrates with high selectivity for use as contrast agents and therapeutic inhibitors. The identified FAPα and DPP4 macrocycles exhibit a selectivity profile that, with further optimization, could facilitate their development for *in vivo* applications. For FAPα substrates, these could be leveraged to prepare fluorescent image-guided surgery probes. By optimizing their optical properties, such as conjugating a near-infrared fluorophore and a quencher, their specificity for FAPα could provide a tumor-specific signal, useful for a variety of solid tumor surgeries. This would result in FAPα-targeting fluorescent contrast agents for surgical applications, similar to those developed for cathepsin proteases.^4,50^ Another avenue for development could involve replacing the ACC fluorophore in our peptides with an active-site-directed warhead, transforming these substrates into covalent inhibitors for theranostic aplications.^51^ Similarly, our DPP4 substrate could serve as a starting point for developing a selective DPP4 inhibitor, potentially offering improved patient outcomes compared to existing commercial inhibitors.^42^ Future studies should focus on assessing its specificity for human DPP4 over homologous microbial enzymes in the gut. Overall, our method has successfully yielded selective, macrocyclic substrates for FAPα and DPP4, establishing a strong foundation for the discovery of clinically relevant neo-substrates.

## Conclusion

We have developed a chemically modified phage display approach for the identification of selective macrocyclic peptides targeting proteases. We demonstrated the value of our method by identifying novel non-canonical fluorescent macrocyclic substrates for two proteases, FAPα and DPP4. These substrates exhibited high selectivity, up to 60-fold, against homologous proteases and showed overall low off-target cleavage in cells, outperforming commercial substrates. Importantly, our phage display approach enables direct screening of native proteases or biological samples with protease activities. Furthermore, screens can be tailored to specific peptidases by introducing defined substrate sequences in the linker. Our approach also enables direct and quantitative fluorescence monitoring of panning efficiency and a direct-to-biology approach for rapid hit validation. These strategies could be applied to enhance other display formats, such as mRNA display. In conclusion, we have developed a phage display platform to efficiently identify highly selective, non-canonical protease substrates that leverage an unnatural macrocyclic structure to achieve high target selectivity.

## Supporting Information

Supporting information includes: supplementary figures, supplementary table, synthetic schemes, biology methods, chemistry methods, HPLC purity analysis, risk statement.

## Supporting information

Supplmental Spectra

## Acknowledgements

This work was supported by “la Caixa” Foundation (ID 100010434) by a Junior Leader grant to M.B.-X. (LCF/BQ/PI22/11910012). NIH grant (R01 EB026285 to M.B.), Stanford Cancer Institute Translational Oncology Program seed grant (to M.B.), the National Science Foundation Graduate Research Fellowship under grant no. DGE-1656518 (F.F.), Stanford ChEM-H O’Leary-Thiry Graduate Fellowship (F.F.), NIH Stanford Graduate Training Program in Biotechnology T32GM141819 (F.F.), International Alliance for Cancer Early Detection (ACED) Graduate Fellowship (F.F.), and Stanford Molecular Imaging Scholars (NIH T32CA118681, to E.D.C.). We would like to thank the group of Jan Konvalinka (IOCB Prague), namely Jana Starkova and Filip Wichterle, for the stable clonal HEK293 tet-off FAP cell line. We thank Jef Callebaut for helping with scripts for data analysis.

## Competing interests

The authors have no competing interests to declare.

## TOC

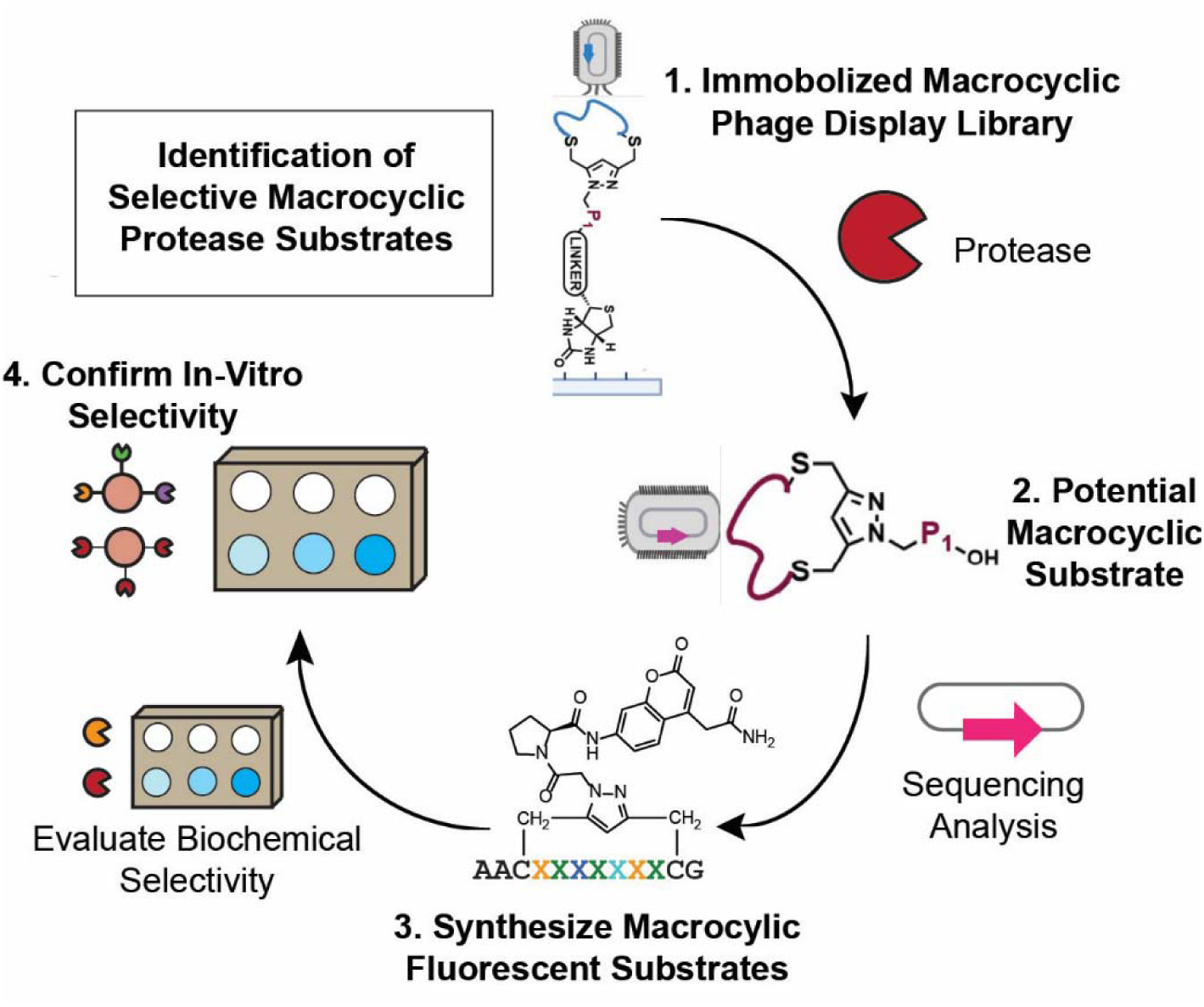

## Notes

### Competing Interest Statement

The authors have declared no competing interest.

