## Supplemental Information and Figures for "Macrocyclic phage display for identification of selective protease substrates"

Supplementary Table.	17

Supplementary [Methods	18](#_Toc157009298)

[**Phage**](#_Toc157009299) **Display Methods.	20**

**Supplementary Figures**

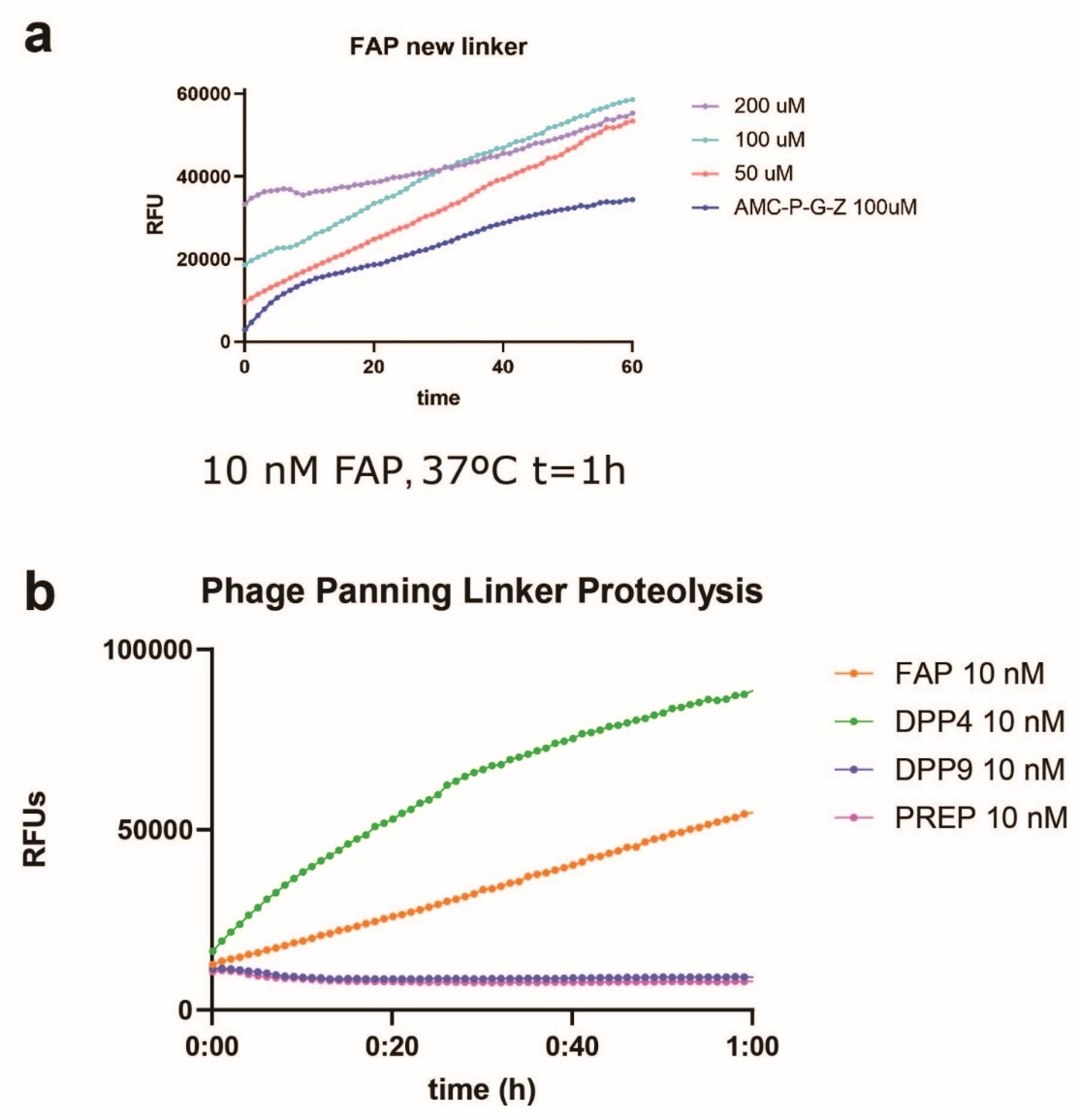

**Supplementary figure 1. Fluorescence assessment of the biorthogonal linker used for phage panning**. **a.** Decreasing concentrations of the biorthogonal linker treated with 10 nM FAPα. FAPα commercial substrate zGP-AMC is included for comparison. Proteolysis is represented in RFUs vs time. Substrate proteolysis was monitored for 1h at 37°C. **b.** The hydrolase activity of selected S9 proteases was monitored using fluorescence. The graph shows hydrolysis of the biorthogonal panning linker at 100 μM, upon incubation with FAPα 10 nM, DPP4 10 nM, DPP9 10 nM, and PREP 10 nM for 1 hour at 37°C.

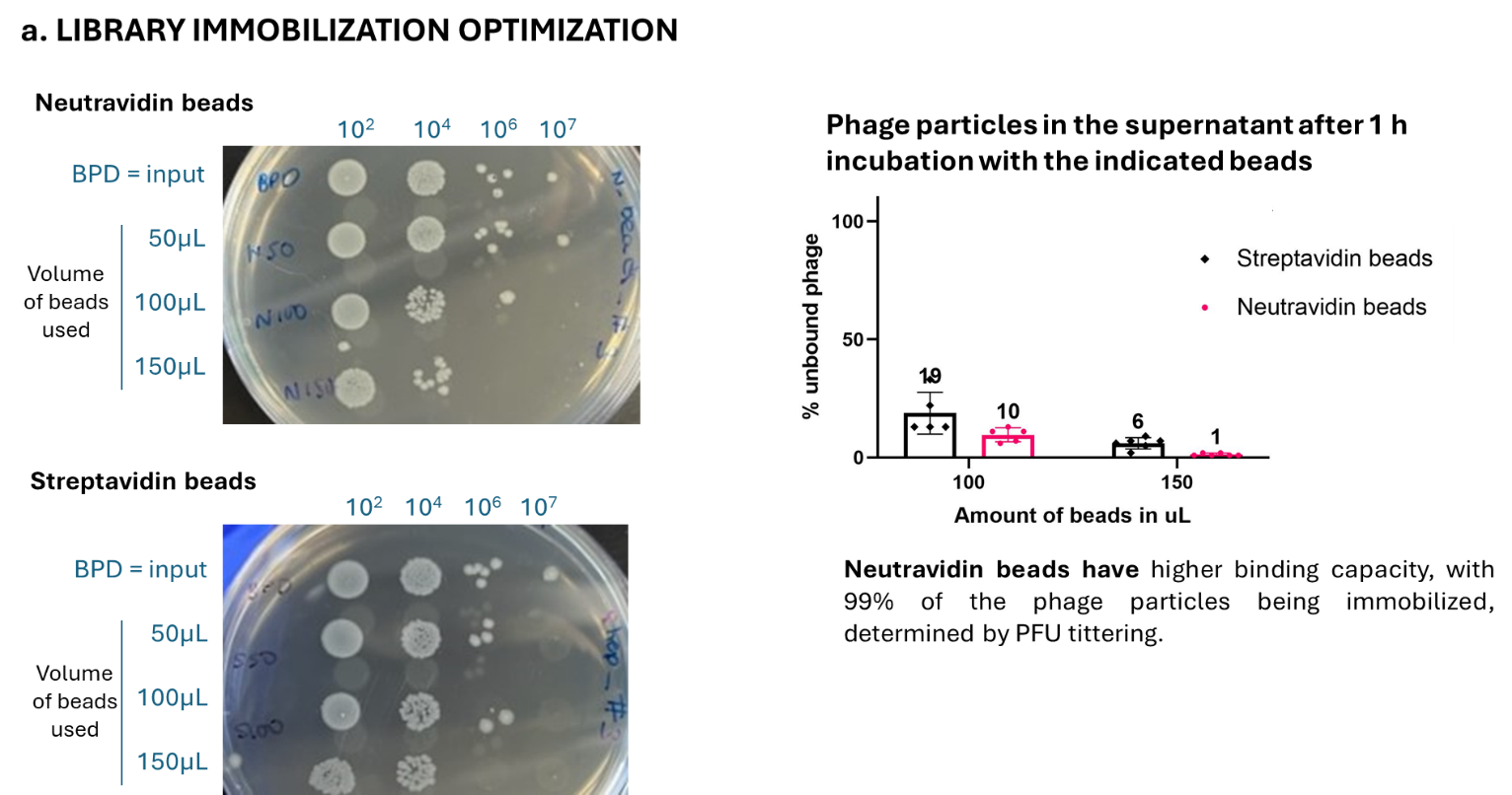

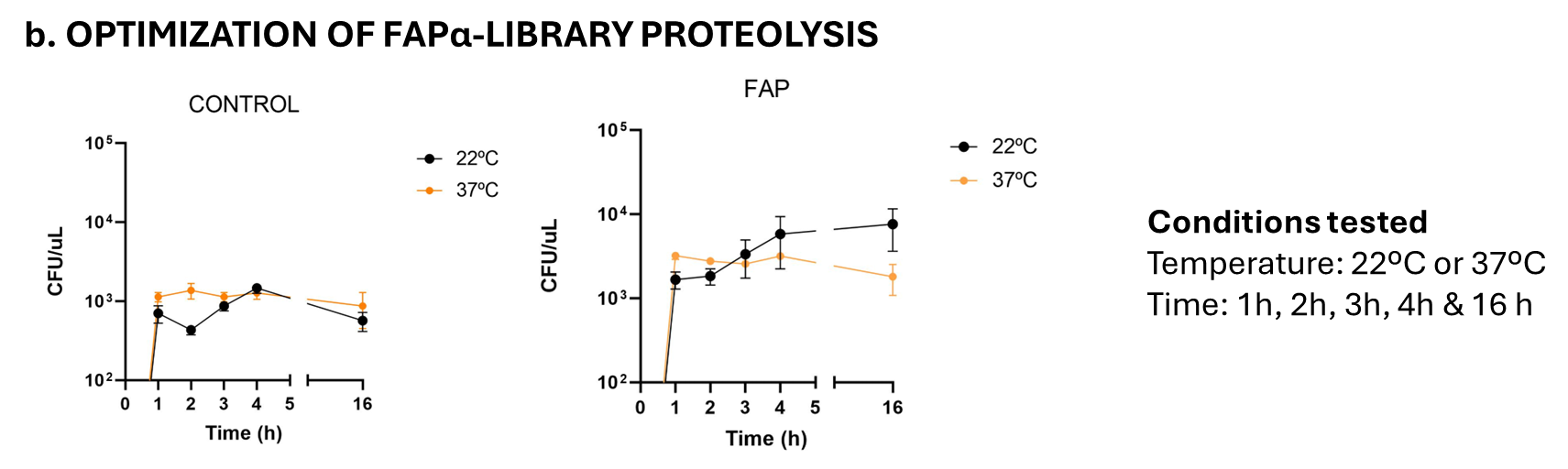

**Supplementary figure** **2**.**Optimization of phage panning conditions. a**. Library immobilization. Comparison of streptavidin and neutravidin beads. PFU colony counting shows that a library of 1.3x10^9^ phages can be fully immobilized (<1% free phage) on-bead after 1 hour incubation with 150 µL of neutravidin beads. Under the same conditions, streptavidin beads leave can only immobilize 94% of the equivalent phage library. **b.** Proteolytic selection. Immobilized peptide library (1.5x10^9^ phages per tested condition) was treated with 90 nM FAPα under the indicated temperatures and times. The optimal conditions are 37ºC at 1 hour, showing high proteolysis and consistency across the replicates, as determined by PFU tittering. The control represents the no-protease treated library. All experiments were performed in triplicates and the error bars represent the standard deviation.

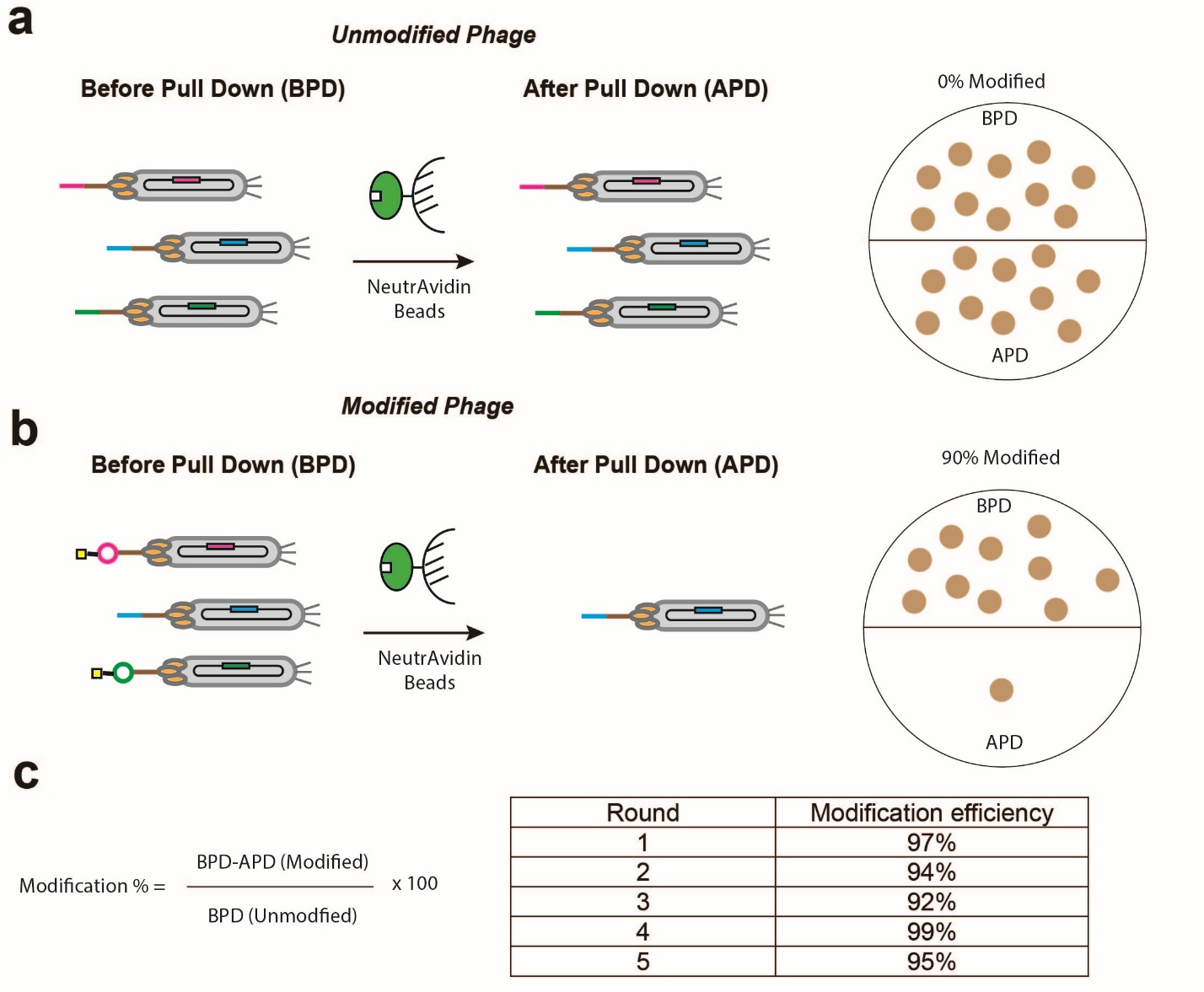

**Supplementary figure 3. Biotin pulse-chase experiments monitor the chemical modification efficiency of displayed peptides**. **a**. The unmodified phage are not captured by the neutravidin beads and can be visualized with a titer. **b**. Phage that have been modified with the biotin linker are captured by the neutravidin beads and do not appear in the titer. **c.** The modification yield (in %) can be calculated using the titer numbers, with the formula indicated. The table of the recorded modification efficiency for the phage display library after each round of chemical modifications is shown here, in all rounds the chemical modification was >90% efficient.

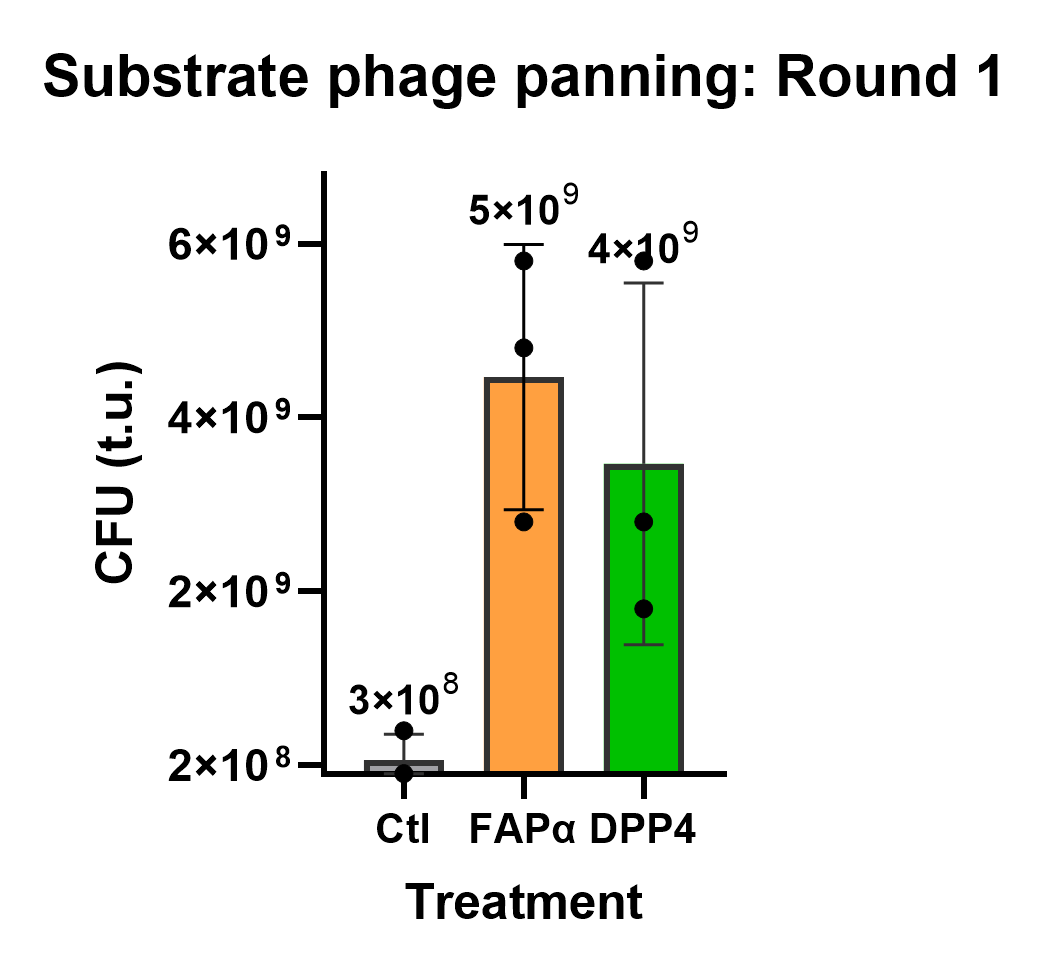

**Supplementary figure 4.** **Library enrichment in the first panning round (R1) monitored using the traditional PFU tittering method**. Treatment with the target proteases FAPα or DPP4 shows a 16- and 13-fold enrichment respectively, compared with the control library (no protease treatment).

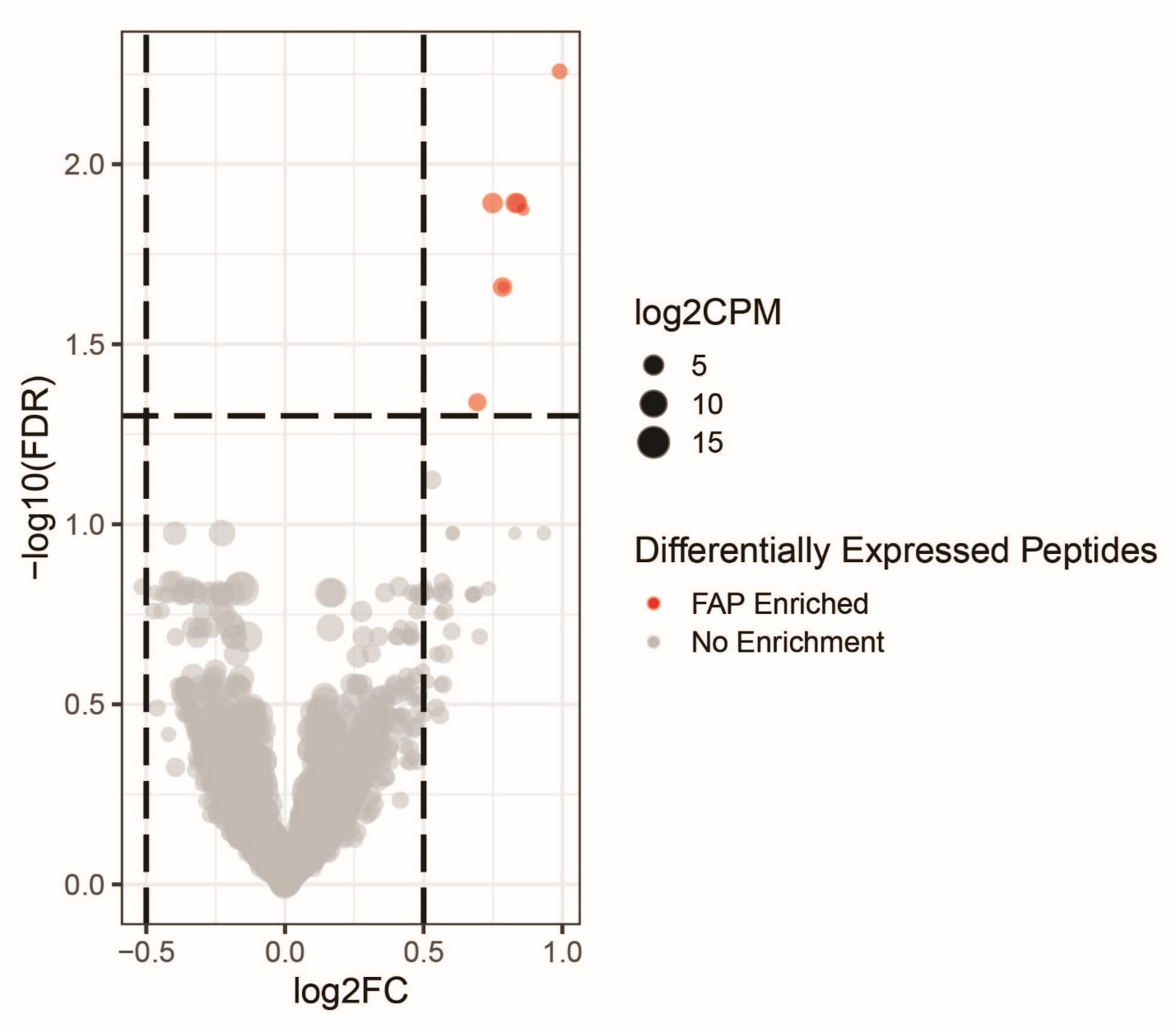

**Supplementary figure 5.** **Enriched peptide sequences after 5 rounds of selection with FAPα**. The volcano plot shows the comparison of the control vs the FAPα treated library, indicating the false discovery rate versus the fold change of unique phage clones. A sequence is considered enriched once it is 1-fold more abundant in one of the libraries (log2FC = 0.5). The circles represent the counts per million (CPM) of each clone: red for FAPα enriched clones and grey for non-enriched clones. No clones were found to be enriched in the control library.

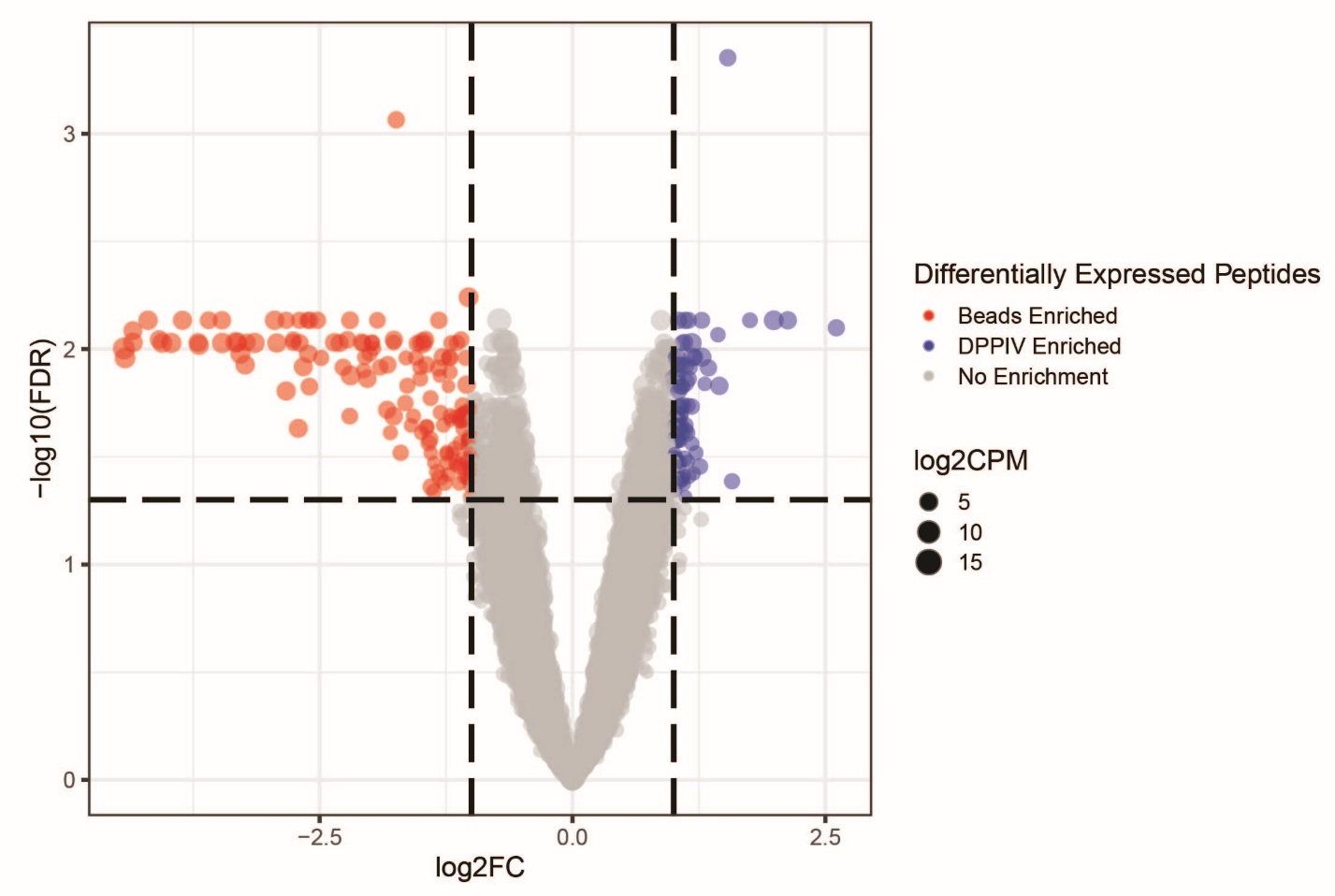

**Supplementary figure 6.** **Enriched peptide sequences found after 5 rounds of selection with DPP4**. The volcano plot shows the comparison of the control vs the DPP4 treated library, indicating the false discovery rate versus the fold change of unique phage clones. A sequence is considered enriched once it is 2-fold more abundant in one of the libraries (log2FC=1). The circles represent the counts per million (CPM) of each clone: blue for DPP4 enriched clones, red for clones enriched in the control library and grey for non-enriched clones.

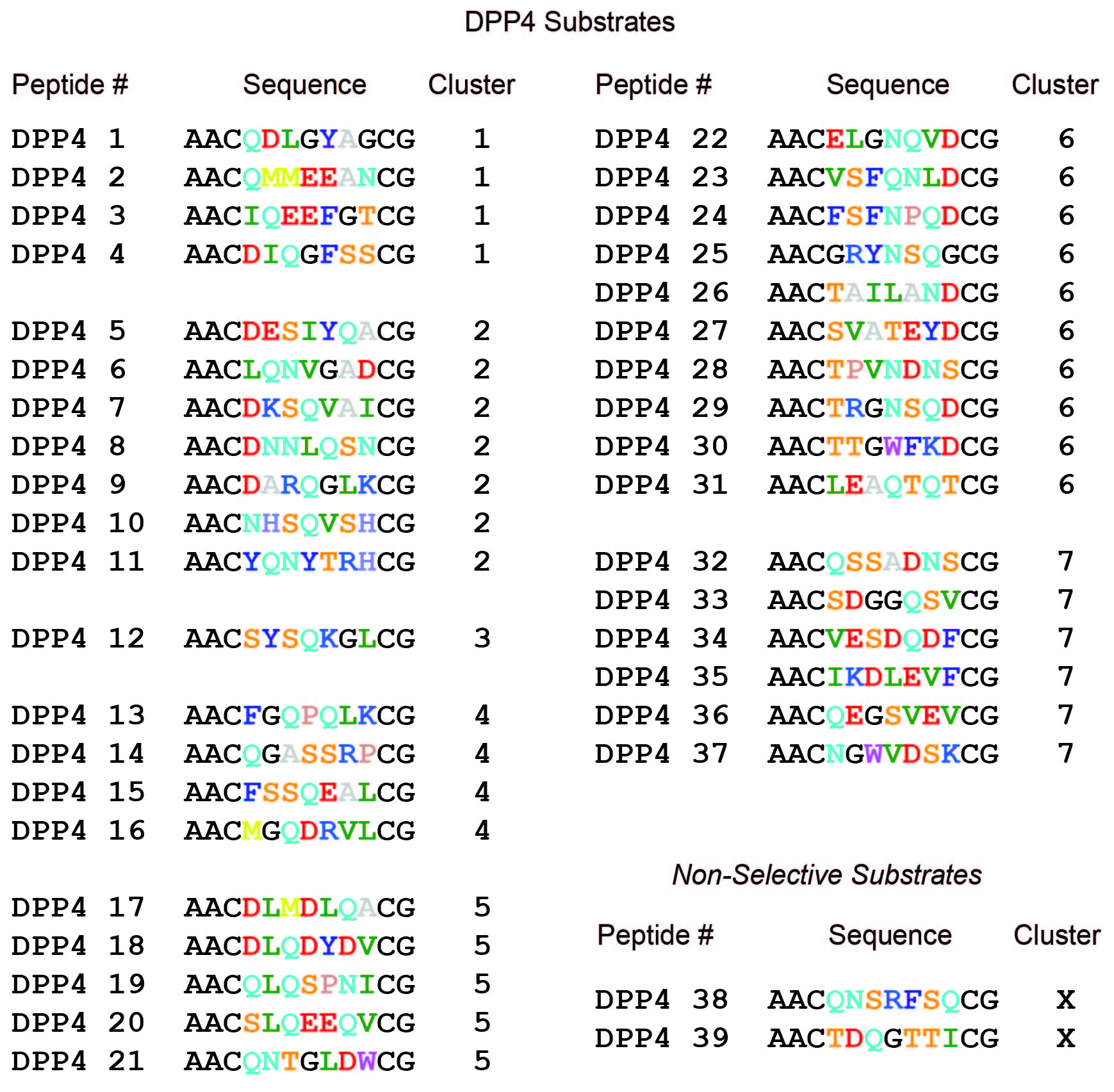

**Supplementary figure 7.** **Linear peptide sequences representing the top hits synthesized for DPP4**. These are representative sequences selected from a larger number of peptide hits, which were classified in 7 different clusters using an unsupervised clustering and alignment algorithm: GibbsCluster.^1^ The number assigned to each peptide represents the enrichment order calculated within each cluster. Constant amino acids at the N and C-terminal ends of the peptides are indicated in black. Variable amino acids are colored using the RasMol coloring scheme, except glycine (G) that is colored black for visual purposes.

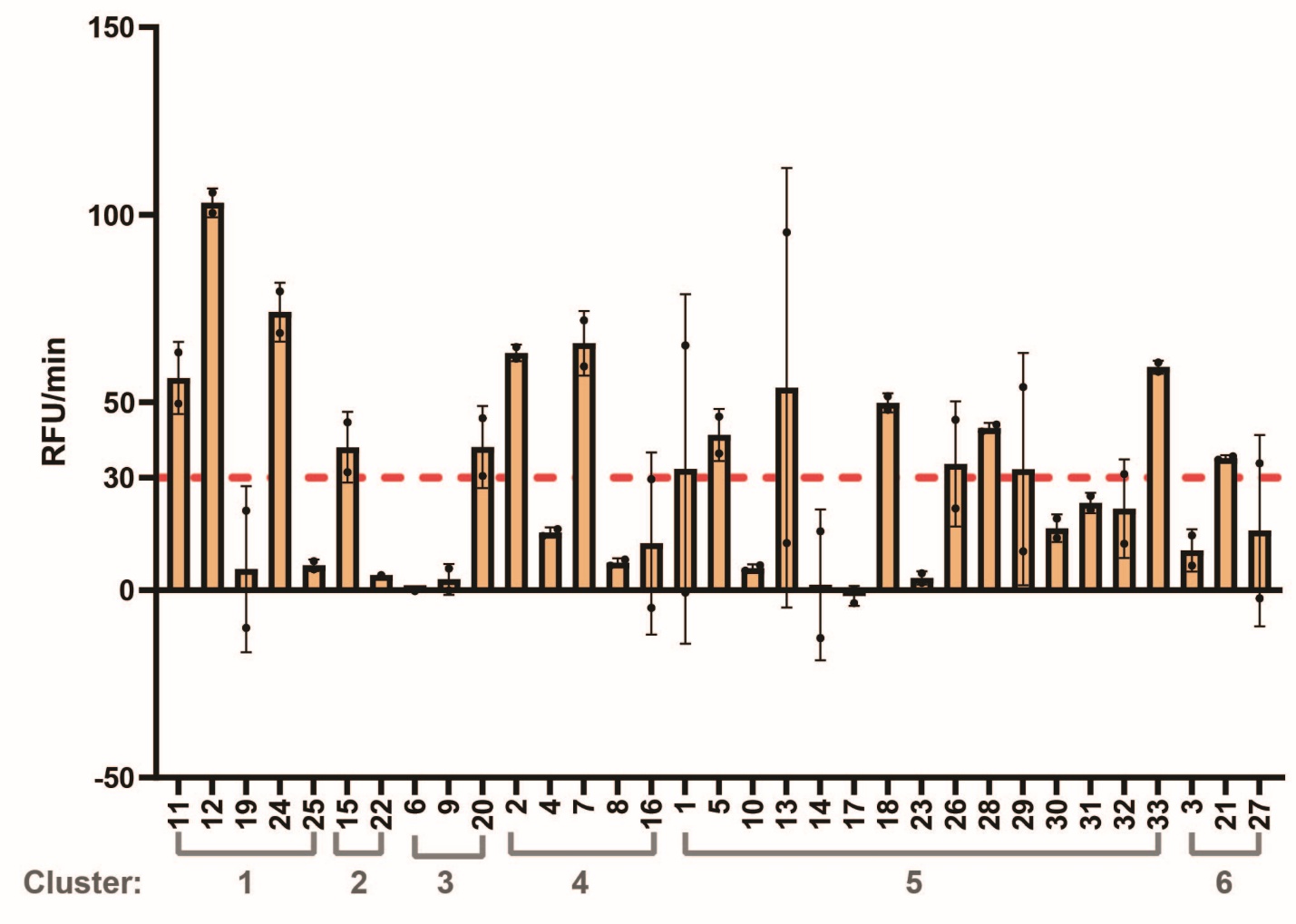

**Supplementary figure 8. Crude testing of FAPα synthesized peptide hits, sorted by sequence similarity clusters**. The graph shows the fluorescence in RFU/min produced by each macrocyclic substrate at 200 μM upon treatment with 10 nM FAPα for 1.5 hours at 37°C. The red dotted line shows the cut-off used to select peptide hits. The experiment was run in triplicates and the error bars represent the standard deviation.

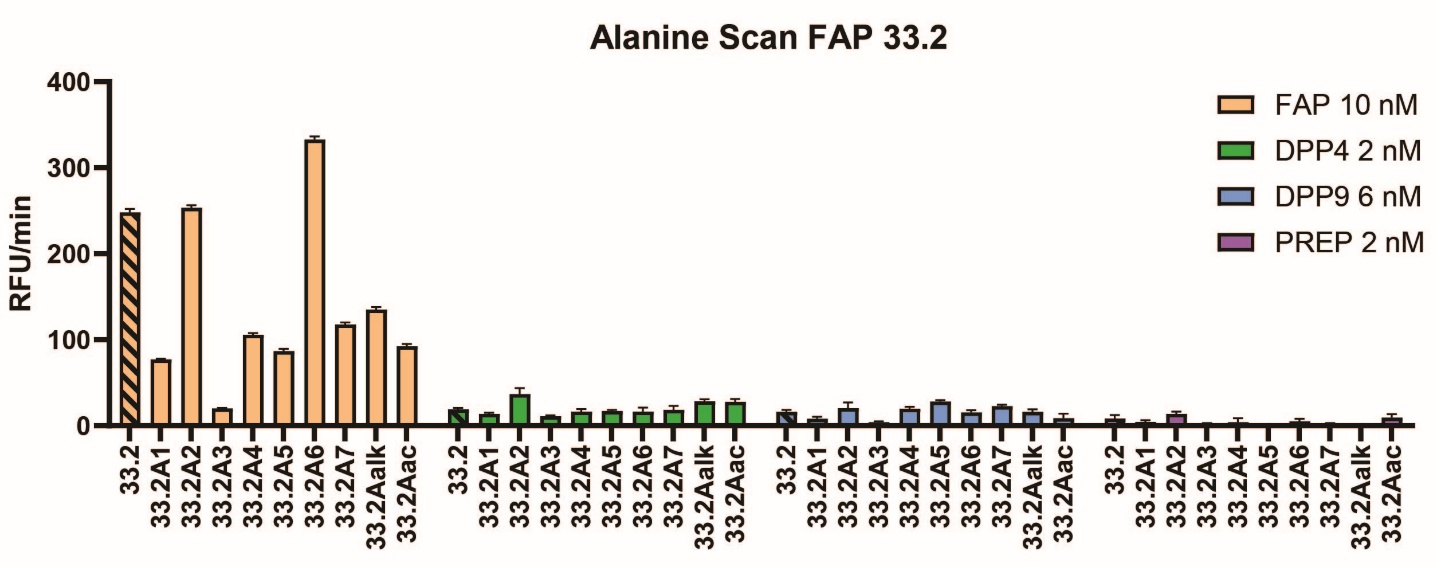

**Supplementary figure 9. Alanine scan of peptide FAP-33.2 shows the potential modification points in the peptide sequence.** The graph shows the fluorescence in in RFU/min for FAP-33.2 analogs upon treatment with 10 nM FAPα, 2 nM DPP4, 6 nM DPP9 and 2 nM PREP. All substrates were tested at 200 μM for 1.5 hours at 37°C in triplicates. Error bars represent the standard deviation.

**FAP-33.2 analogs featuring unnatural amino acids**

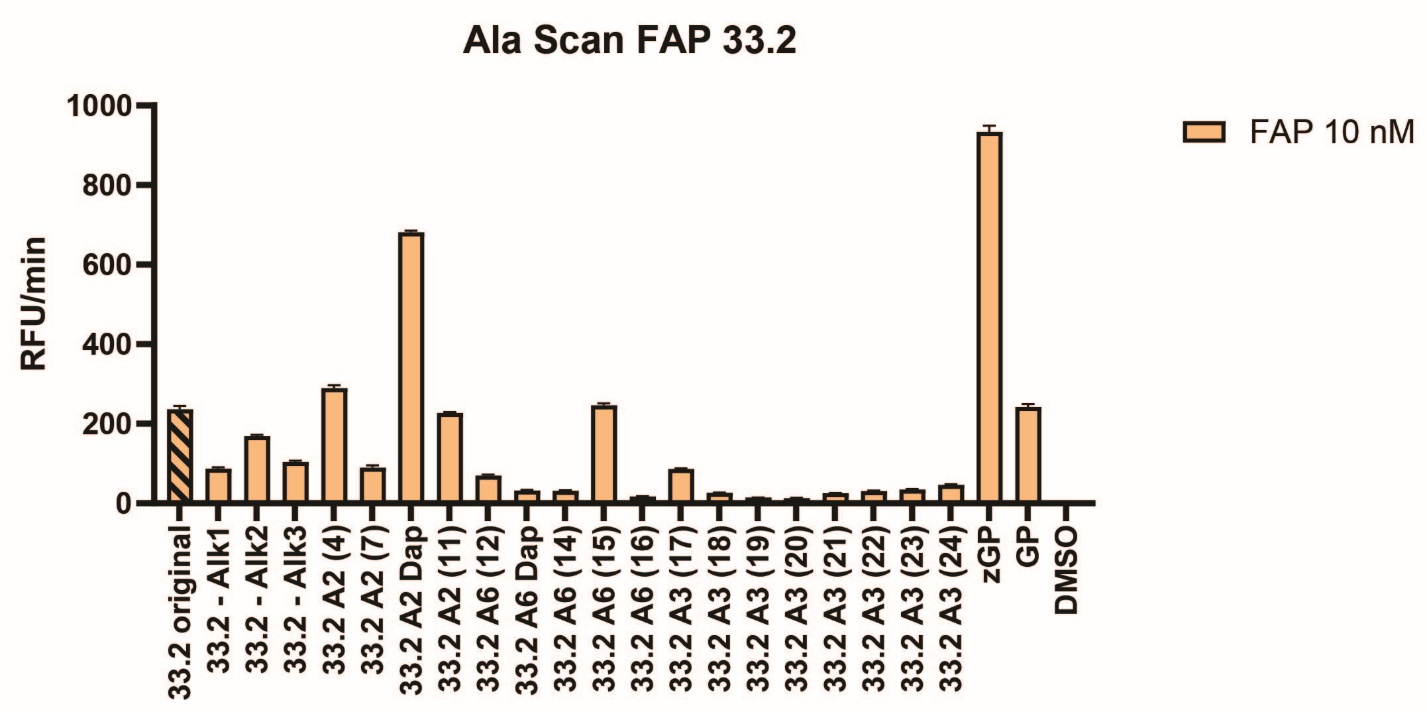

**Unnatural Amino Acid Number (X) Side Chain Structure**

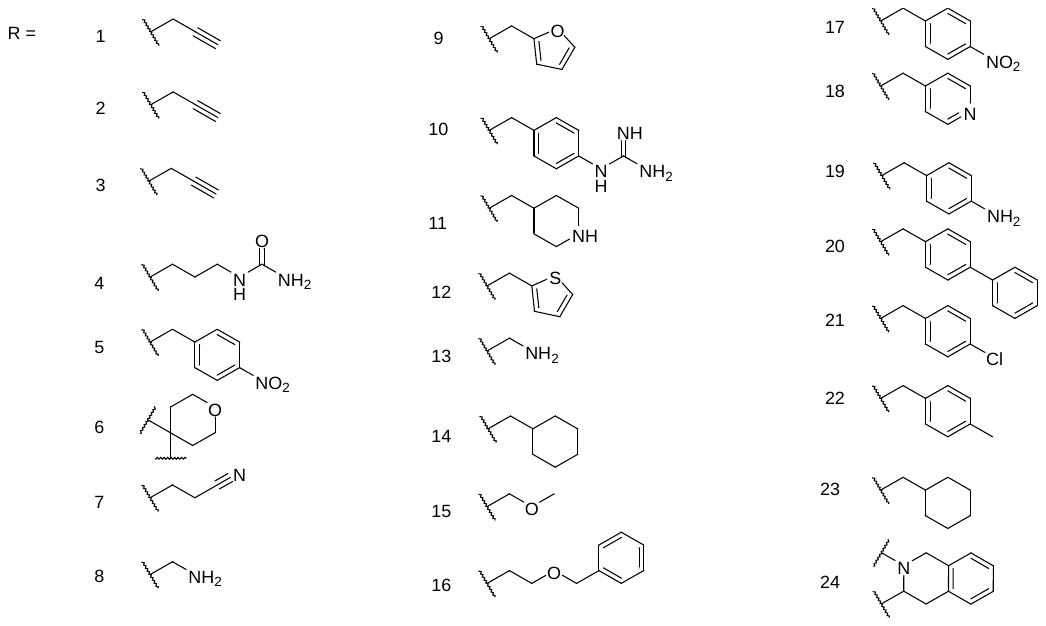

**Supplementary figure 10. Activity of FAP-33.2 analogs measured by fluorescence.** The graph shows the tested peptides. The modified position in each of the analogs is indicated as “A” followed by a number, which indicates the position in the peptide sequence. The unnatural amino acid (UAA) used is identified in between brackets by a number or a 3 letter code. The correlation between the numbers and the UAA side chain is shown in the bottom panel. The fluorescence is depicted as RFU/min. Each substrate was tested at a concentration of 200 μM and incubated with 10 nM FAPα for 1.5 hours at 37°C. The experiments were performed in triplicates. Error bars represent the standard deviation.

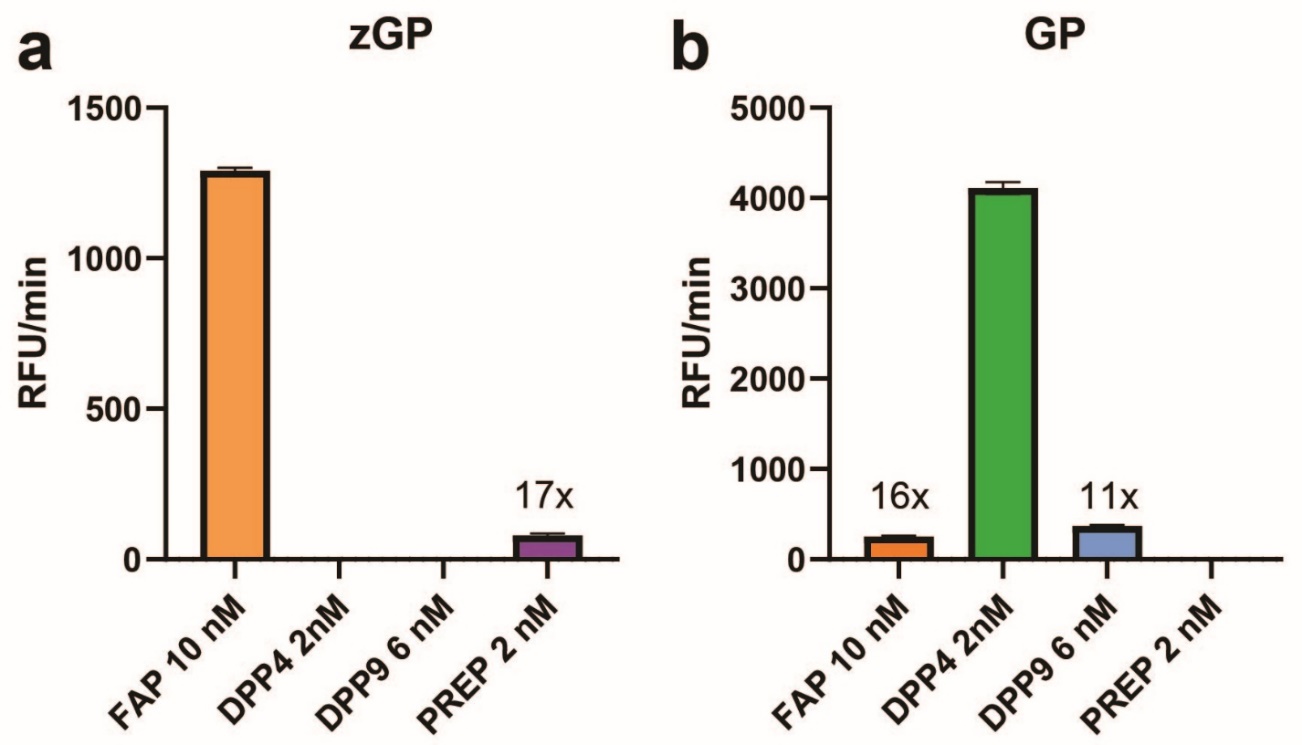

**Supplementary figure 11. Selectivity of the DPP4 and FAPα commercial substrates.** The activity of each substrate upon incubation with selected S9 proteases is indicated as RFU/min. **a**. Substrate zGP is hydrolyzed by FAPα and PREP, being 17-fold more selective for FAPα. **b**. Substrate GP is hydrolyzed by FAPα, DPP9 and DPP4, being 16- and 11-fold more selective for DPP4. The commercial substrates were tested at 200 μM for 1 hour at 37°C with 10 nM FAPα, 2 nM DPP4, 6 nM DPP9 and 6 nM PREP. The error bars represent the standard deviation.

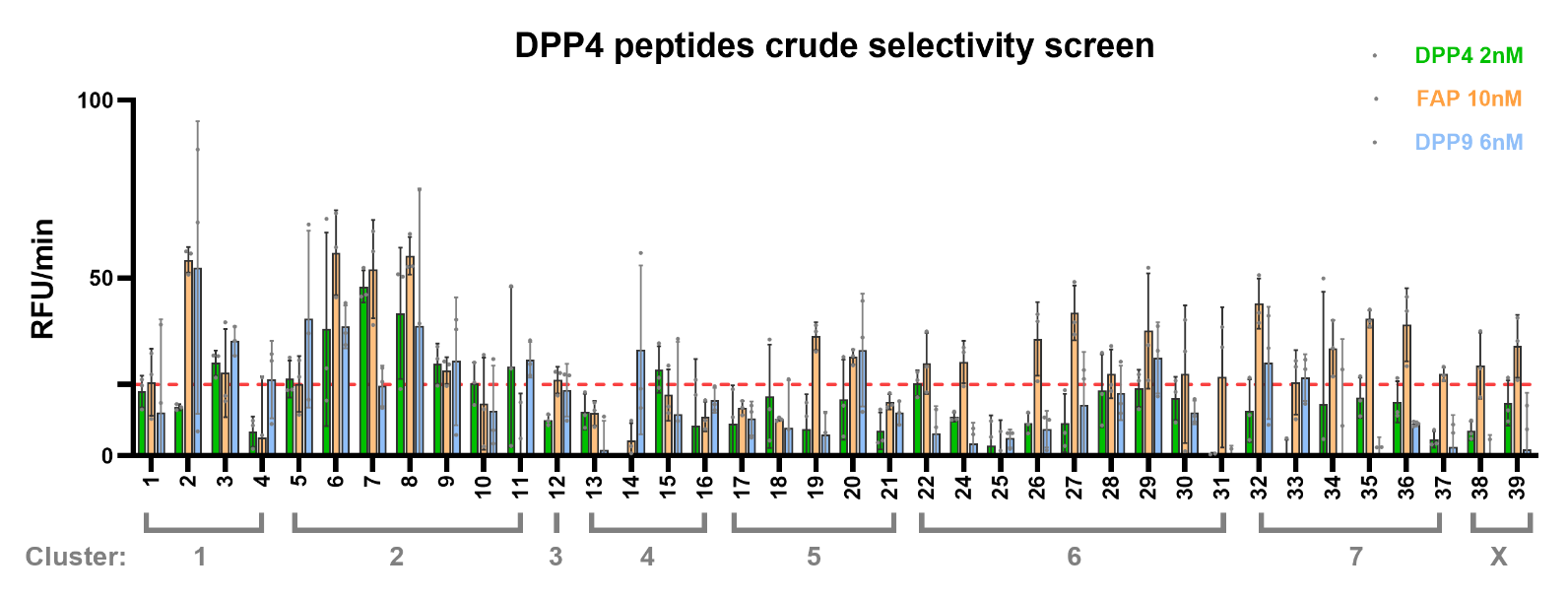

**Supplementary figure 12. Selectivity assessment of crude DPP4 peptide hits, sorted by sequence similarity clusters.** The graph shows the fluorescence in RFU/min produced by each macrocyclic substrate at 200 μM upon treatment with either 2 nM DPP4, 10 nM FAPα or 6 nM DPP9 for 1.5 hours at 37°C. The red dotted line shows the DPP4 activity cut-off used to select peptide hits. The experiment was run in triplicates and the error bars represent the standard deviation.

**Evaluation of FAPα and DPP4 commercial substrates in cells**

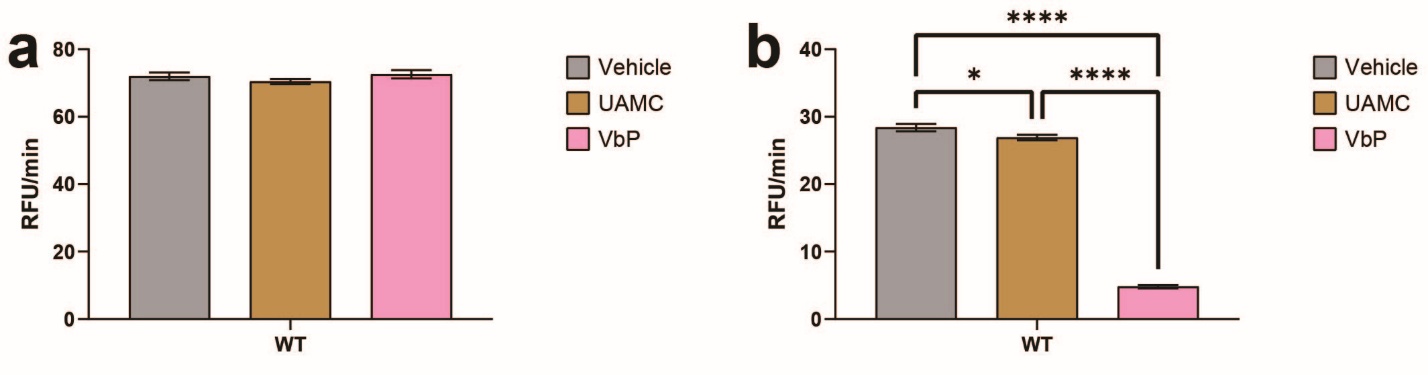

**Supplementary figure 13. *In cellulo* assays of FAPα and DPP4 commercial substrates tested in wild type (wt) HEK293 cells**. Both fluorogenic substrates were used at 200 µM: **a.** zGP-AMC, **b**. GP-AMC. Prior to the addition of the indicated substrate, the cells were treated with either: vehicle DMSO (in grey) or 1 µM covalent inhibitor, UAMC-1100 for FAPα inhibition (in ochre) or VbP for DPP4 inhibition (in pink). All experiments were performed at 37ºC in triplicates, the mean slope in RFUs per minute is plotted where the error bars show one standard deviation. A one-way ANOVA was performed with Tukey’s test. Non-significant values are not shown. P values represent: *<.05, **<.005, **<.0005,****<.0001.

***In cellulo* evaluation of FAP-33.2 macrocyclic substrate**

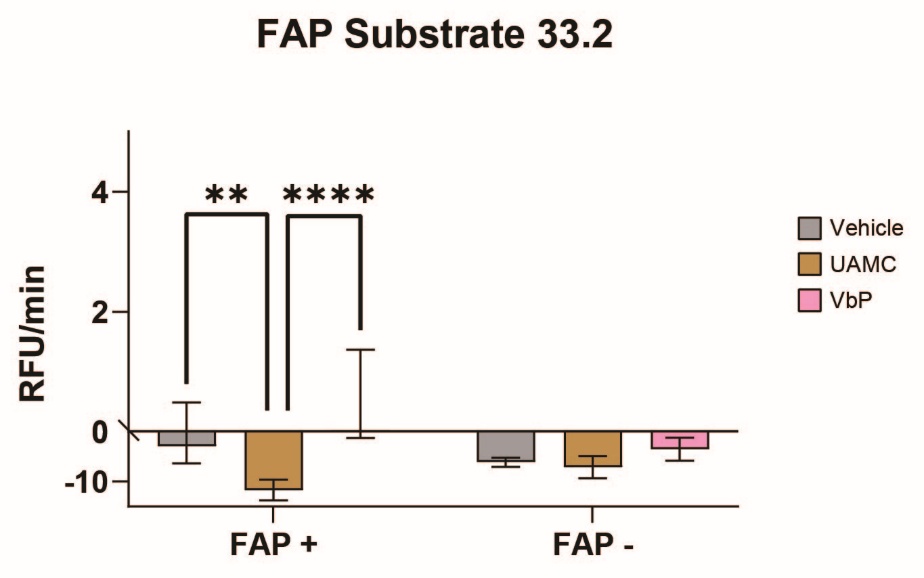

**Supplementary figure 14. FAP-33.2 macrocyclic substrate tested in HEK293 cells with inducible FAPα expression.** HEK293 cells expressing FAPα (FAP+) or with doxycycline-supressed FAPα expression (FAP-) were used to assess *in vitro* substrate cleavage. Prior to the addition of the indicated substrate, the cells were treated with either: vehicle DMSO (in grey) or 1 µM covalent inhibitor, UAMC-1100 for FAPα inhibition (in ochre) or VbP for DPP4 inhibition (in pink). All experiments were performed at 37ºC in triplicates, the mean slope in RFUs per minute is plotted where the error bars show one standard deviation. A two-way ANOVA was performed with Tukey’s. Non-significant values are not shown. P values represent: **<.005, ***<.0005,****<.0001.

**Unnatural Amino Acid (X) General Name and CAS**

| Unnatural Amino Acid Number (X) | General Name | CAS # |
| --- | --- | --- |
| 1 | Fmoc-Pra-OH | 198561-07-8 |
| 2 | Fmoc-Pra-OH | 198561-07-8 |
| 3 | Fmoc-Pra-OH | 198561-07-8 |
| 4 | Fmoc-Cit-OH | 133174-15-9 |
| 5 | Fmoc-Phe(4-NO_2_)-OH | 95753-55-2 |
| 6 | Fmoc-4-amino-tetrahydropyran-4-OH | 285996-72-7 |
| 7 | Fmoc-Cba-OH | 913253-24-4 |
| 8 | Fmoc-Dap(Boc)-OH | 162558-25-0 |
| 9 | Fmoc-(2-Furyl)Ala-OH | 159611-02-6 |
| 10 | Fmoc-Phe(4-guanidino-Boc2)-OH | 187283-25-6 |
| 11 | Fmoc-Ala-4Pip(Boc)-OH | 204058-25-3 |
| 12 | Fmoc-Ala(2-thienyl)-OH | 130309-35-2 |
| 13 | Fmoc-Dap(Boc)-OH | 162558-25-0 |
| 14 | Fmoc-Cha-OH | 135673-97-1 |
| 15 | Fmoc-Ser(Me)-OH | 159610-93-2 |
| 16 | Fmoc-Hse(Bzl)-OH | 1185841-92-2 |
| 17 | Fmoc-Phe(4-NO_2_)-OH | 95753-55-2 |
| 18 | Fmoc-4-Pal-OH | 169555-95-7 |
| 19 | Fmoc-Phe(4-NHBoc)-OH | 174132-31-1 |
| 20 | Fmoc-Bip-OH | 199110-64-0 |
| 21 | Fmoc-Phe(4-Cl)-OH | 175453-08-4 |
| 22 | Fmoc-Phe(4-Me)-OH | 199006-54-7 |
| 23 | Fmoc-Cha-OH | 135673-97-1 |
| 24 | Fmoc-Tic-OH | 136030-33-6 |

**Supplementary table 1**. **Unnatural amino acids used to prepare FAP-33.2 analogs.** The amino acids were purchased from a variety of suppliers, indicated in the chemistry methods section. All amino acids have L stereochemistry except for Fmoc-4-amino-tetrahydropyran-4-OH where the convention does not apply.

**Methods**

**Biology Methods**

**Fluorogenic Substrate Cleavage Assay**

The following proteases were used for the activity assay: Recombinant Human FAPα Protein, CF (R&D Systems, Catalog No.: 3715-SE-010), Recombinant Human Prolyl Oligopeptidase/PREP Protein, CF (R&D Systems, Catalog No.: 4308-SE-010). Recombinant Human DPP9 Protein, CF (R&D Systems, Catalog No.: 5419-SE-010). Recombinant Human DPP4 Protein (Sino Biological, Catalog No.: 10688-HNCH-10). The buffers used for the activity assay were: FAPα - 50 mM Tris, 1 M NaCl, 1 mg/mL BSA, pH 7.5, PREP- 25 mM Tris, 250 mM NaCl, 2.5 mM DTT, pH 7.5 (DTT was freshly added immediately before use), DPP9- 25 mM Tris, pH 7.5, DPP4- 25 mM Tris, pH 7.5. The following commercial substrates were used: zGP-AMC (Bachem,Catalog No.: 4002518.0050) and GP-AMC (ChemScene, Catalog No.: CS-0142320). All assays were conducted in Greiner Bio-One black microplates of 384 wells and polystyrene F-bottom (Catalog No.: 784076). All compounds were dissolved in DMSO at a concentration of 10 mM and then diluted into the respective buffer to the desired final concentration. The protease was added to the specified concentration before the fluorescence measurements. For fluorescence measurements, 2 μL of substrate was further diluted into 6 μL of buffer, followed by addition of 2 μL of enzyme in buffer. For no enzyme controls 2 μL of substrate was diluted into 8 μL of buffer. The fluorescence was measured using a Cytation 3 plate reader (BioTek). All experiments have at least three technical replicates. All assays were conducted at 37°C for a minimum of 90 min, unless otherwise noted. Fluorescence measurements were performed at ex/em, 355/460 nm for ACC substrates. The substrates velocity was quantified by calculating the slope of the linear phase of the reaction as relative fluorescence units (RFU) per min.

**General cell culture**

HEK tet-off FAPα stable clonal cell line, received from the group of Jan Konvalinka, IOCB Prague, was cultured in IMDM medium with L-glutamine (Sigma-Aldrich, 51471C-1000ML), and supplemented with 10% FAS and antibiotics – Geneticin G418 400 μg/ml (Roche, G418-RO) and Puromycin 0.5 μg/ml (Corning, MT61385RA) or Doxycycline 100 ng/ml (Sigma-Aldrich, D9891-1G). In general, cells were passaged at least two times after thawing before their use in cellular assays. FAPα expression was suppressed with Doxycycline for at least a week prior to an experiment.

**Live-cell substrate assay**

Cells were grown to near confluency and harvested via trypsinization. After dissociation, extra media with serum was added to prevent further trypsinization. To collect the cells, those were centrifuged for 3 min at 500G and media removed. Cells were washed a total of three times with PBS and resuspended in PBS to 100 cells per μL. To perform the activity assay 14 μL of suspended cells in PBS (100 cell/μL) were plated in a black 384-well low volume “V” bottom plate (total of 1400 cells per well) and pre-treated with 2 μL of either vehicle or inhibitors (1 μM final concentration) for 30 min at 37 °C. DMSO was used as the vehicle. The inhibitors used were UAMC-1100 and Talabostat (VbP, valine boroproline) which semi-selectively inhibited FAPα and DPP4 activity, respectively, in addition to other homologous proteases. After this time, 4 μL of the desired substrate was added to a final concentration of 200 μM and the fluorescence was recorded immediately. Fluorescence was measured using a Cytation 3 plate reader (BioTek). All experiments have at least three technical replicates. All assays were conducted at 37°C for 90 min, unless otherwise noted. Fluorescence measurements were performed at ex/em, 355/460 nm for ACC substrates. The substrates velocity was quantified by calculating the slope of the linear phase of the reaction as relative fluorescence units (RFU) per min.

**Phage Display Methods**

The library and panning protocol was adapted from Ekanayake et. al.^2^

**Library composition**

The phage encoded library has the terminal sequence AACXXXXXXXCG. The complexity of the library is 1x10^9^. In general a minimum of 5x10^10^ PFU were used for panning to have 50 copies of each sequence represented at the beginning of panning.

**General reagents for phage display**

*YT media*

15.5 g of 2X YT broth (1.6% tryptone, 1.0% yeast extract, 0.5% sodium chloride) 500mL of MilliQ water in a 2L flask, autoclaved.

*YT media with 30 μg/mL final concentration of chloramphenicol (CA)*

15.5 g of 2X YT broth (1.6% tryptone, 1.0% yeast extract, 0.5% sodium chloride) 500 mL of MilliQ water in a 2L flask, autoclaved. Let it cool to room temperature. Add 500 μL of 30 mg/mL chloramphenicol in ethanol.

Agar plates of YT media with chloramphenicol (*30 μg/mL CA*)

Weight out 31 g of 2X YT broth (1.6% tryptone, 1.0% yeast extract, 0.5% sodium chloride), 15 g of agar and add 1000 mL of MilliQ water. Autoclave. Let it cool down to approximately 55°C and add 1000 μL of 30 mg/mL CA in ethanol. Mix and pour into plates under sterile conditions.

*20% PEG 8000 Solution*

Bring 200 g of PEG 8000 and 145 g of NaCl to 1L using MiliQ water and autoclave in a 2L container. (Using PEG between 5000 and 10,000 will also afford phage precipitation)

*Buffers used*:

Reaction buffer 1: 1M NaHCO_3_, 200 mM of TCEP in MilliQ H_2_O at pH 8.5 (TCEP freshly added before use)

Reaction buffer 2: 1M Ammonium acetate, 0.5 M EDTA in MilliQ H_2_O at pH 4

Wash buffer 1: 0.1% Tween in PBS.

Wash buffer 2: 1.0% Tween in PBS.

Blocking Solution: Add 1% solid Bovine Serum Albumin (BSA) to 0.1% Tween in PBS.

FAPα reaction buffer: 50 mM Tris, 1 M NaCl, pH 7.5

PREP reaction buffer: 25 mM Tris, 250 mM NaCl, 2.5 mM DTT, pH 7.5 (DTT was freshly added immediately before use)

DPP9 reaction buffer: 25 mM Tris, pH 7.5

DPP4 reaction buffer: 50 mM Tris, 1 M NaCl, pH 7.5

**Substrate phage panning protocol**

**Phage panning optimization.** To determine optimal experimental conditions, we tested the following parameters: a. Library immobilization: We compared streptavidin and neutravidin magnetic beads to identify the highest binding capacity, optimal incubation time, and bead volume for immobilizing a library of 10⁹ phage-displayed peptides. PFU colony counting revealed that the best binding occurred with 150 µL neutravidin beads and a 1-hour incubation (**Figure S2**). b. Proteolytic selection. To optimize FAPα-library selection, we evaluated two temperatures (22ºC and 37ºC) and five incubation times (1, 2, 3, 4, or 16 hours). PFU colony counting indicated that optimal conditions were 1 hour at 37ºC. For practical reasons, we adopted 2 hours at 37ºC, as it showed no significant difference from 1 hour at 37ºC (**Figure S2**). c. Washing conditions. We optimized washing buffer composition, temperature at which the washes are performed, and the number of washes using PFU colony counting. The best conditions were seven washes with 1% BSA and 1% Tween in PBS (5× at room temperature, 2× at 37ºC) followed by two washes with 0.1% Tween in PBS (1× at 37ºC, 1× at room temperature). We tested various conditions, including Tween concentrations (0.1%–5%), different temperatures (RT or 37ºC), and up to 12 total washes.

**Phage-encoded library propagation.** Phage harboring the cysteine-rich peptide library (CX_7_C) were used for selecting substrate peptides for FAPα and DPP4 proteases. The randomized DNA sequences encoding the peptide library were inserted into the phage DNA between the pelB signal peptide and the disulfide-free pIII protein of the fdg3p0ss phage vector^1^. To overcome the low infectivity of the disulfide-free phage strain used in our screens, we used a large volume of 2YT rich medium (2 x 500mL) for the production of phage. The phage peptide libraries in TG1 E.*coli* bacteria (Lucigen, 60502-1) were thawed from stock and used for inoculating 2YT medium containing 30 μg/ml^–1^ chloramphenicol, which were incubated at 30 °C with shaking at 250 rpm for 16 h. After this time, the medium rich with secreted phage was separated from the host bacteria by centrifuging the solution at 12,785G for 30 min at 4°C, and the supernatant containing the phage was decanted into a new container with ice cold 20% PEG 8000, 2.5 M NaCl solution and thoroughly mixed (4 bottles: 250 mL phage supernatant x 65 mL PEG solution). Following a 1-h incubation on ice, mixing every 15 min, the precipitated phage were spun at 12,785G for 45 min at 4°C. The container was carefully removed from the centrifuge and the phage pellet identified. The supernatant was decanted into a waste flask in one motion, without disrupting the phage pellet (facing opposite from the supernatant decantation route). The containers were turned upside down, and the excess remaining liquid was removed using tweezers and an absorbent material to wipe the liquid without touching the phage pellet. Per container, 1 mL of PBS (cell culture grade) was used to dissolve the phage pellets. This was transferred to a new 15 mL conical tube. The same areas were rewashed with another 1 mL PBS and added to the conical tube (in total 2mL per container). The sample was vortexed. The 15 mL conical tube was allowed to sit on ice for 10 min. The solution was centrifuged at 6,000G for 15 min at 4°C to pellet the leftover cell debris. The supernatant was carefully transferred using a pipet into a new 15 mL conical tube, and 2 mL of cold 20% PEG 8000 2.5 M NaCl solution was added. The whole was mixed, vortexed, and left it on ice for 30 min to precipitate the phage. The precipitated phage was centrifuged at 6,000G for 60 min at 4°C. The supernatant was decanted into the waste. Using tweezers and an absorbent material, the excess drops of liquid were wiped off without touching the phage pellet. The pellet was dissolved in 1 mL of PBS and transferred to a 1.5 mL Eppendorf. The sample was vortexed and allowed to sit on ice for 15 min, after which it was centrifuged for 20 min at 4°C and max speed to pellet the remaining cell debris. The phage-containing supernatant (1 mL) was transferred to a new tube. This afforded the unmodified native (NL) phage library. Of this sample, 50 µL were kept for tittering (NL) and the remaining 950 µL were used in the next step.

**Library modification.** The NL isolated in the previous step (950 µL) was treated with 150 µL NaHCO_3_ 1M (final concentration 100 mM) pH=8.0, 15 µL 200 mM TCEP (final concentration 20 mM) and brought to 1.5 mL total volume by adding miliQ autoclaved H_2_O. The mixture was left rotating at room temperature for 30-60 min to afford the reduction of the cysteines. After this time, 15 µL of 200 mM DPD in DMF were added. After vortexing, the whole was left rotating for exactly 30 min after which the solution turned light yellow. The excess DPD was removed by adding the phage reaction mixture to a Zeba Protein Dissolving column (7k, MWCO), pre-washed 5 times with 1 mL of reaction buffer, and centrifuged dry. The Zeba column containing the phage mixture was placed in a collection tube and centrifuged for 4 min at 1000G. The filtrated ~1500uL of phage solution was placed in a 5 mL Eppendorf and 50 µL sample were taken in order to perform chemical modification pulse-chase experiments (*DPD_C7C*). To the phage solution were added 490 μL of Reaction Buffer 2 and 40 μL of 20 mM bio-orthogonal linker in acetonitrile (40 μM reaction concentration) and the whole was vortexed and left mixing for 2 hours at room temperature. After this time, 600 μL of 20% PEG 8000 solution were added to precipitate the phage. The whole was mixed by inversion and left sitting on ice for 30 min, after which it was centrifuged at 6,000G at 4°C for 30 min to collect the phage. The supernatant was removed and the excess drops of liquid were dried, using tweezers and an absorbent wipe, without touching the phage pellet. The phage pellet was resuspended in 1mL of ice cold PBS, vortexed briefly and left sitting on ice for 15 min. Next, the solution was centrifuged at 4°C for 10 min at max speed. The phage containing supernatant was transferred into a new tube and kept on ice or stored at 4ºC overnight. This is the modified phage library, “Modified Library”. Of this supernatant, 50 μL were kept for the titer/ biotin *pulse-chase* experiment.

**Phage panning. Substrate selection.** In order to immobilize the modified phage library, 150 μL of neutravidin magnetic beads (New England Biolabs) per condition were placed into an Eppendorf tube. In order to wash the beads, the tubes were placed on a magnetic stand and the solution was allowed to clear. All the liquid was removed from the tube. Three washes were performed. To perform a wash, 600 μL of the 0.1% Tween Blocking Solution were added and the beads were broken up with a pipette. The beads were placed in the bead magnetic holder. The liquid was removed. To block the beads, 600 μL of 1% BSA 0.1%Tween Blocking Solution were added and the whole was left rocking for 1h at room temperature. After this, 3 more washes were performed with 600 μL of PBS. The modified library was split in 3, adding 333 μL of modified phage to the beads and left rotating for 1h at room temperature to allow on-bead phage library immobilization. After this time, the supernatant was removed and 7 washes with 1% BSA and 1% Tween in PBS (5x at room temperature and 2x at 37ºC) and 2 washes with 0.1% Tween in PBS (1x at 37ºC and 1x at room temperature) were performed. For the library selection, 100 μL of activity buffer (50 mM Tris, 1 M NaCl, pH=7.5) containing the desired protease or no-protease control were added to the phage-libraries immobilized on beads and mixed by thoroughly inverting. Specifically, for each protease and panning round the following concentrations were used: FAPα: 90 nM (R1), 30 nM (R2), 30 nM (R3), 30 nM (R4), 10 nM (R5) and DPP4: 30 nM (R1), 10 nM (R2), 10 nM (R3), 5 nM (R4), 5 nM (R5). The whole was incubated for 2 h at 37ºC, except for R5 when the incubation time was reduced to 90 min. After the indicated time, the tubes were placed on a magnetic stand. The solution was allowed to clear and the supernatant, containing the selected phage displaying fluorogenic substrates, was removed and placed in a separate Eppendorf which was kept on ice. For R5 the selection was performed in triplicates (3x target protease, 3x control library).

Before performing R5 of selection a negative-selection step was performed, for this a mixture of S9 homolog proteases was incubated with the immobilized library, prior to the treatment with the target protease. Specifically, the library was incubated for 30 min at 37ºC with 100 μL of activity buffer containing a mix of 10 nM DPP4, 10 nM DPP9 and 10 nM PREP (for the FAPα panning) and 10 nM FAPα, 10 nM DPP9 and 10 nM PREP (for the DPP4 panning). After the indicated time, the tubes were placed on a magnetic stand, the solution was allowed to clear and the supernatant was removed and placed in a separate Eppendorf which was kept at 4ºC. The beads containing the phage library were washed 5 times with PBS prior incubation with the target protease to perform R5 of selection.

**Stock generation from the selected phage after R5 selection.** To generate glycerol stocks of TG1 cells infected with the protease selected phage of R5, the following steps were taken: An overnight culture of naïve TG1 cells in YT media without antibiotic was prepared by adding a pipette tip that had been used to scrape into a frozen TG1 glycerol stock. After overnight shaking at 37°C at 200 rpm, 5 mL of this overnight culture were used to inoculate fresh 500 mL of YT media under sterile conditions. These cultures were then incubated at 37°C and shaking at 200 rpm for 90 min or until the OD_600_=0.4. After this time, the TG1 culture was split in 45 mL aliquots placed in sterile 50 mL falcon tubes. Each was then infected with 90 µL of one of panning selection outputs, each containing a specific phage library (control library or protease enriched). These cultures were incubated at 37ºC without shaking for 90 min. The 50 mL falcon tubes were taken and centrifuged at 2000 G for 15 min at room temperature. The supernatant was decanted, leaving about 400 μL of media which was used to resuspend the pellet. This suspension was split in 200 μL aliquots and plated into two 15-cm 2YT agar plates with chloramphenicol (30 μg/mL of CA). Under sterile conditions, the bacteria were plated on the plate using a sterile Drigalski spatula until all the media was absorbed. The plates were incubated upside down overnight at 37°C. On the next day, under sterile conditions, 5 mL of sterile YT media was added to one petri dish with a lawn of bacteria. The bacteria were removed from the plate into the YT media using a sterile Drigalski spatula. Once all bacteria were suspended, the 5 mL of media with bacteria was transferred onto the second petri dish with bacteria infected with the same phage library and the same procedure was repeated. Once all bacteria were suspended, the 5 mL of media with bacteria was transferred to a 15 mL falcon tube and 5 mL of autoclaved sterile 50% glycerol/50% water solution was added and mixed. This mixture was aliquoted into 5 cryogenic vials (2 mL in each) and frozen in liquid nitrogen immediately. Each cryogenic vial contains TG1 cells infected with the protease-selective substrate library.

**Library DNA Preparation and NGS.** To sequence the phage-encoded libraries, 200 µL of the glycerol stocks previously prepared (containing the selected libraries after R5 of panning), were used for phage DNA extraction. The DNA was isolated and purified using a Zyppy Plasmid Miniprep Kit [Zymo research], following the kit instructions with the following modifications: DNA elution was performed with 20 µL DNAse free water for 5 min. Of the extracted phage plasmid DNA, 200 ng were used for each sample for PCR amplification, using the following reaction conditions:

400nM Forward Primer Mix (20 nM of each of the individual oligos final conc): 2.5 µL

400nM Reverse Primer Mix (20 nM of each of the individual oligos final conc): 2.5 µL

10mM dNTP : 1 µL

5xHF Buffer : 10 µL

Phage Plasmid : 4 µL of 50 ng/µL (or adjust accordingly)

DMSO: 1 µL

Water : 28.5 µL

The whole was placed on ice while the thermocycler was warming up. After carefully mixing and spinning down, 0.5 µL of Phusion Polymerase were added. Once the Thermocycler reached the right temperature, the samples were added and the PCR was carried out as indicated: [Lid 105ºC Vol 50uL] 95ºC for 5:00 min / 98ºC for 15 sec / 67ºC for 30 sec / 72ºC for 15 sec / GOTO 2, 25 times / 72ºC for 5:00 min / 4C for forever. Next, the samples were evaluated by agarose gel electrophoresis to confirm that the amplified products matched the calculated mobility shift. A second PCR amplification was performed to barcode samples directly from the first PCR procedure, using different combination pairs of forward and reverse primers and according to the following reaction conditions:

2uM Forward adapter: 2.5 µL

2uM Reverse adapter: 2.5 µL

10mM dNTP: 1 µL

5x HF: 10 µL

DNA PCR 1 (crude reaction): 2 µL

DMSO: 1 µL

Water: 30.5 µL

The whole was placed on ice while the thermocycler was warming up and after carefully mixing and spinning down, 0.5 µL of Phusion Polymerase were added. Once the Thermocycler reached the right temperature, the samples were added and the PCR was carried out as indicated: [Lid 105ºC Vol 50uL] 95ºC for 5:00 min / 95ºC for 15 sec / 56ºC for 30 sec / 72ºC for 30 sec / GOTO step 2, 30 times / 72ºC for 5:00 min / 4C for forever. The amplified samples were checked using agarose gel electrophoresis to confirm that the new products had the calculated molecular weight. The gel bands with the expected molecular weight after amplification were cut, using clean razorblades, for DNA extraction and were subsequently purified using the commercial gel DNA recovery kit (Zymo Research, #D4008). The final barcoded purified DNA samples were submitted for NGS. NGS of the was performed by MedGenome Inc. (Foster City, CA), where the samples were pooled and read on the NovaSeq, with 2 million paired reads (paired end 150 reads) for each sample.

*Primers for PCR1
**Step 1 – Forward**

TCGTCGGCAGCGTCAGATGTGTATAAGAGACAGCAACAGTTTCAGCGCCAGAACCGCC

TCGTCGGCAGCGTCAGATGTGTATAAGAGACAGNCAACAGTTTCAGCGCCAGAACCGCC

TCGTCGGCAGCGTCAGATGTGTATAAGAGACAGNNCAACAGTTTCAGCGCCAGAACCGCC

TCGTCGGCAGCGTCAGATGTGTATAAGAGACAGNNNCAACAGTTTCAGCGCCAGAACCGCC

TCGTCGGCAGCGTCAGATGTGTATAAGAGACAGNNNNCAACAGTTTCAGCGCCAGAACCGCC

**Step 2 - Reverse**

GTCTCGTGGGCTCGGAGATGTGTATAAGAGACAGGCTATGCGGCCCAGCCGGCC

GTCTCGTGGGCTCGGAGATGTGTATAAGAGACAGGNCTATGCGGCCCAGCCGGCC

GTCTCGTGGGCTCGGAGATGTGTATAAGAGACAGGNNCTATGCGGCCCAGCCGGCC

GTCTCGTGGGCTCGGAGATGTGTATAAGAGACAGGNNNCTATGCGGCCCAGCCGGCC

GTCTCGTGGGCTCGGAGATGTGTATAAGAGACAGGNNNNCTATGCGGCCCAGCCGGCC

**Phage infectivity measurement (phage tittering)**

The phage was quantified by counting the number of infected TG1 colonies. For this, the desired dilutions were prepared via serial dilutions of phage using PBS. Of each solution, 20 µL were incubated with exponentially growing TG1 bacterial cells (OD_600_ = 0.4) for 90 minutes at 37°C. After the 90-minute infection, 10 µL of each tittered bacterial culture was added in triplicates onto 2YT agar plates (30 µg/mL CA) and incubated overnight at 37°C. Phage titers were calculated by colony counts and dilution factors.

Post-translational chemical modification efficiency / Biotin pulse-chase experiments: In order to assess the efficiency of the chemical modification, pull down experiments were carried out with the aliquots from the DPD-treated phage (*DPD_C7C*) and linker-treated phage (*Modified_C7C*) saved from the previous library preparations. The samples were serially diluted to 10^6^. [990 µL of PBS was added to 10 µL of the *DPD_C7C* and *Modified_C7C* to make 10^2^ dilutions, and then repeated twice to make 10^4^ and 10^4^ dilutions]. Of each 10^6^ dilution, 10 µL were set aside and labelled *DPD_BPD* and *Modified_BPD* (BPD: before pulldown). Next, magnetic neutravidin beads (20 µL bead swirl per condition) were equilibrated and washed (200 µL 3x 0.1% Tween PBS). The beads were then blocked with 200µL of 1%BSA in PBS for 15 min. After this time, one more wash with 200µL PBS was performed. Next, 950 µL of the 10^6^ dilution samples were incubated with the neutravidin beads for 30-minute at room temperature with tumbling. Next, the beads were placed on a magnetic holder, and the supernatant was transferred to new tubes labeled as *DPD_APD* and *Modified_APD* (APD: after pulldown). Per sample, 10 µL of the diluted phage was transferred to 190 µL of exponentially growing TG1 bacterial cells for a 90-minute infection at 37°C. Of each tittered bacterial culture, 200 µL were plated in triplicates onto 2YT agar (30 µg/mL CA) and incubated overnight at 37°C. Phage titers were calculated by colony counts and dilution factors. Chemical modification efficiency was calculated as: yield% = 100 x (Modified library BPD − APD) / Unmodified_BPD.

Post-translational chemical modification toxicity: In order to ensure that the chemical modification was indeed biorthogonal with the phage, traditional PFU counting was carried out before and after each of the chemical reactions. In short, samples containing unmodified phage library (NL= native library), DPD_C7C and Modified_C7C were serially diluted with PBS to afford 10^6^, 10^7^ and 10^8^ dilutions. From those, 20 µL were transferred to 180 µL of exponentially growing TG1 bacterial cells (OD_600_= 0.3-0.4) for a 90-minute infection at 37°C. Of each tittered bacterial culture 10 µL were spotted in triplicates onto 2YT agar plates (30 µg/mL CA) and incubated overnight at 37°C. Chemical modification toxicity to the phage was calculated by PFU counting and dilution factors. Toxicity step 1 (%) = DPD_C7C/NL and Overall toxicity (%) = Modified_C7C/NL.

Phage panning monitoring by PFU counting: For the first panning round (R1) the library selection was monitored using classical PFU counting. For this, 10 µL of phage solution was saved for titer measurement before and after protease incubation, and dilutions 10^6^, 10^7^ and 10^8^ were prepared as previously indicated. TG1 cells in 5 mL YT media were grown overnight at 37°C and 200 rpm. Of this culture, 50 μL were transferred into 5 mL of fresh YT media and grown at 37°C for 90 minutes or until (OD_600_=0.4). Per treatment condition, 20 μL of the diluted phage were added to 180 μL of the bacterial culture in an Eppendorf tube and incubated at 37°C for 90 minutes. After incubation, the cultures were briefly mixed, 10 μL per sample were spotted in triplicates onto 2YT agar plates (30 µg/mL CA). The spots were allowed to dry and the plates were incubated overnight at 37°C. Colonies were counted and multiplied by the dilution factor. The phage recovery was calculated as the percentage of post-panning titer divided by pre-panning titer. For each panning cycle, a no-protein control panning (beads only) was used to calculate the fold change of phage recovery of the sample (protein target) over the control (beads only).

Phage panning monitoring by fluorescence: For rounds R2 to R5, the library selection by the target protease, was monitored using a microplate reader (Cytation 3 from BioTek). To do so, in a 96-well Greiner Bio-One black microplate F-bottom were placed in 10 µL of activity buffer, 10 µL of neutravidin beads slurry either: unused (washed as previously described, “beads autofluorescence control”); prior to protease incubation (no-protein control) and after protease incubation (panned library). The fluorescence measurements were performed at ex/em, 355/460 nm at 37ºC. The plate was vigorously mixed before each reading. All experiments have at least three technical replicates.

**Computational Pipeline**

Paired reads were merged using the fastq_mergepairs command of the vsearch package. Next, adapters were clipped from the 5’ and 3’ end of merged reads using cutadapt. DNA sequences were translated into protein sequences using the translate function of Biostrings.

Using RStudio, data was filtered to exclude peptides with mutated constant amino acid positions

or with cysteines in variable amino acid positions. We employed the DESeq2 package for differential enrichment analysis, following standard procedures (<https://bioconductor.org/packages/release/bioc/vignettes/DESeq2/inst/doc/DESeq2.html>). ^3^ This method employs the prior distribution of log fold changes (LFC) to calculate LFC and

statistical significance for all peptides, even when they are not detected in all samples.

**Chemistry Methods**

**General Materials and Synthetic Methods**

All reactions were performed at room temperature and under atmospheric conditions unless otherwise noted. Light sensitive ACC derived compounds were protected from light whenever possible. All commercial solvents and reagents were used without further purification including N,N’- dimethylformamide (DMF), acetonitrile (ACN), dichloromethane (DCM), methanol (MeOH), and molecular biology grade dimethyl sulfoxide (DMSO). An analytical LC-MS was used to track reaction completion and purity of the final compounds. The LC-MS systems used were: a 1200 HPLC equipped with a Zorbax SB-C18 column (1.8 μm, 2.1 x 50 mm) and coupled to a 6125B Single Quad Mass Spectrometer from Agilent or an Agilent 1100 Series HPLC equipped with a Luna 4251-E0 C_18_ column (3 μm, 4.6 x 150 mm) and coupled to a PE SCIEX API 3000 mass spectrometer (wavelengths monitored = 215, 254 nm). All final compounds and intermediates were purified using either a CombiFlash 200 Rfi (Teledyne Isco) with a 4 or 12 g reverse phase C_18_ RediSep Rf Gold column (wavelengths monitored = 215 & 254 nm) or with a normal phase silica hand packed columns. Additionally, all macrocycles were purified using a preparatory Agilent 1260 Infinity HPLC system equipped with a Luna Omega 5 µm C18 100A LC column 250 x 10 mm (Catalog No.: 00G-4785-N0) (wavelengths monitored = 215 & 254 nm). The gradients used are described in the synthetic methods section. The identity of all reaction intermediates was confirmed by LC-MS.

**Solid Phase Peptide Synthesis**

All amino acids have L stereochemistry. Standard Fmoc-protected amino acids Fmoc-Ala-OH, Fmoc-Arg(Pbf)-OH, Fmoc-Asn(Trt)-OH, Fmoc-Asp(tBu)-OH, Fmoc-Cys(Trt)-OH, Fmoc-Gln(Trt)-OH, Fmoc-Glu(tBu)-OH, Fmoc-Gly-OH, Fmoc-His(Trt)-OH, Fmoc-Ile-OH, Fmoc-Leu-OH, Fmoc-Lys(Boc)-OH, Fmoc-Met-OH, Fmoc-Phe-OH, Fmoc-Pro-OH, Fmoc-Ser(But)-OH, Fmoc-Thr(But)-OH, Fmoc-Trp(Boc)-OH, Fmoc-Tyr(tBu)-OH), Fmoc-Val-OH, were purchased from Advanced ChemTech, Chem-Impex, AAPPTec and Novabiochem.

Linear peptides were synthesized on Rink amide resin (loading 100-200 mesh, 0.68 meq/g, 1% DVB) (Chem-Impex Int’L INC, Cat. # 02900) using standard Fmoc chemistry. The peptides were synthesized using A Syro II (Biotage) fully automated parallel synthesizer with a standard reactor block using a 2 mL reaction vessel (PP-Reactor, 2 mL, with PE Frit, Cat. # V020PE051) (2mL plunger, Cat. # V020ST020). Scale up reactions were performed manually.

Peptides were cleaved from the resin using a modified reagent K mixture: 90% v/v trifluoroacetic acid (TFA) from Chem-Impex Int’l Inc. (Catalog No. 00289), 2.5% w/v phenol from Sigma-Aldrich (Catalog No. 328111), 2.5% v/v MilliQ water, 2.5% v/v thioanisole from Sigma-Aldrich (Catalog No. 28002), and 2.5% v/v 1,2-ethanedithiol from Sigma-Aldrich (Catalog No. 02390), for 3 hours at room temperature (RT) with minor agitation. The mixture was separated from the beads and collected into a 15-mL conical tube. The resin was washed with an additional minimal amount of the modified reagent K mixture and collected. Diethyl ether was then added to the reagent K mixture to precipitate the peptide. This mixture was allowed to cool for 2 hours at -20°C, then centrifuged at 275 G, and the diethyl ether was removed. This process was repeated two more times, for a total of three ether washes. Residual ether was allowed to evaporate briefly, followed by the addition of 2 mL of 50/50 ACN/H_2_O (0.1% TFA). The linear peptides were then lyophilized. In general, most peptides were synthesized on a 30 mg resin scale and were cleaved with a total volume of 1.2 mL of reagent K, with 0.5 mL used for washing, not exceeding 2 mL.

**General procedure A for Fmoc deprotection:** A 40% piperidine in DMF solution was added to the resin for 3 minutes with shaking, then drained. Following this, a 20% piperidine in DMF solution was added to the resin for 10 minutes with shaking, then drained. The reaction vessel was fully drained.

**General procedure B for resin washing**: Three washing steps were performed between couplings and deprotections using DMF. DMF was added to the reaction vessel and allowed to sit for 1 minute with gentle shaking. The reaction vessel was then fully drained.

**General procedure C for amide bond couplings**: 2-(1H-Benzotriazol-1-yl)-1,1,3,3-tetramethyluronium hexafluorophosphate (HBTU) (Peptides International, Catalog No.: KHB-1065-PI), 2,4,6-collidine (Alfa Aesar, Catalog No.: A11058), and the desired amino acids were pre-dissolved in DMF. To the reaction vessel 5 equivalents (eq) of HBTU, 10 eq of 2,4,6-collidine, and 5 eq of amino acid were added, using minimal DMF, and allowed to react for 40 minutes at RT, gently shaking it twice. The reaction vessel was then drained.

**Synthetic procedure for fluorogenic macrocyclic peptide substrates preparation**

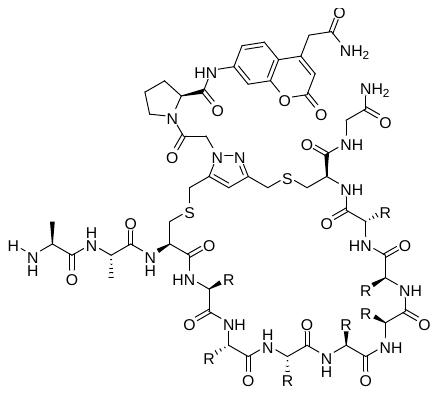

To a 2mL Eppendorf tube, ~6 mg of lyophilized linear peptide (1 eq, 5 µmol) was added and dissolved using the reducing reaction buffer (2 mM TCEP, 50 mM NaHCO_3_ in MilliQ H_2_O, pH=8). For hydrophobic peptides, ACN was added to increase solubility, for most cases 700 µL of reducing reaction buffer and 300 µL of ACN were used. In all cases, the percentage of ACN did not exceed 40% nor did the total reaction volume exceed 1.2 mL. The pH was verified to be basic (~8 pH) before proceeding. The peptides were rotated for 30 min at RT. Then 0.85 mg of 1,5-dichloropentane-2,4-dione (DPD) in 22 µL of DMF (1 eq, 5 µmol; Ambeed Inc. A1340600) was added and allowed to react for exactly 30 min with rotation at RT. Next, 100 µL of acidic reaction buffer (1M NH_4_Ac pH = 4.0 in MilliQ H_2_O) was added. The pH was verified to be acidic (~4 pH) before proceeding. When not acidic, subsequent volumes (50 µL) of acidic buffer were added. For insoluble peptides, additional ACN was added not exceeding 40% of the total volume nor increasing the total volume over 1.5mL. Then 1.7 mg (0.9 eq, 4.5 µmol) of biorthogonal linker dissolved in 30 µL of MilliQ H_2_O were added and reacted for 2 hours at 37°C with rotation. The reaction was then lyophilized. For crude macrocycles, the resulting solid was dissolved in 500 µL of DMSO to make a 10 mM crude stock. For purified macrocycles, the solid was purified via preparatory HPLC using a gradient of 5-95% ACN (0.1% TFA) in H_2_O (0.1% TFA) over 20 min. The resulting purified macrocycle was dissolved in 10 mM DMSO stocks.

**Synthetic procedure for biorthogonal panning linker preparation**

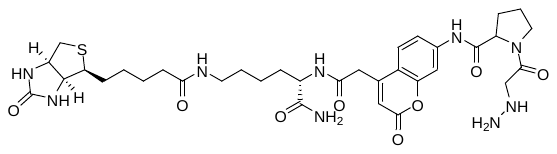

Rink amide resin (1 eq, 1 g) was treated with DMF for 1 hour to swell the resin. The resin was washed, deprotected, and then washed again. Next, Fmoc-Lys(Biotin)-OH (3 eq, 2.4 mmol, 859 mg) was dissolved in DMF with heating (using a heat gun) and O-(7-Azabenzotriazolyl)-tetramethyluronium hexafluorophosphate (HATU) (2.5 eq, 1.2 mmol, 456 mg) (Oakwood Chemical, Catalog No.: 023926) and collidine (2.5 eq, 1.2 mmol, 156 µL) (Chem-Impex, Catalog No.: 00110) were added and agitated for 10 min, before addition to the resin. The whole was allowed to react overnight at RT with agitation. Next, the resin was washed, the Fmoc deprotected and washed again. In a falcon tube Fmoc-ACC (2.5 eq, 1.2 mmol, 529.7 mg), HATU (2.5 eq, 1.2 mmol, 456 mg) and collidine (2.5 eq, 1.2 mmol, 156 µL) were allowed to react for 10 min and added to the resin to react for 2h. After this time, subsequent washes with DMF and DCM were performed and the coupling completion was confirmed in a Kaiser test. The resin was then treated according to procedure A to deprotect the Fmoc and washed. In a falcon tube Fmoc-Pro-OH (2.5 eq, 1.2 mmol, 404.8 mg), HATU (2.5 eq, 1.2 mmol, 456 mg) and collidine (2.5 eq, 1.2 mmol, 156 µL) were allowed to react for 10 min and added to the resin to react for 3h at 37ºC. After this time, an additional portion of preactivated Fmoc-Pro-OH (1.25 eq, 0.6 mmol, 202.4 mg), HATU (1.23 eq, 0.6 mmol, 228 mg) and collidine (1.25 eq, 0.6 mmol, 78 µL) was added and allowed to react overnight at RT. After this time subsequent washes with DMF and DCM were performed and the coupling completion was confirmed in a Kaiser test. The resin was then treated according to general procedure A to deprotect the Fmoc and washed. Next, Tri-Boc-hydrazinoacetic acid (2.5 eq, 1.2 mmol, 468.5 mg) (Santa Cruz Biotechnology, Catalog No.: sc-237235), HATU (2.5 eq, 1.2 mmol, 456 mg) and collidine (2.5 eq, 1.2 mmol, 156 µL) dissolved in a minimal amount of DMF were agitated for 3 min to pre-activate before addition to the resin and the whole reacted for 24 h at RT. After this time, subsequent washes with DMF and DCM were performed and the coupling completion was confirmed in a Kaiser test. The resin was then treated according to procedure B to deprotect the Fmoc and washed. To the resin, 10mL of modified reagent B mixture (95% TFA v/v, MilliQ water 2.5% v/v, triisopropylsilane 2.5% v/v from Sigma-Aldrich (Catalog No.: 233781) were added and allowed to react for 2 h at RT. The modified reagent B mixture was collected from the resin, which was re-washed. Cold diethyl ether was then added and kept for 2 h at -20°C. After this, it was centrifuged at 275 G and the ether removed to collect the precipitated peptide. This process was repeated two more times. The residual yellow solid was then dried and dissolved in DCM with minimal MeOH and purified via silica gel flash chromatography using a gradient of 0-25 % MeOH in DCM over 40 min. The purified fractions were combined and evaporated. The residual off-yellow solid was dissolved in 50/50 ACN/H_2_O and lyophilized. (323 mg, 36 % yield). MS(ESI)+ calculated for C_34_H_47_N_9_O_8_S [M+H]^+1^: 742.3302, found 742.5.

**Synthetic procedure for ACC linker preparation**

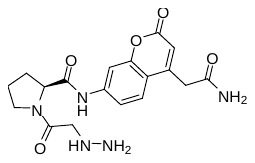

Rink amide resin (1 eq, 960 µmol, 2 g) was swelled with DCM for 10 min, washed, deprotected and washed again. Fmoc-ACC (2.5 eq, 2.4 mmol, 1060 mg) (Aapptec, Catalog No.: CTZ017) and Hydroxybenzotriazole hydrate (HOBt) (2.5 eq, 2.4 mmol, 444 mg) (Oakwood Chemical, Catalog No.: MO2875) were dissolved in minimal DMF. Then DIC (2.5 eq, 2.4 mmol, 3.8mL) (Chem-Impex, Catalog No.: 00110) was added and the whole agitated for 3 min. The pre-activated mixture was added to the resin and reacted for 24 h at RT. The whole was washed, Fmoc deprotected and washed. Next, preactivated Fmoc-Pro-OH (5 eq, 4.8 mmol, 1620 mg), HATU (5 eq, 4.8 mmol, 1824 mg) and collidine (10 eq, 9.6 mmol, 1.2 mL) in DMF added to the resin and reacted for 24 h at RT. After this time, the mixture was drained and the procedure repeated and allowed to react for another 24 h at RT. After this time, the coupling was still not completed and the resin was treated with Fmoc-Pro-OH (5 eq, 4.8 mmol, 1620 mg), HATU (5 eq, 4.8 mmol, 1824 mg), and N,N-Diisopropylethylamine (DIPEA) (7.2 eq, 6.9 mmol, 1.2 mL) in a minimal amount of DMF. The whole was allowed to react for 3 h at 37°C with agitation. After this, the coupling was complete and the resin was washed, Fmoc deprotected and washed. To the whole, preactivated tri-boc-hydrazinoacetic acid (2.5 eq, 2.4 mmol, 936 mg), HATU (2.5 eq, 2.4 mmol, 912 mg) and collidine (10 eq, 9.6 mmol, 1.2 mL) in minimal amount of DMF were added and reacted for 24 h at RT with agitation. After this time, subsequent washes with DMF and DCM were performed and the coupling completion was confirmed in a Kaiser test. The resin was then treated according to procedure B to deprotect the Fmoc and washed. To the resin, 10 mL of modified reagent B mixture (95% TFA v/v, MilliQ water 2.5% v/v, triisopropylsilane 2.5% v/v) were added and allowed to react for 2 h at RT. The modified reagent B mixture was collected from the resin, which was re-washed. Cold diethyl ether was then added and kept for 2 h at -20°C. After this, it was centrifuged at 275 G and the ether removed to collect the precipitated peptide. This process was repeated two more times. The residual yellow solid was then dried and dissolved in DCM with minimal MeOH and purified via silica gel flash chromatography using a gradient of 0-25 % MeOH in DCM over 40 min. The purified fractions were combined and evaporated. The residual off-yellow solid was dissolved in 50/50 ACN/H_2_O and lyophilized. (280 mg, 56 % yield). MS(ESI)+ calculated for C_18_H_21_N_5_O_5_ [M+H]^+1^: 388.1576, found 388.0. All washes were performed according to general procedure B and the Fmoc deprotections, following the general procedure A.
