## Supplementary material for "Macrocyclic phage display for identification of selective protease substrates": Supplmental Spectra

Supplementary Spectra

Crude Molecules

DPP4 1 Crude

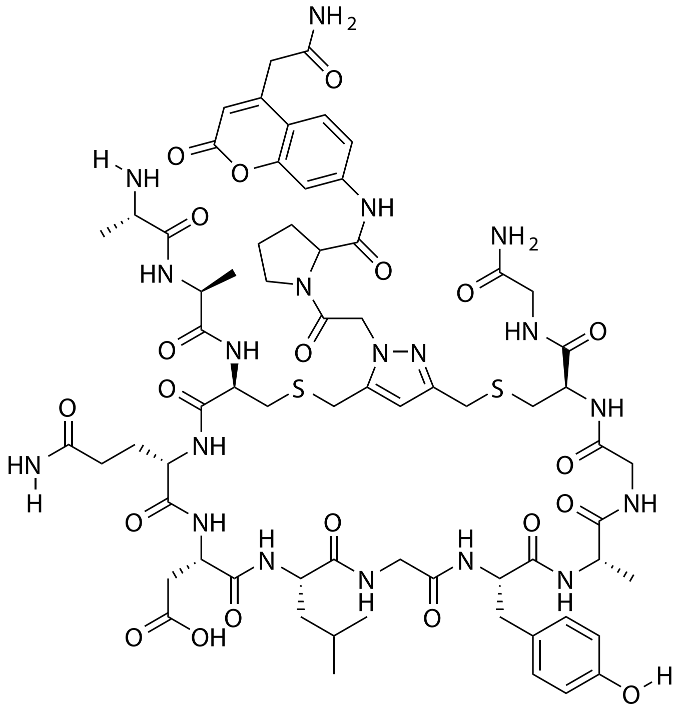

| Probe Name | Sequence | Expected Mass $[M+1]^{+1}$ or $[M+2]^{+2}$ | Found Mass |
|---|---|---|---|
| DPP4 1 | AACQDLGYAGCG | 1574.6112 | 1574.9 |

Total Ion Chromatogram

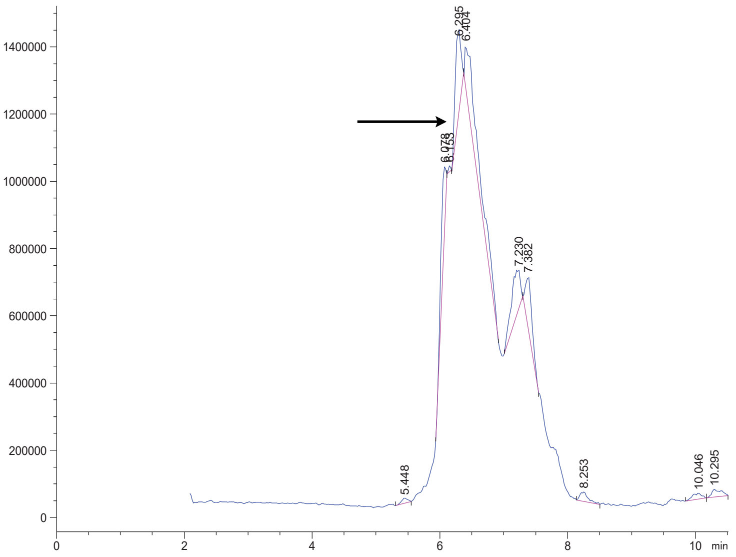


Variable Wavelength Detector: 220 nm

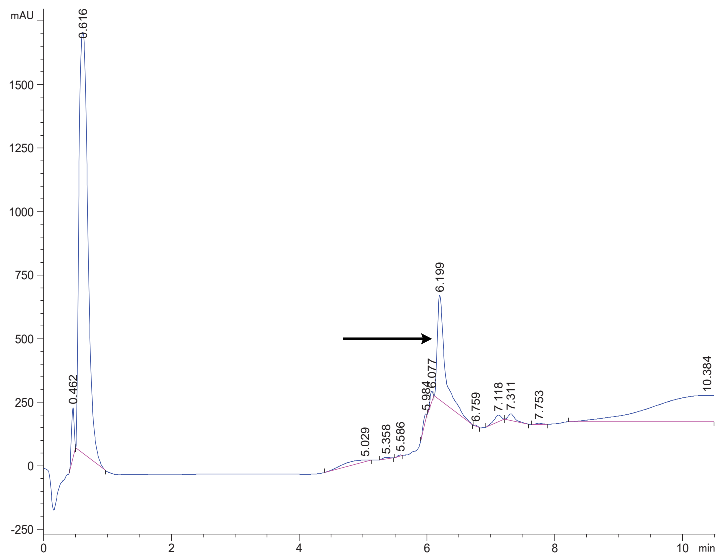

DPP4 2 Crude

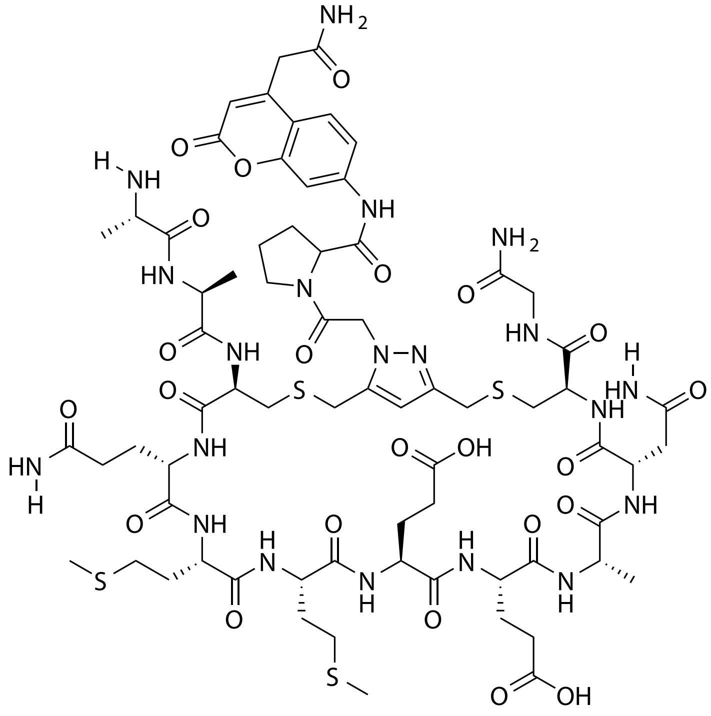


| Probe Name | Sequence | Expected Mass [M+1]$^{+1}$ or [M+2]$^{+2}$ | Found Mass |
|---|---|---|---|
| DPP4 2 | AACQMMEEANCG | 1703.6030 | 1703.9 |

Total Ion Chromatogram

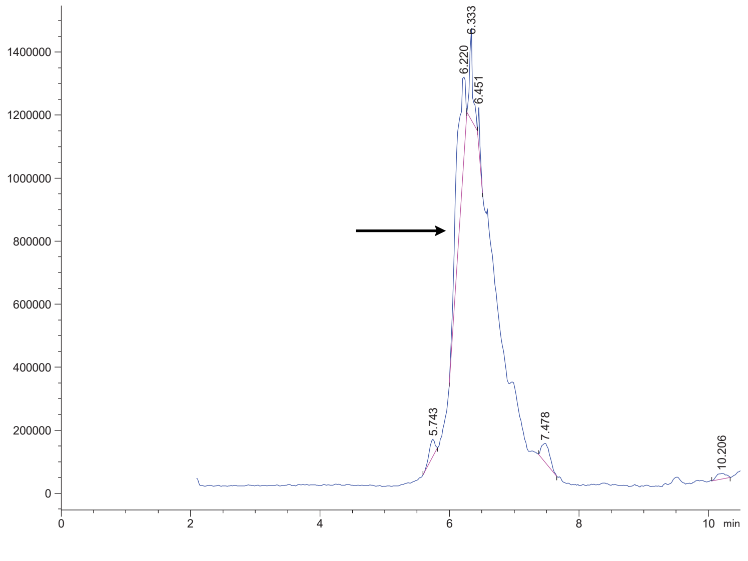


Variable Wavelength Detector: 220 nm

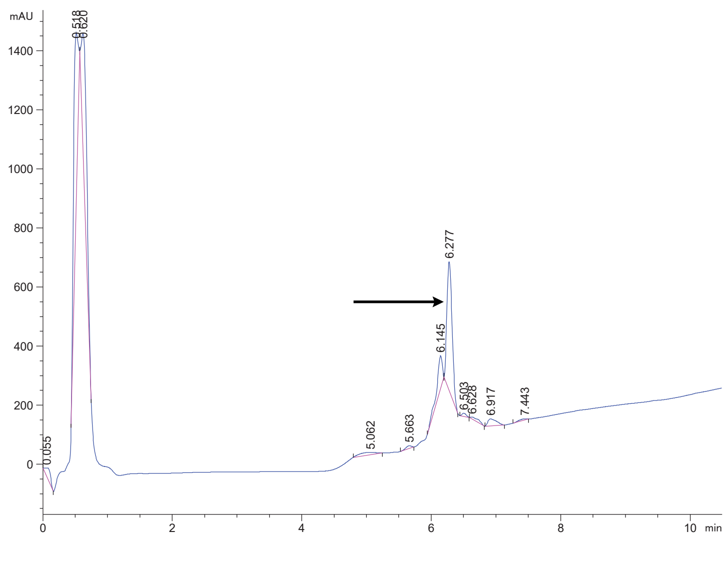

DPP4 3 Crude

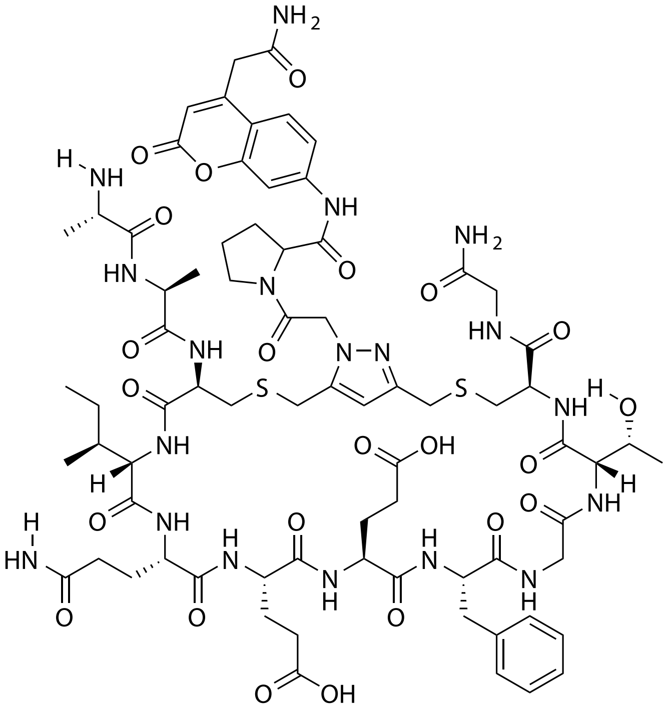

| Probe Name | Sequence | Expected Mass $[M+1]^{+1}$ or $[M+2]^{+2}$ | Found Mass |
|---|---|---|---|
| DPP4 3 | AACIQEEFGTCG | 1674.6636 | 1674.9 |

Total Ion Chromatogram

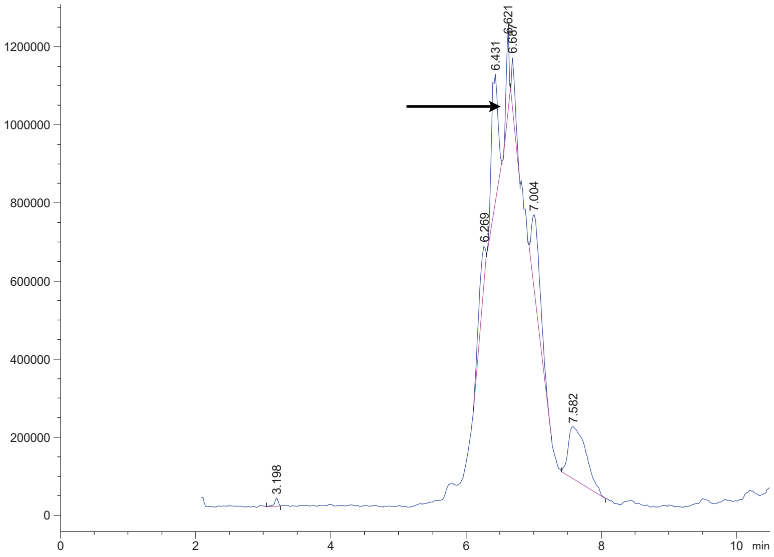


Variable Wavelength Detector: 220 nm

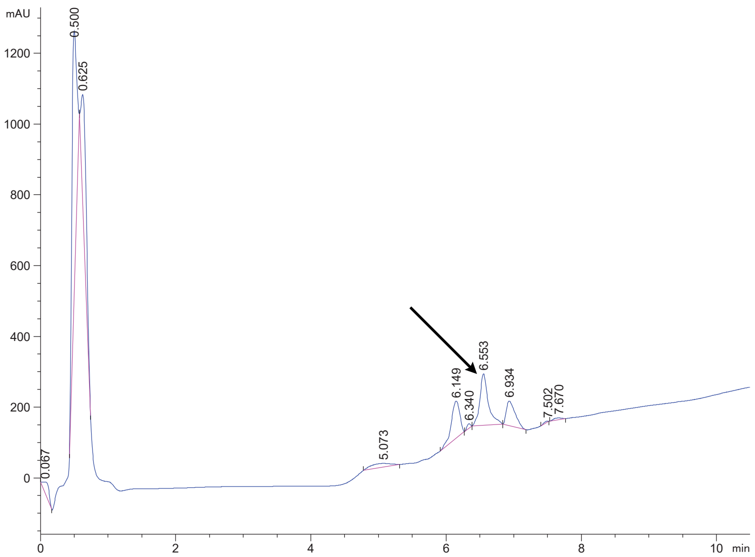

DPP4 4 Crude

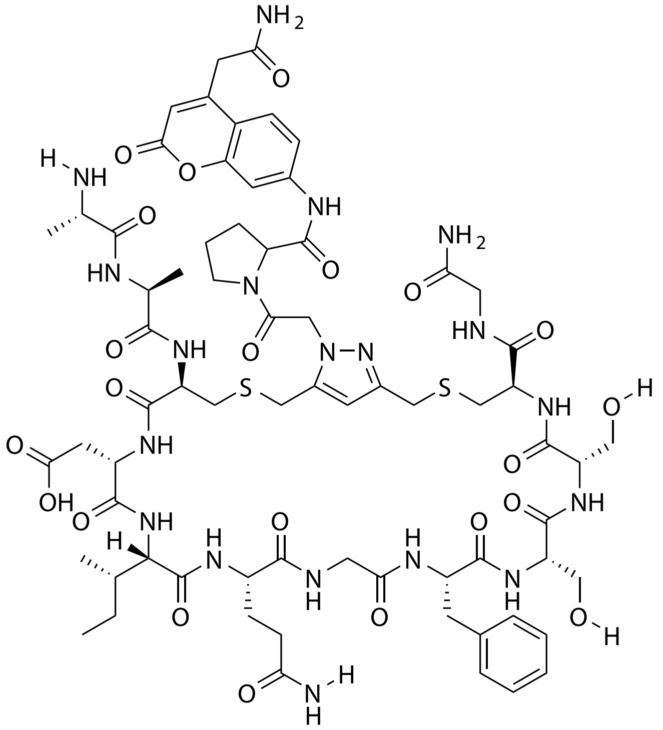


| Probe Name | Sequence | Expected Mass $[M+1]^{+1}$ or $[M+2]^{+2}$ | Found Mass |
|---|---|---|---|
| DPP4 4 | AACDIQGFSSCG | 1604.6218 | 1604.9 |

Total Ion Chromatogram

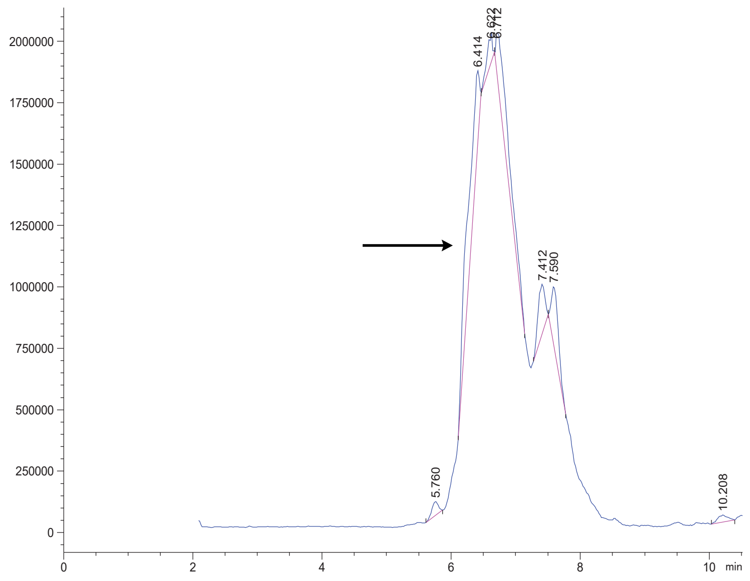


Variable Wavelength Detector: 220 nm

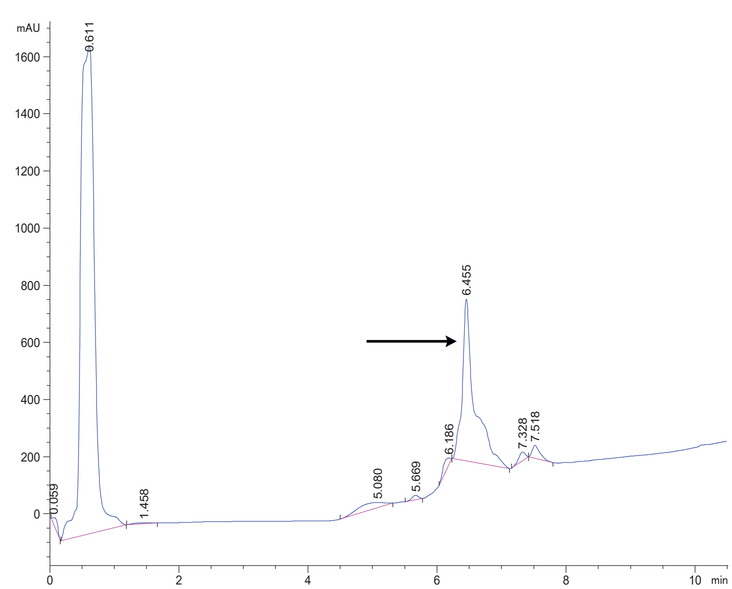

### DPP4 5 Crude

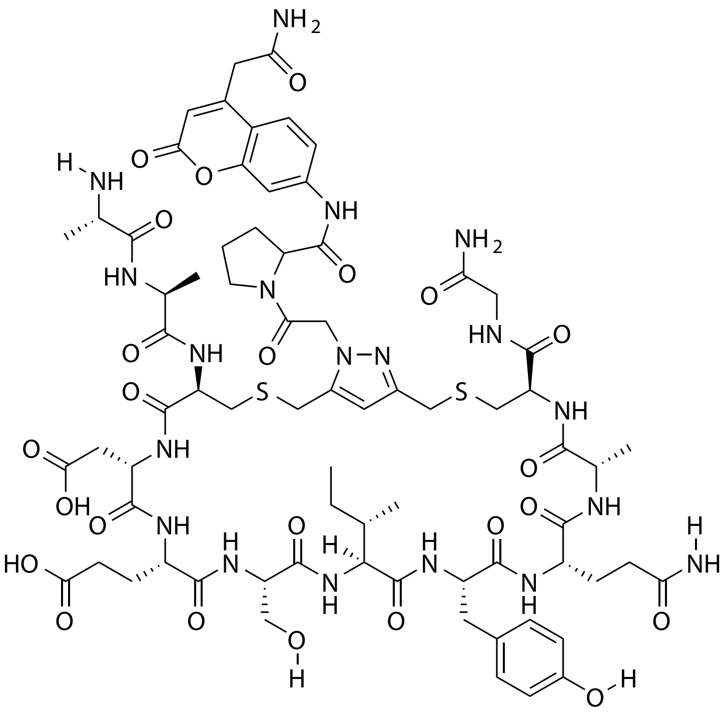

| **Probe Name** | **Sequence** | **Expected Mass $[M+1]^{+1}$ or $[M+2]^{+2}$** | **Found Mass** |
|---|---|---|---|
| DPP4 5 | AACDESIYQACG | 1676.6429 | 1676.9 |

Total Ion Chromatogram

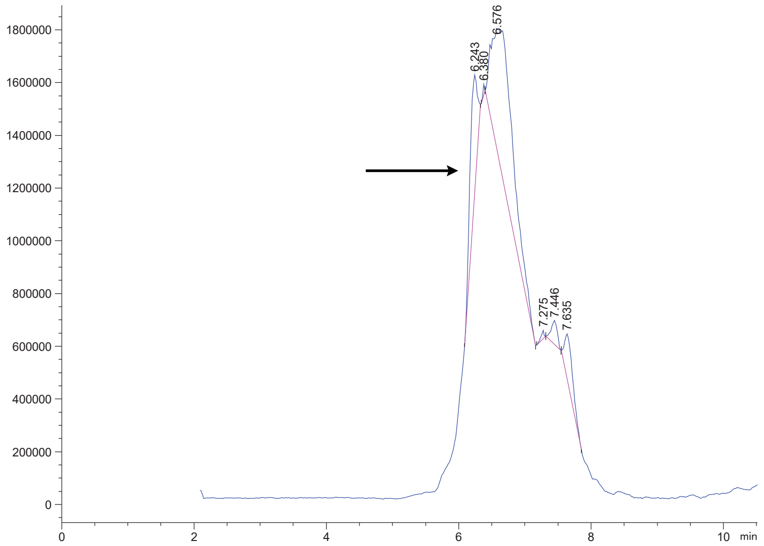


Variable Wavelength Detector: 220 nm

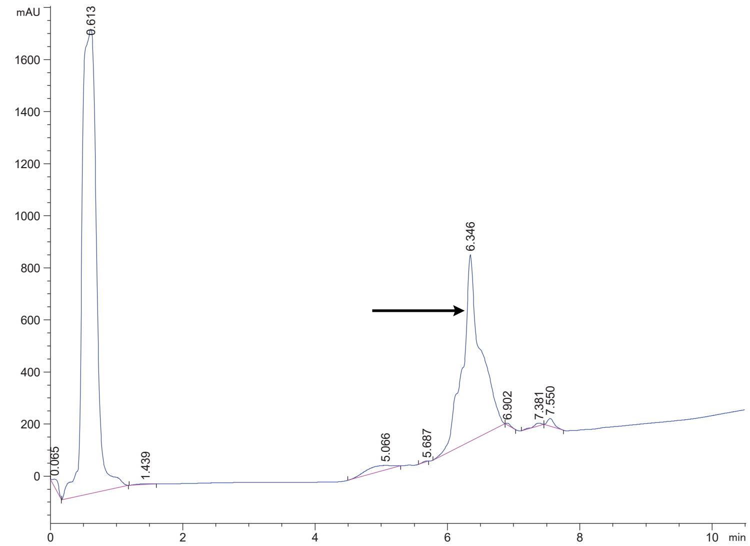

### DPP4 6 Crude

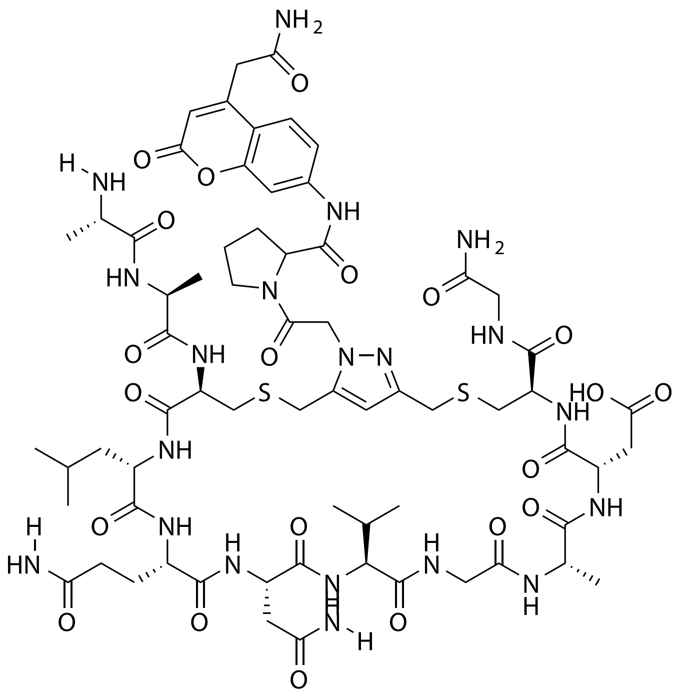

| Probe Name | Sequence | Expected Mass $[M+1]^{+1}$ or $[M+2]^{+2}$ | Found Mass |
|---|---|---|---|
| DPP4 6 | AACLQNVGADCG | 1567.6377 | 1567.9 |

Total Ion Chromatogram

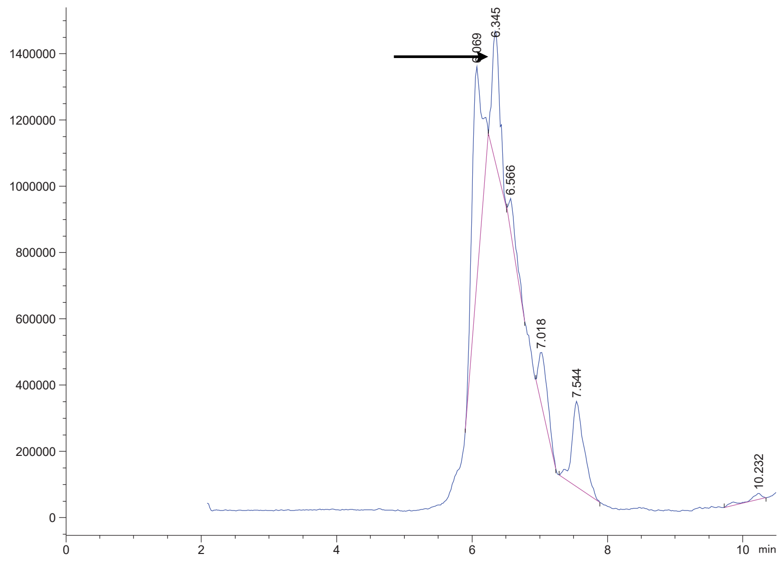

Variable Wavelength Detector: 220 nm

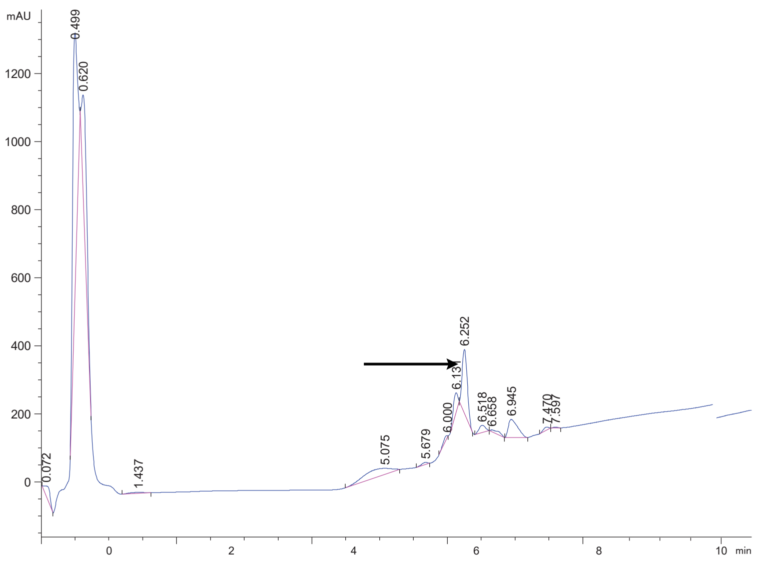

DPP4 7 Crude

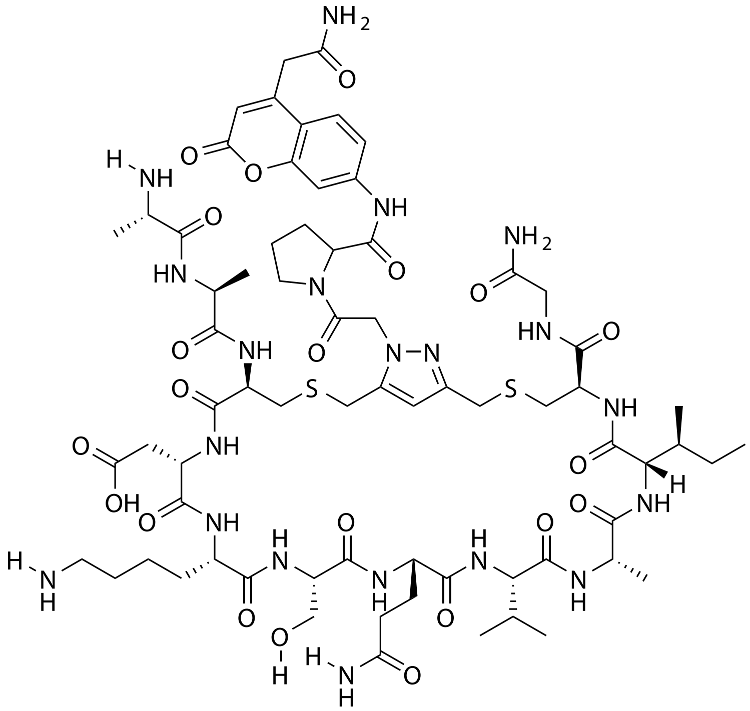

| Probe Name | Sequence | Expected Mass $[M+1]^{+1}$ or $[M+2]^{+2}$ | Found Mass |
|---|---|---|---|
| DPP4 7 | AACDKSQVAICG | 806.3540 $[M+2]^{+2}$ | 806.3 $[M+2]^{+2}$ |

Total Ion Chromatogram

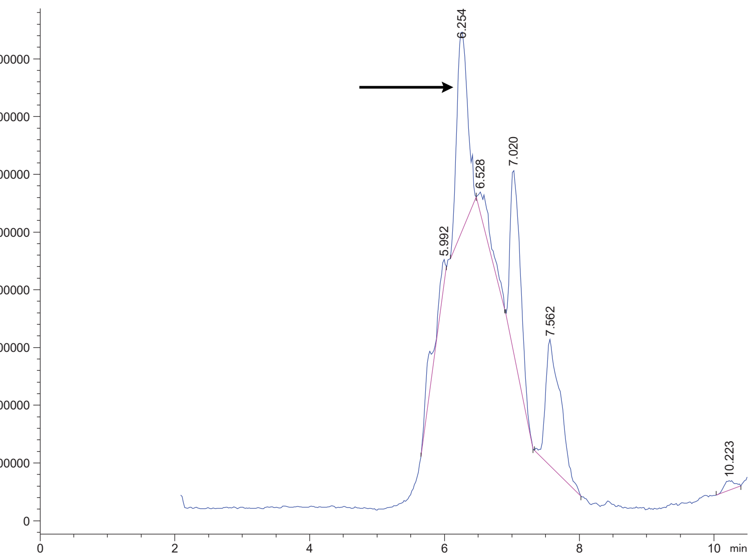


Variable Wavelength Detector: 220 nm

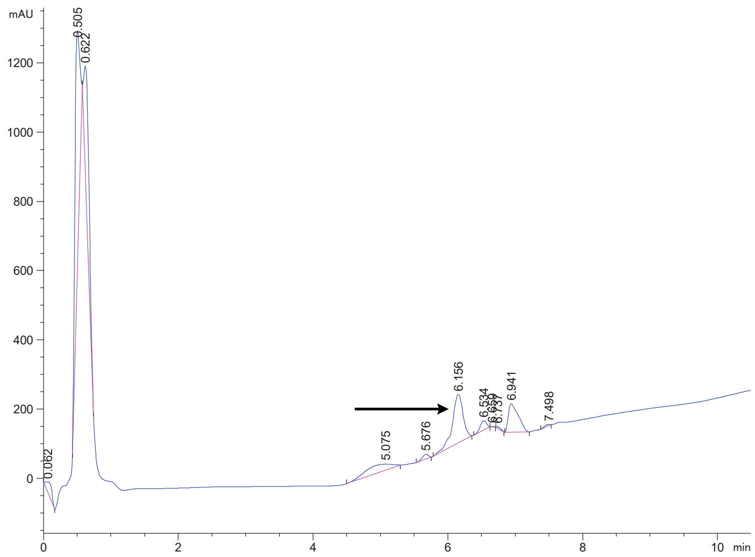

### DPP4 8 Crude

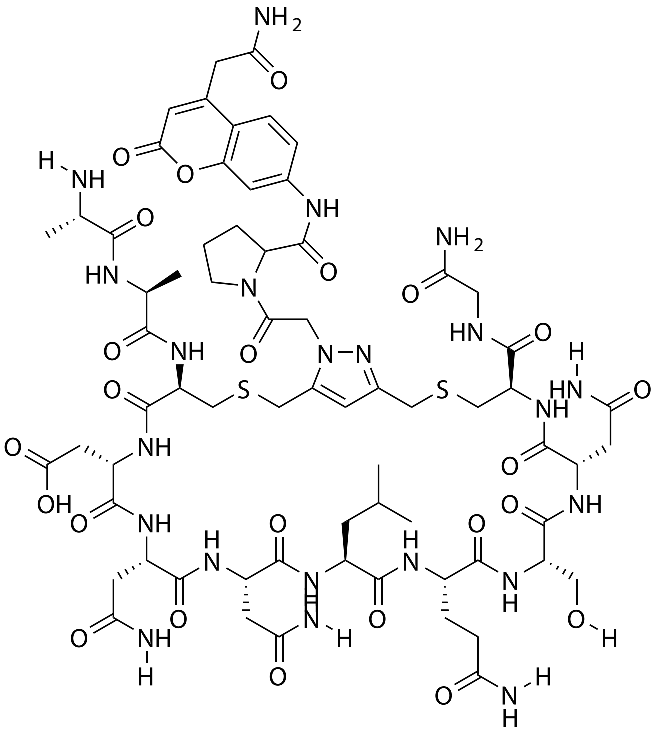

| Probe Name | Sequence | Expected Mass $[M+1]^{+1}$ or $[M+2]^{+2}$ | Found Mass |
|---|---|---|---|
| DPP4 8 | AACDNNLQSNCG | 1655.6286 | 1655.9 |

Total Ion Chromatogram

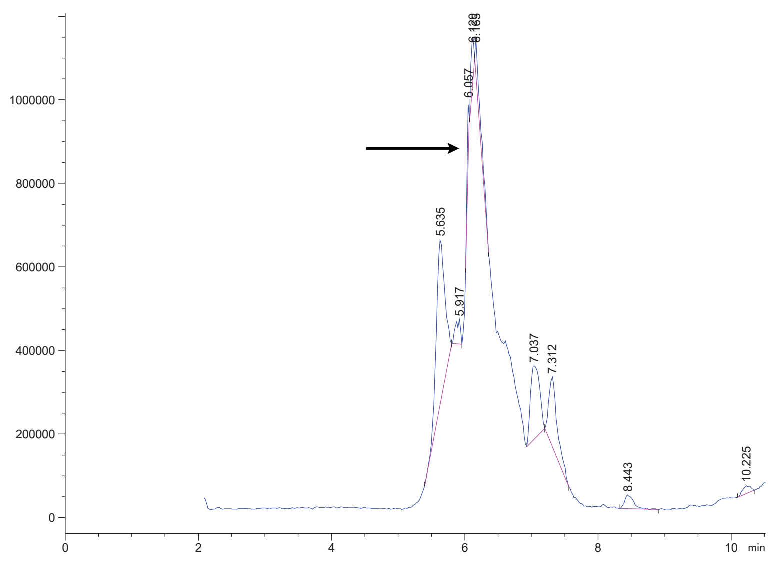


Variable Wavelength Detector: 220 nm

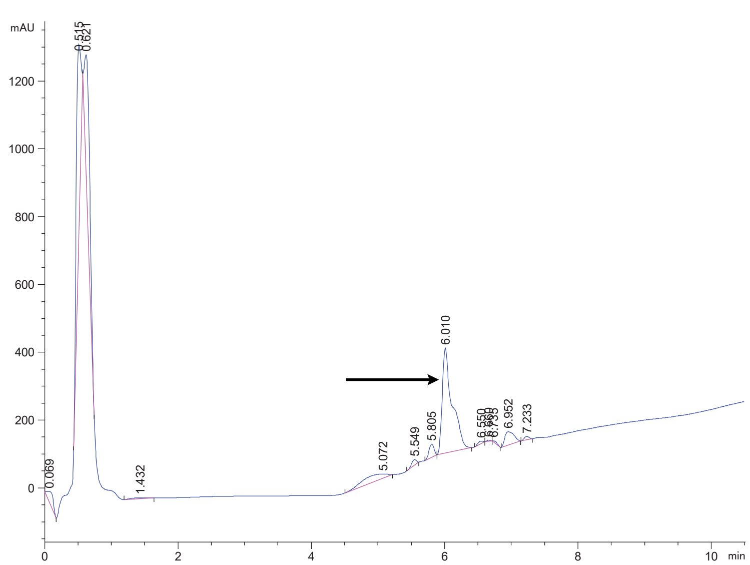

### DPP4 9 Crude

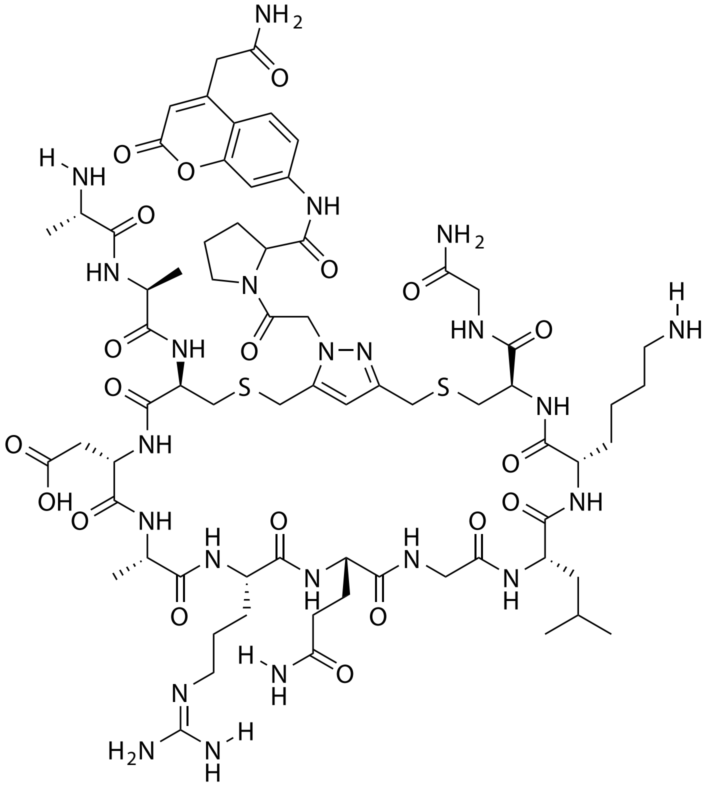

| Probe Name | Sequence | Expected Mass $[M+1]^{+1}$ or $[M+2]^{+2}$ | Found Mass |
|---|---|---|---|
| DPP4 9 | AACDARQGLKCG | 1638.7225 | 1638.0 |

Total Ion Chromatogram

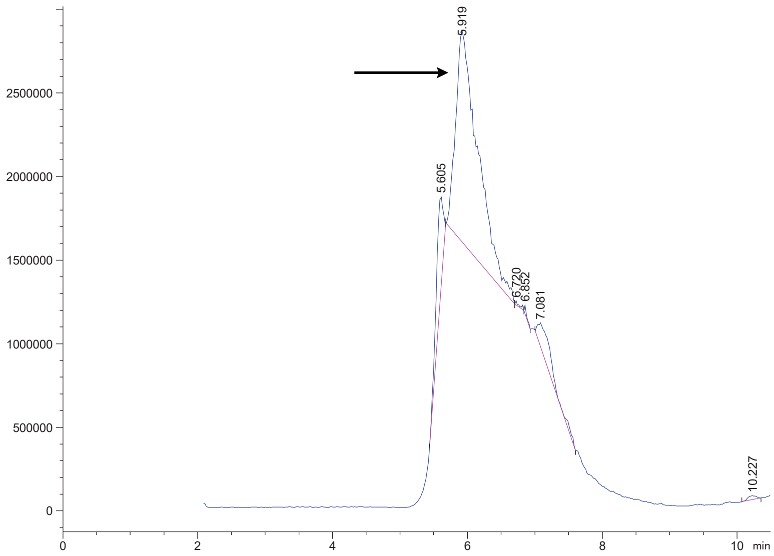


Variable Wavelength Detector: 220 nm

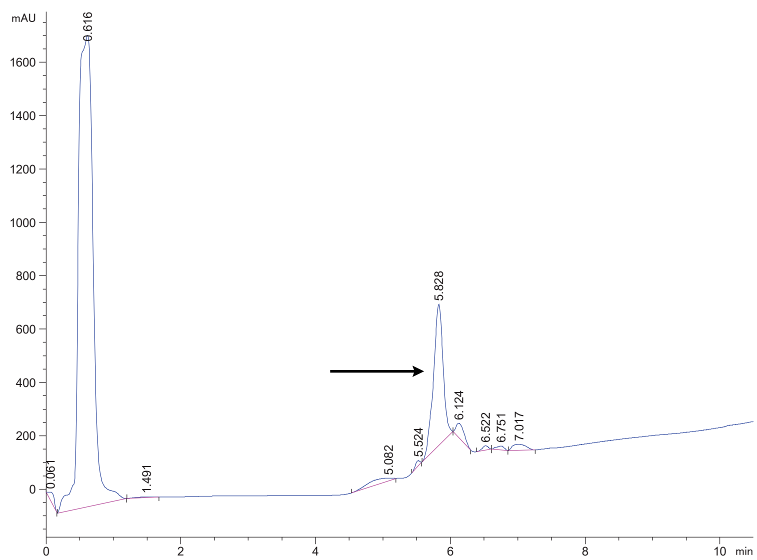

DPP4 10 Crude

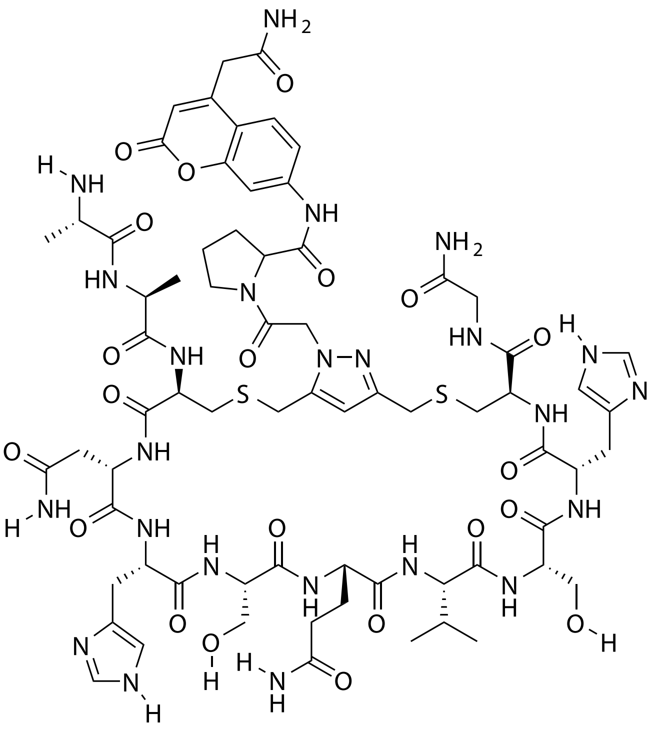

| Probe Name | Sequence | Expected Mass $[M+1]^{+1}$ or $[M+2]^{+2}$ | Found Mass |
|---|---|---|---|
| DPP4 10 | AACNHSQVSHCG | 1659.6500 | 1659.9 |

Total Ion Chromatogram

Variable Wavelength Detector: 220 nm

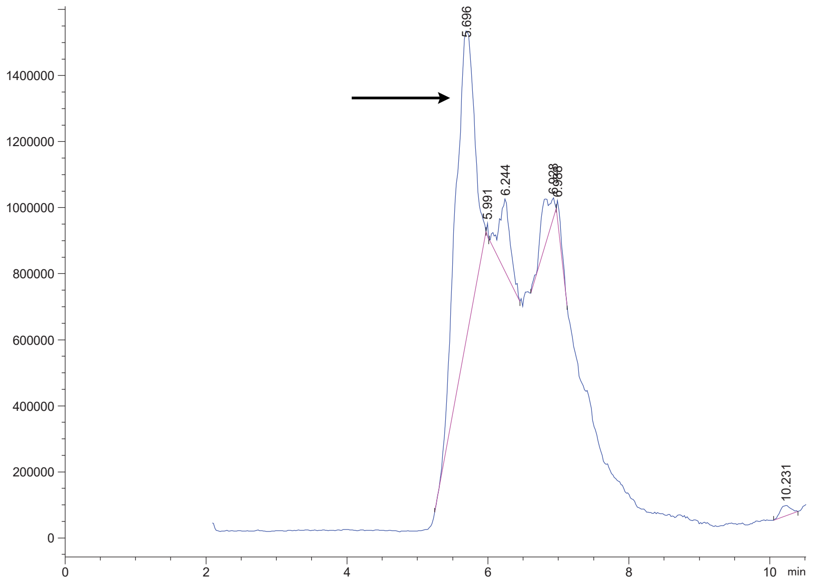

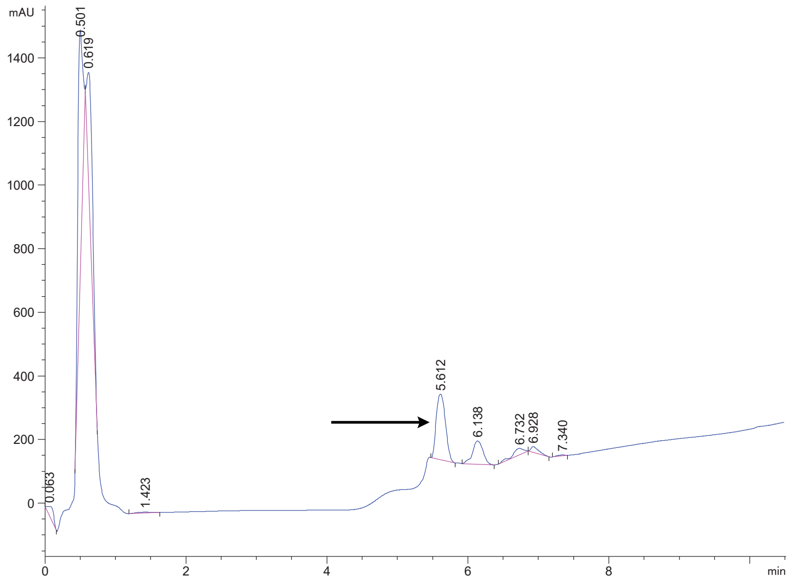

DPP4 11 Crude

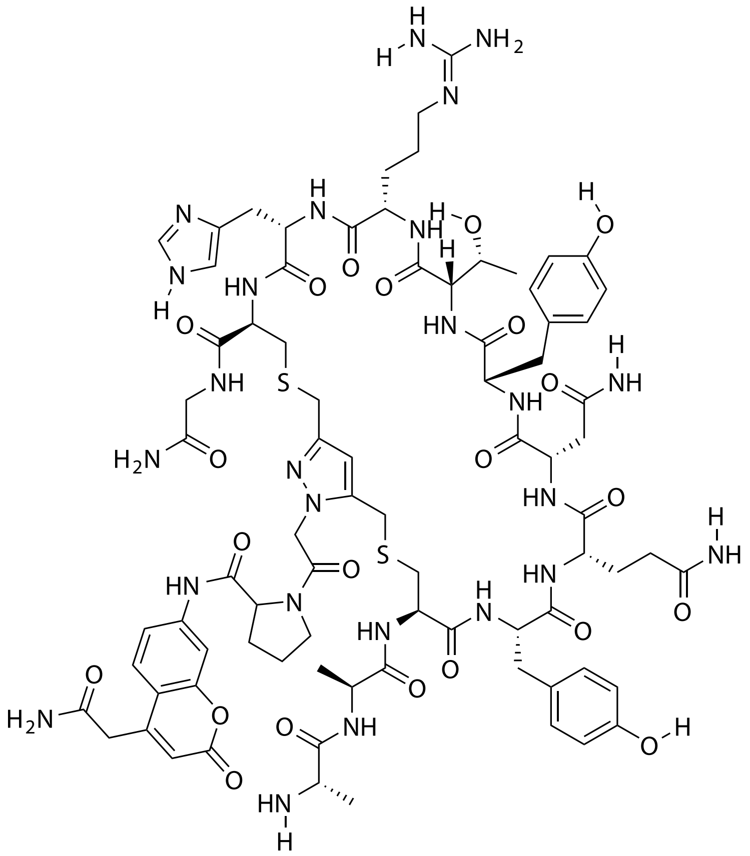

| Probe Name | Sequence | Expected Mass $[M+1]^{+1}$ or $[M+2]^{+2}$ | Found Mass |
|---|---|---|---|
| DPP4 11 | AACYQNYTRHCG | 1832.7341 $[M+2]^{+2}$ | 916.7 $[M+2]^{+2}$ |

Total Ion Chromatogram

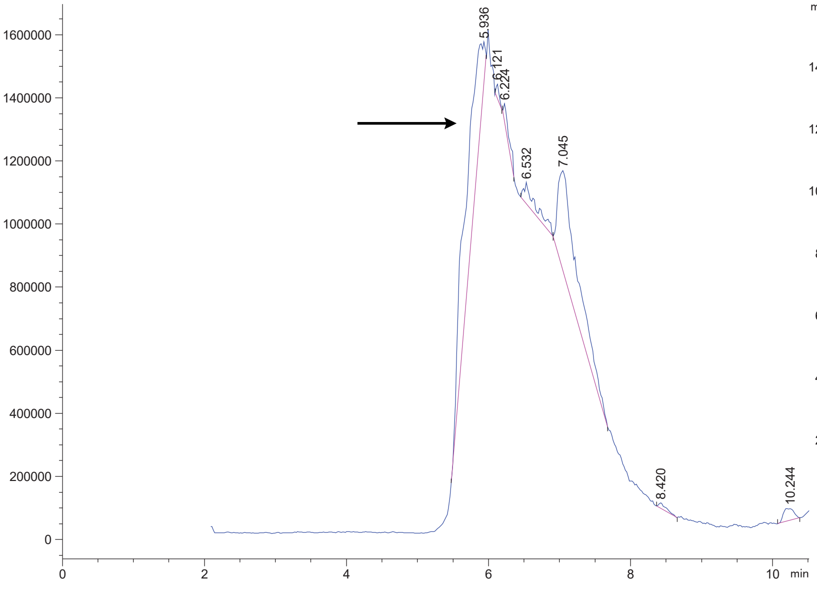

Variable Wavelength Detector: 220 nm

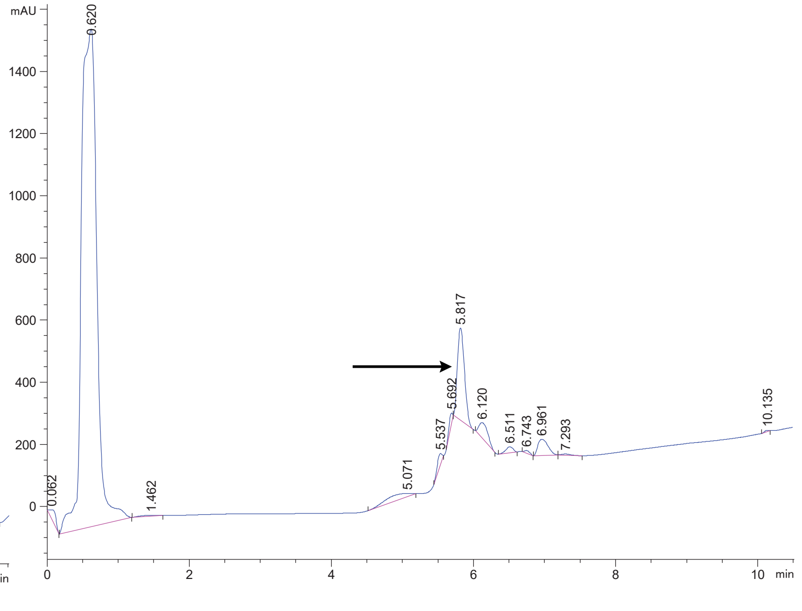

### DPP4 12 Crude

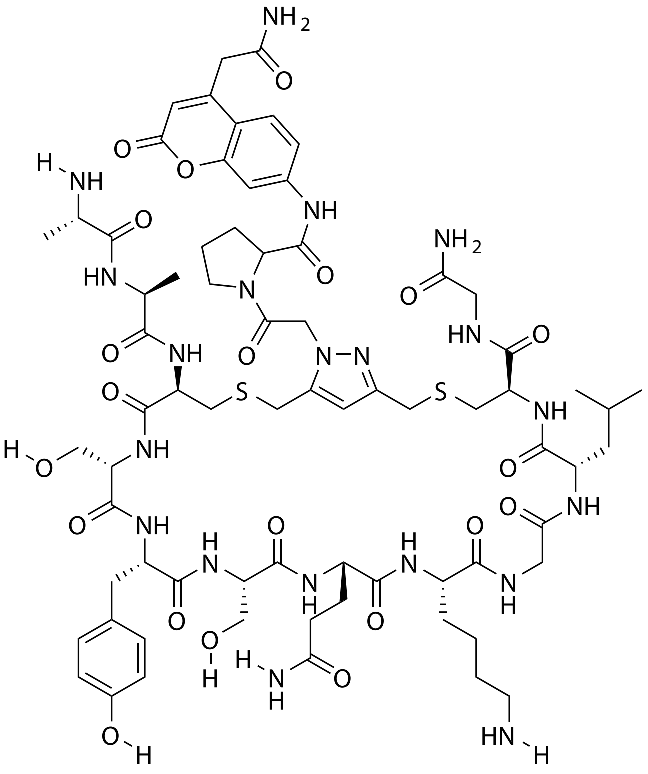

| Probe Name | Sequence | Expected Mass $[M+1]^{+1}$ or $[M+2]^{+2}$ | Found Mass |
|---|---|---|---|
| DPP4 12 | AACSYSQKGLCG | 1633.6847 | 1634.0 |

Total Ion Chromatogram

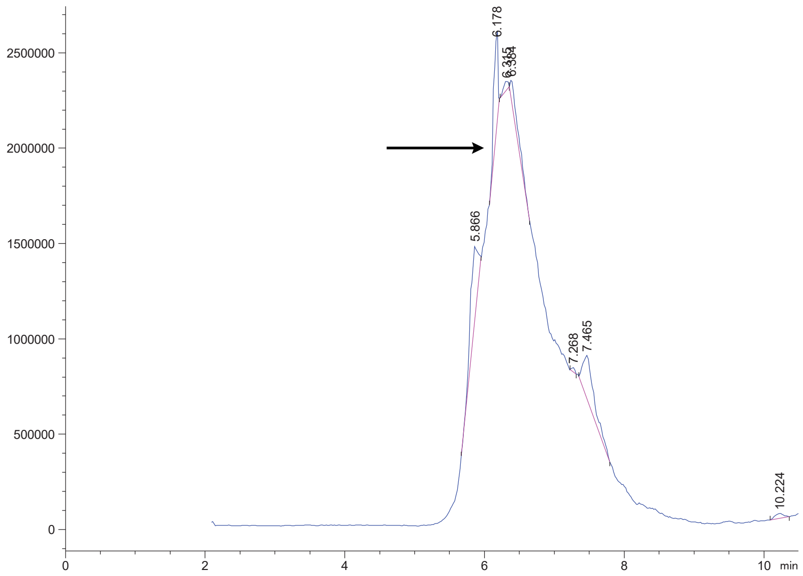


Variable Wavelength Detector: 220 nm

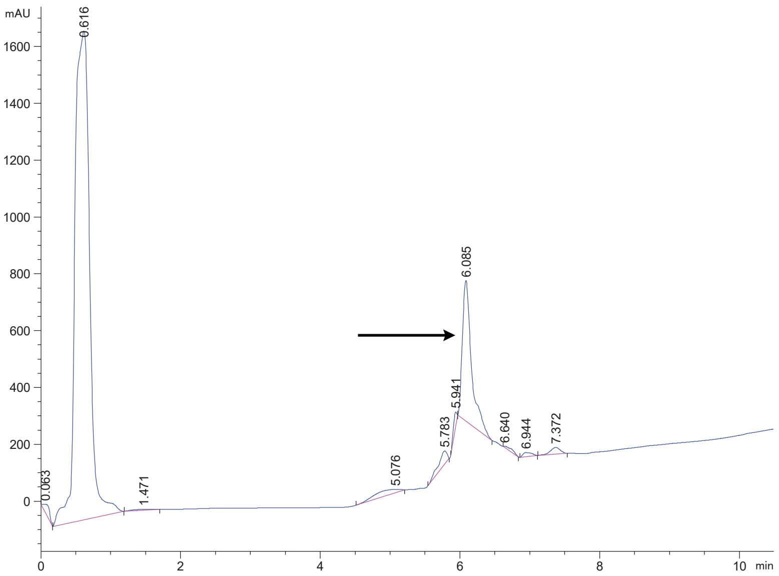

### DPP4 13 Crude

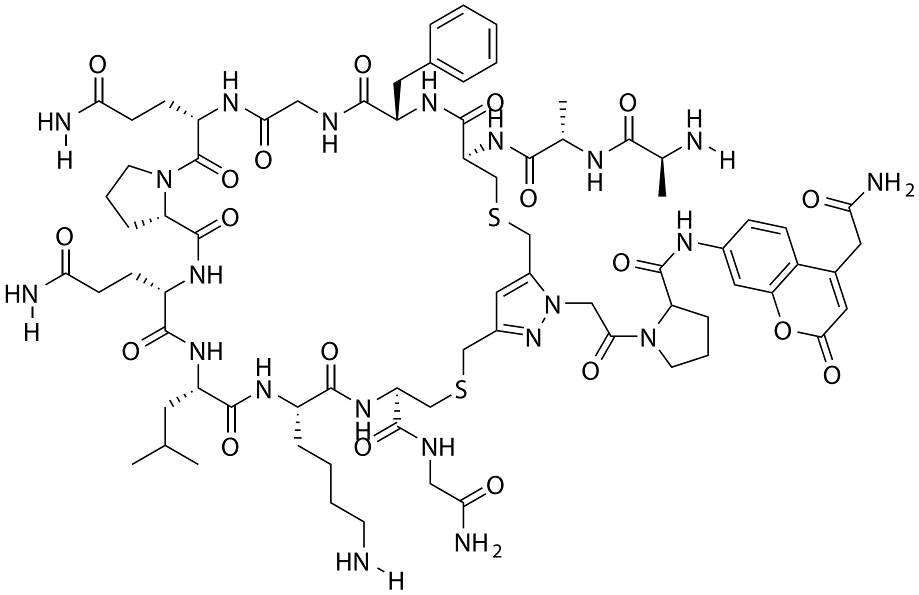

| **Probe Name** | **Sequence** | **Expected Mass $[M+1]^{+1}$ or $[M+2]^{+2}$** | **Found Mass** |
|---|---|---|---|
| DPP4 13 | AACFGQPQLKCG | 1668.7371 | 1669.0 |

Total Ion Chromatogram

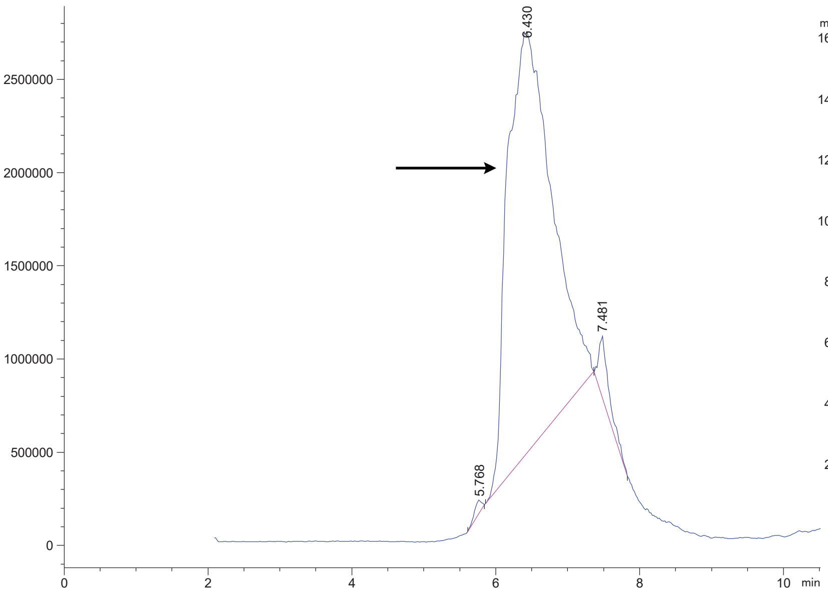


Variable Wavelength Detector: 220 nm

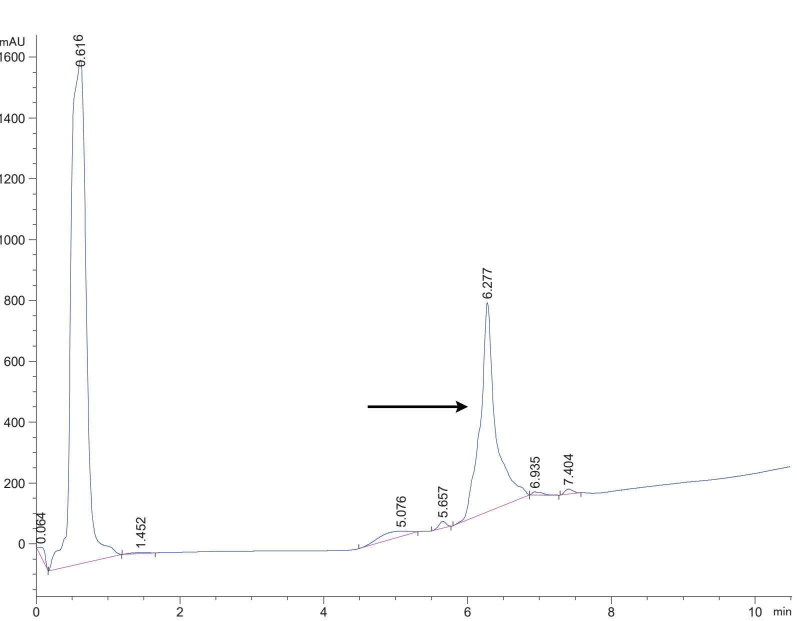

DPP4 14 Crude

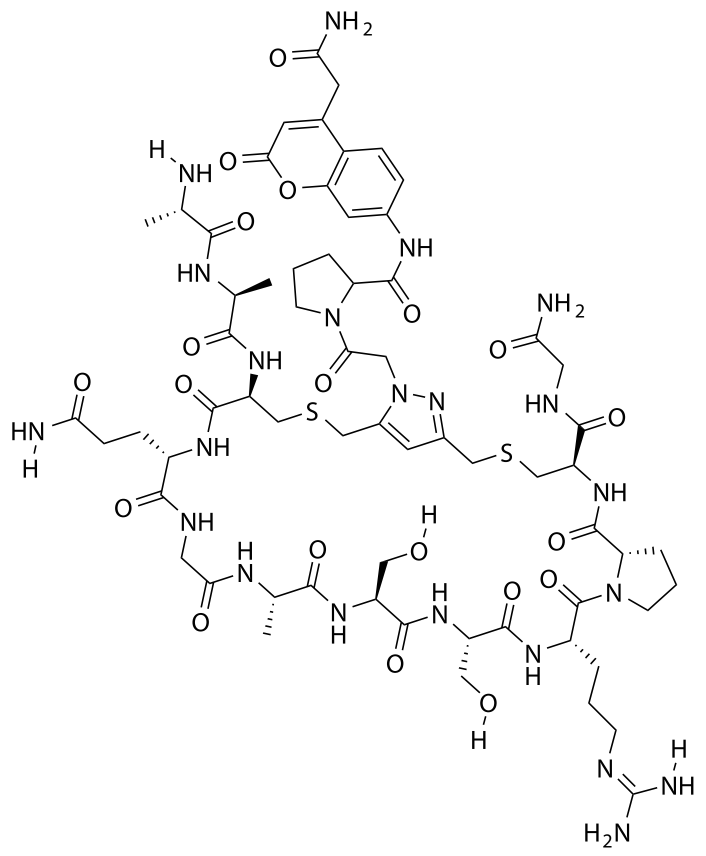

| Probe Name | Sequence | Expected Mass $[M+1]^{+1}$ or $[M+2]^{+2}$ | Found Mass |
|---|---|---|---|
| DPP4 14 | AACQGASSRPCG | 1553.6333 | 1553.9 |

Total Ion Chromatogram

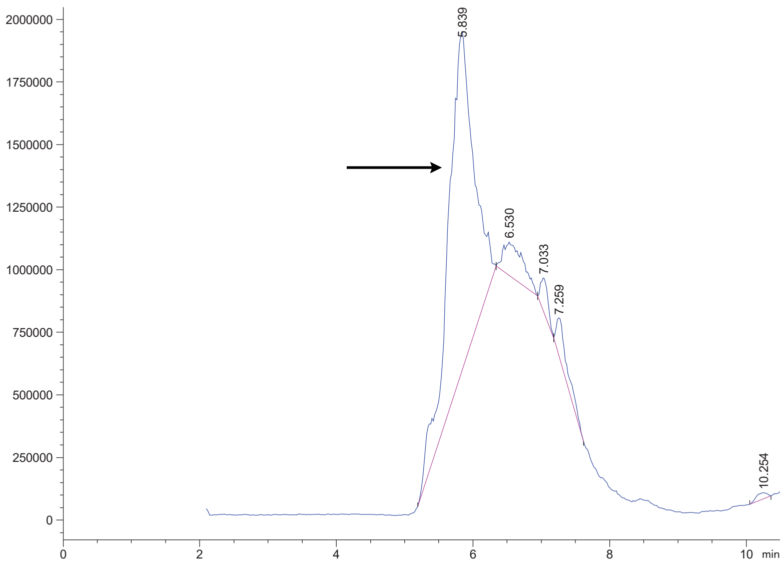

Variable Wavelength Detector: 220 nm

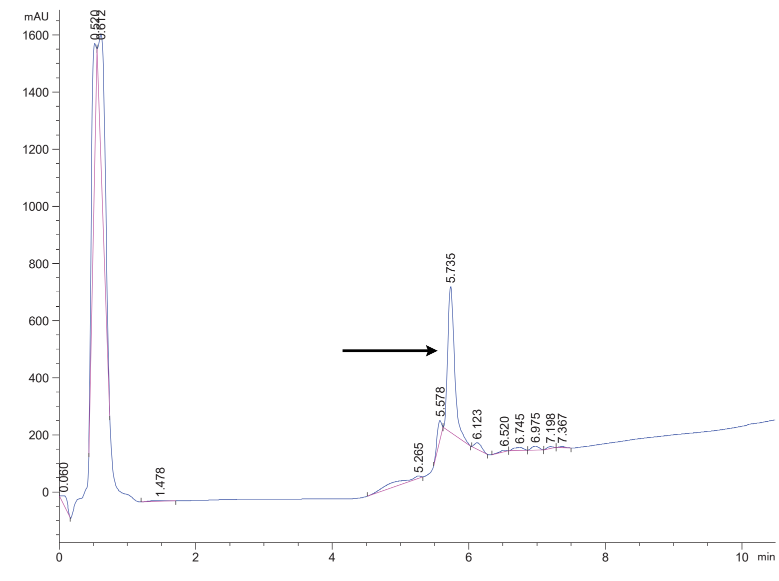

### DPP4 15 Crude

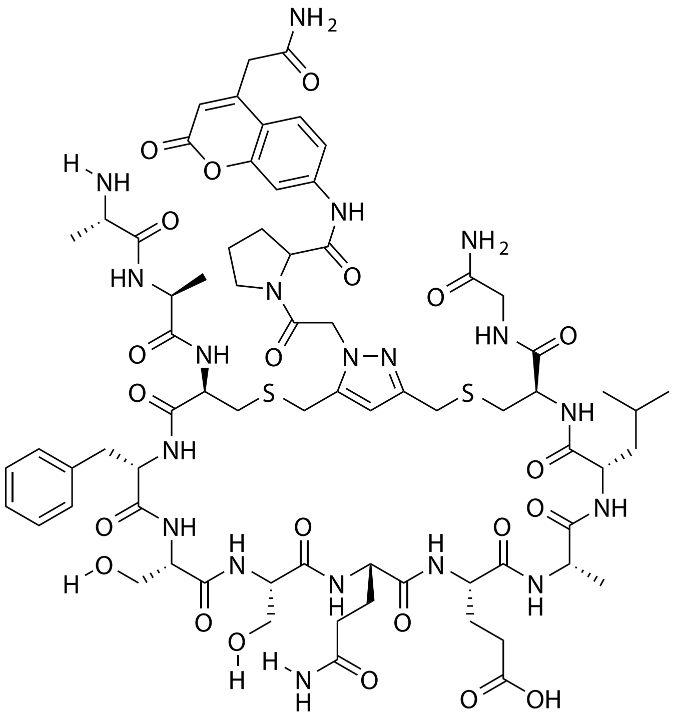

| Probe Name | Sequence | Expected Mass [M+1]$^{+1}$ or [M+2]$^{+2}$ | Found Mass |
|---|---|---|---|
| DPP4 15 | AACFSSQEALCG | 1632.6531 | 1632.9 |

Total Ion Chromatogram

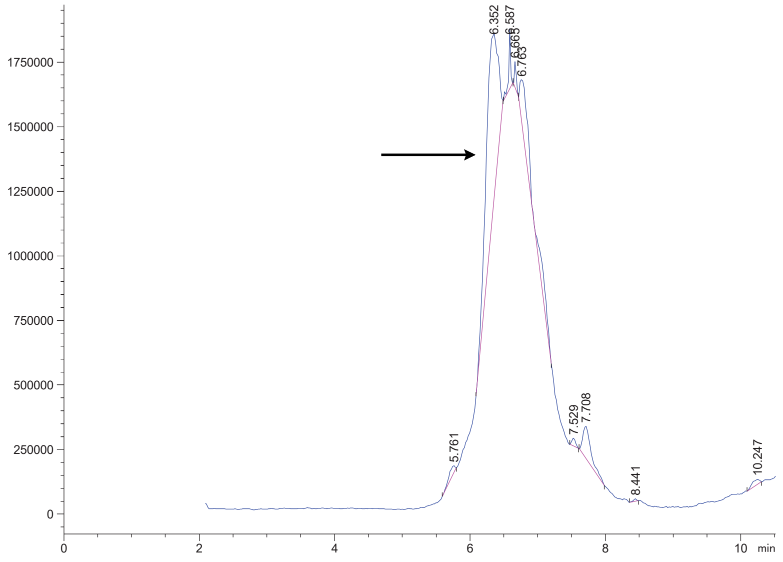

Variable Wavelength Detector: 220 nm

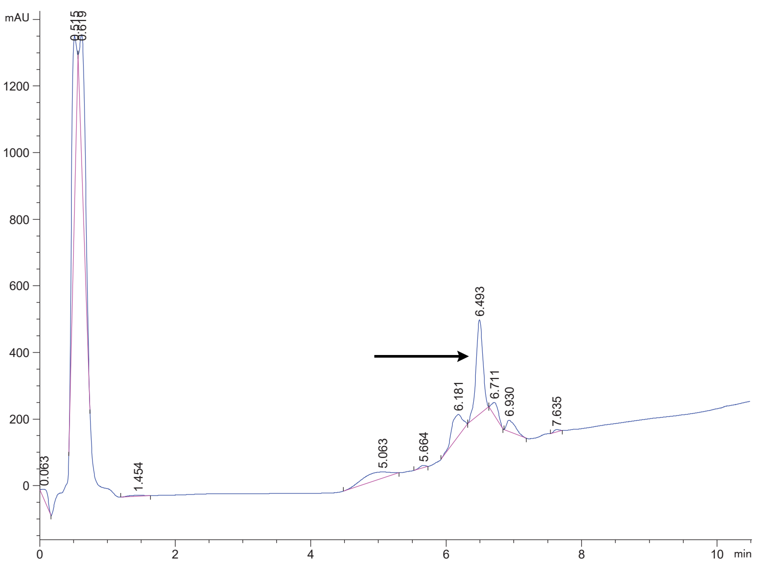

### DPP4 16 Crude

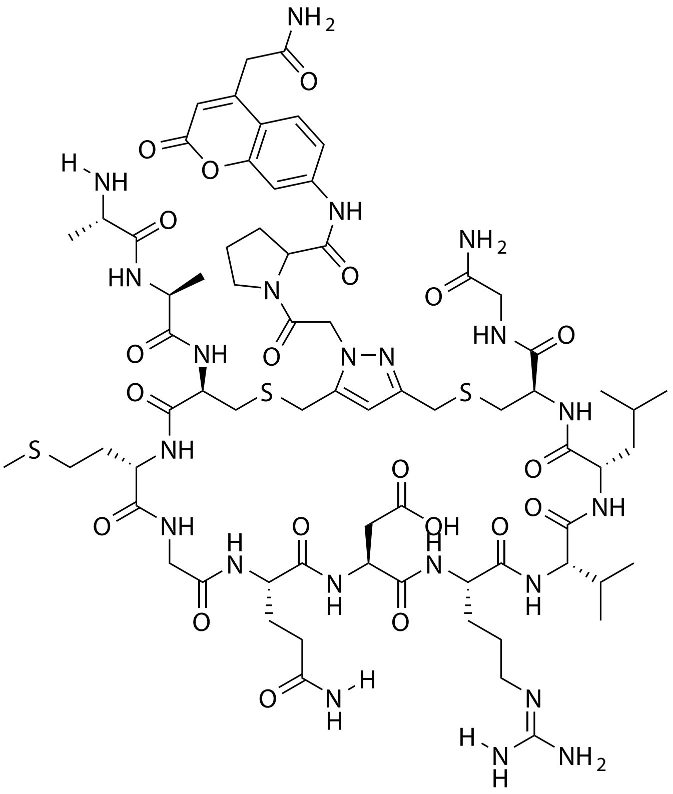

| Probe Name | Sequence | Expected Mass $[M+1]^{+1}$ or $[M+2]^{+2}$ | Found Mass |
|---|---|---|---|
| DPP4 16 | AACMGQDRVLCG | 1669.6993 | 1670.0 |

Total Ion Chromatogram

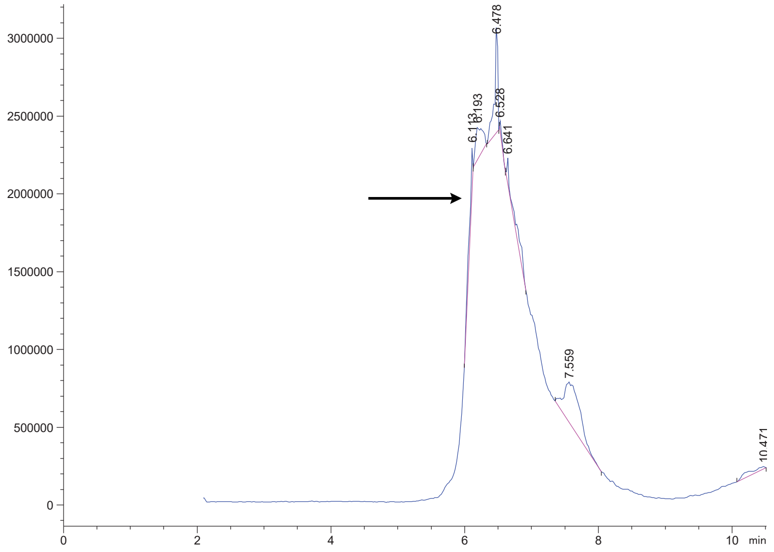

Variable Wavelength Detector: 220 nm

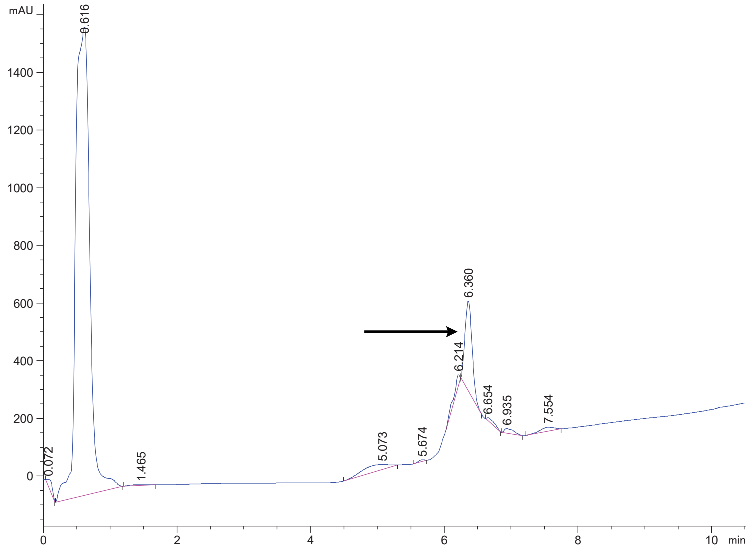

### DPP4 17 Crude

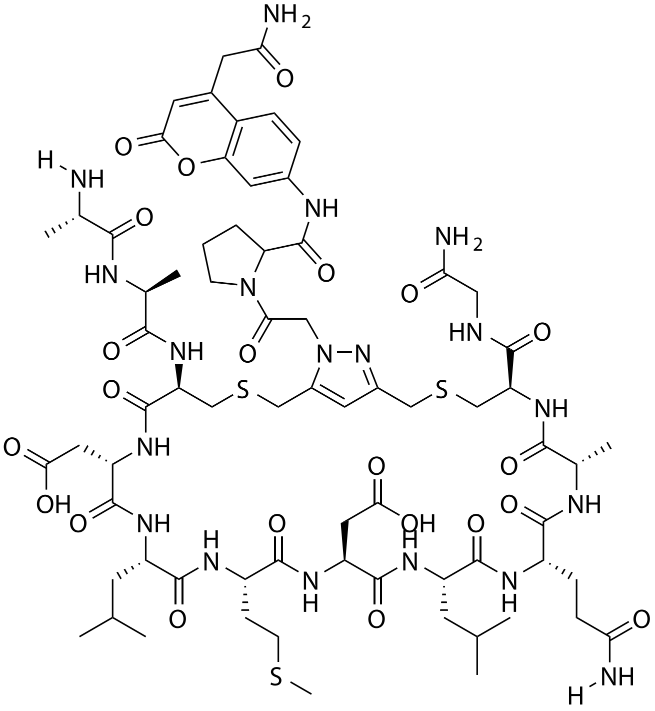

| Probe Name | Sequence | Expected Mass $[M+1]^{+1}$ or $[M+2]^{+2}$ | Found Mass |
|---|---|---|---|
| DPP4 17 | AACDLMDLQACG | 1656.6564 | 1656.9 |

Total Ion Chromatogram

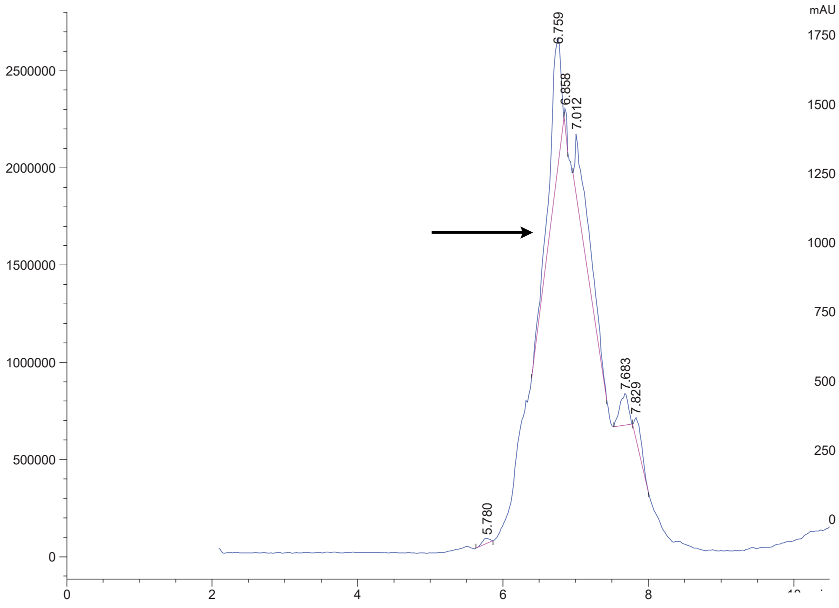


Variable Wavelength Detector: 220 nm

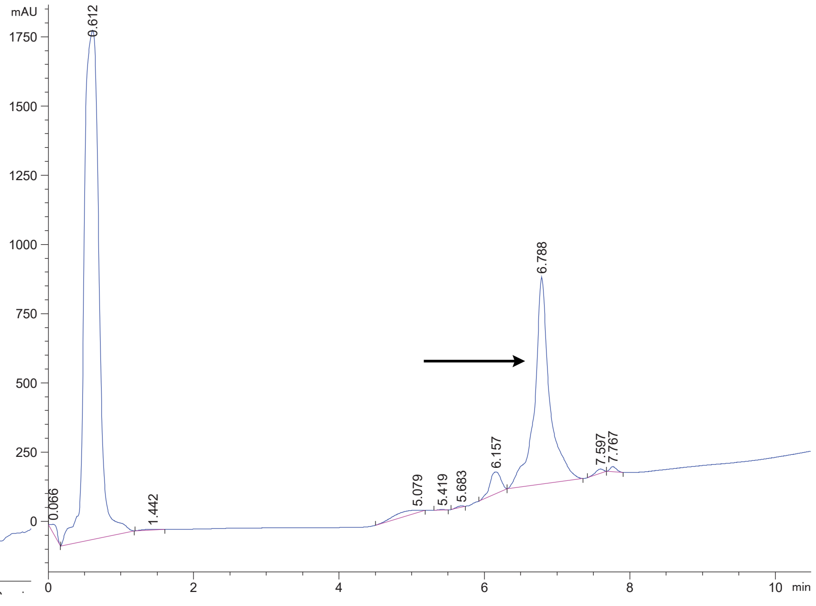

DPP4 18 Crude

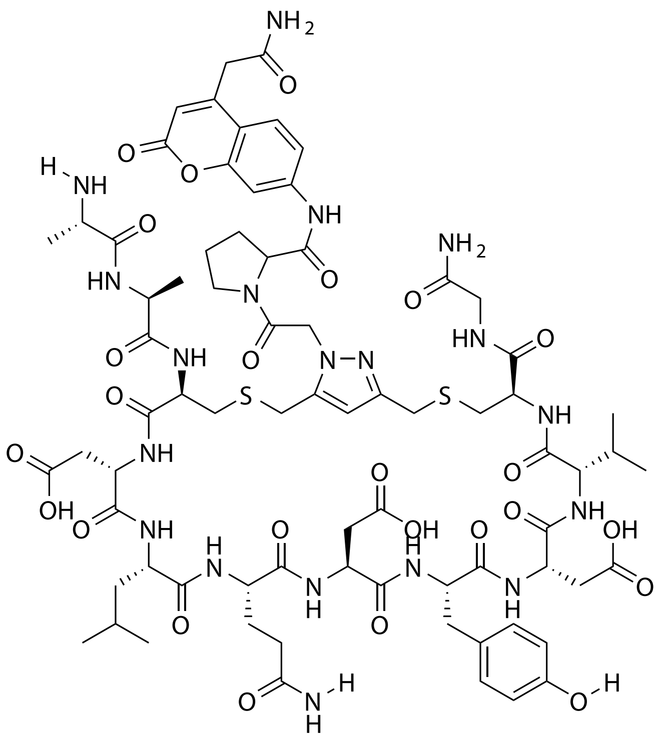

| Probe Name | Sequence | Expected Mass $[M+1]^{+1}$ or $[M+2]^{+2}$ | Found Mass |
|---|---|---|---|
| DPP4 18 | AACDLQDYDVCG | 1718.6534 | 1718.9 |

Total Ion Chromatogram

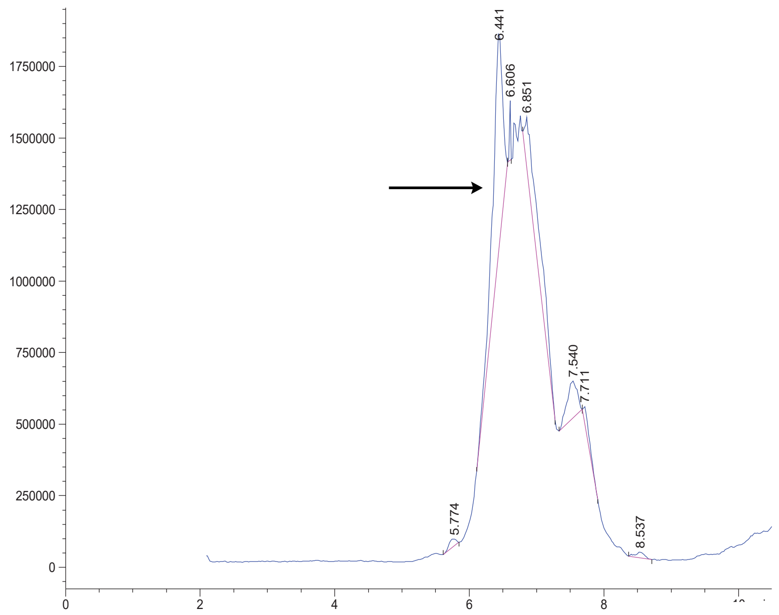

Variable Wavelength Detector: 220 nm

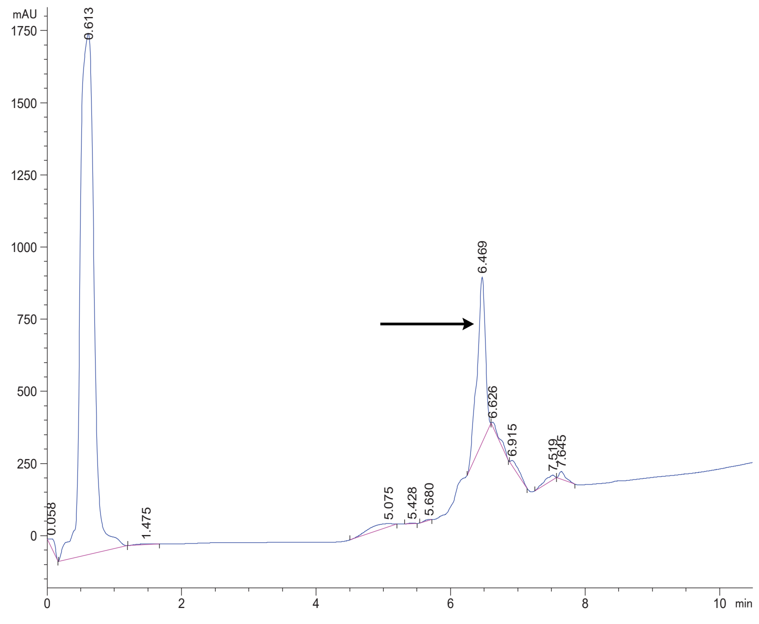

### DPP4 19 Crude

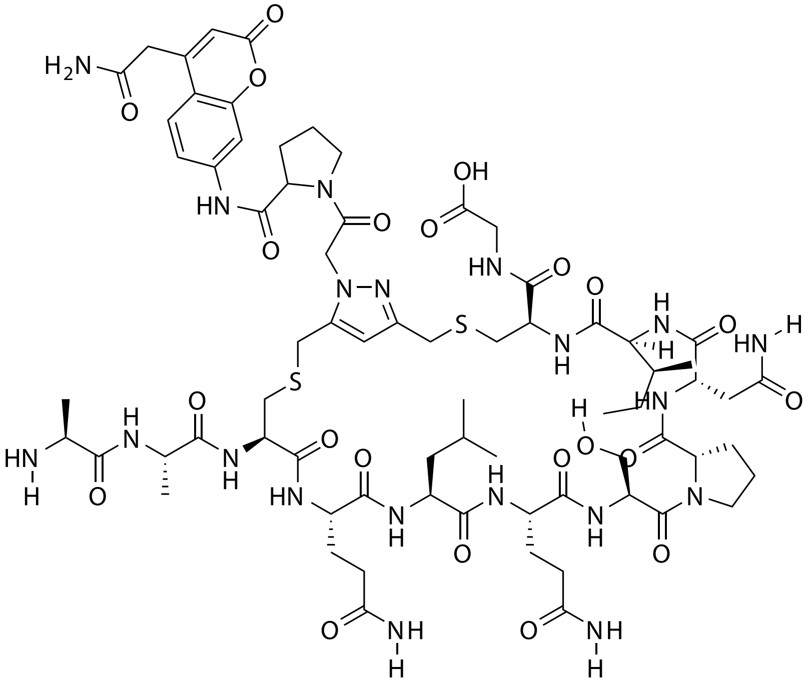

| **Probe Name** | **Sequence** | **Expected Mass $[M+1]^{+1}$ or $[M+2]^{+2}$** | **Found Mass** |
|---|---|---|---|
| DPP4 19 | AACQLQSPNICG | 1651.6953 | 1652.0 |

Total Ion Chromatogram

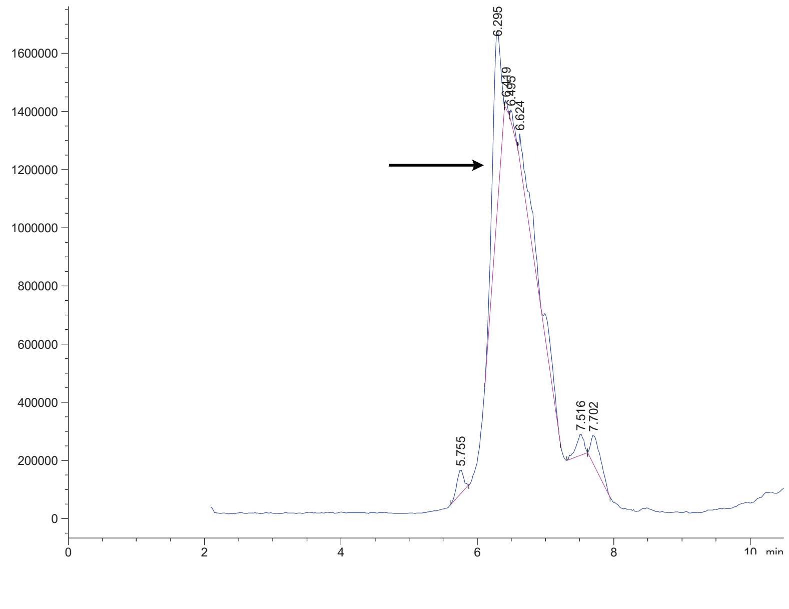


Variable Wavelength Detector: 220 nm

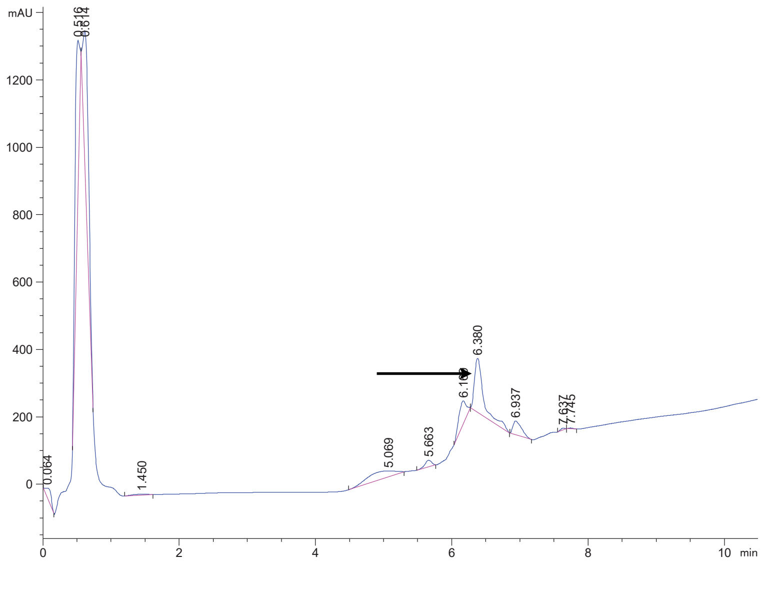

DPP4 20 Crude

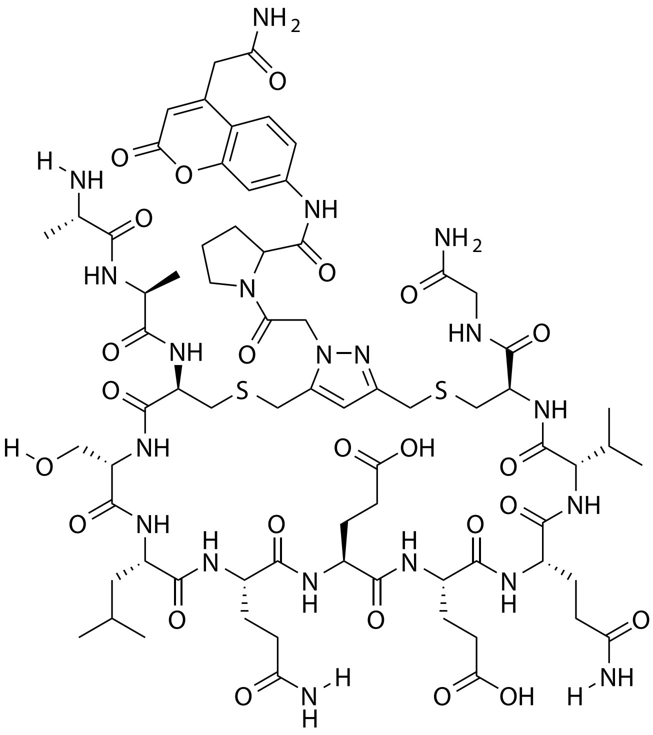


| Probe Name | Sequence | Expected Mass $[M+1]^{+1}$ or $[M+2]^{+2}$ | Found Mass |
|---|---|---|---|
| DPP4 20 | AACSLQEEQVCG | 1683.6851 | 1683.9 |

Total Ion Chromatogram

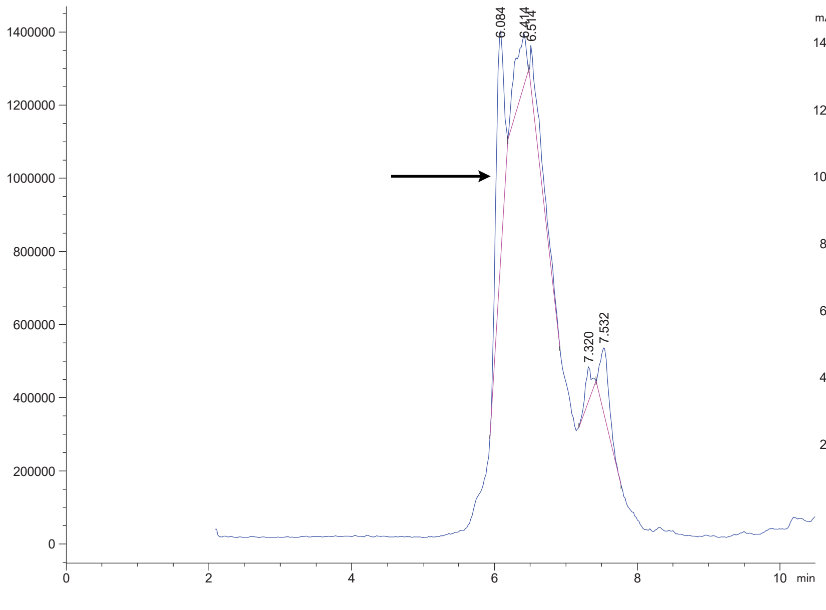


Variable Wavelength Detector: 220 nm

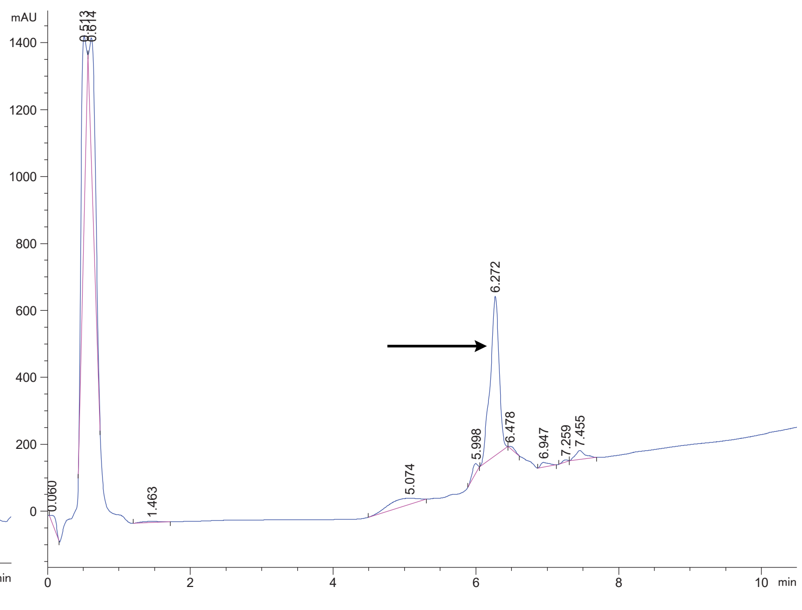

DPP4 21 Crude

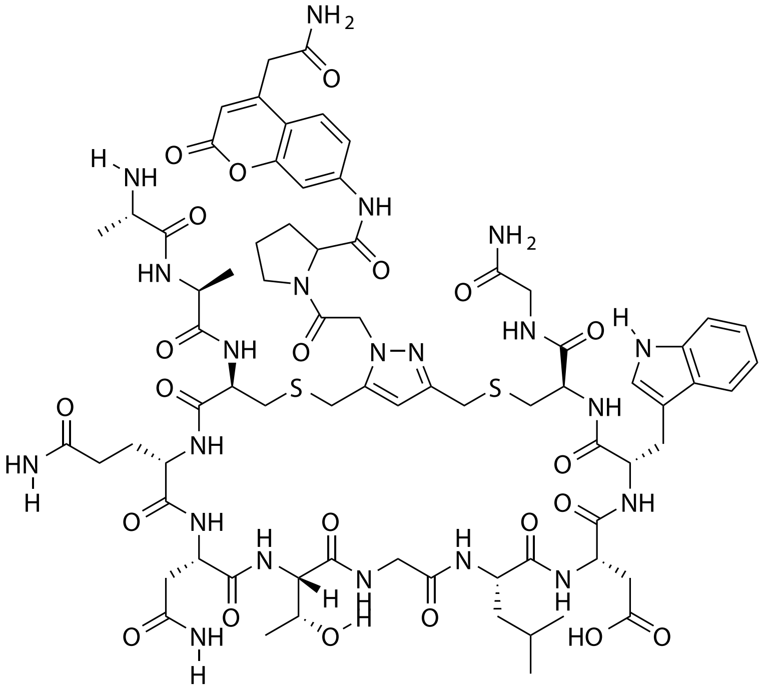

| Probe Name | Sequence | Expected Mass [M+1]$^{+1}$ or [M+2]$^{+2}$ | Found Mass |
|---|---|---|---|
| DPP4 21 | AACQNTGLDWCG | 1684.6592 | 1684.9 |

Total Ion Chromatogram

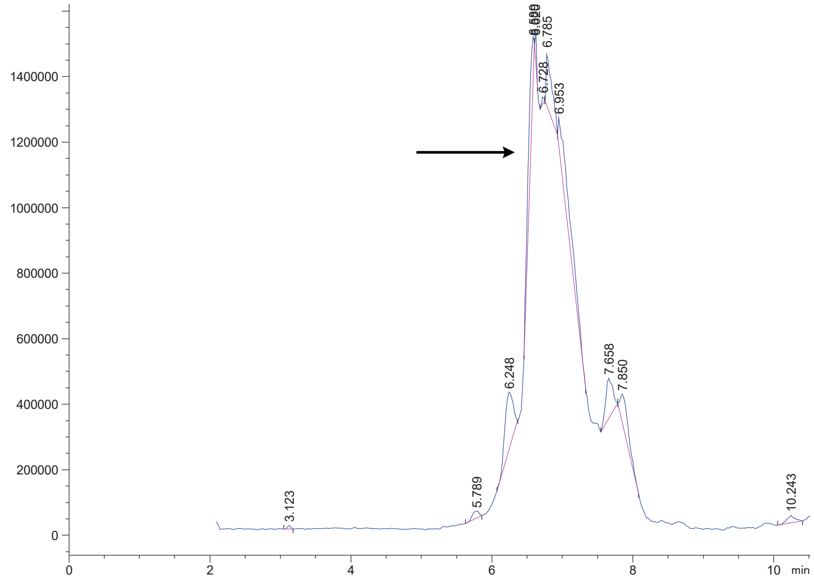

Variable Wavelength Detector: 220 nm

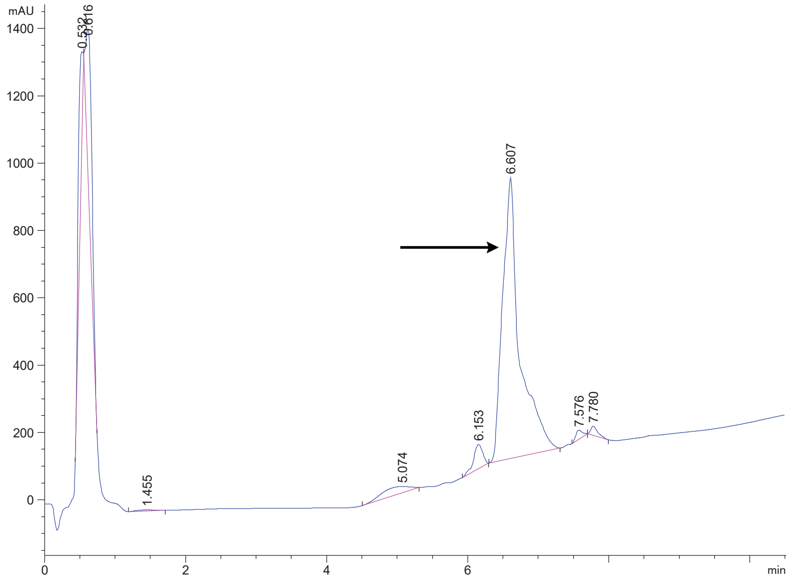

DPP4 22 Crude

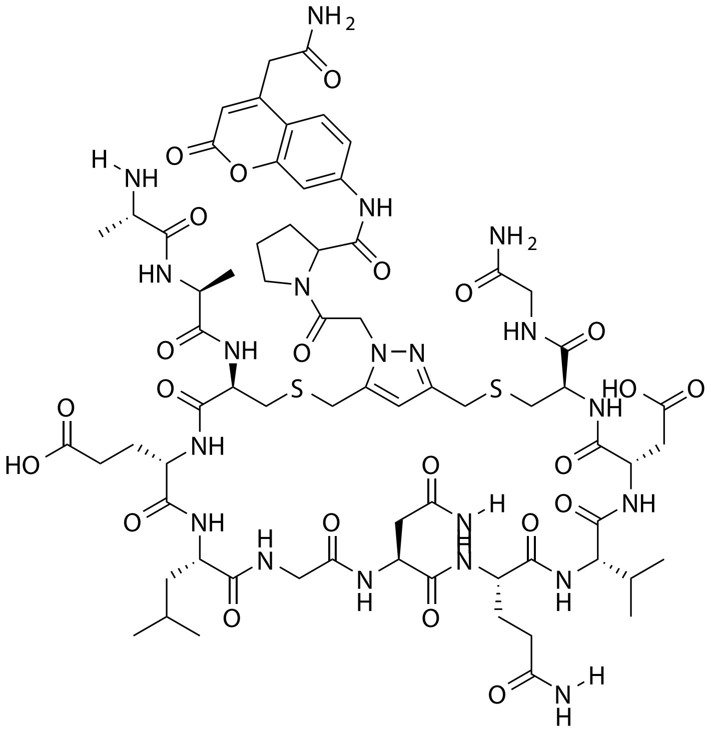

| Probe Name | Sequence | Expected Mass $[M+1]^{+1}$ or $[M+2]^{+2}$ | Found Mass |
|---|---|---|---|
| DPP4 22 | AACELGNQVDCG | 1625.6432 | 1625.9 |

Total Ion Chromatogram

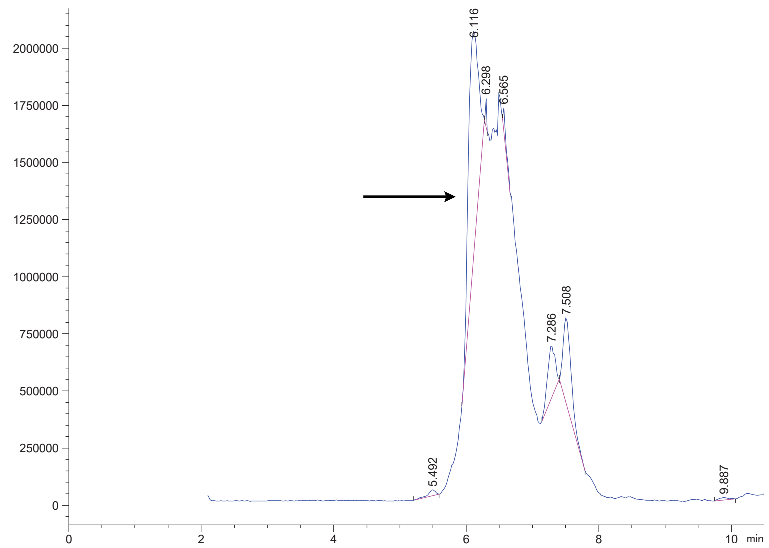


Variable Wavelength Detector: 220 nm

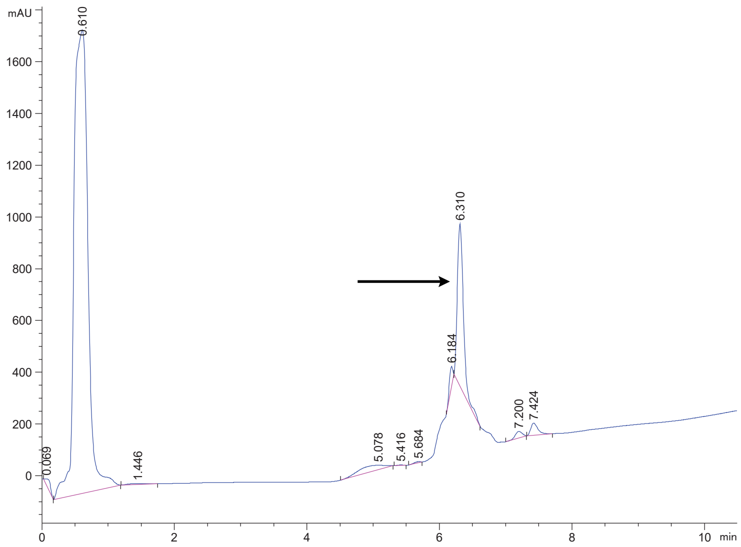

DPP4 23 Crude

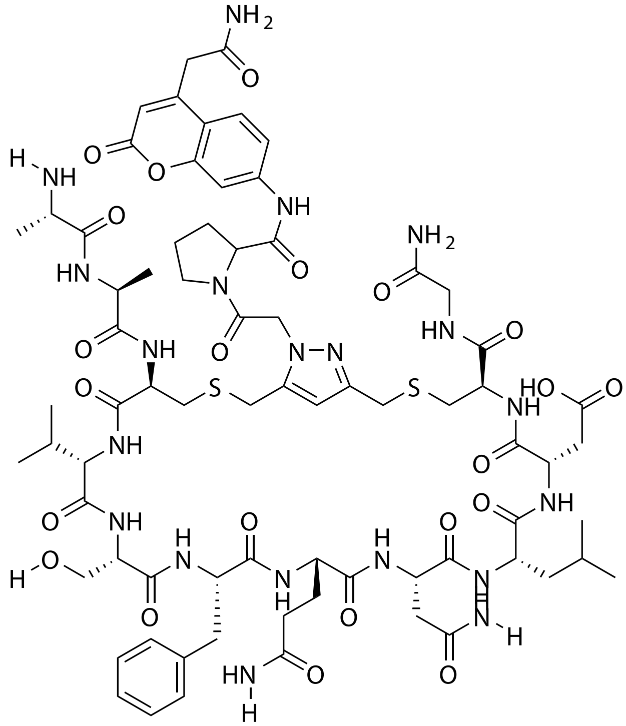

| Probe Name | Sequence | Expected Mass $[M+1]^{+1}$ or $[M+2]^{+2}$ | Found Mass |
|---|---|---|---|
| DPP4 23 | AACVSFQNLDCG | 837.3437 $[M+2]^{+2}$ | 837.2 $[M+2]^{+2}$ |

Total Ion Chromatogram

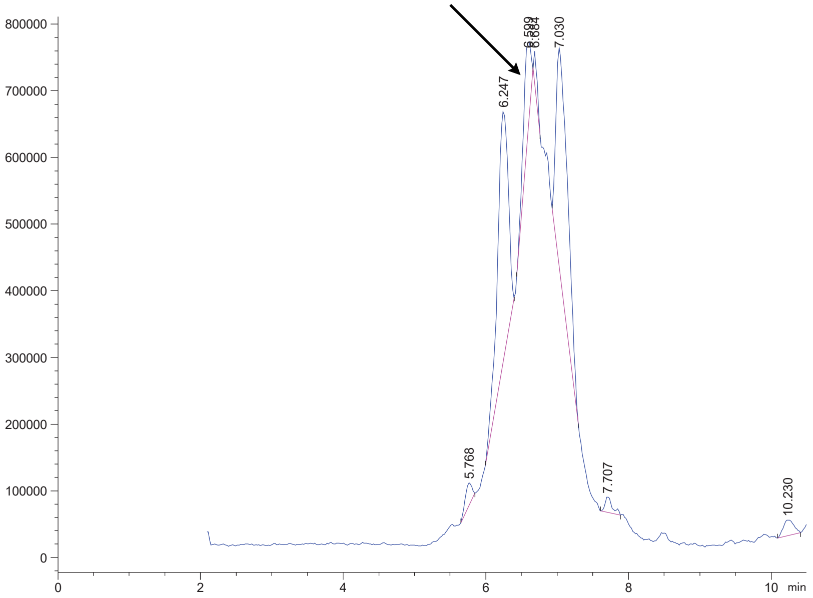

Variable Wavelength Detector: 220 nm

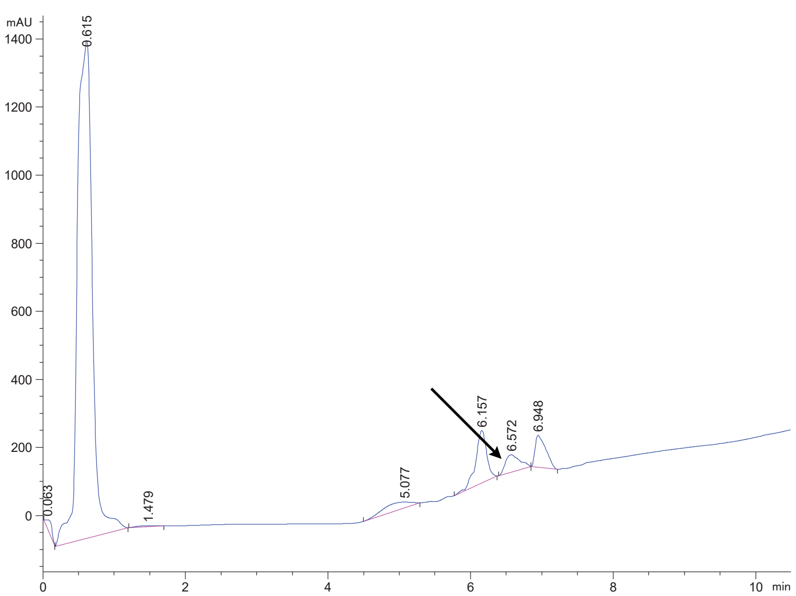

### DPP4 24 Crude

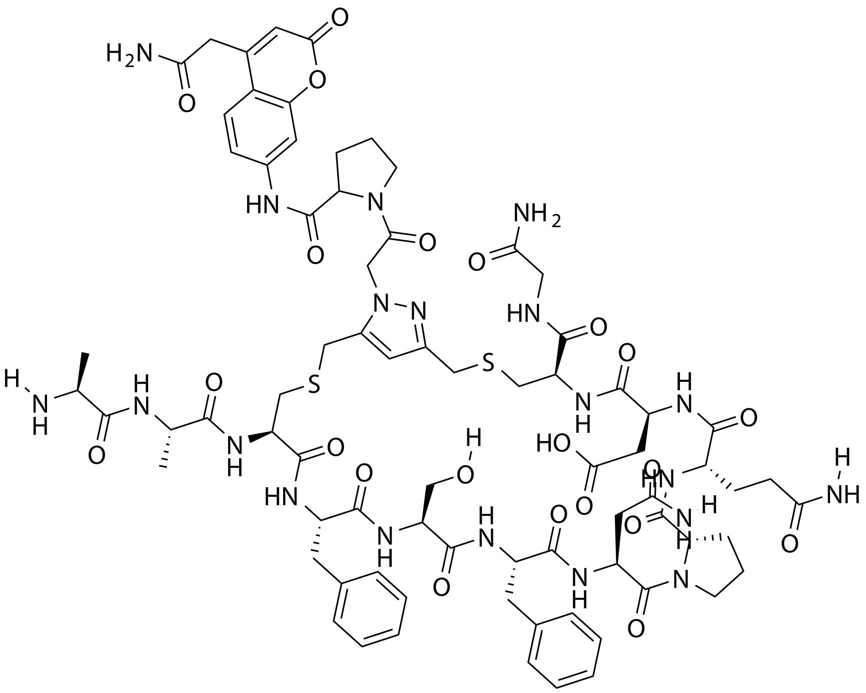

| Probe Name | Sequence | Expected Mass $[M+1]^{+1}$ or $[M+2]^{+2}$ | Found Mass |
|---|---|---|---|
| DPP4 24 | AACFSFNPQDCG | 1705.6483 | 1705.9 |

Total Ion Chromatogram

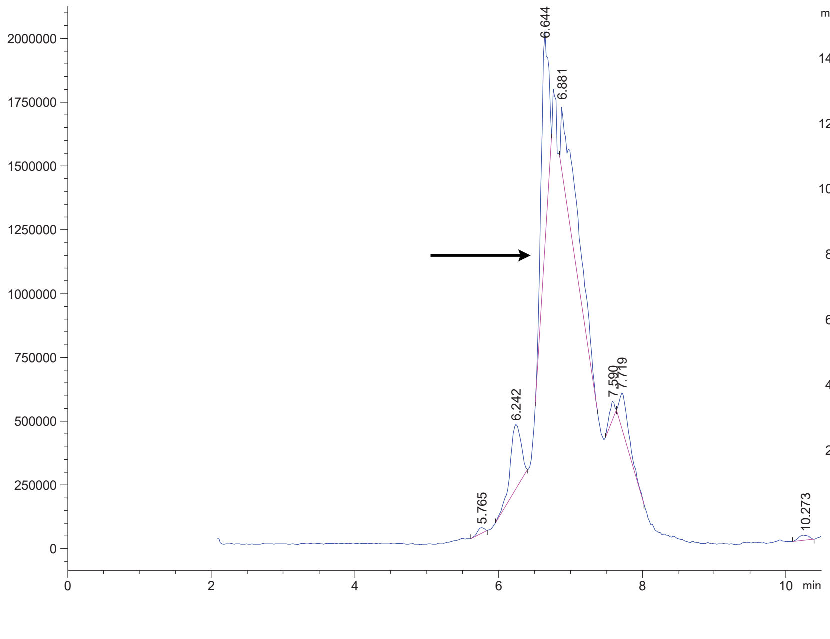


Variable Wavelength Detector: 220 nm

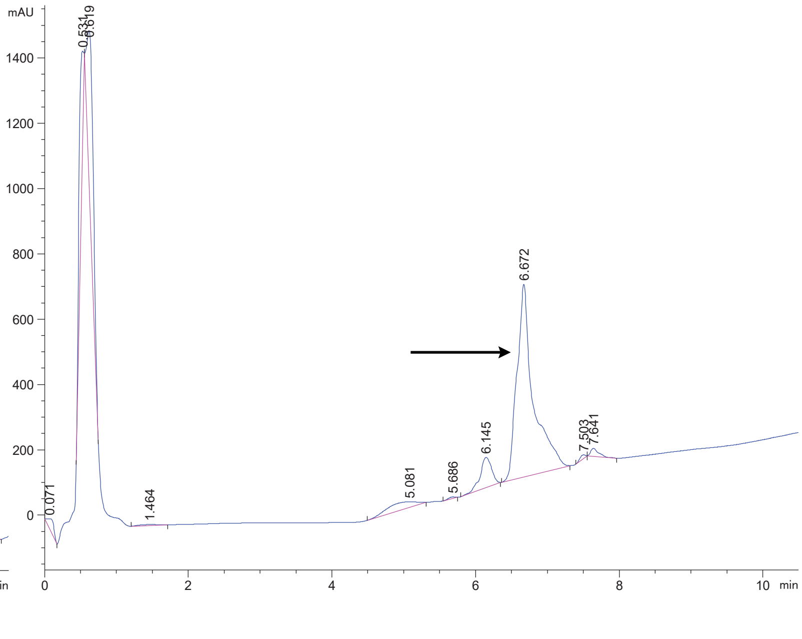

### DPP4 25 Crude

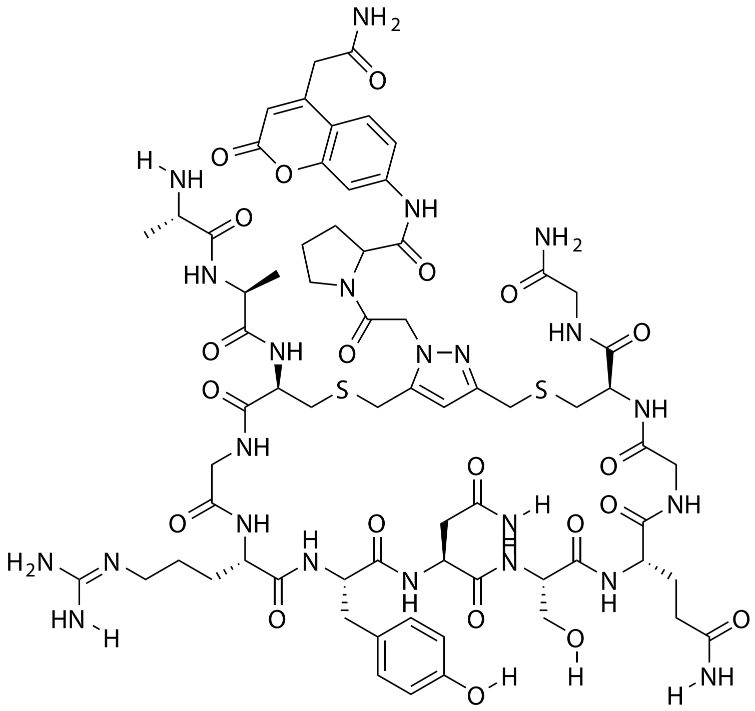


| Probe Name | Sequence | Expected Mass [M+1]$^{+1}$ or [M+2]$^{+2}$ | Found Mass |
|---|---|---|---|
| DPP4 25 | AACGRYNSQGCG | 1632.6391 | 1632.9 |

Total Ion Chromatogram

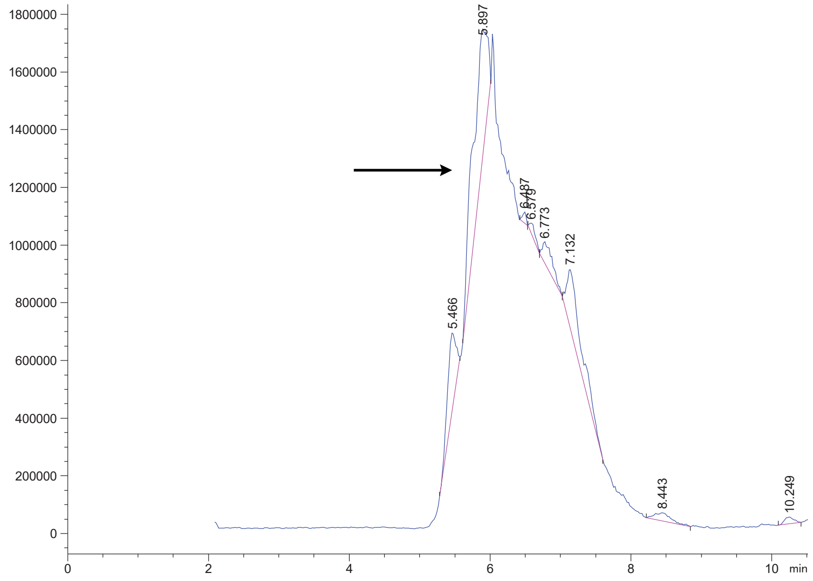


Variable Wavelength Detector: 220 nm

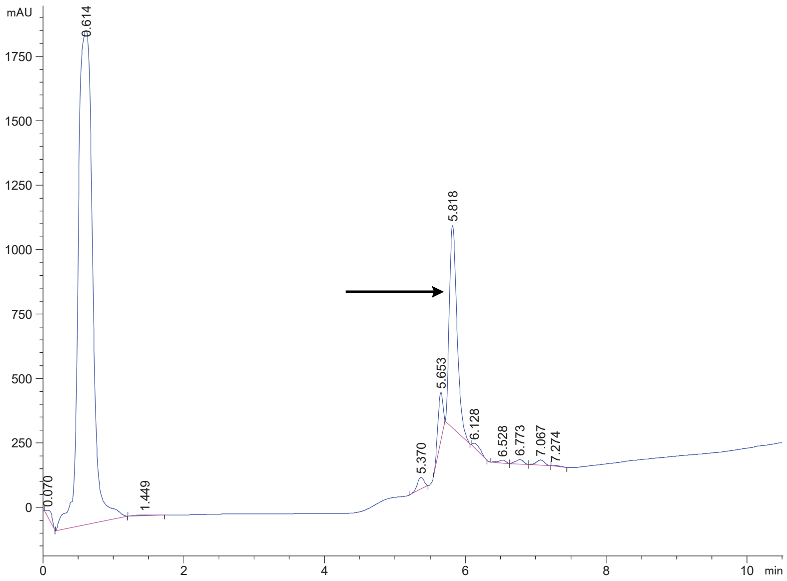

### DPP4 26 Crude

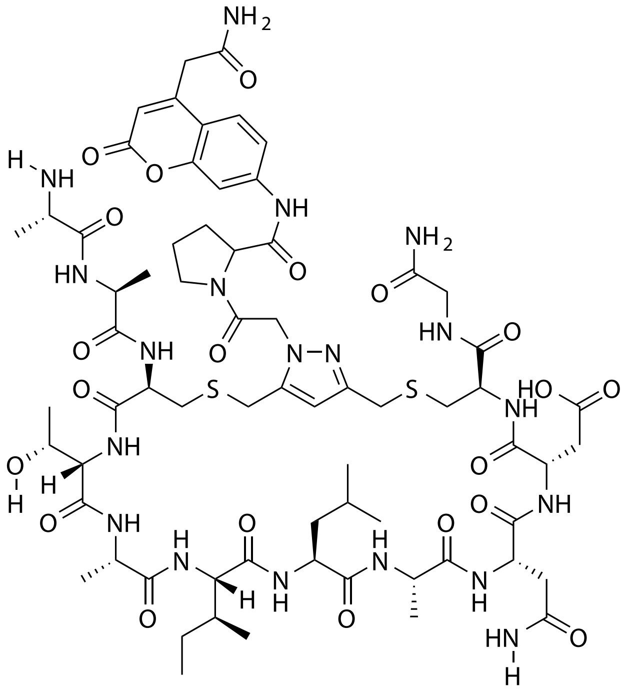

| Probe Name | Sequence | Expected Mass $[M+1]^{+1}$ or $[M+2]^{+2}$ | Found Mass |
|---|---|---|---|
| DPP4 26 | AACTAILANDCG | 1568.6581 | 1568.9 |

Total Ion Chromatogram

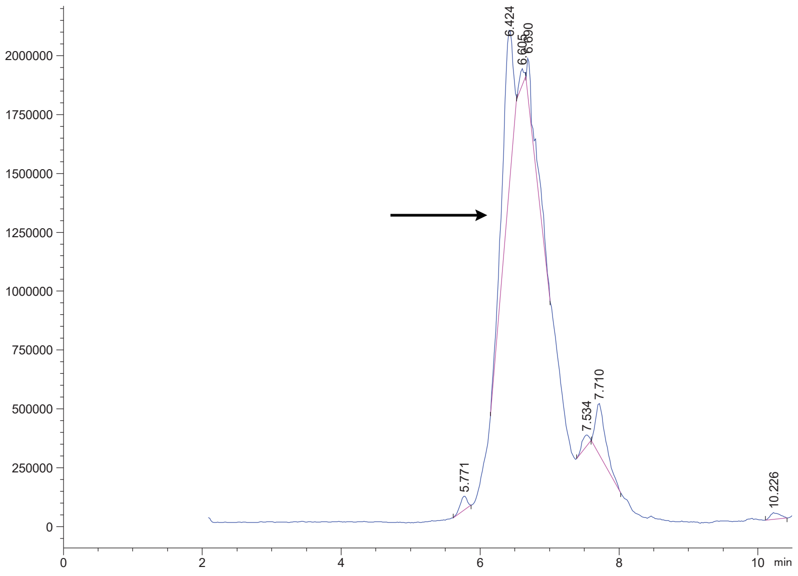


Variable Wavelength Detector: 220 nm

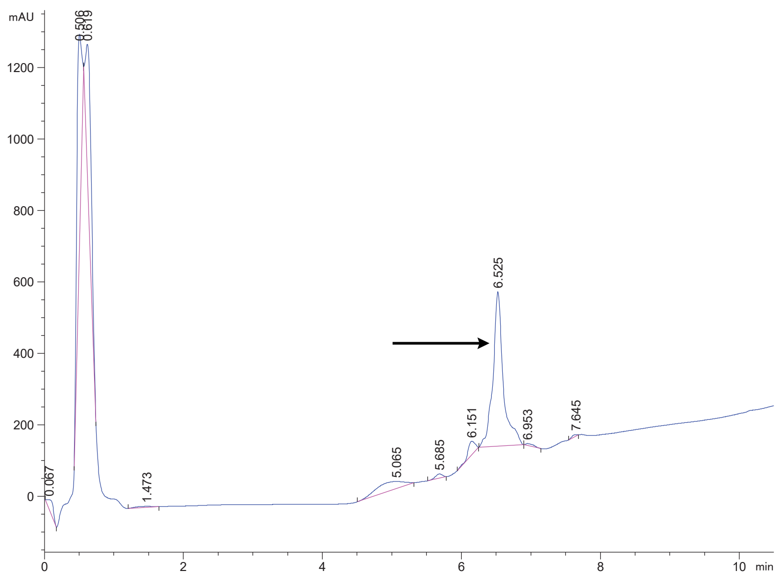

DPP4 27 Crude

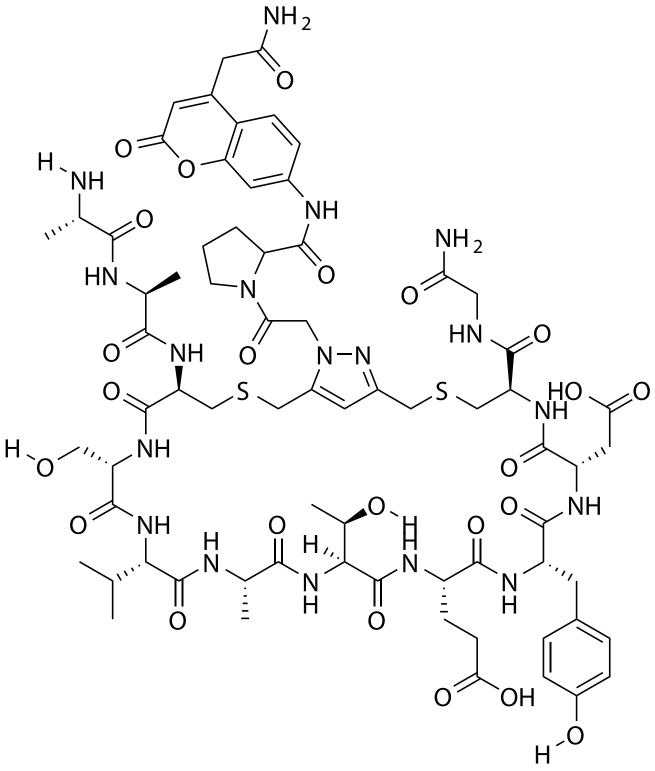

| Probe Name | Sequence | Expected Mass $[M+1]^{+1}$ or $[M+2]^{+2}$ | Found Mass |
|---|---|---|---|
| DPP4 27 | AACSVATEYDCG | 1635.6163 | 1635.9 |

Total Ion Chromatogram

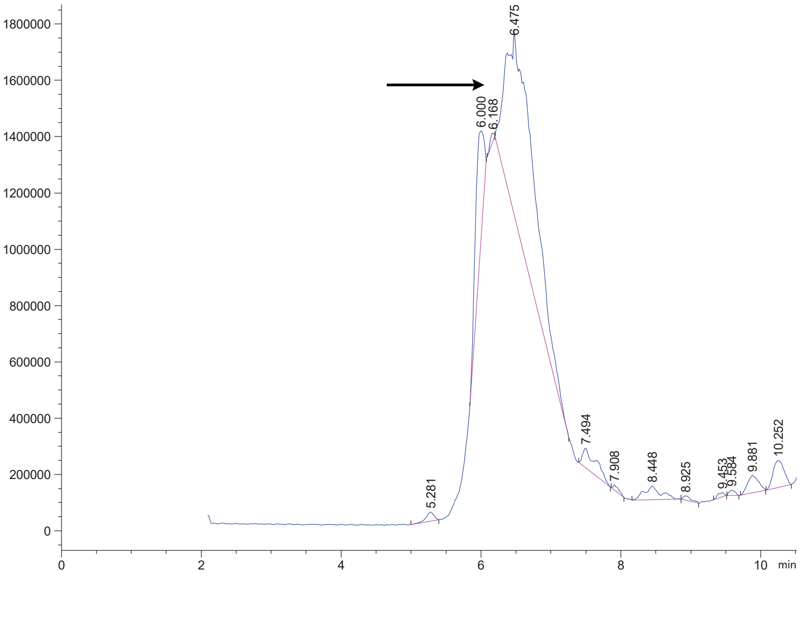


Variable Wavelength Detector: 220 nm

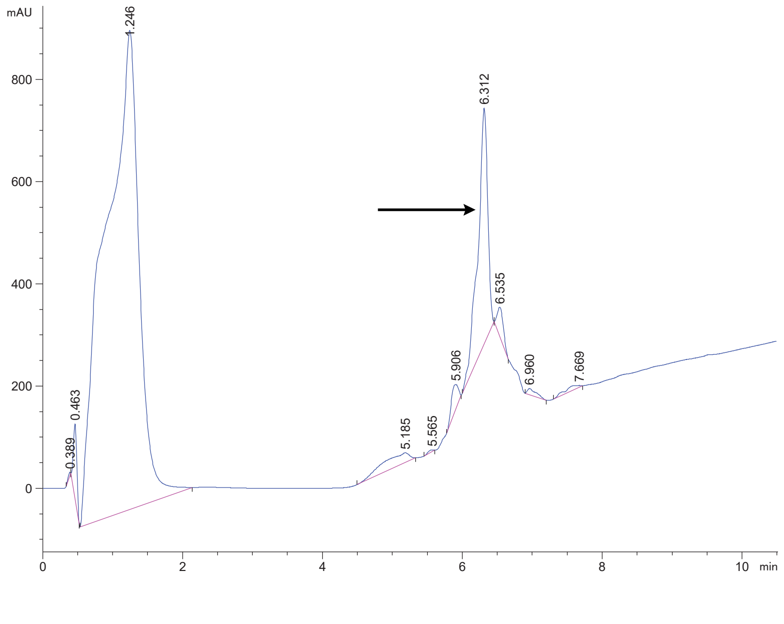

### DPP4 28 Crude

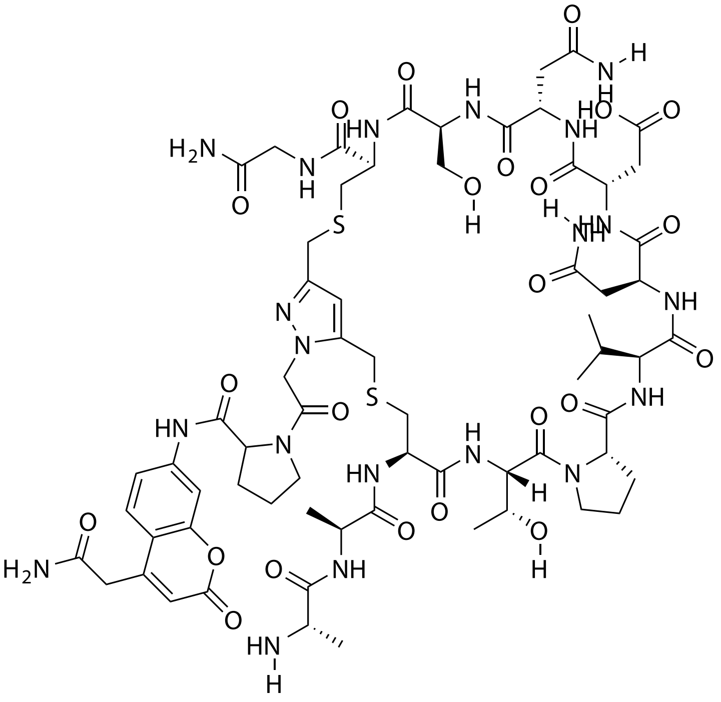

| **Probe Name** | **Sequence** | **Expected Mass $[M+1]^{+1}$ or $[M+2]^{+2}$** | **Found Mass** |
|---|---|---|---|
| DPP4 28 | AACTPVNDNSCG | 1597.6119 | 1597.9 |

Total Ion Chromatogram

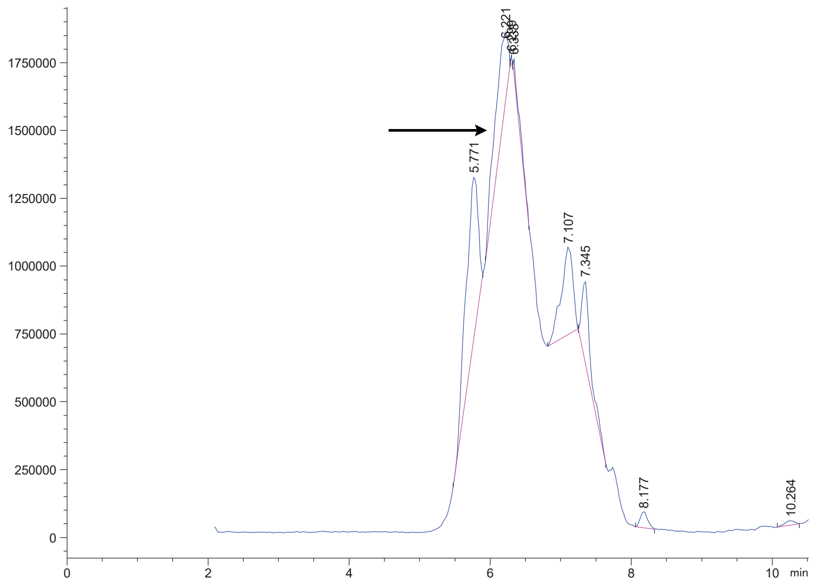


Variable Wavelength Detector: 220 nm

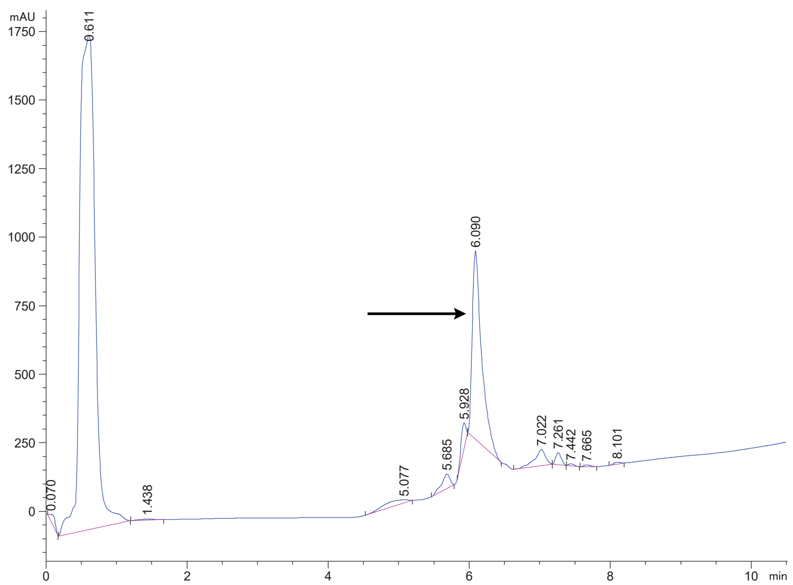

DPP4 29 Crude

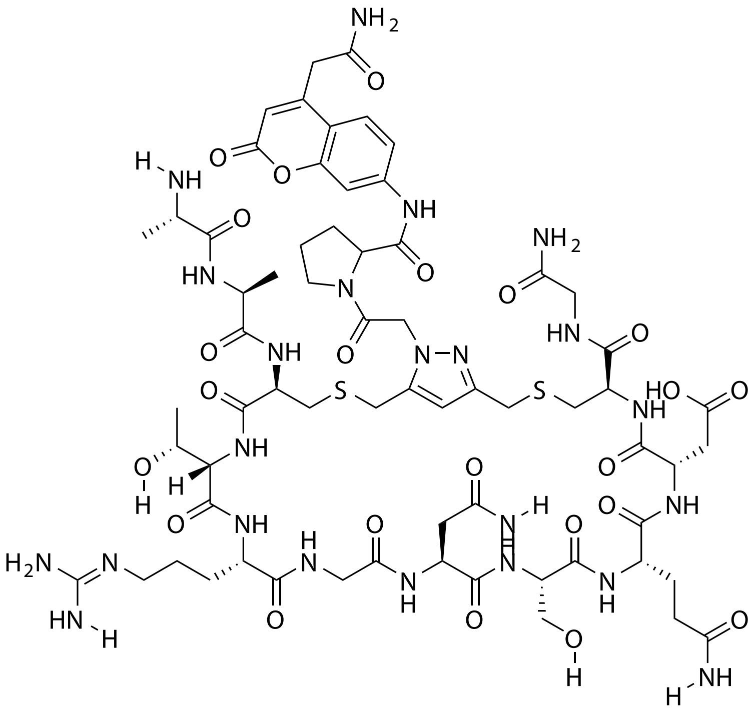

| Probe Name | Sequence | Expected Mass $[M+1]^{+1}$ or $[M+2]^{+2}$ | Found Mass |
|---|---|---|---|
| DPP4 29 | AACTRGNSQDCG | 1628.6290 | 1628.9 |

Total Ion Chromatogram

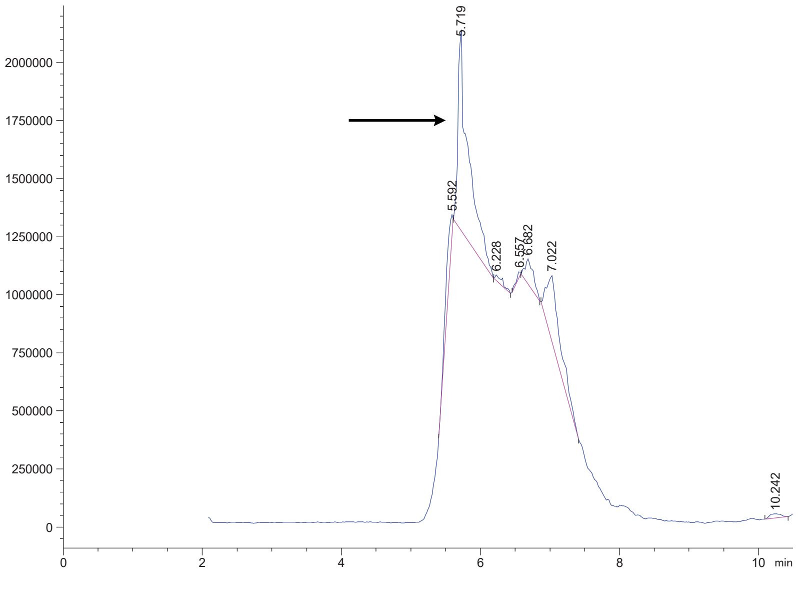

Variable Wavelength Detector: 220 nm

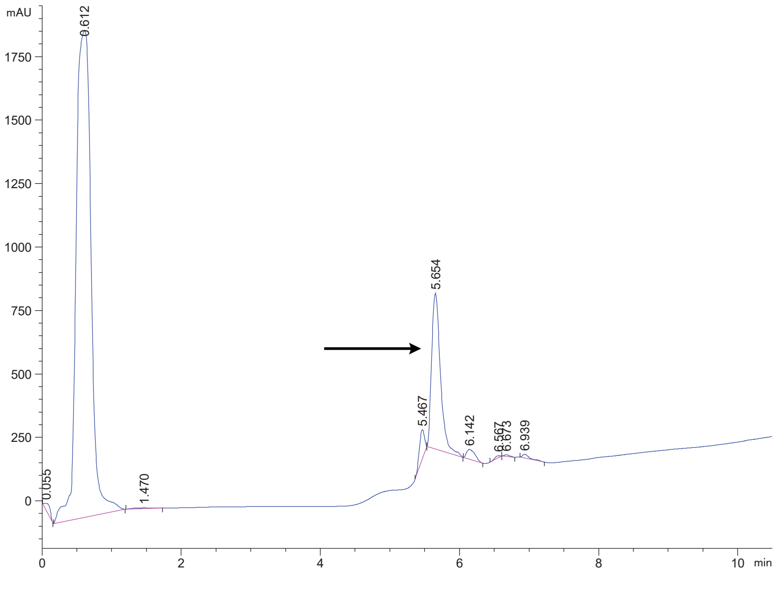

DPP4 30 Crude

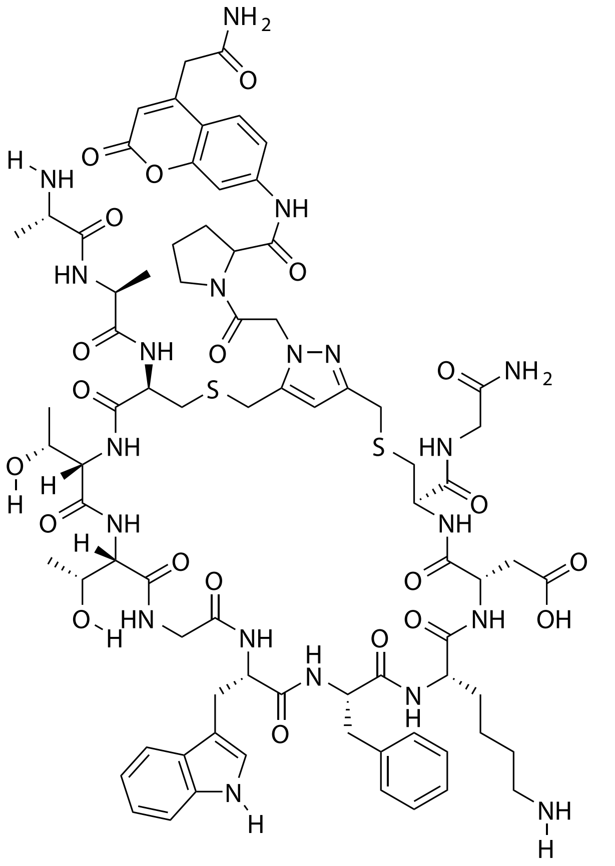

| Probe Name | Sequence | Expected Mass $[M+1]^{+1}$ or $[M+2]^{+2}$ | Found Mass |
|---|---|---|---|
| DPP4 30 | AACTTGWFKDCG | 1705.6847 | 1705.9 |

Total Ion Chromatogram

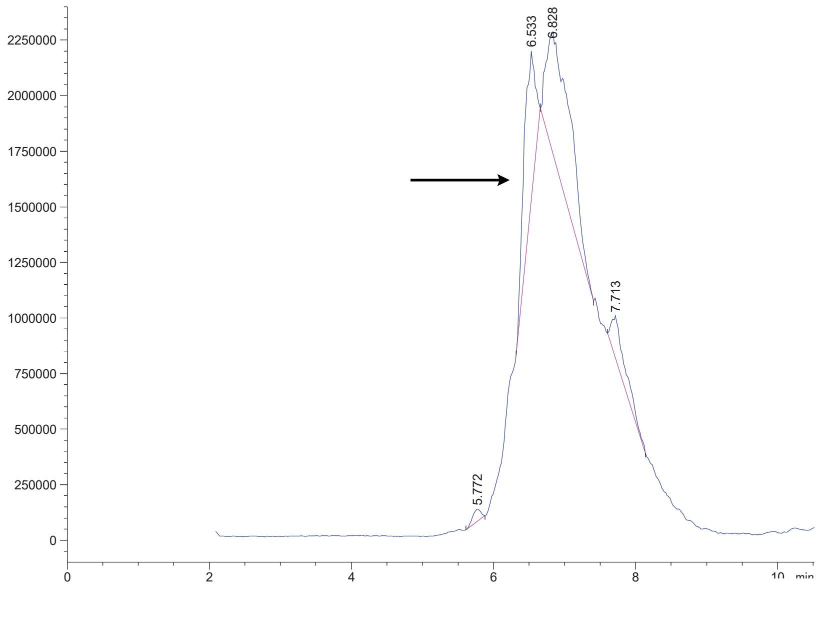


Variable Wavelength Detector: 220 nm

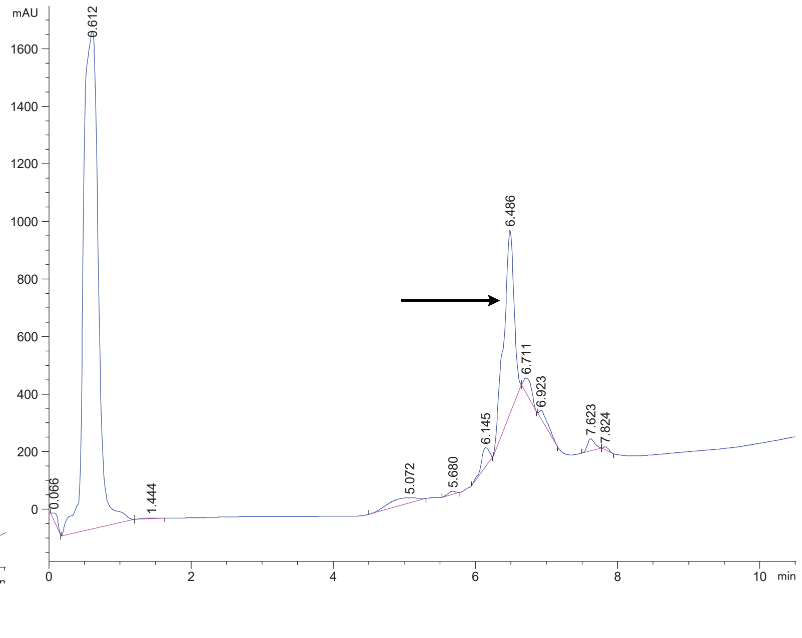

DPP4 31 Crude

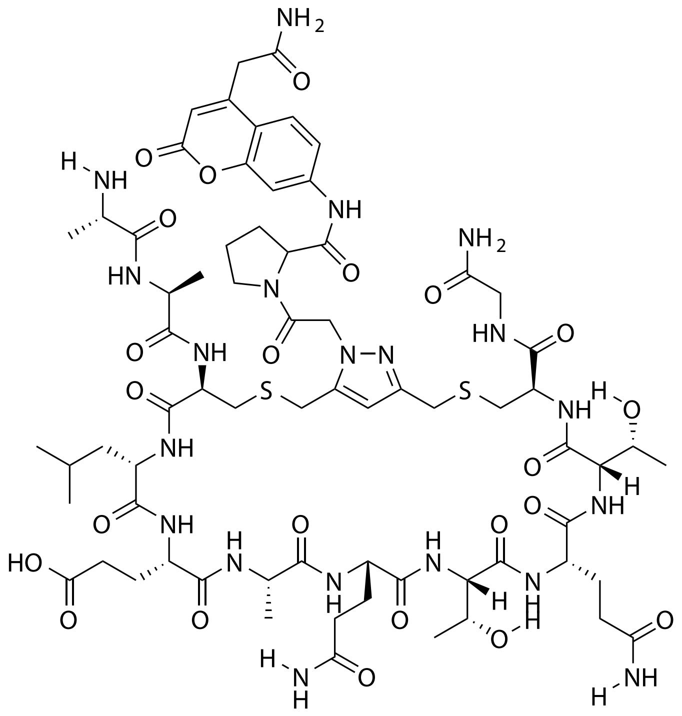


| Probe Name | Sequence | Expected Mass [M+1]+1 or [M+2]+2 | Found Mass |
|---|---|---|---|
| DPP4 31 | AACLEAQTQTCG | 1641.6745 | 1642.0 |

Total Ion Chromatogram

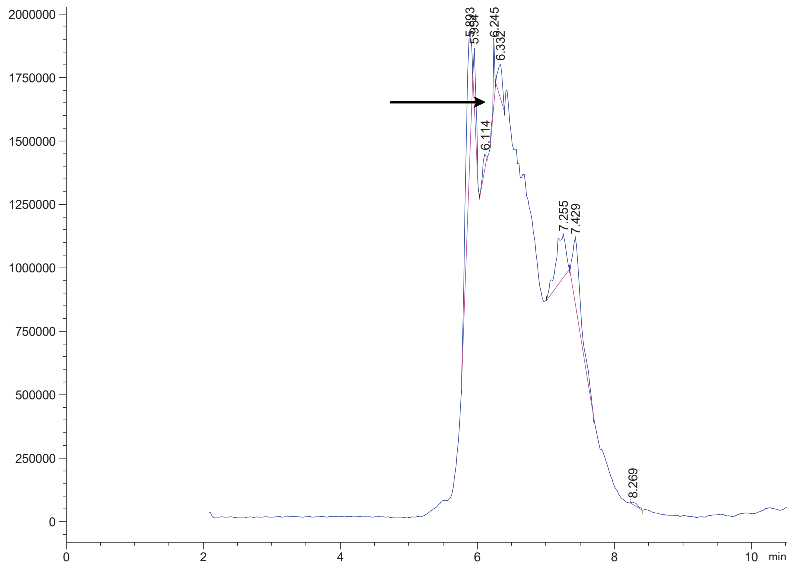


Variable Wavelength Detector: 220 nm

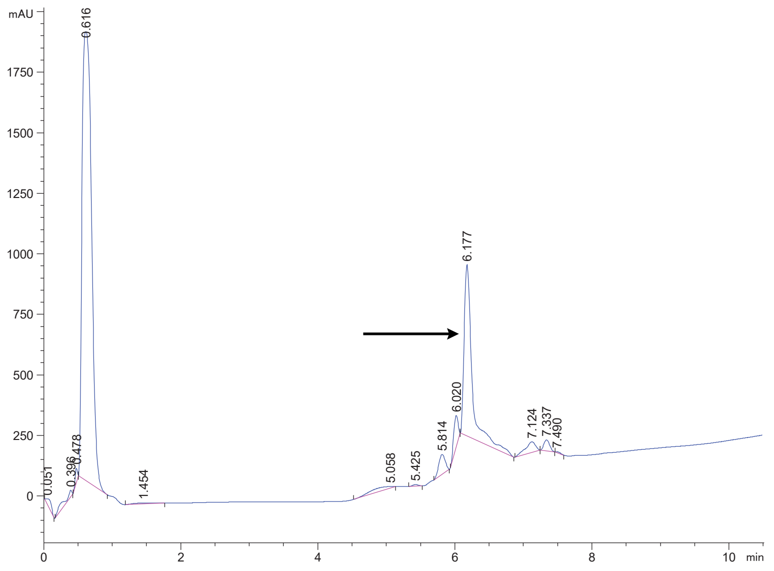

DPP4 32 Crude

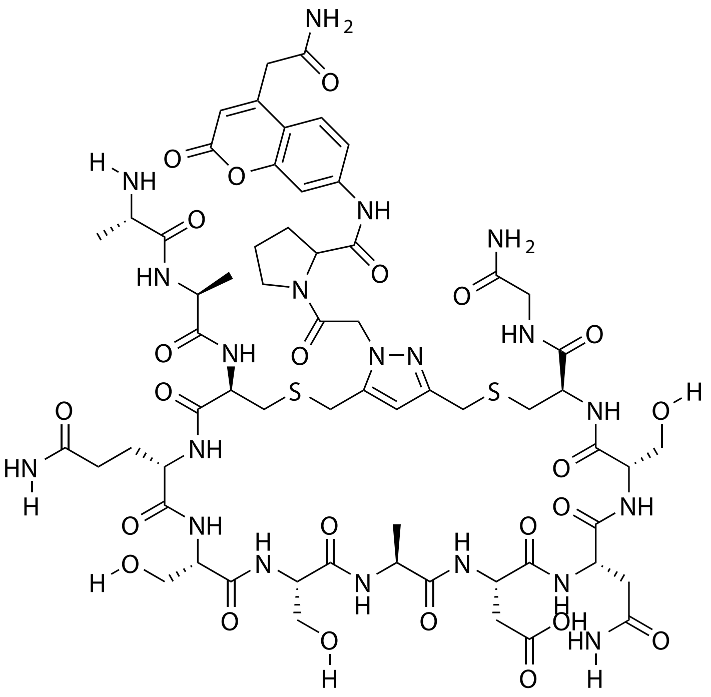

| Probe Name | Sequence | Expected Mass $[M+1]^{+1}$ or $[M+2]^{+2}$ | Found Mass |
|---|---|---|---|
| DPP4 32 | AACQSSADNSCG | 1559.5599 | 1559.8 |

Total Ion Chromatogram

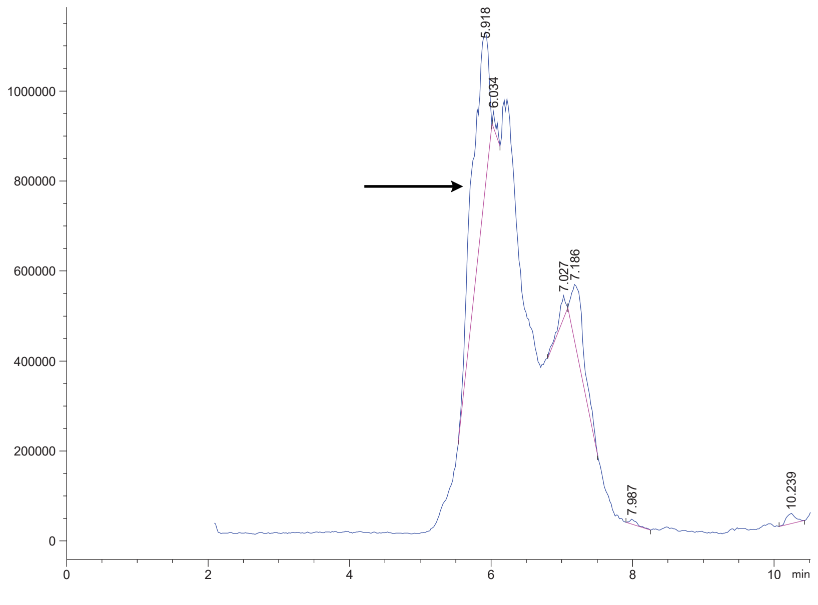

Variable Wavelength Detector: 220 nm

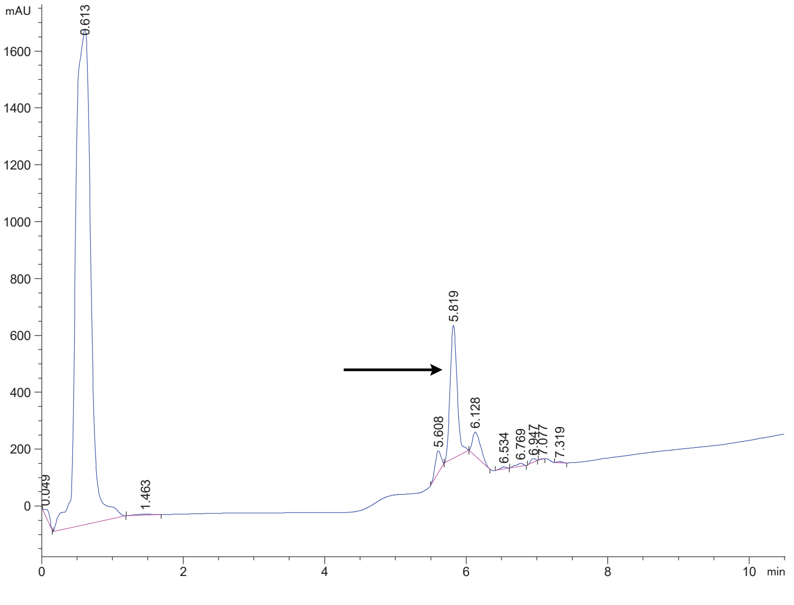

### DPP4 33 Crude

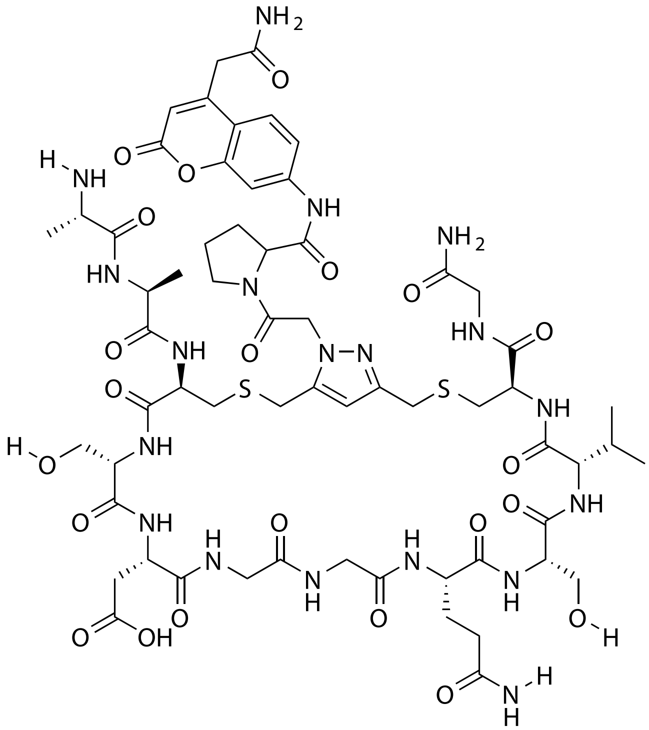

| Probe Name | Sequence | Expected Mass $[M+1]^{+1}$ or $[M+2]^{+2}$ | Found Mass |
|---|---|---|---|
| DPP4 33 | AACSDGGQSVCG | 1500.5592 | 1500.9 |

Total Ion Chromatogram

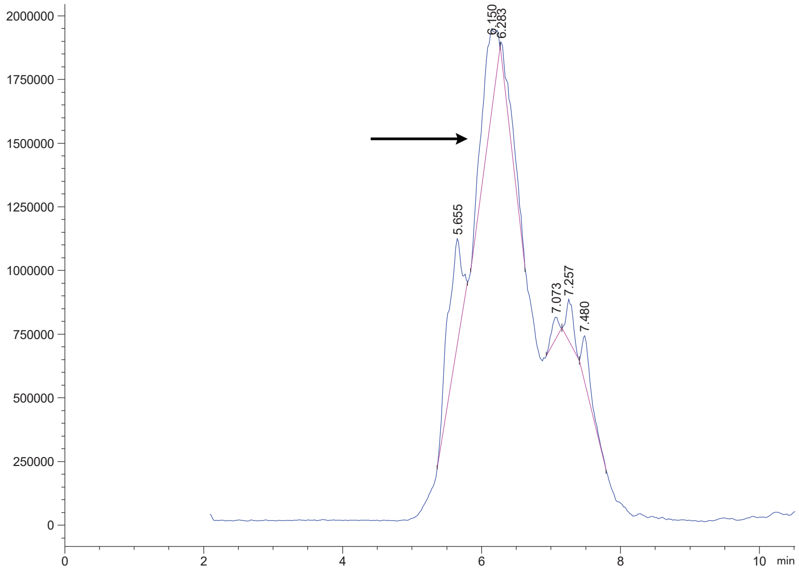


Variable Wavelength Detector: 220 nm

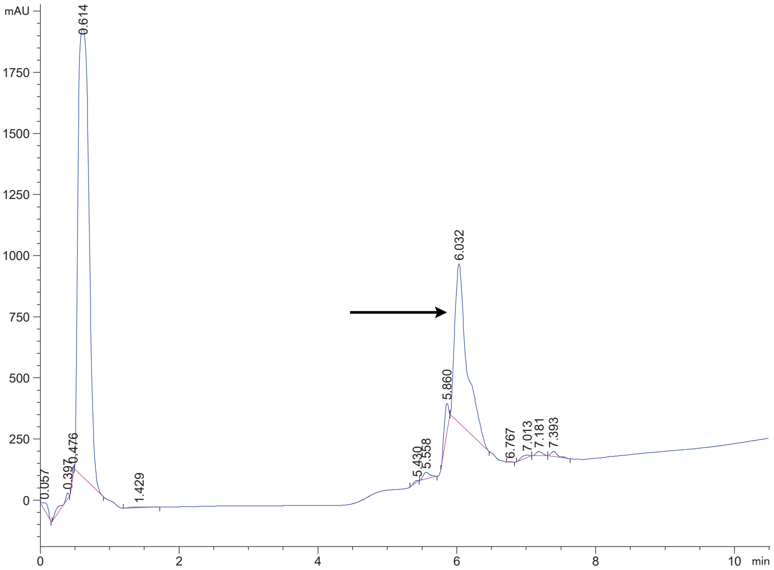

DPP4 34 Crude

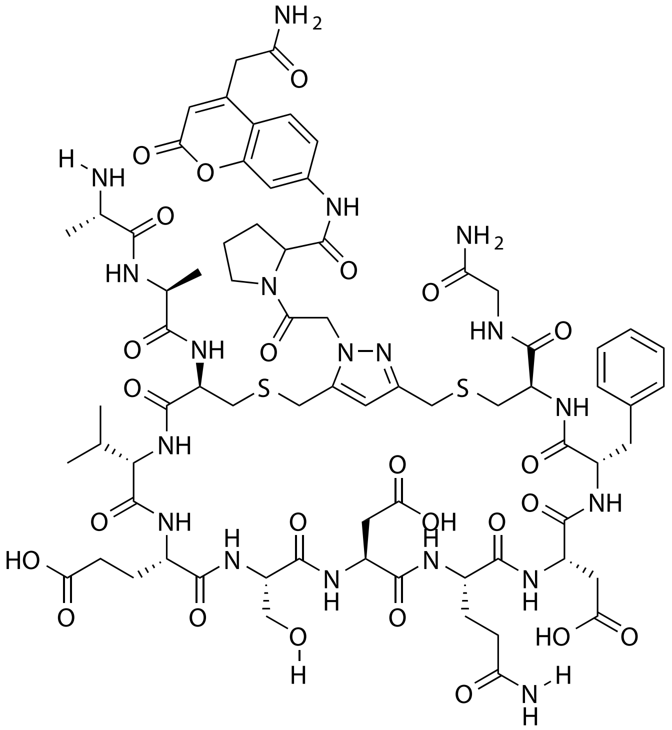


| Probe Name | Sequence | Expected Mass $[M+1]^{+1}$ or $[M+2]^{+2}$ | Found Mass |
|---|---|---|---|
| DPP4 34 | AACVESDQDFCG | 1690.6211 | 1690.9 |

Total Ion Chromatogram

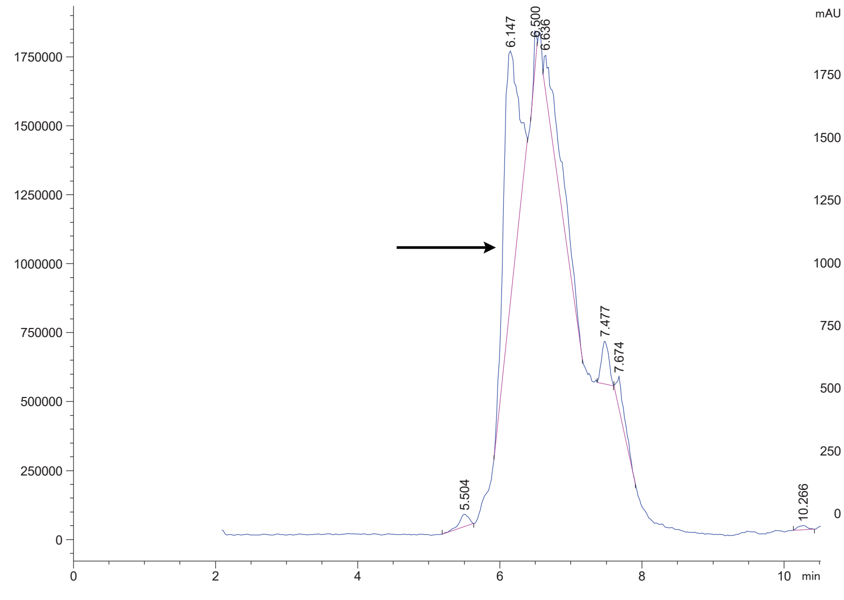


Variable Wavelength Detector: 220 nm

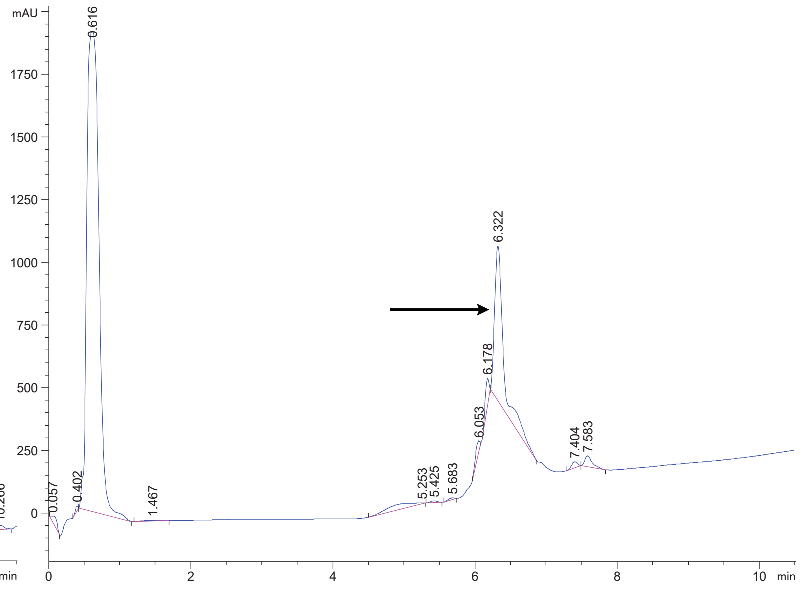

DPP4 35 Crude

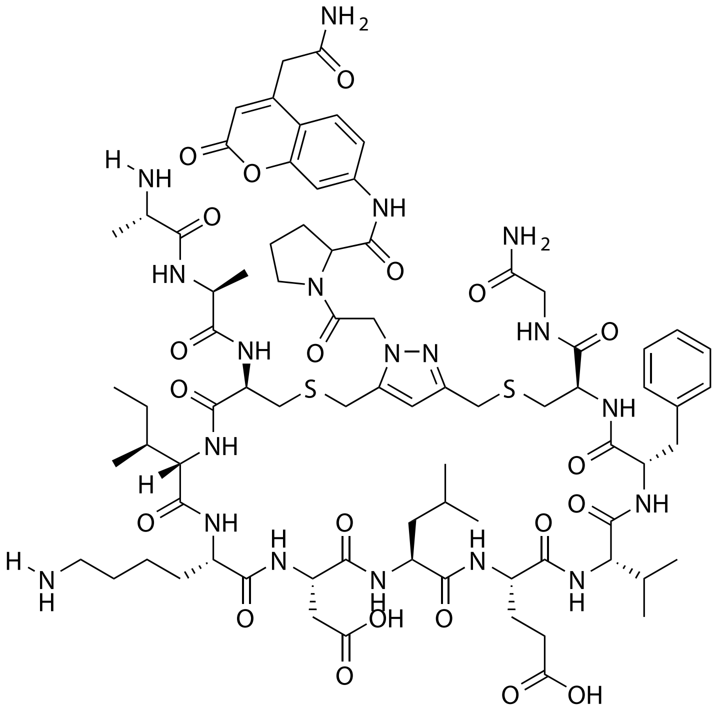

| Probe Name | Sequence | Expected Mass $[M+1]^{+1}$ or $[M+2]^{+2}$ | Found Mass |
|---|---|---|---|
| DPP4 35 | AACIKDLEVFCG | 1714.7677 | 1715.0 |

Total Ion Chromatogram

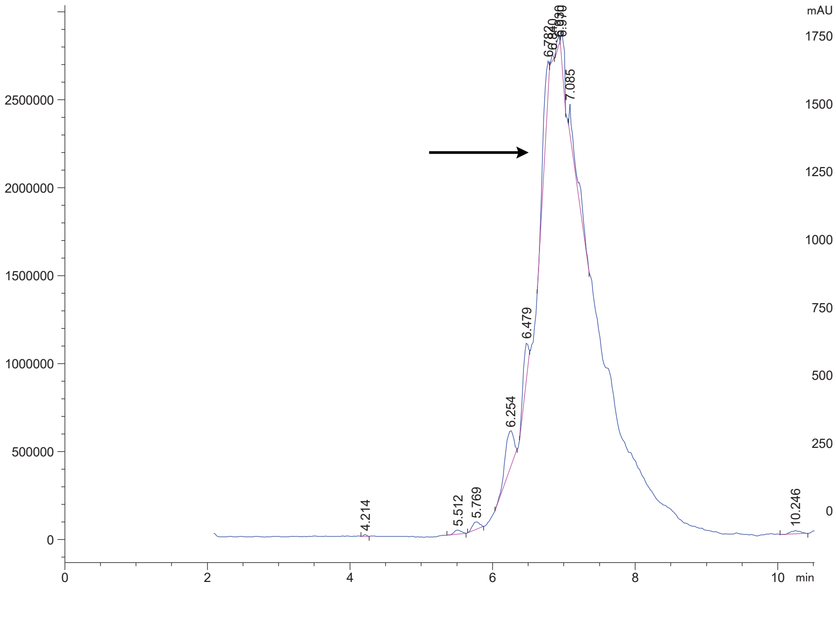

Variable Wavelength Detector: 220 nm

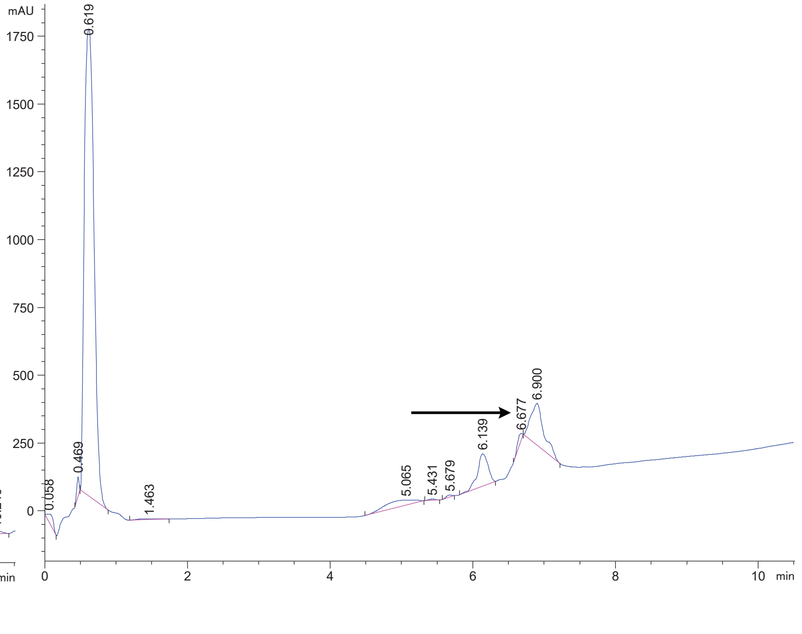

### DPP4 36 Crude

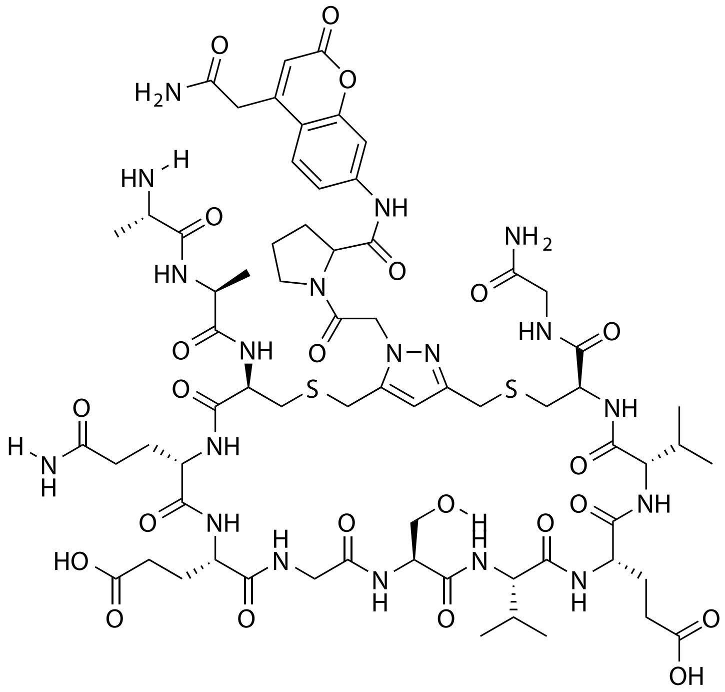


| Probe Name | Sequence | Expected Mass [M+1]^{+1} or [M+2]^{+2} | Found Mass |
|---|---|---|---|
| DPP4 36 | AACQEGSVEVCG | 1598.6323 | 1598.9 |

Total Ion Chromatogram

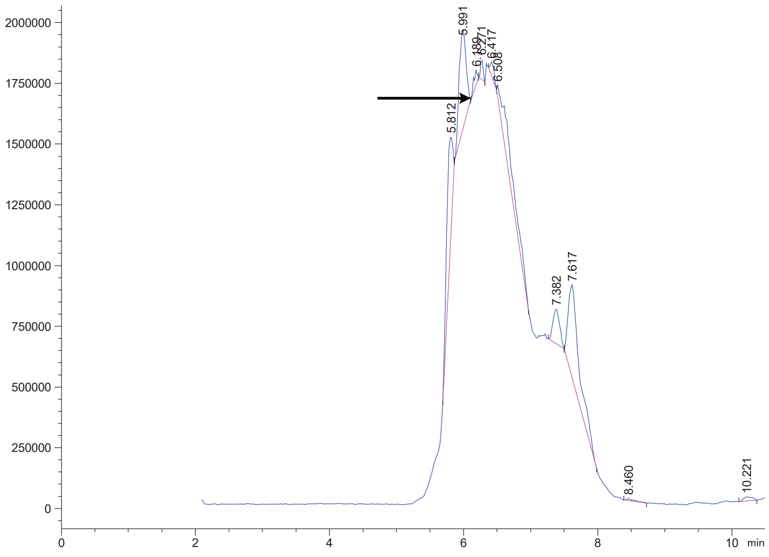


Variable Wavelength Detector: 220 nm

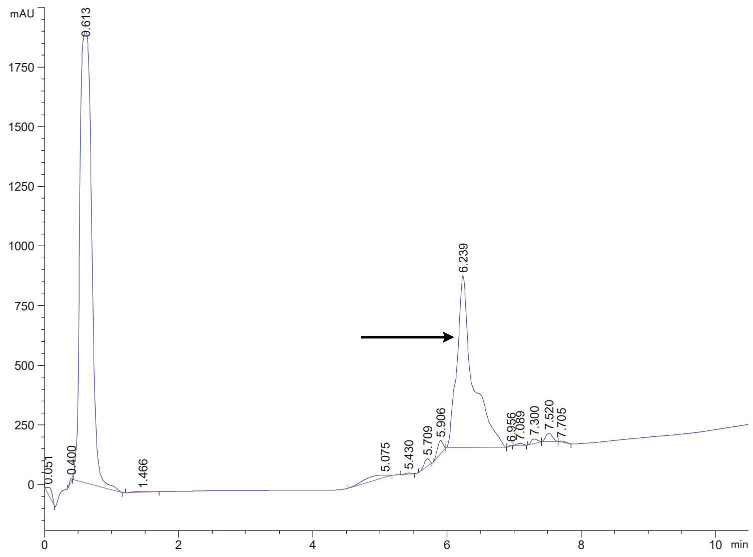

DPP4 37 Crude

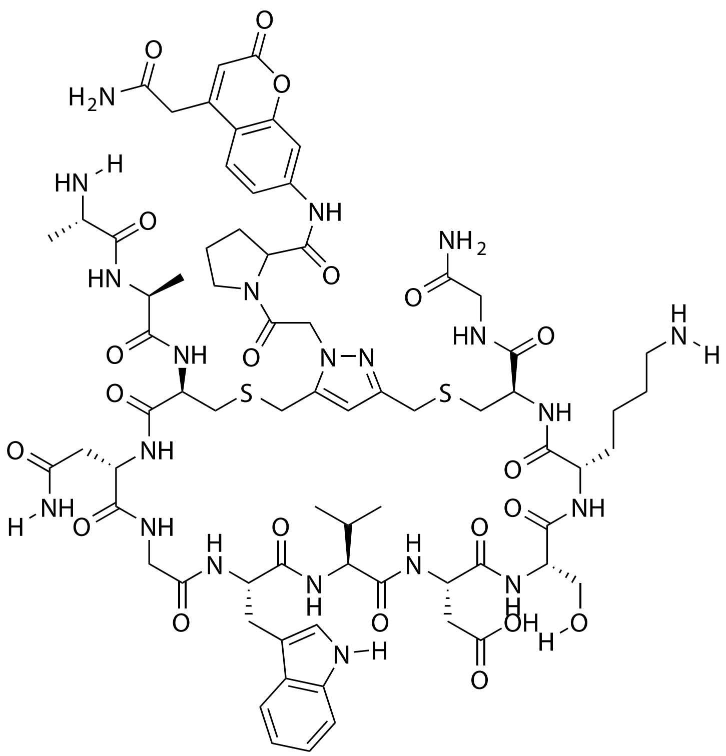

| Probe Name | Sequence | Expected Mass $[M+1]^{+1}$ or $[M+2]^{+2}$ | Found Mass |
|---|---|---|---|
| DPP4 37 | AACNGWVDSKCG | 1656.6643 | 1656.9 |

Total Ion Chromatogram

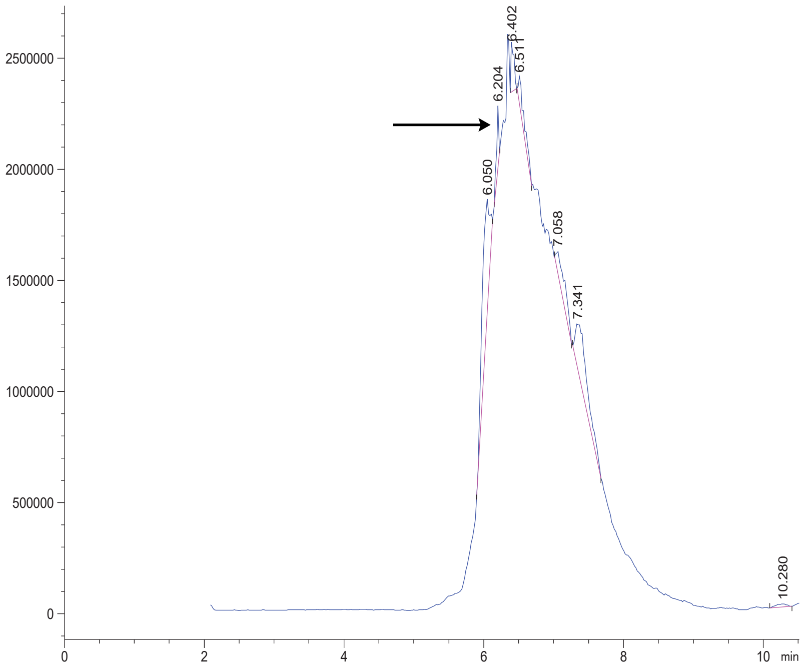

Variable Wavelength Detector: 220 nm

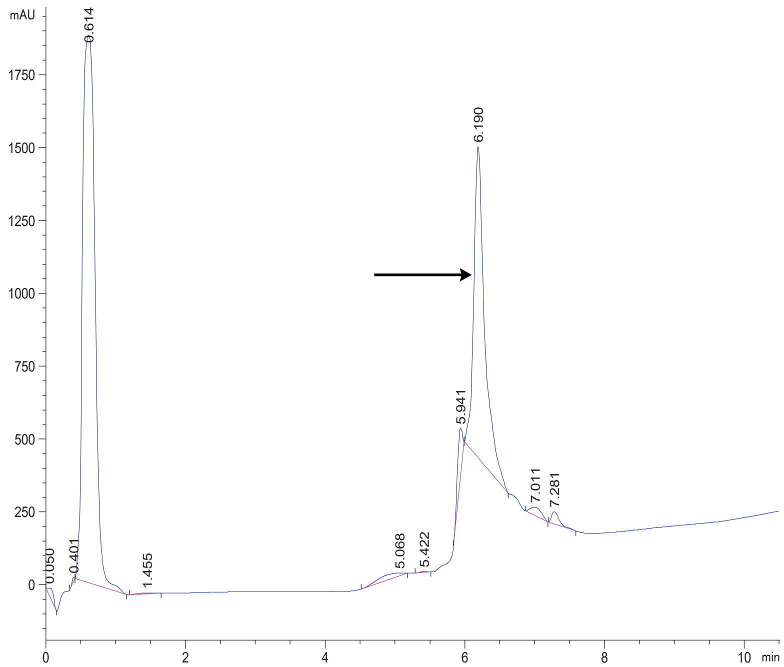

DPP4 38 Crude

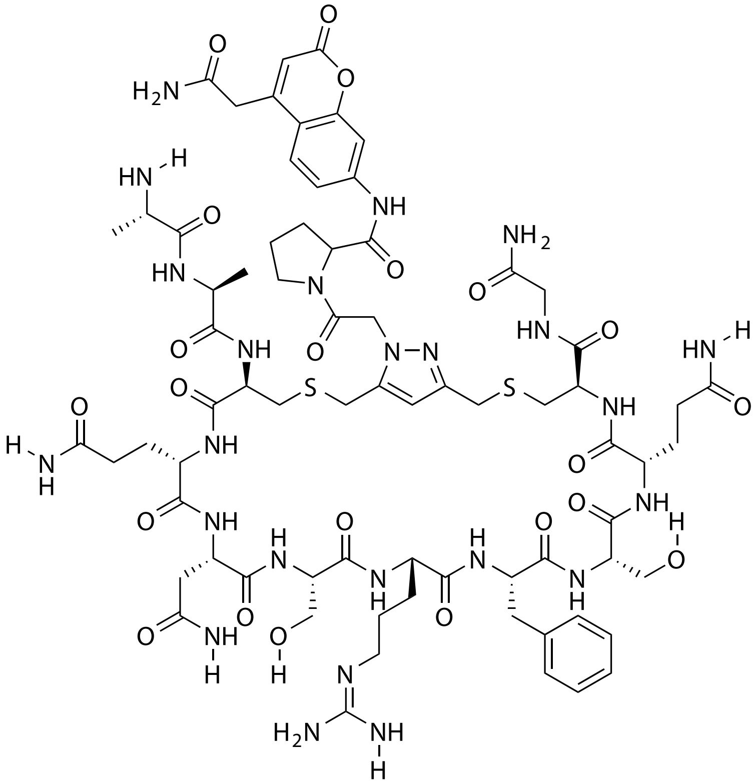

| Probe Name | Sequence | Expected Mass $[M+1]^{+1}$ or $[M+2]^{+2}$ | Found Mass |
|---|---|---|---|
| DPP4 38 | AACQNSRFSQCG | 1717.6919 | 1717.9 |

Total Ion Chromatogram

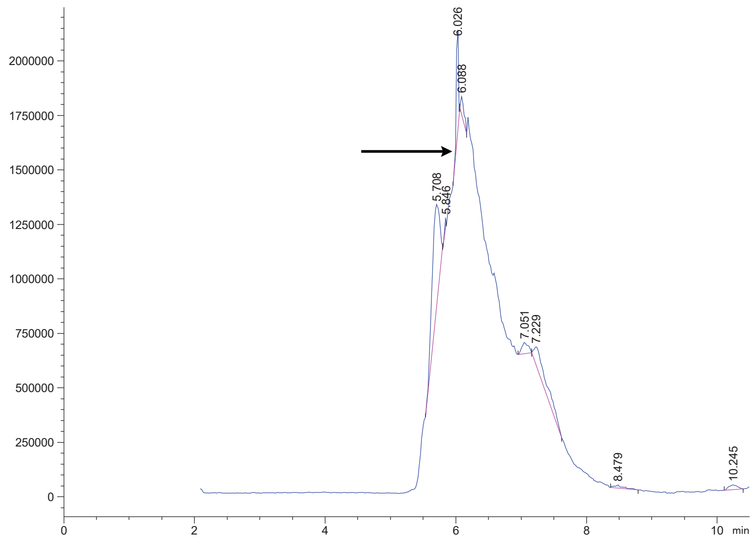

Variable Wavelength Detector: 220 nm

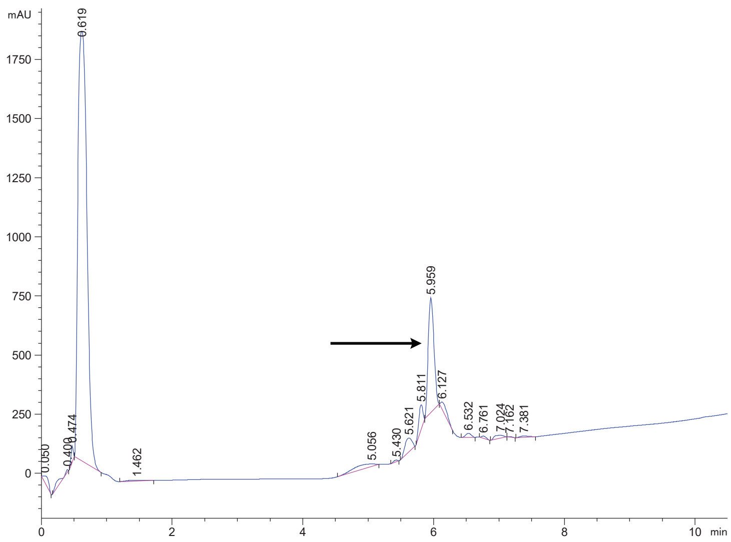

### DPP4 39 Crude

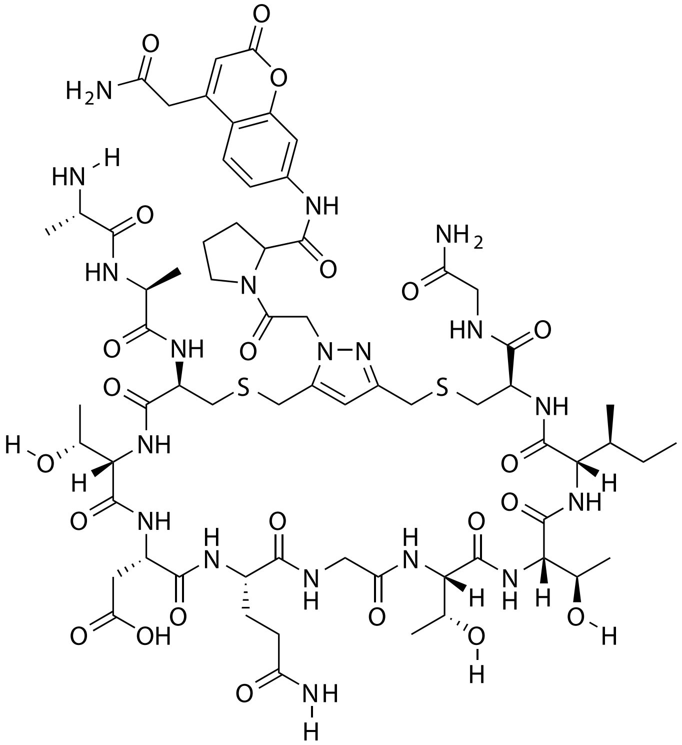


| Probe Name | Sequence | Expected Mass $[M+1]^{+1}$ or $[M+2]^{+2}$ | Found Mass |
|---|---|---|---|
| DPP4 39 | AACTDQGTTICG | 1586.6323 | 1586.9 |

Total Ion Chromatogram

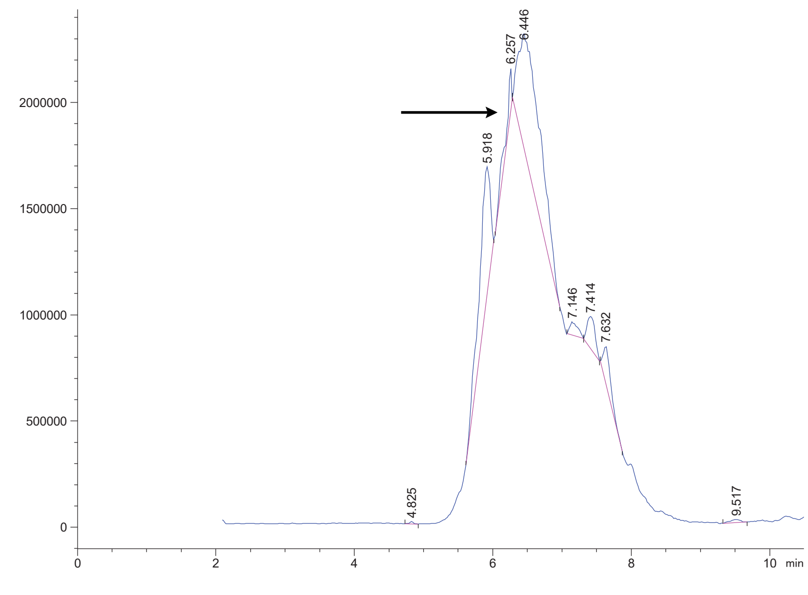


Variable Wavelength Detector: 220 nm

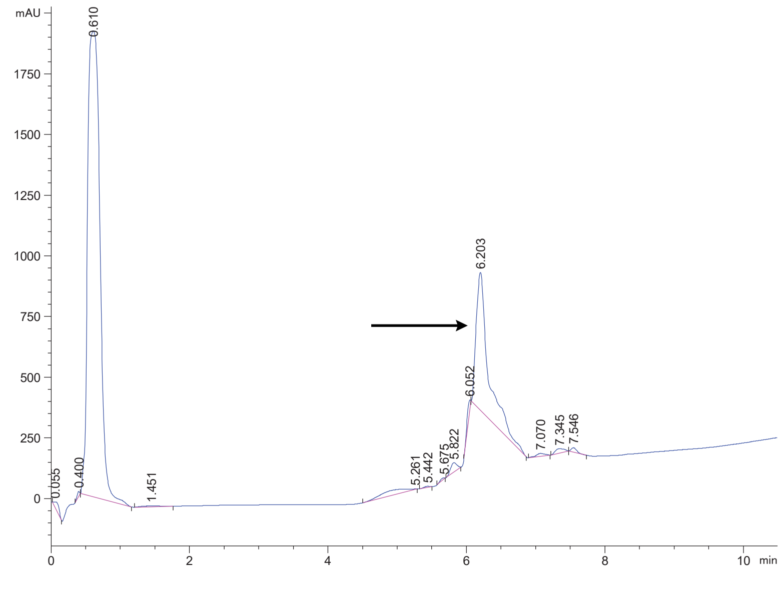

FAP 1 Crude

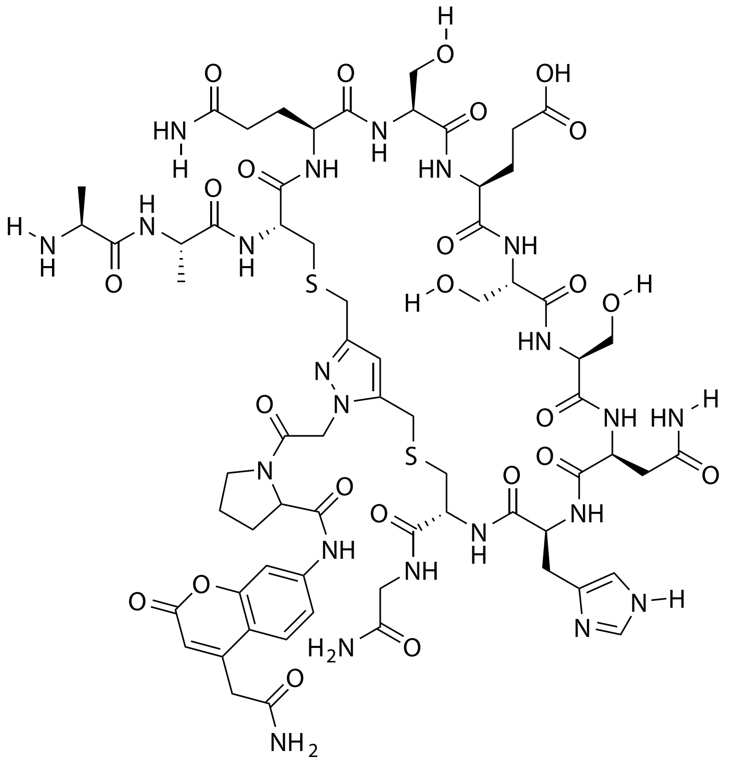

| Probe Name | Sequence | Expected Mass $[M+1]^{+1}$ or $[M+2]^{+2}$ | Found Mass |
|---|---|---|---|
| FAP 1 | AACQSESSNHCG | 820.3026 $[M+2]^{+2}$ | 820.2 $[M+2]^{+2}$ |

Total Ion Chromatogram

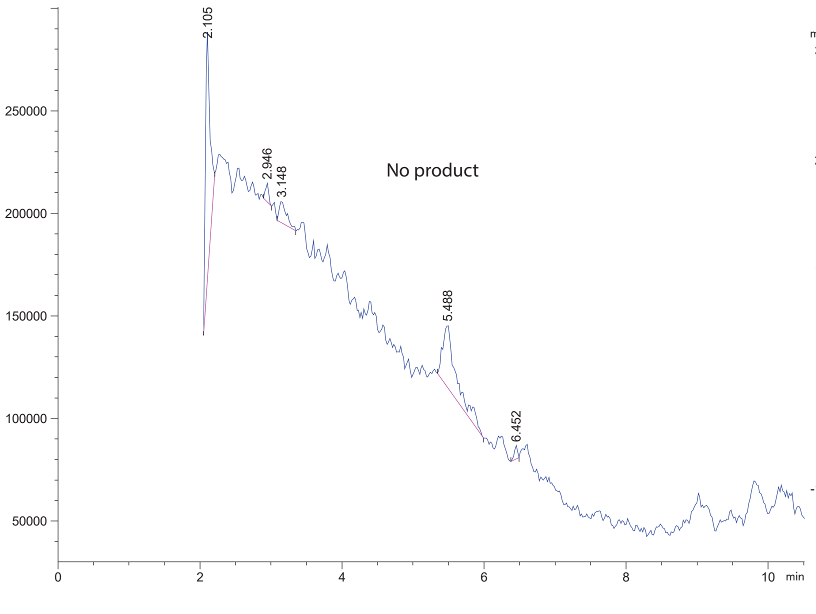


Variable Wavelength Detector: 220 nm

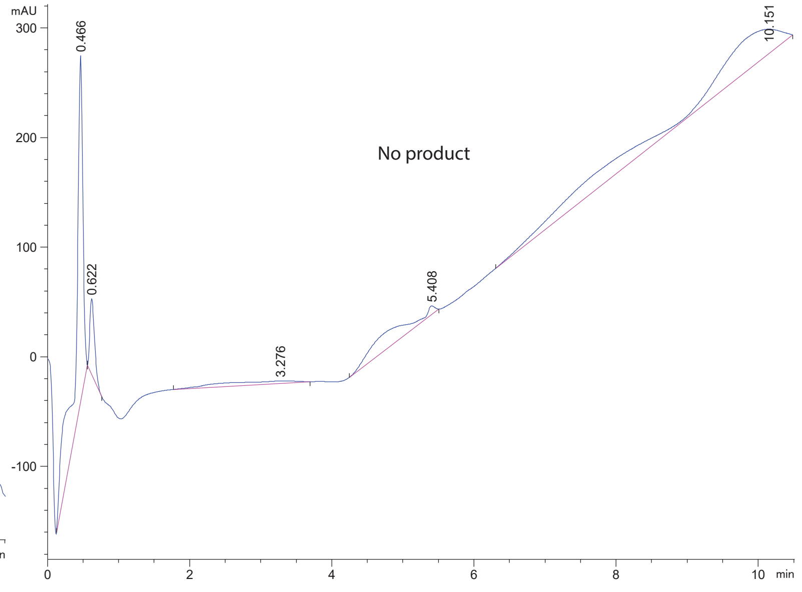

FAP 2 Crude

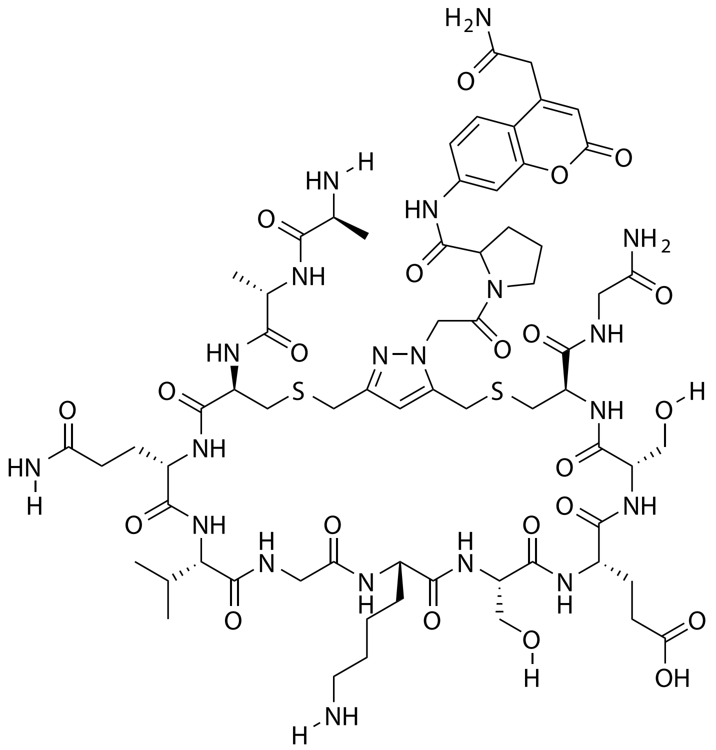

| Probe Name | Sequence | Expected Mass [M+1]$^{+1}$ or [M+2]$^{+2}$ | Found Mass |
|---|---|---|---|
| FAP 2 | AACQVGKSESCG | 1585.6483 | 1585.9 |

Total Ion Chromatogram

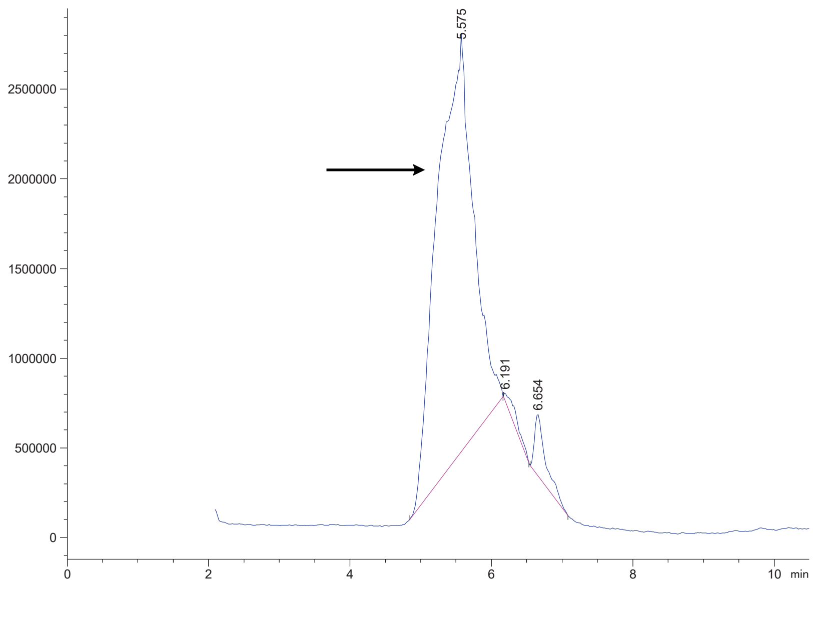

Variable Wavelength Detector: 220 nm

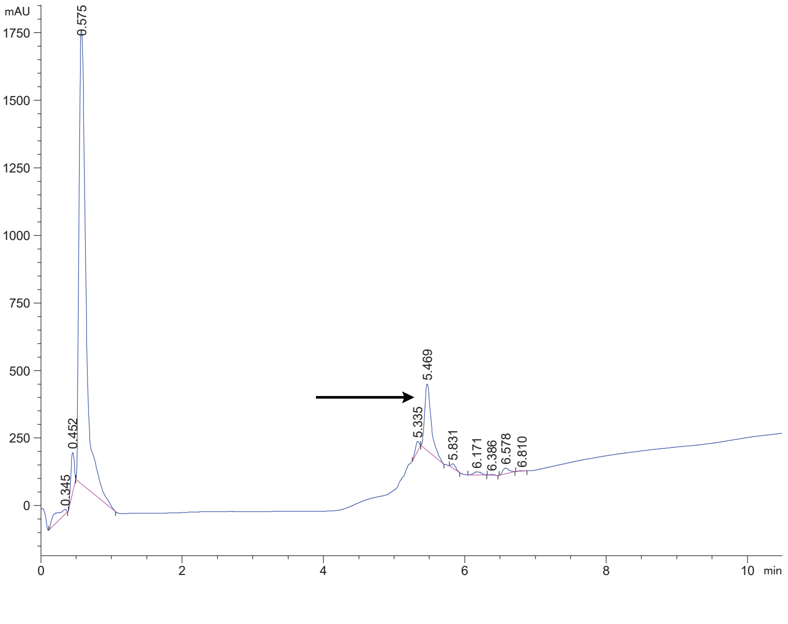

FAP 3 Crude

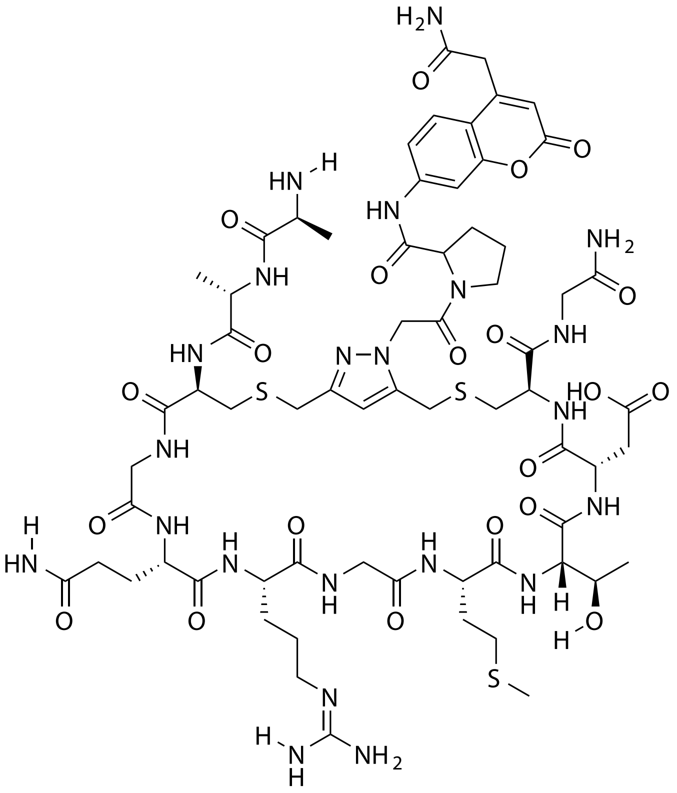

| Probe Name | Sequence | Expected Mass $[M+1]^{+1}$ or $[M+2]^{+2}$ | Found Mass |
|---|---|---|---|
| FAP 3 | AACGQRGMTDCG | 1615.6160 | 1615.8 |

Total Ion Chromatogram

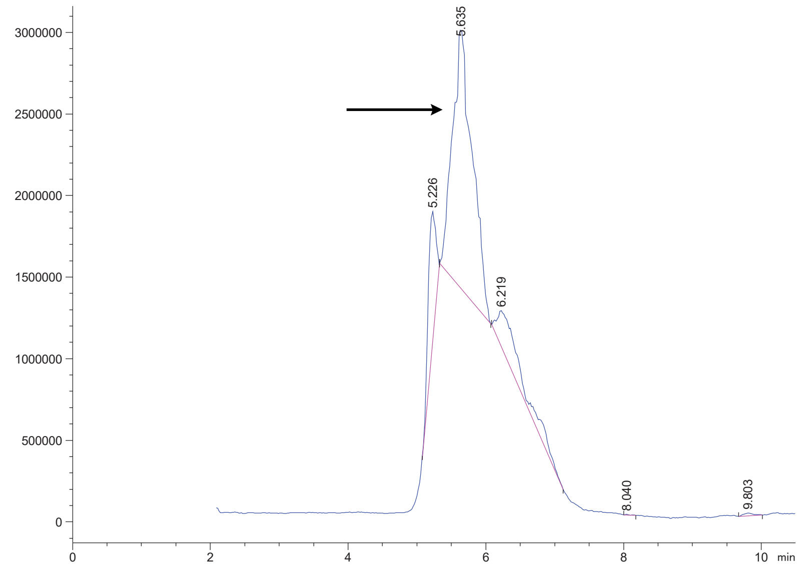

Variable Wavelength Detector: 220 nm

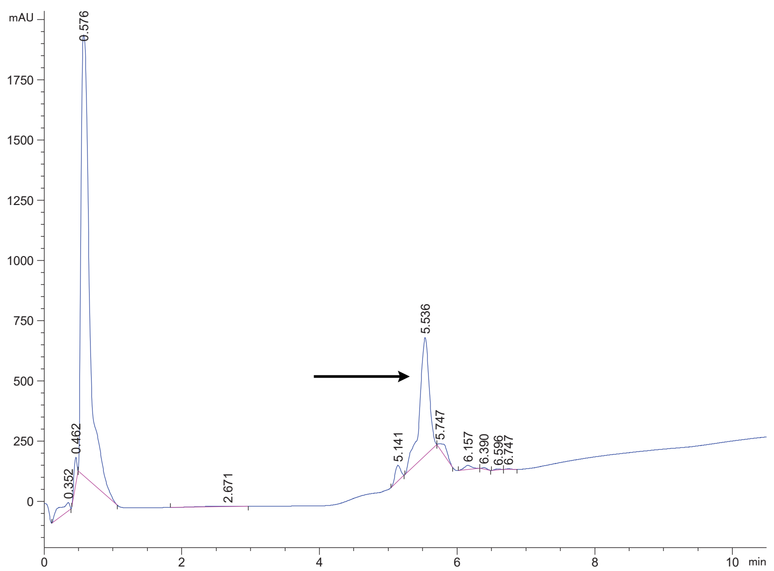

FAP 4 Crude

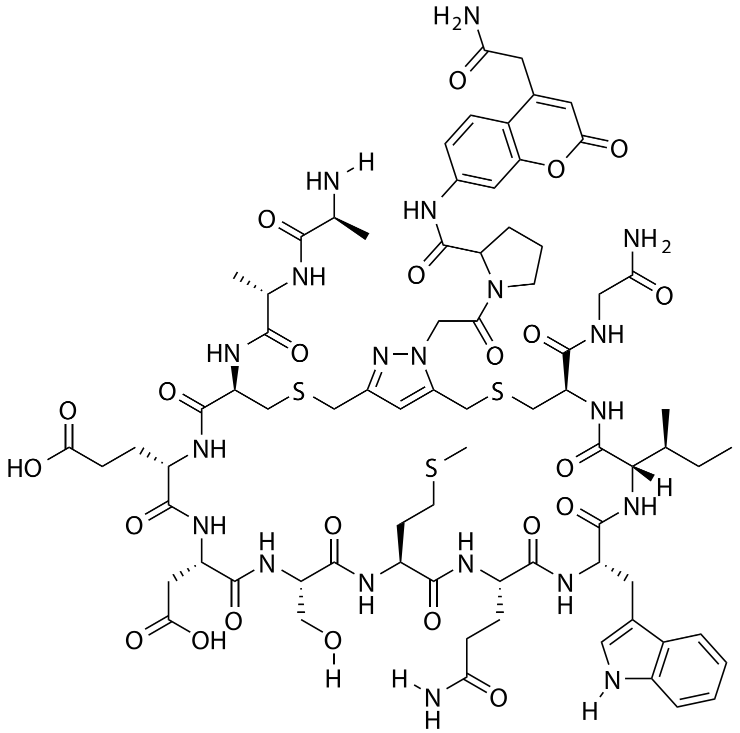

| Probe Name | Sequence | Expected Mass $[M+1]^{+1}$ or $[M+2]^{+2}$ | Found Mass |
|---|---|---|---|
| FAP 4 | AACEDSMQWICG | 880.3350 $[M+2]^{+2}$ | 880.2 $[M+2]^{+2}$ |

Total Ion Chromatogram

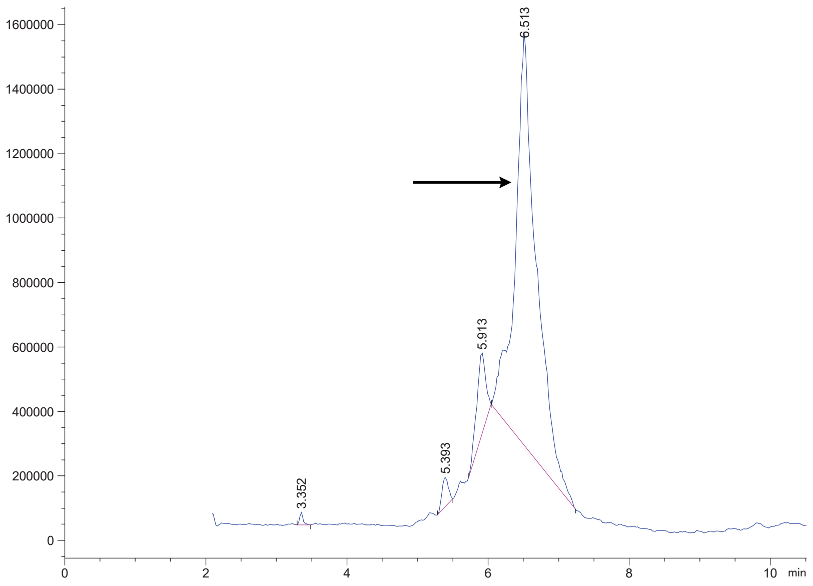


Variable Wavelength Detector: 220 nm

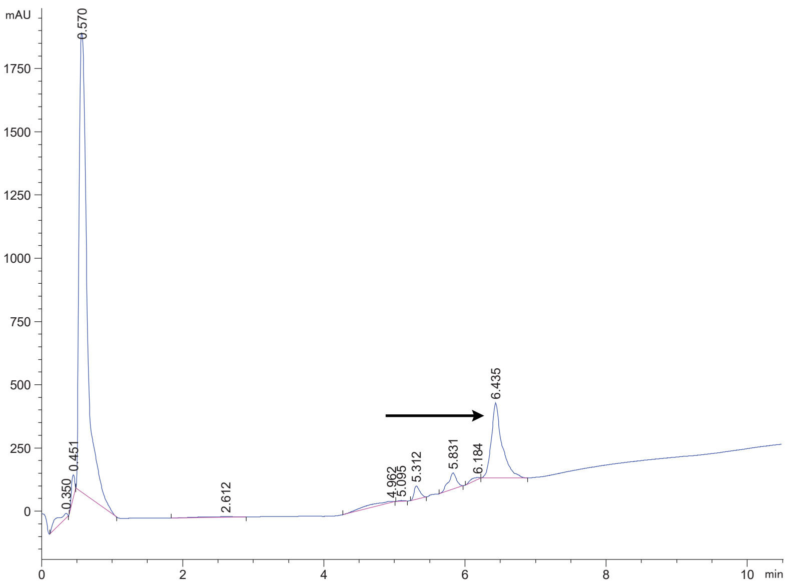

FAP 6 crude

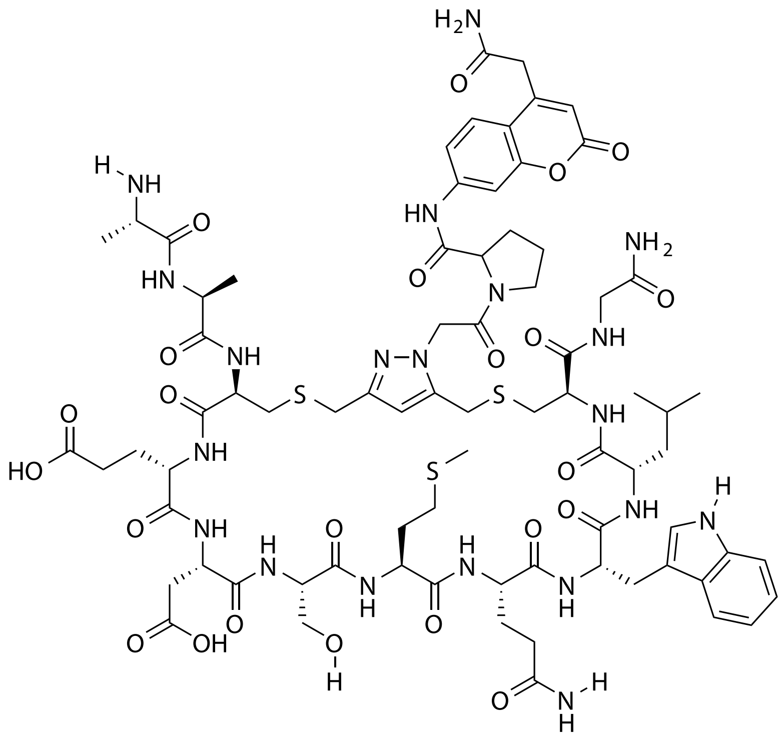


| Probe Name | Sequence | Expected Mass $[M+1]^{+1}$ or $[M+2]^{+2}$ | Found Mass |
|---|---|---|---|
| FAP 6 | AACEDSMQWLCG | 1759.6622 | 1758.8 |

Total Ion Chromatogram

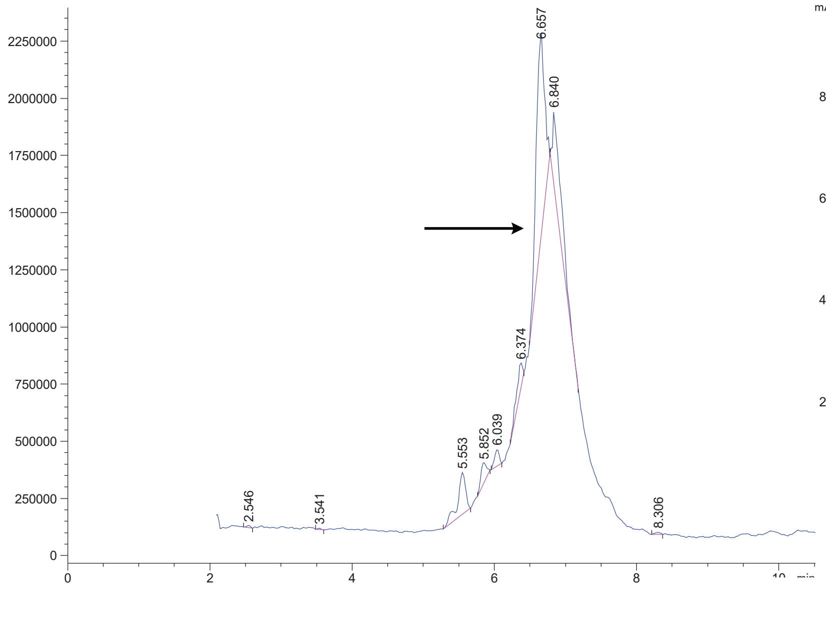


Variable Wavelength Detector: 220 nm

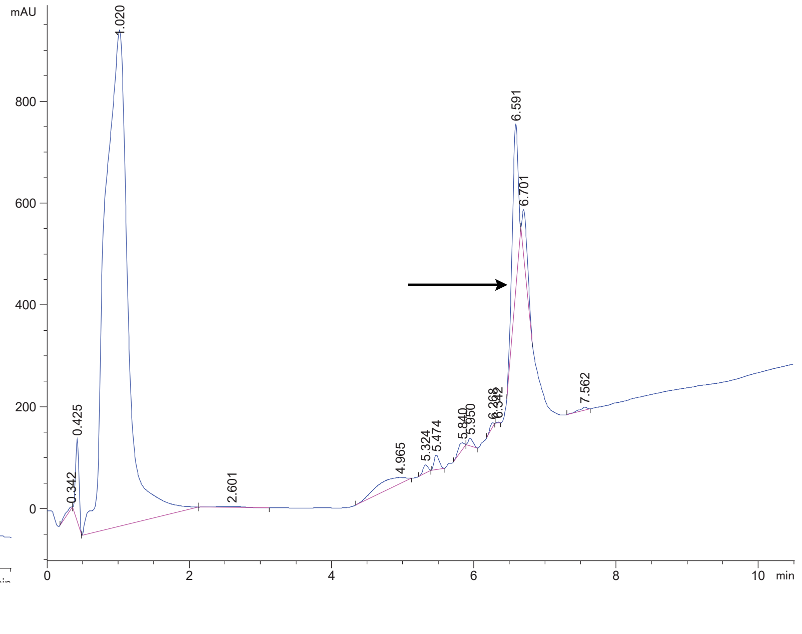

FAP 5 Crude

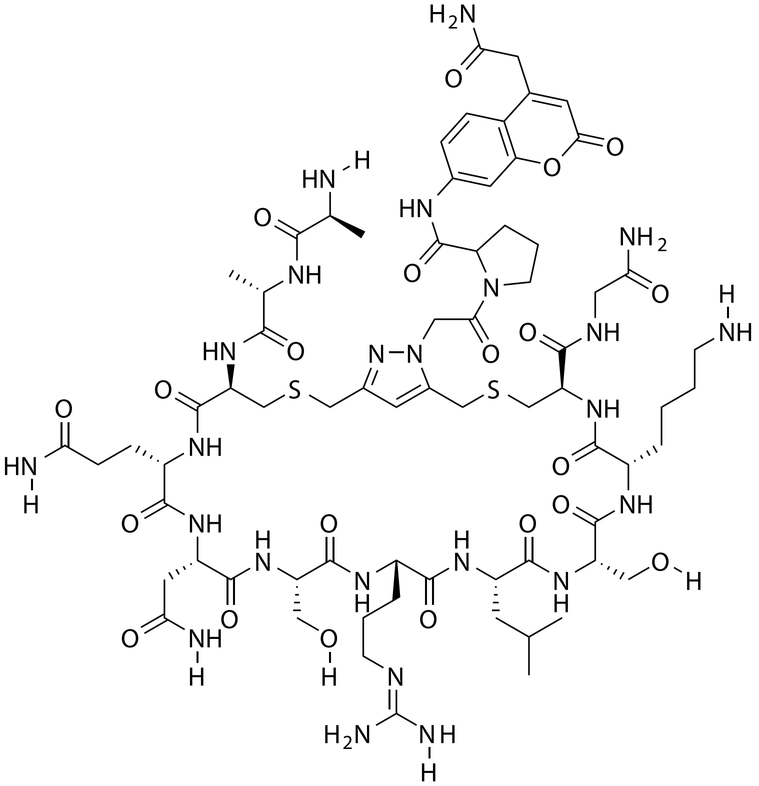

| Probe Name | Sequence | Expected Mass $[M+1]^{+1}$ or $[M+2]^{+2}$ | Found Mass |
|---|---|---|---|
| FAP 5 | AACQNSRLSKCG | 1683.7439 | 1683.0 |

Total Ion Chromatogram

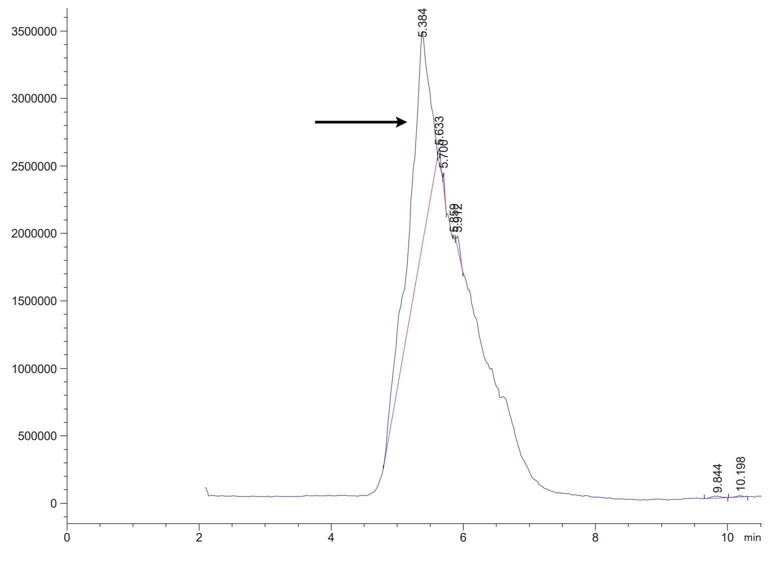

Variable Wavelength Detector: 220 nm

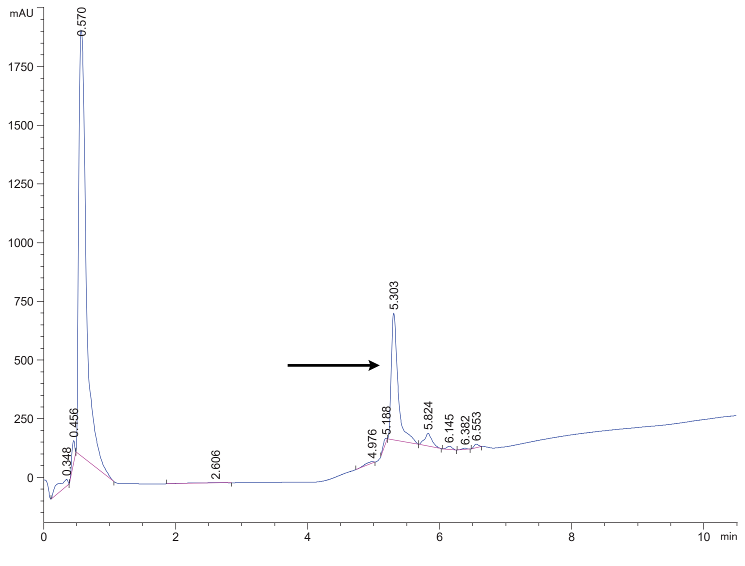

FAP 7 crude

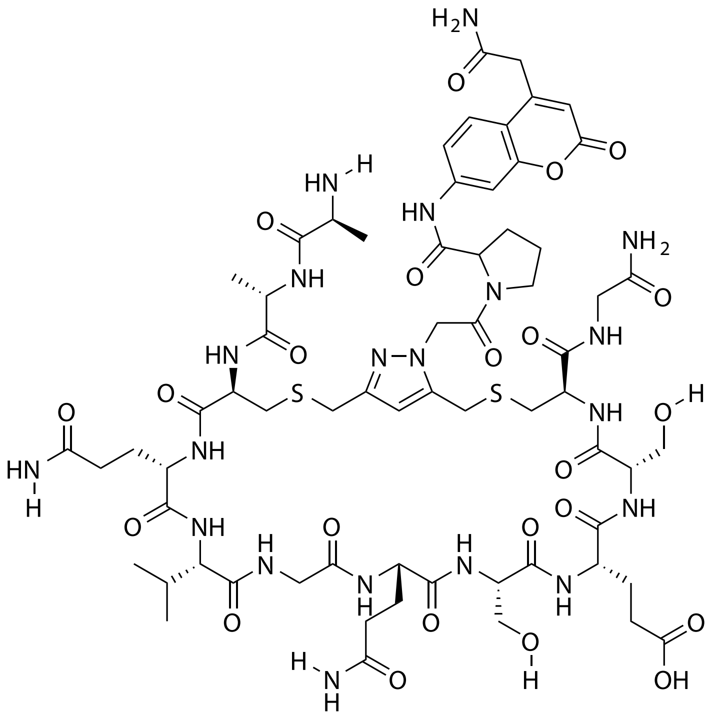


| Probe Name | Sequence | Expected Mass $[M+1]^{+1}$ or $[M+2]^{+2}$ | Found Mass |
|---|---|---|---|
| FAP 7 | AACQVGQSESCG | 1585.6119 | 1585.9 |

Total Ion Chromatogram

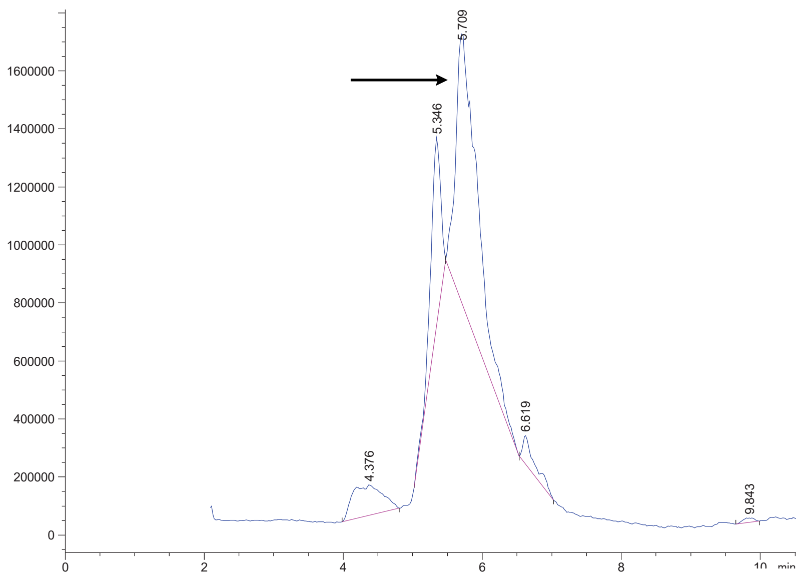


Variable Wavelength Detector: 220 nm

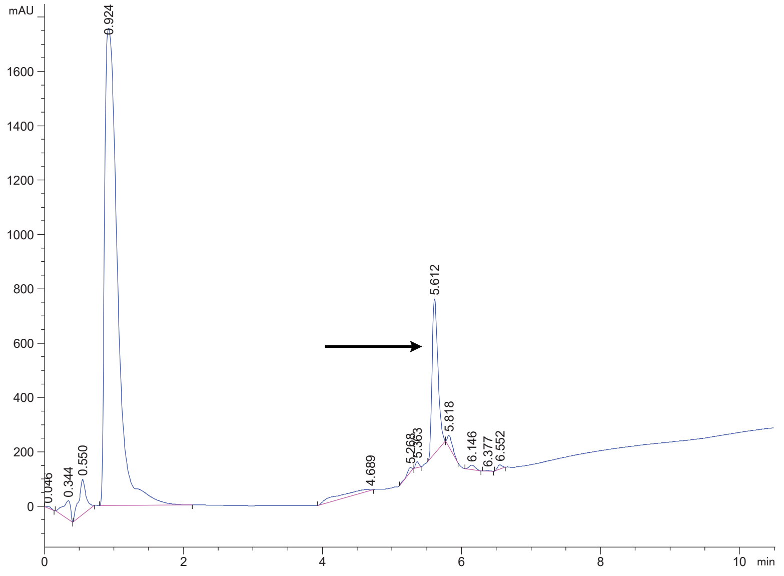

FAP 8 crude

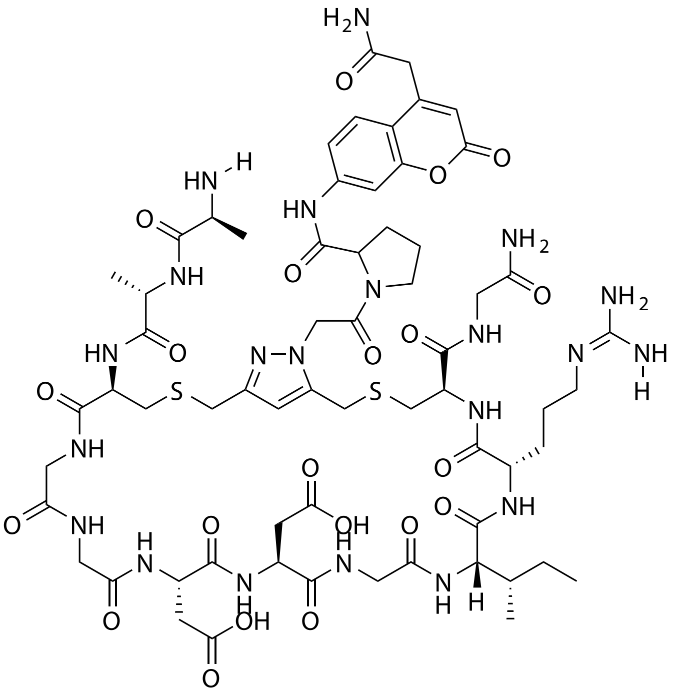

| Probe Name | Sequence | Expected Mass $[M+1]^{+1}$ or $[M+2]^{+2}$ | Found Mass |
|---|---|---|---|
| FAP 8 | AACGGDDGIRCG | 1540.6017 | 1540.8 |

Total Ion Chromatogram

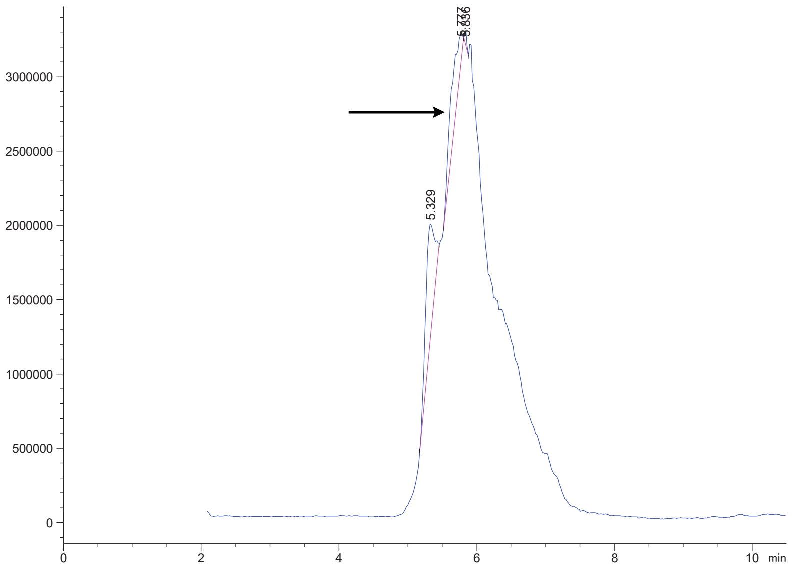


Variable Wavelength Detector: 220 nm

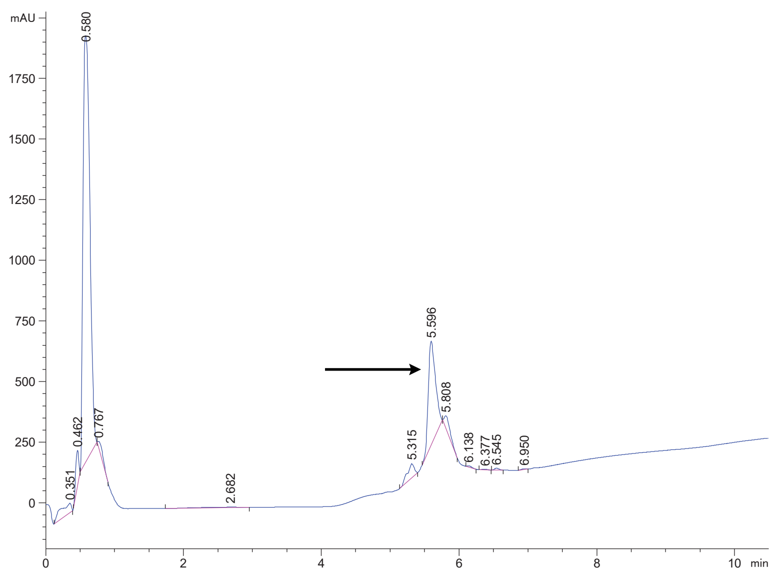

### FAP 9 crude

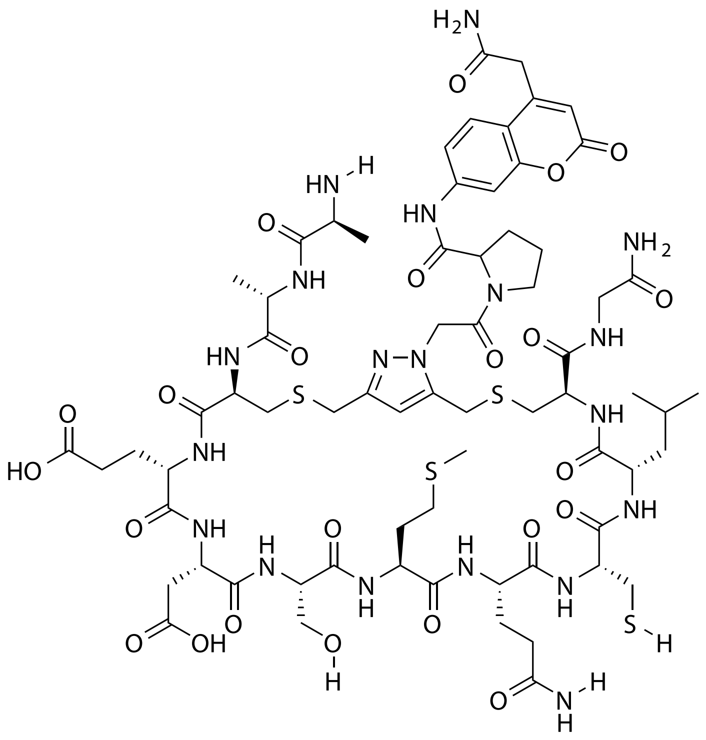

| Probe Name | Sequence | Expected Mass [M+1]$^{+1}$ or [M+2]$^{+2}$ | Found Mass |
|---|---|---|---|
| FAP 9 | AACEDSMQCLCG | 1676.5921 | 1676.8 |

Total Ion Chromatogram

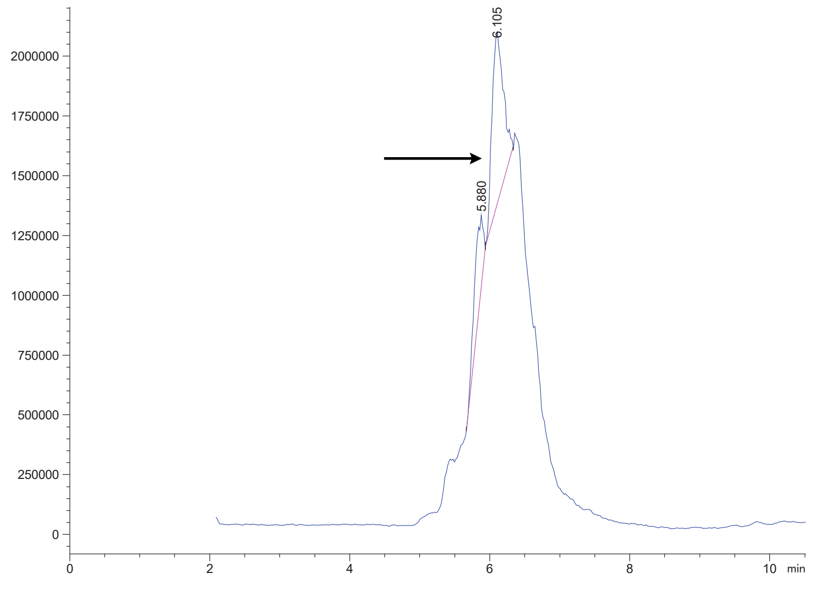


Variable Wavelength Detector: 220 nm

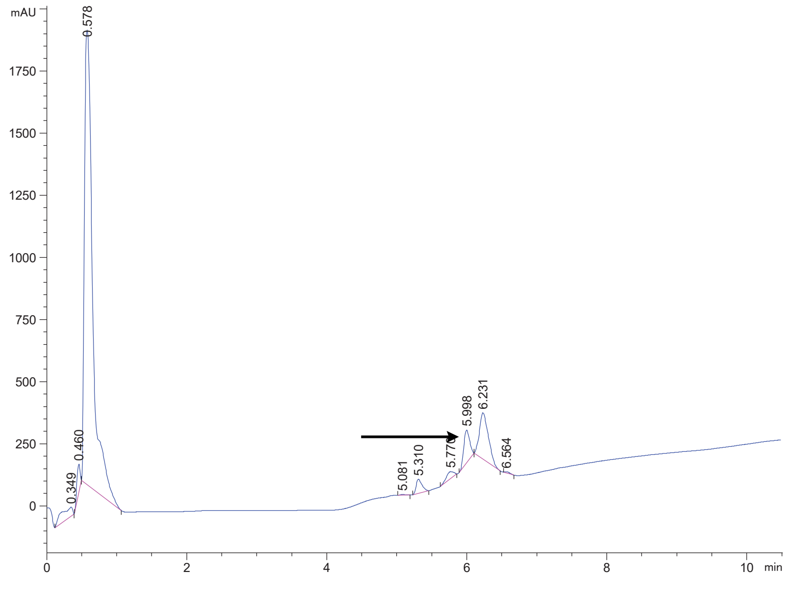

FAP 10 crude

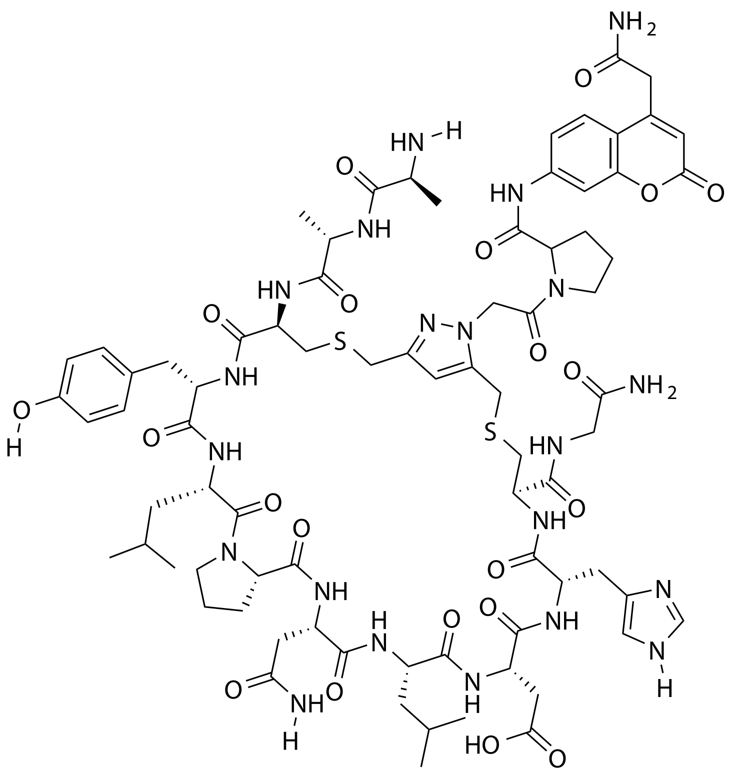

| Probe Name | Sequence | Expected Mass $[M+1]^{+1}$ or $[M+2]^{+2}$ | Found Mass |
|---|---|---|---|
| FAP 10 | AACYLPNLDHCG | 1722.7112 | 1722.9 |

Total Ion Chromatogram

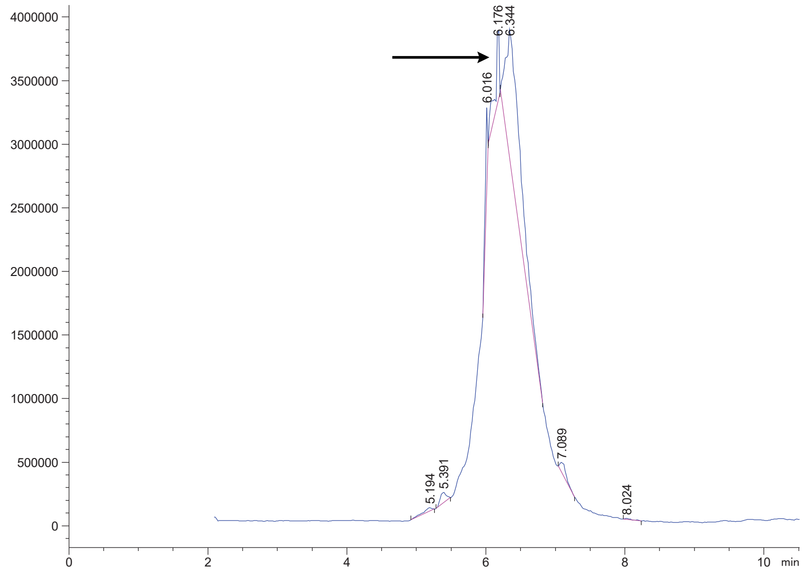


Variable Wavelength Detector: 220 nm

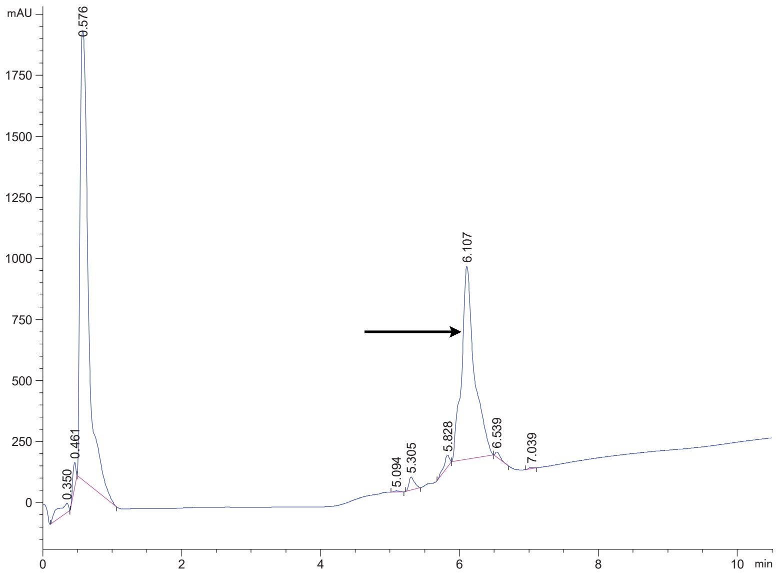

### FAP 11 crude

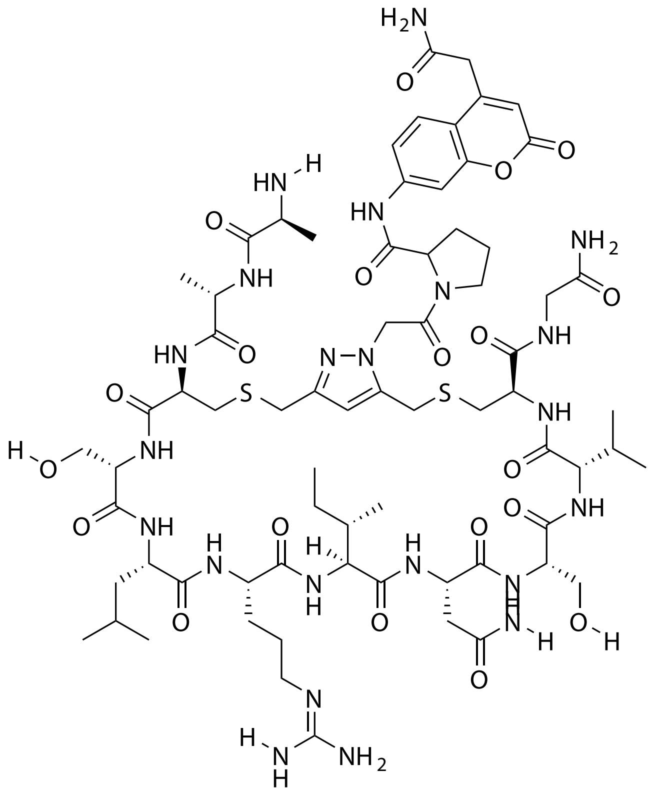

| **Probe Name** | **Sequence** | **Expected Mass $[M+1]^{+1}$ or $[M+2]^{+2}$** | **Found Mass** |
|---|---|---|---|
| FAP 11 | AACSLRINSVCG | 1639.7429 | 1639.0 |

Total Ion Chromatogram

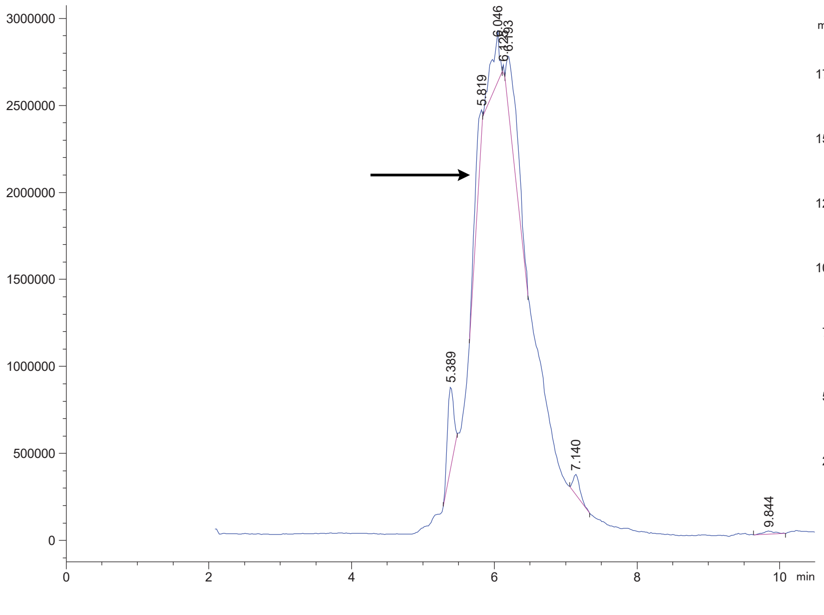


Variable Wavelength Detector: 220 nm

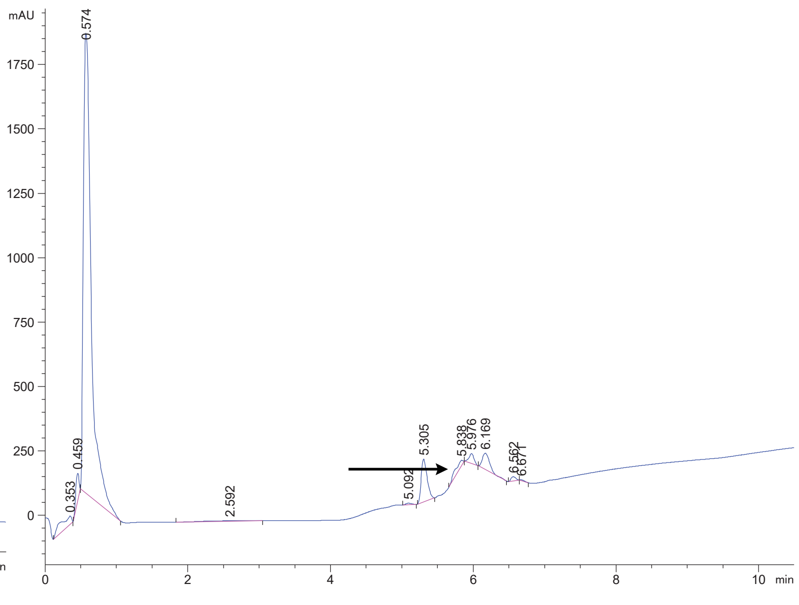

FAP 12 crude

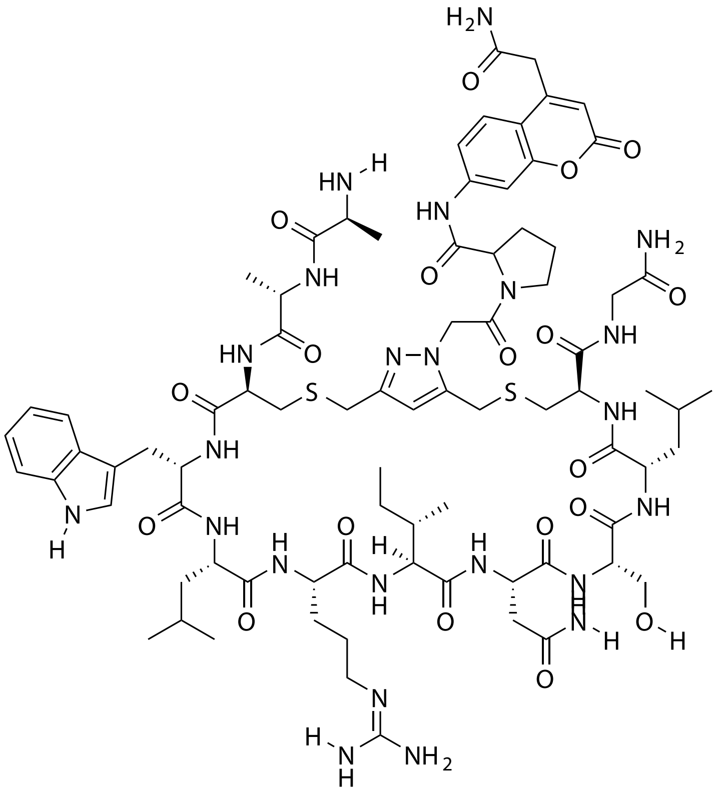

| Probe Name | Sequence | Expected Mass $[M+1]^{+1}$ or $[M+2]^{+2}$ | Found Mass |
|---|---|---|---|
| FAP 12 | AACWLRINSLCG | 876.9068 $[M+2]^{+2}$ | 876.4 $[M+2]^{+2}$ |

Total Ion Chromatogram

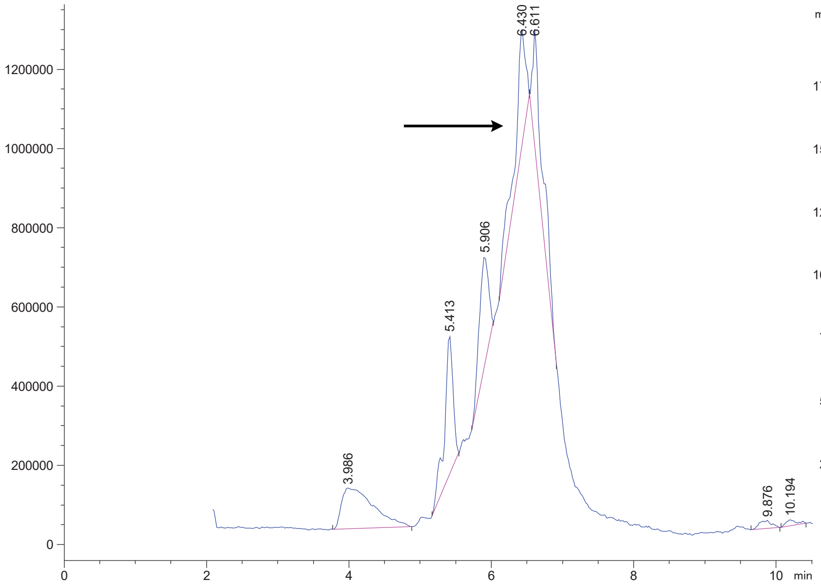


Variable Wavelength Detector: 220 nm

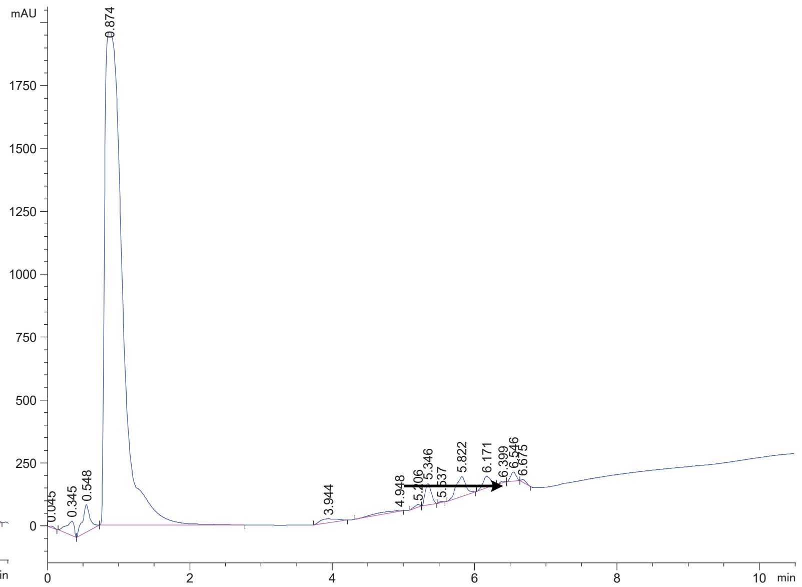

FAP 13 crude

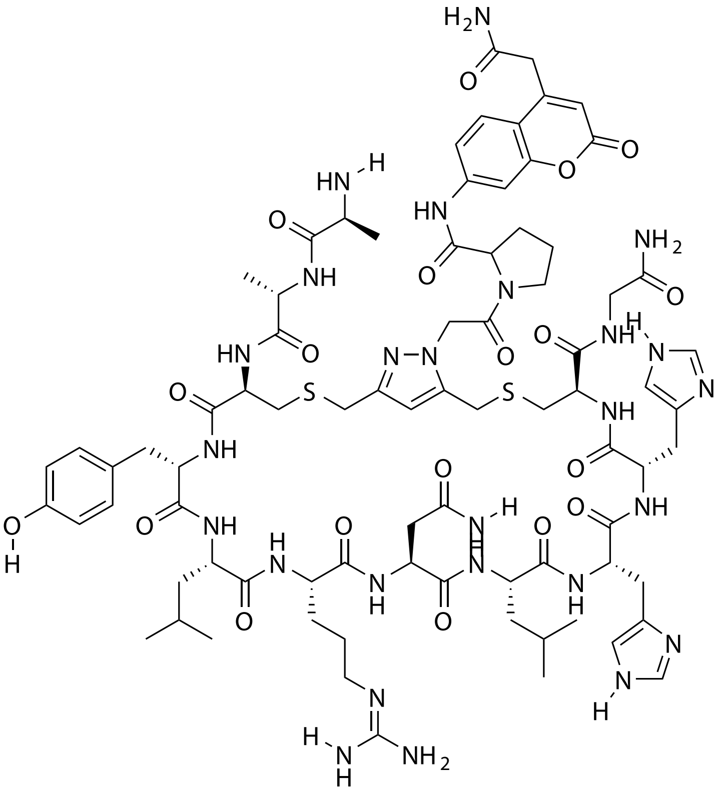

| Probe Name | Sequence | Expected Mass $[M+1]^{+1}$ or $[M+2]^{+2}$ | Found Mass |
|---|---|---|---|
| FAP 13 | AACYLRNLHHCG | 902.3997 $[M+2]^{+2}$ | 902.2 $[M+2]^{+2}$ |

Total Ion Chromatogram

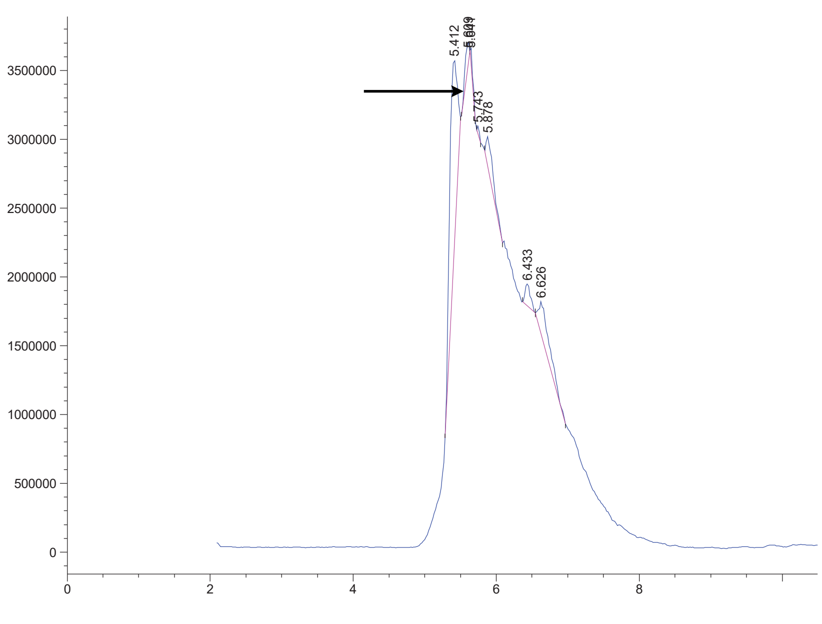


Variable Wavelength Detector: 220 nm

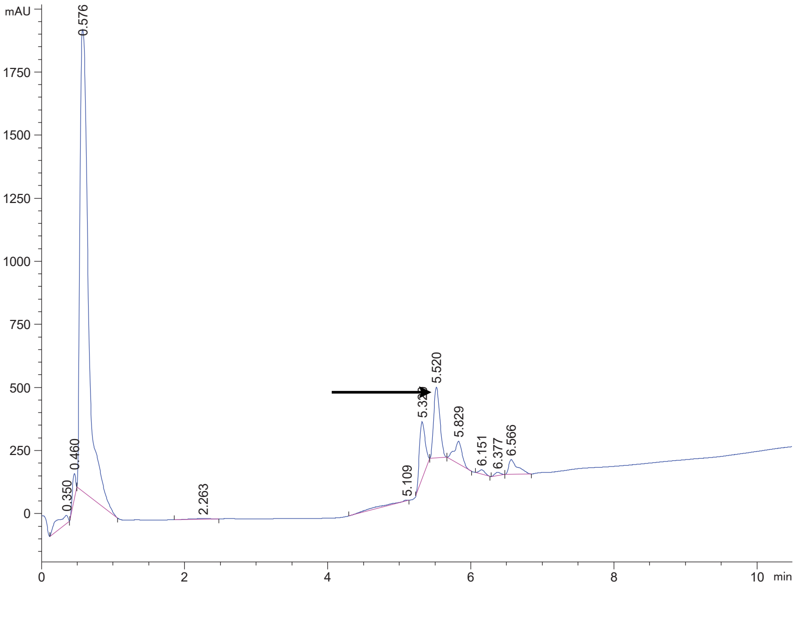

FAP 14 crude

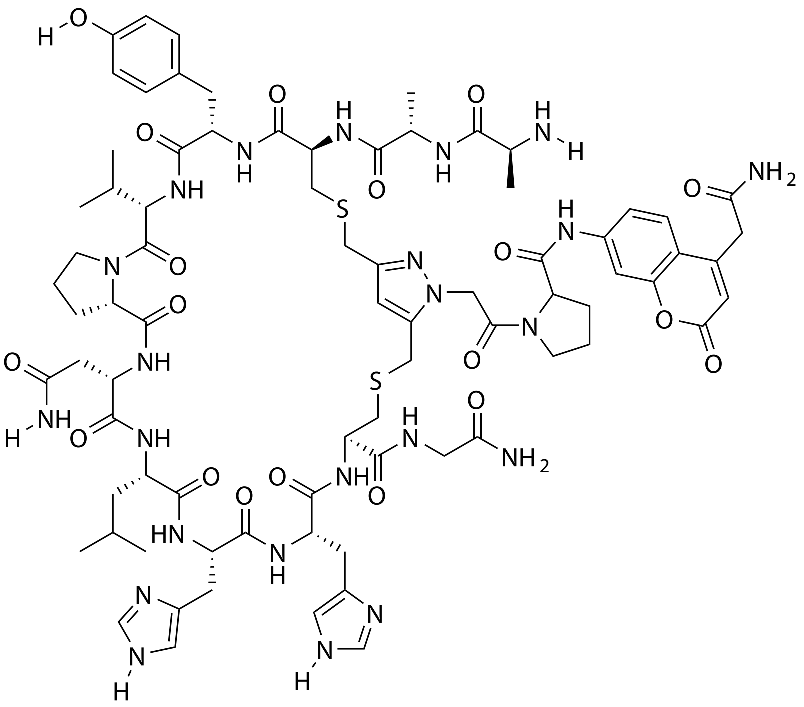

| Probe Name | Sequence | Expected Mass $[M+1]^{+1}$ or $[M+2]^{+2}$ | Found Mass |
|---|---|---|---|
| FAP 14 | AACYVPNLHHCG | 865.8677 $[M+2]^{+2}$ | 865.7 $[M+2]^{+2}$ |

Total Ion Chromatogram

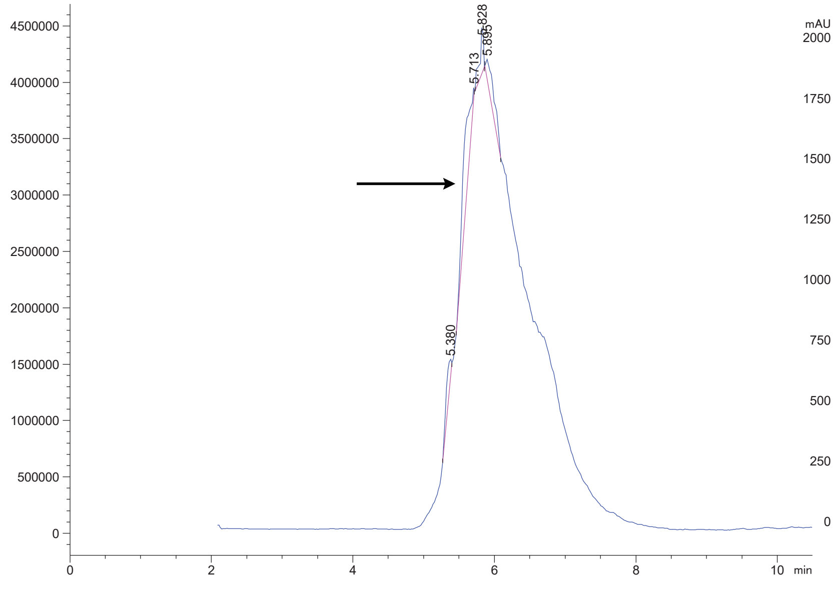


Variable Wavelength Detector: 220 nm

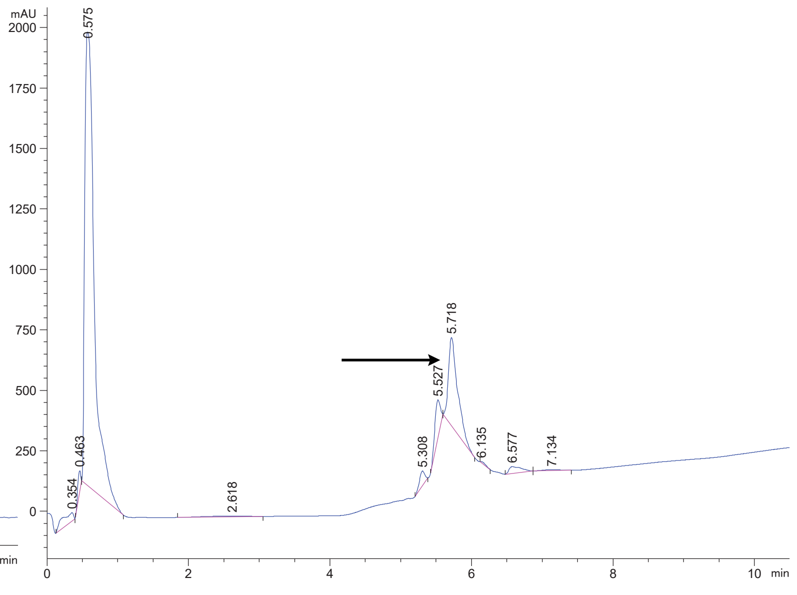

### FAP 15 crude

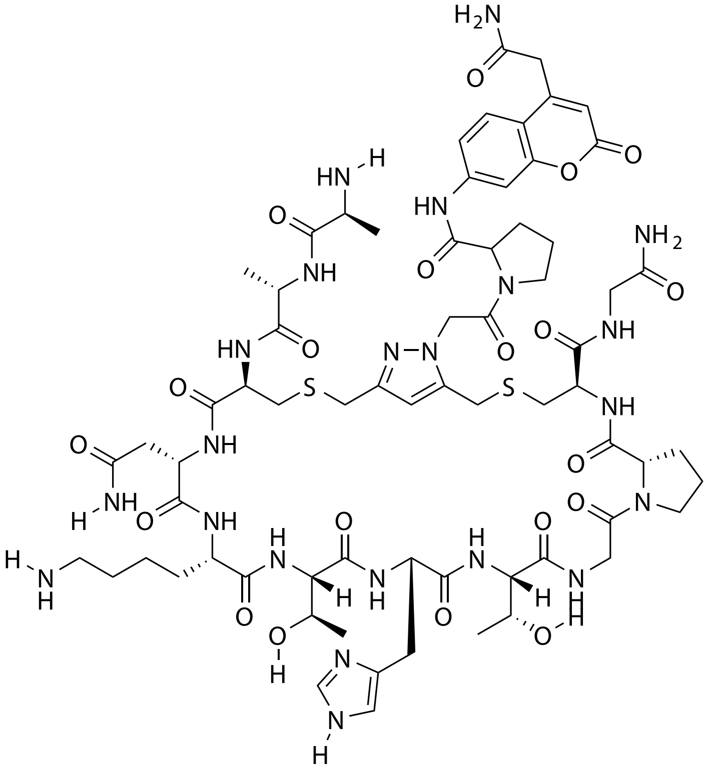

| Probe Name | Sequence | Expected Mass [M+1]$^{+1}$ or [M+2]$^{+2}$ | Found Mass |
|---|---|---|---|
| FAP 15 | AACNKTHTGPCG | 1605.6646 | 1604.9 |

Total Ion Chromatogram

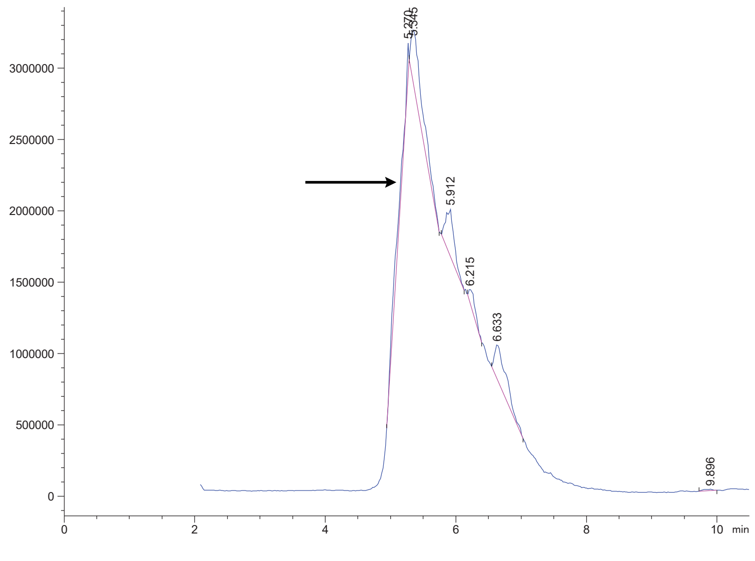


Variable Wavelength Detector: 220 nm

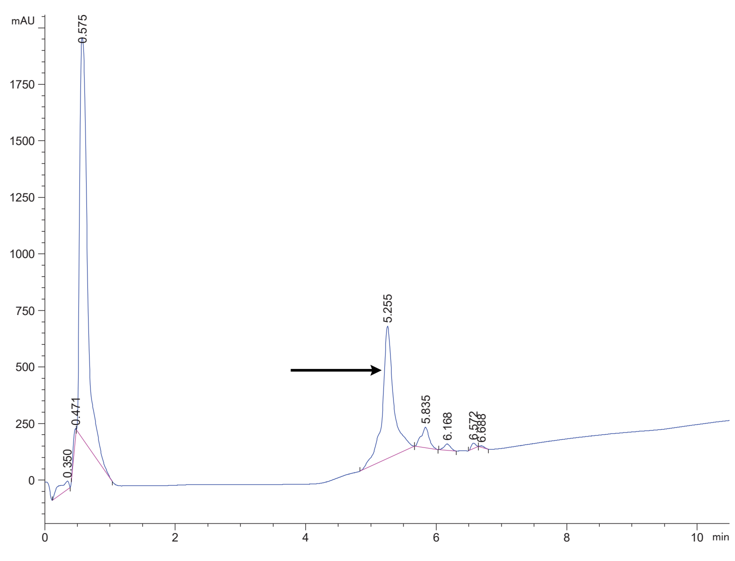

### FAP 16 crude

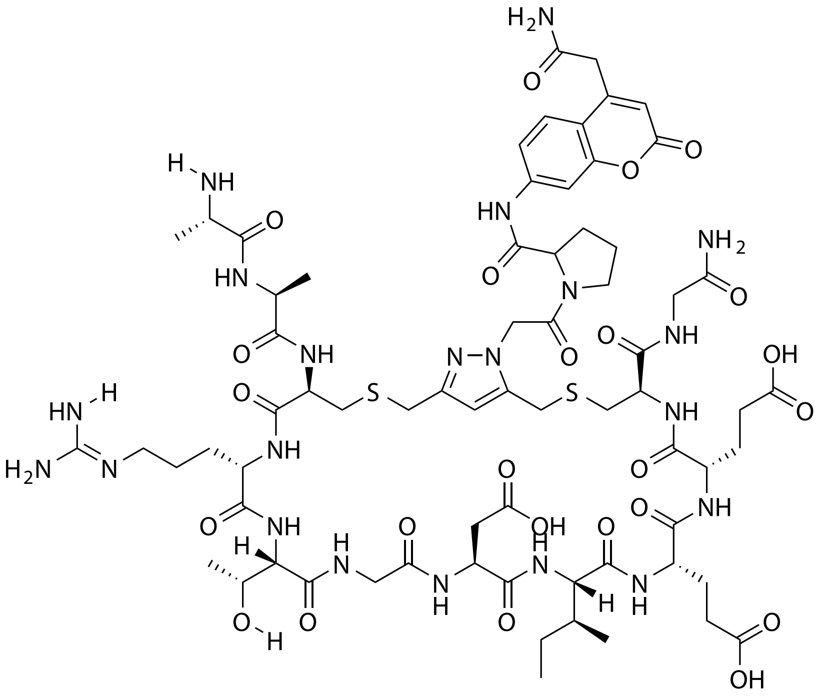

| Probe Name | Sequence | Expected Mass $[M+1]^{+1}$ or $[M+2]^{+2}$ | Found Mass |
|---|---|---|---|
| FAP 16 | AACRTGDIEECG | 1670.6647 | 1670.9 |

Total Ion Chromatogram

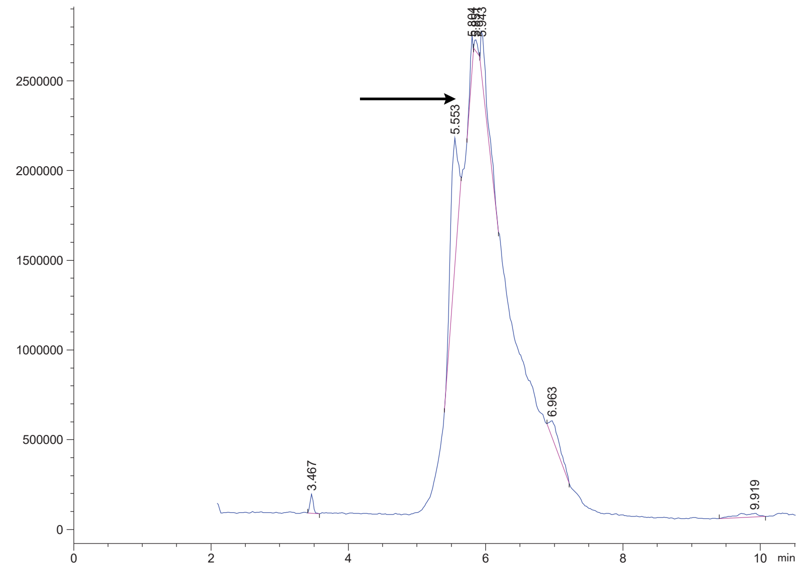


Variable Wavelength Detector: 220 nm

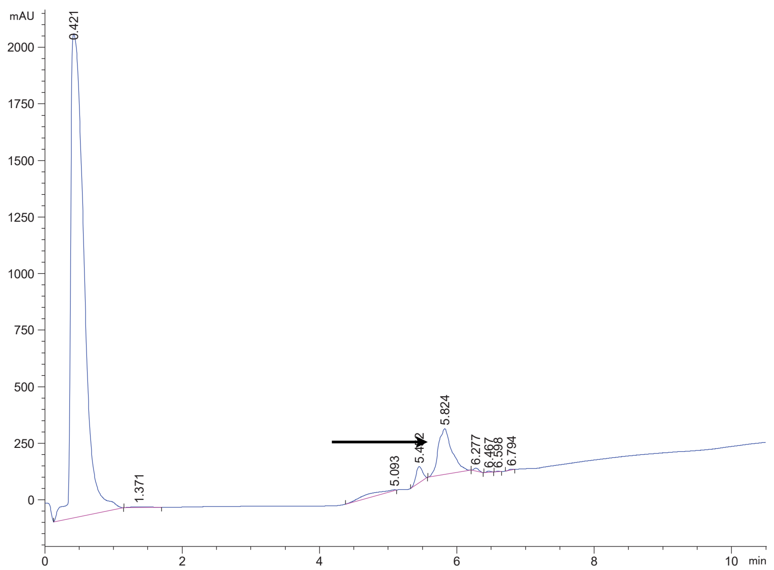

FAP 17 crude

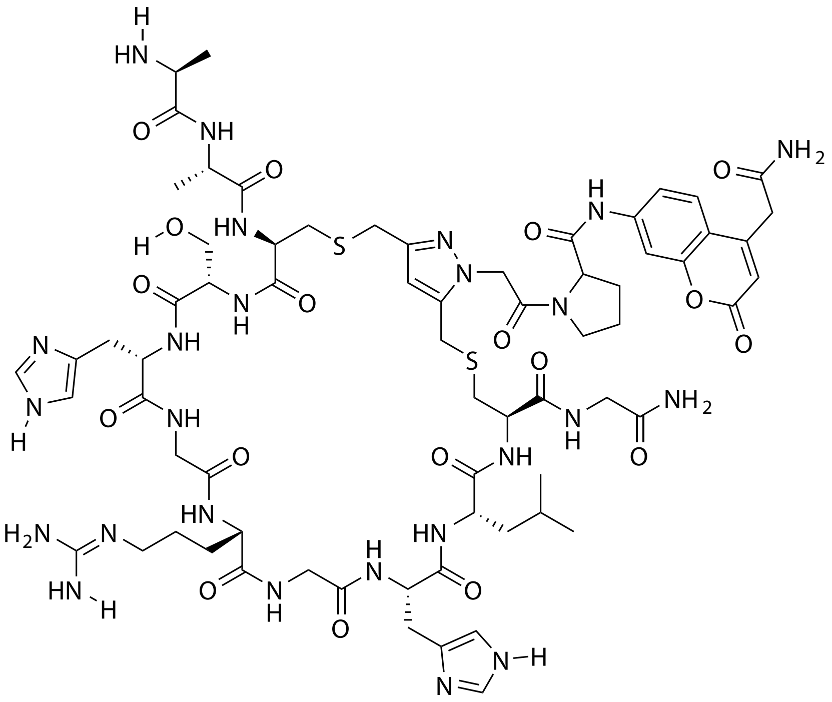

| Probe Name | Sequence | Expected Mass $[M+1]^{+1}$ or $[M+2]^{+2}$ | Found Mass |
|---|---|---|---|
| FAP 17 | AACSHGRGHLCG | 1614.6762 | 1614.9 |

Total Ion Chromatogram

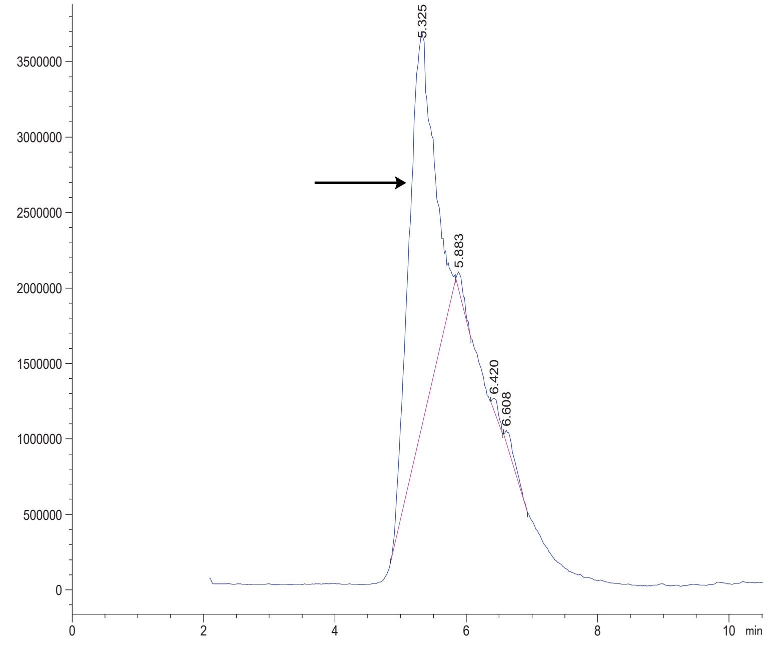


Variable Wavelength Detector: 220 nm

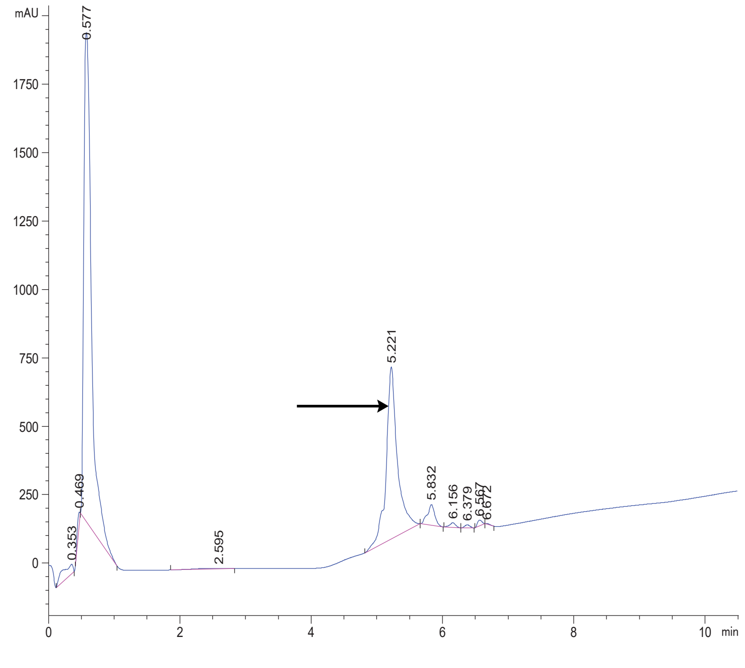

FAP 18 crude

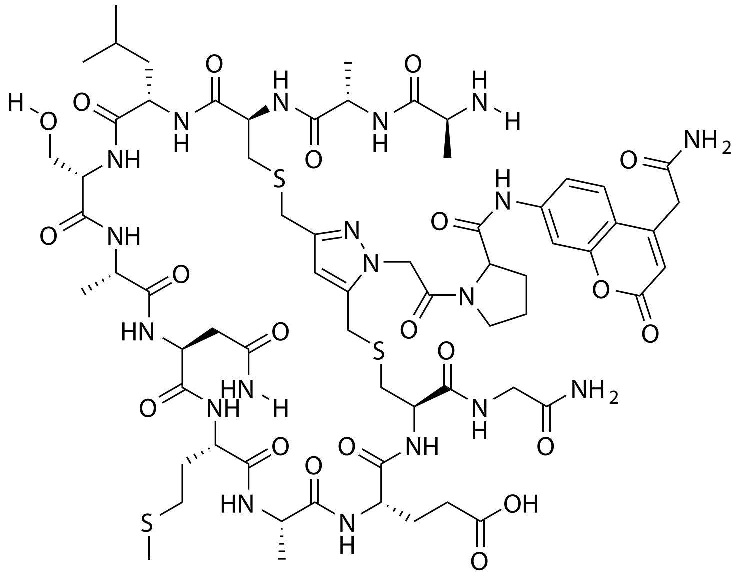

| Probe Name | Sequence | Expected Mass $[M+1]^{+1}$ or $[M+2]^{+2}$ | Found Mass |
|---|---|---|---|
| FAP 18 | AACLSANMAECG | 1586.6146 | 1586.9 |

Total Ion Chromatogram

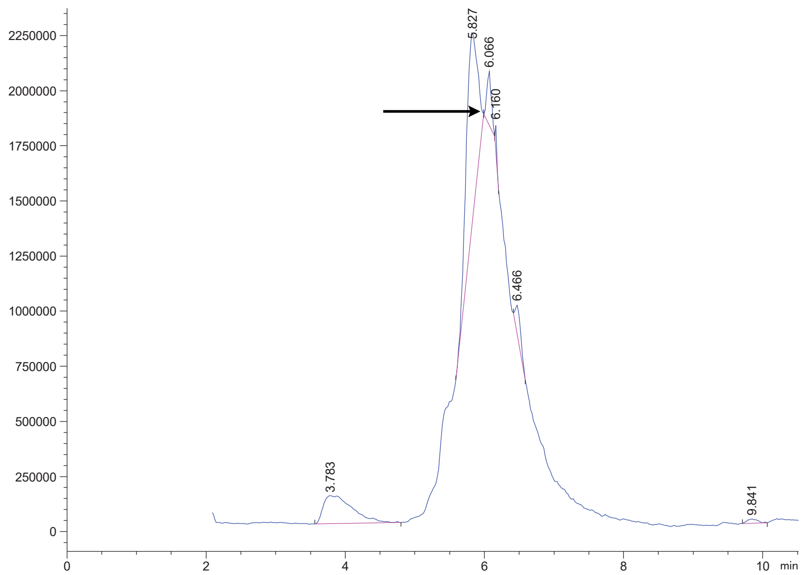


Variable Wavelength Detector: 220 nm

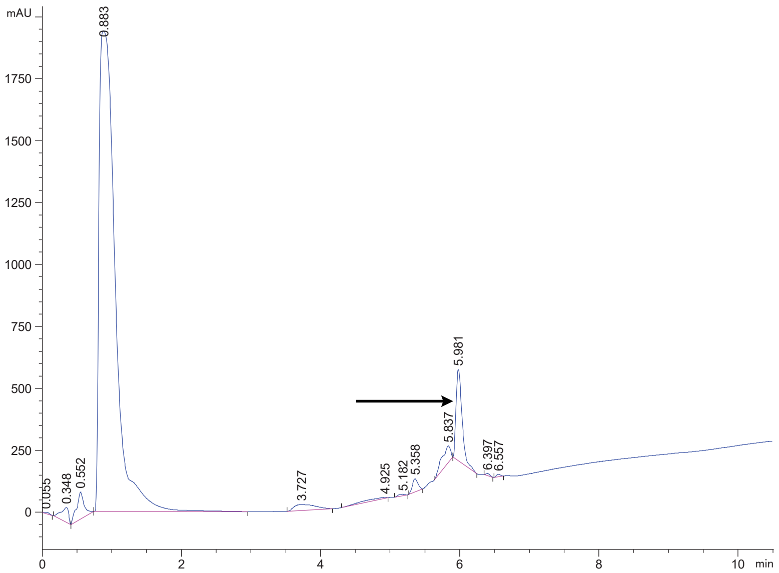

FAP 19 crude

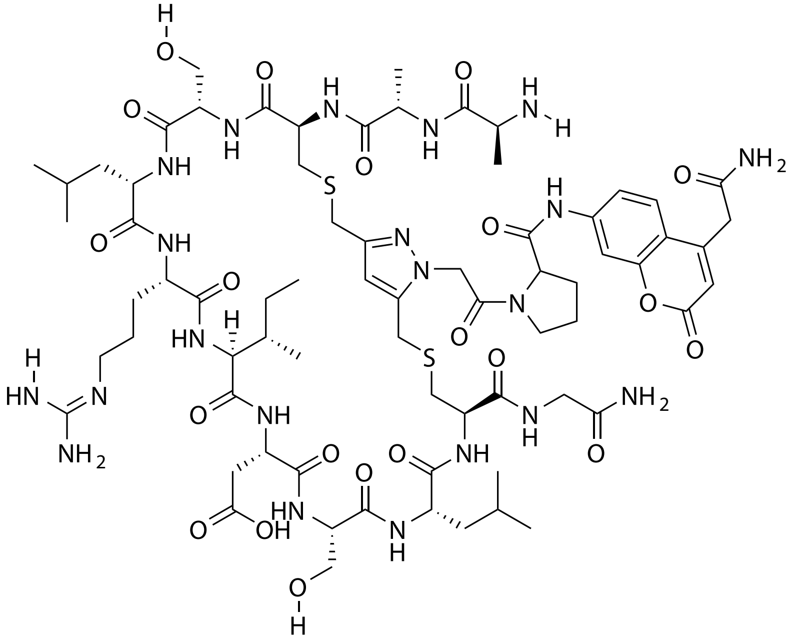

| Probe Name | Sequence | Expected Mass $[M+1]^{+1}$ or $[M+2]^{+2}$ | Found Mass |
|---|---|---|---|
| FAP 19 | AACSLRIDSLCG | 1654.7425 | 1655.0 |

Total Ion Chromatogram

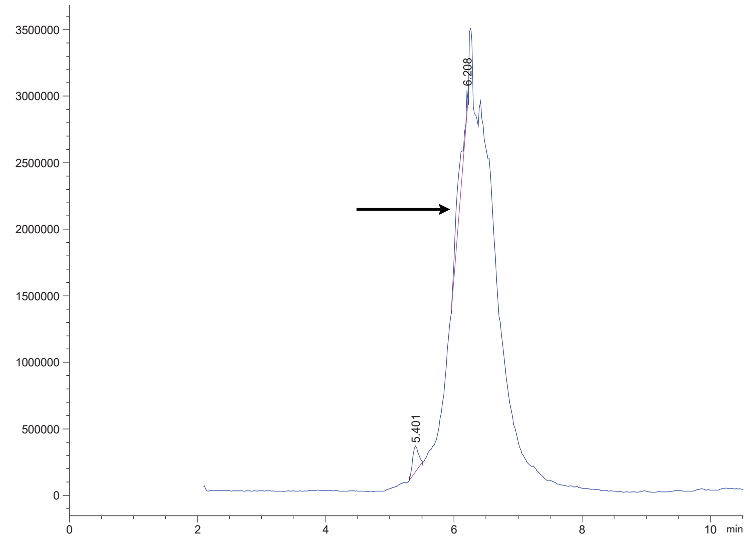


Variable Wavelength Detector: 220 nm

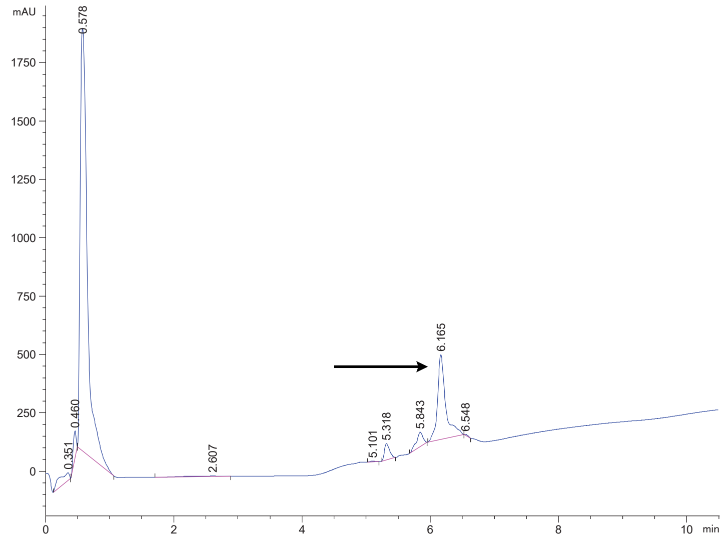

FAP 20 crude

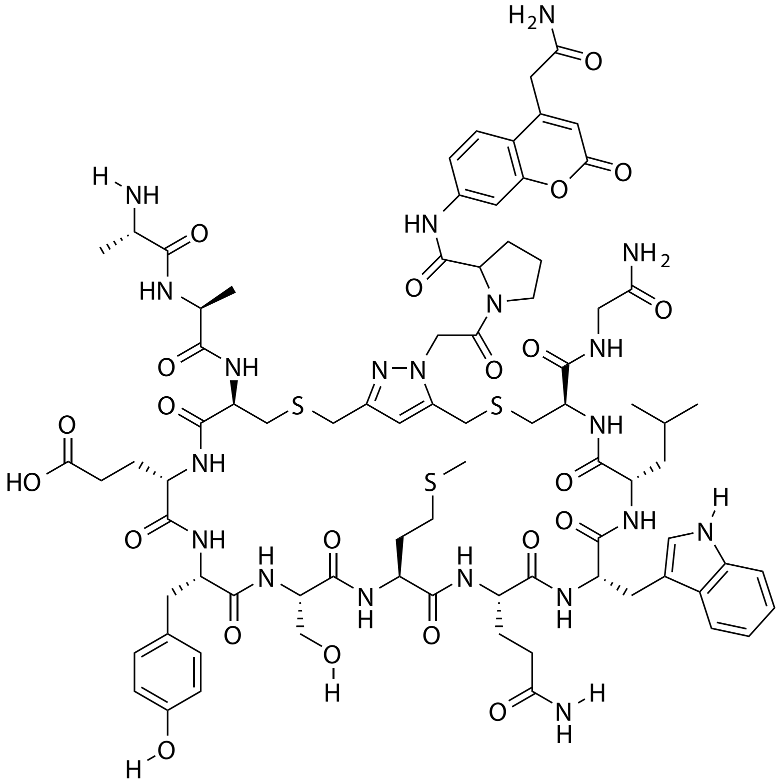

| Probe Name | Sequence | Expected Mass $[M+1]^{+1}$ or $[M+2]^{+2}$ | Found Mass |
|---|---|---|---|
| FAP 20 | AACEYSMQWLCG | 904.3533 $[M+2]^{+2}$ | 904.6 $[M+2]^{+2}$ |

Total Ion Chromatogram

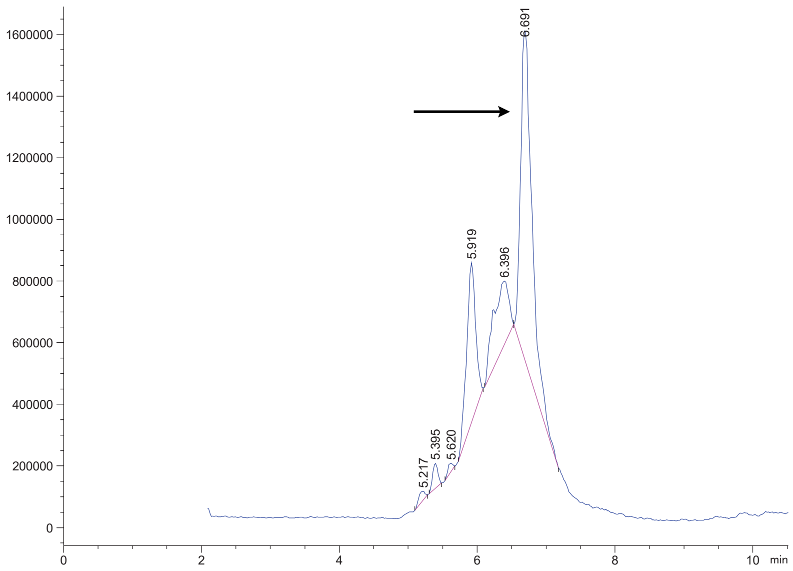

Variable Wavelength Detector: 220 nm

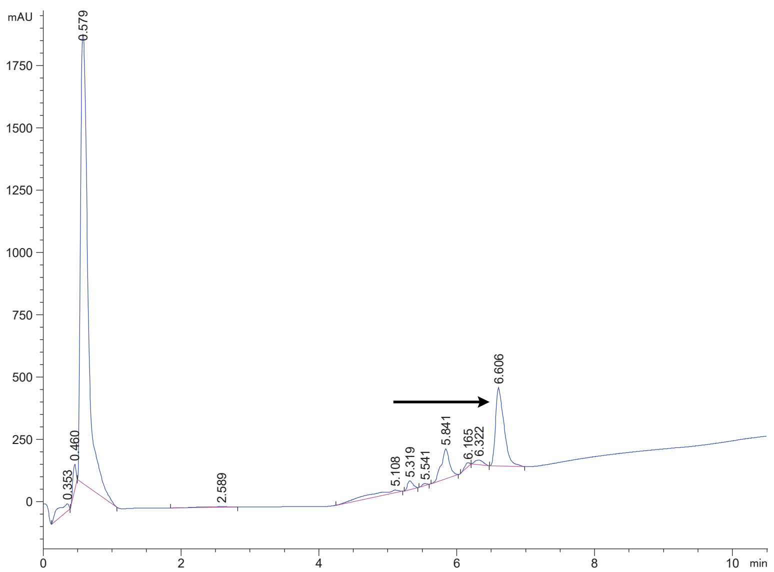

FAP 21 crude

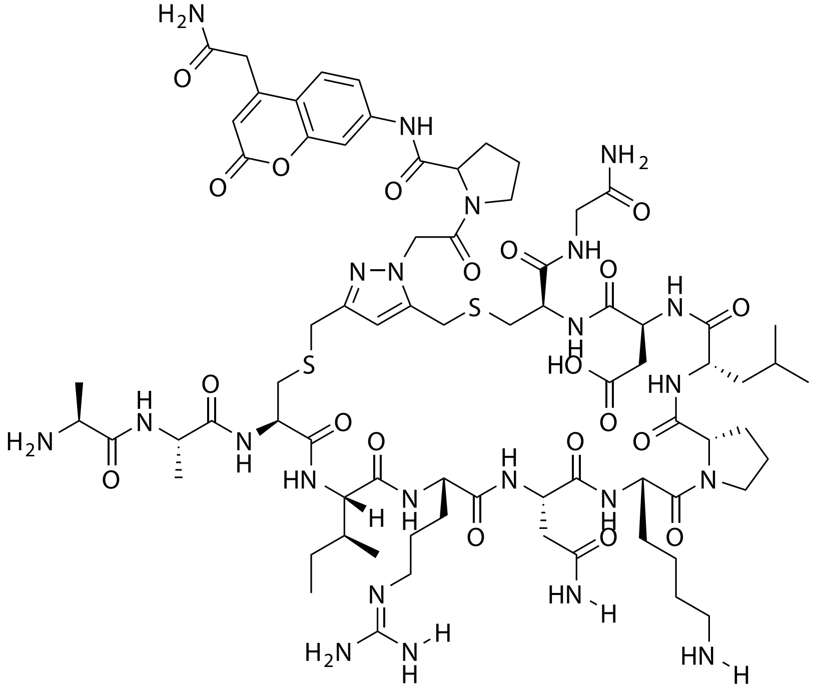

| Probe Name | Sequence | Expected Mass $[M+1]^{+1}$ or $[M+2]^{+2}$ | Found Mass |
|---|---|---|---|
| FAP 21 | AACIRNKPLDCG | 853.8965 $[M+2]^{+2}$ | 853.8 $[M+2]^{+2}$ |

Total Ion Chromatogram

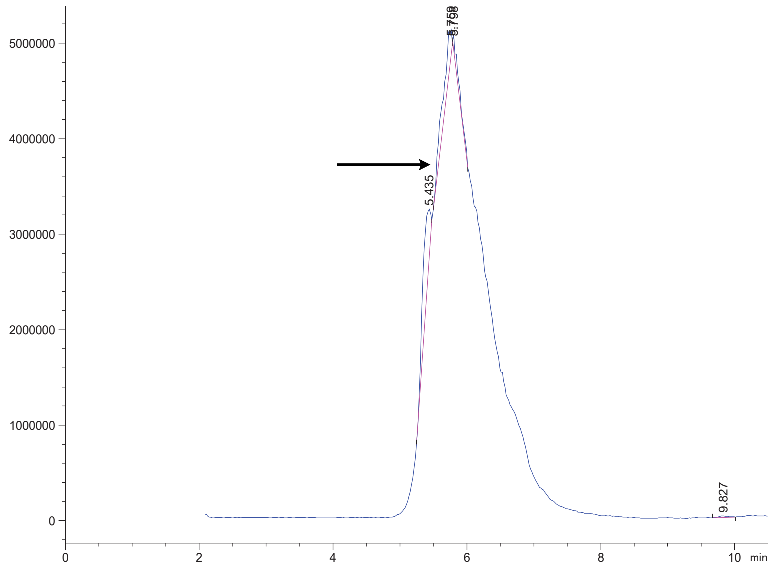

Variable Wavelength Detector: 220 nm

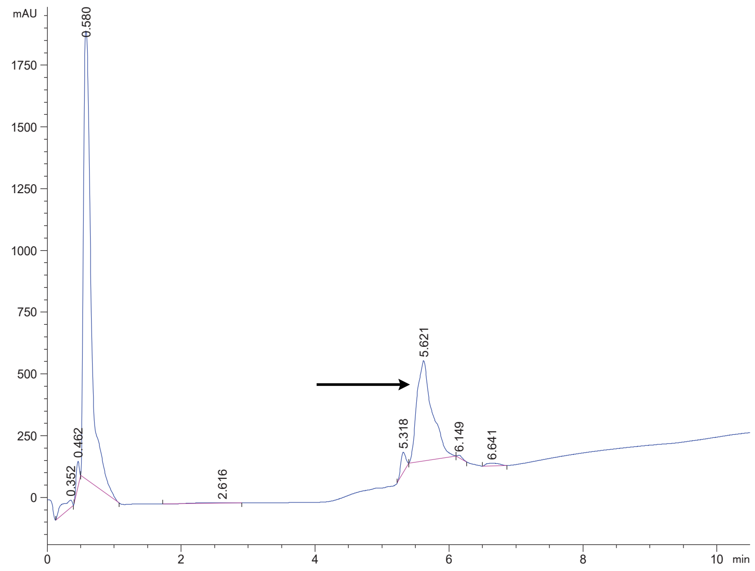

FAP 22 crude

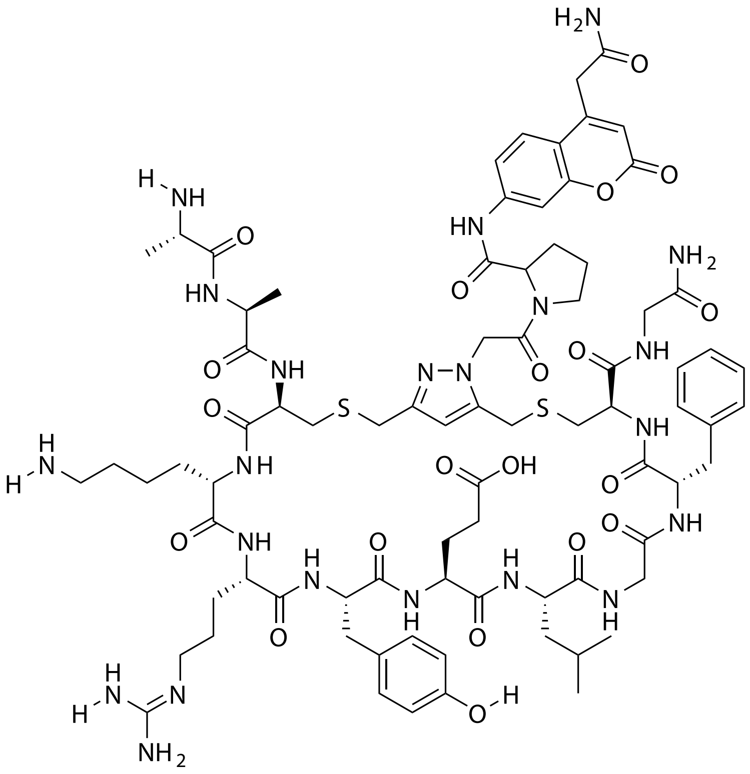

| Probe Name | Sequence | Expected Mass [M+1]$^{+1}$ or [M+2]$^{+2}$ | Found Mass |
|---|---|---|---|
| FAP 22 | AACKRYELGFCG | 882.3910 [M+2]$^{+2}$ | 882.2 [M+2]$^{+2}$ |

Total Ion Chromatogram

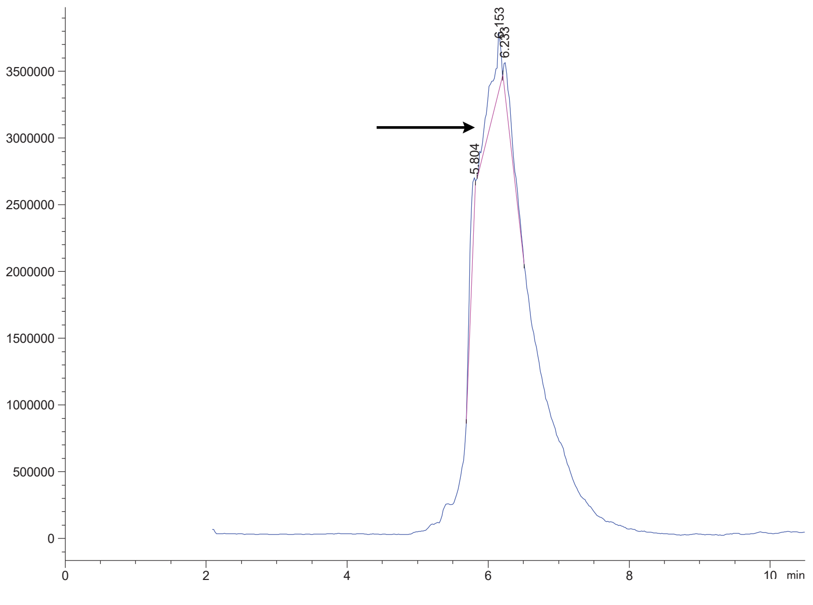

Variable Wavelength Detector: 220 nm

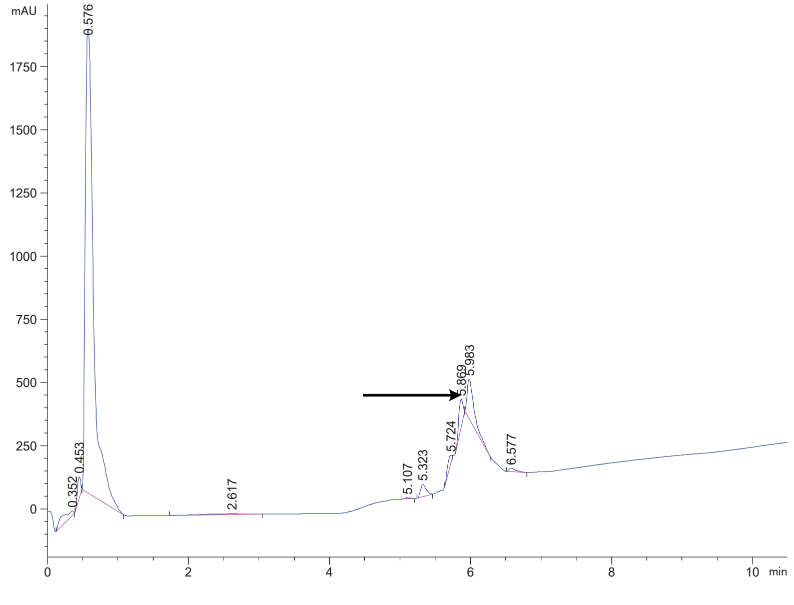

FAP 23 crude

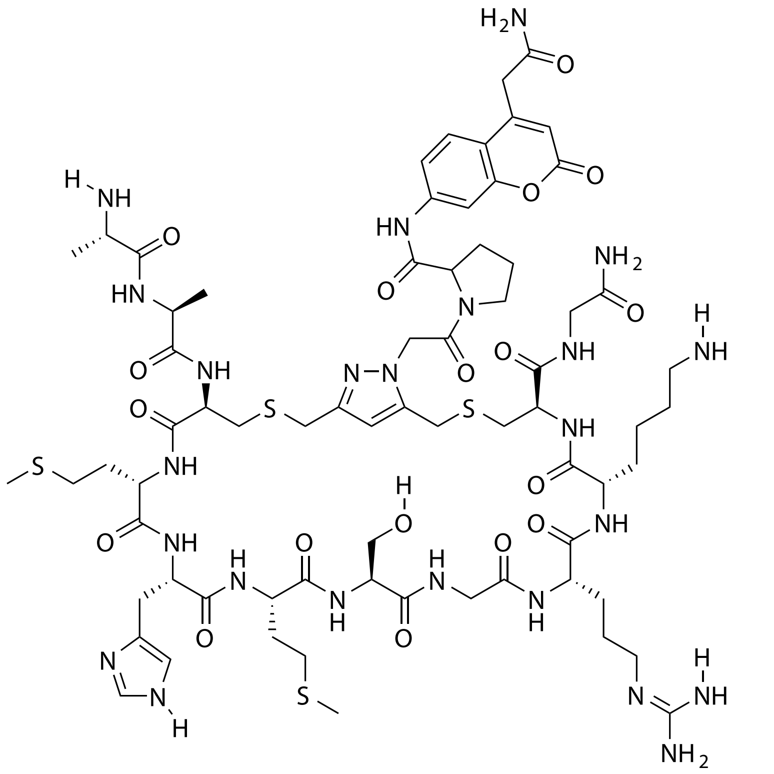

| Probe Name | Sequence | Expected Mass $[M+1]^{+1}$ or $[M+2]^{+2}$ | Found Mass |
|---|---|---|---|
| FAP 23 | AACMHMSGRKCG | 849.3478 $[M+2]^{+2}$ | 849.2 $[M+2]^{+2}$ |

Total Ion Chromatogram

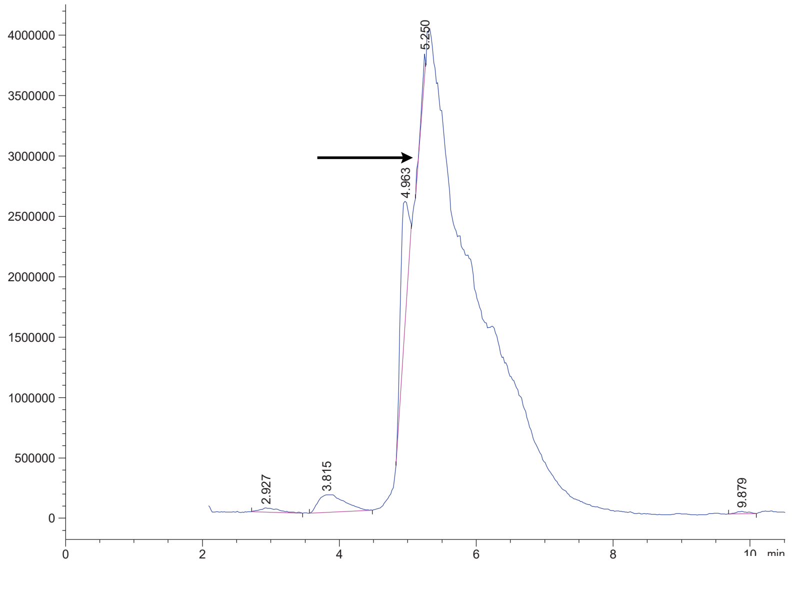

Variable Wavelength Detector: 220 nm

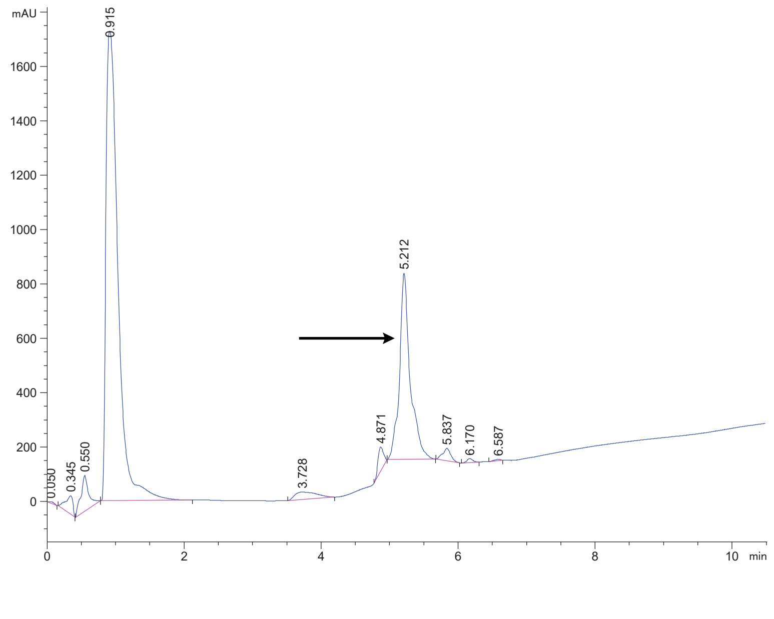

FAP 24 crude

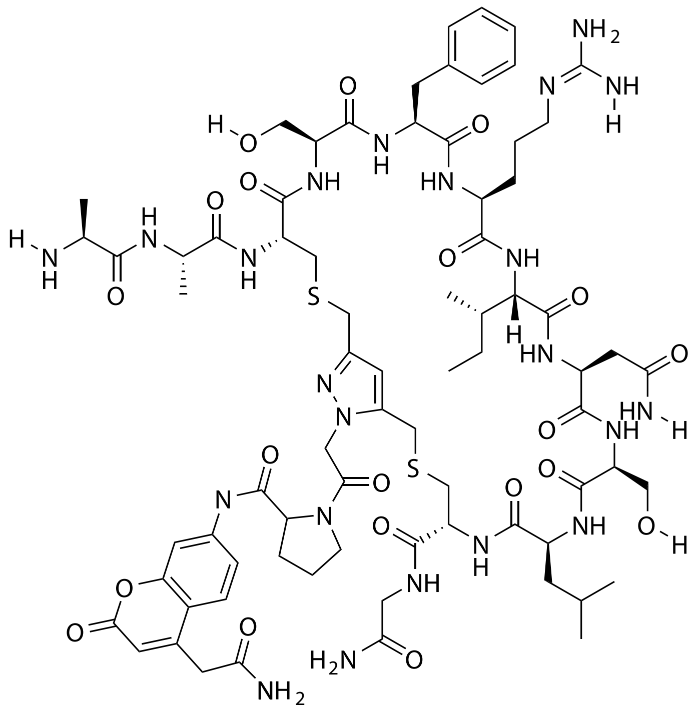

| Probe Name | Sequence | Expected Mass [M+1]$^{+1}$ or [M+2]$^{+2}$ | Found Mass |
|---|---|---|---|
| FAP 24 | AACSFRINSLCG | 844.3753 [M+2]$^{+2}$ | 844.2 [M+2]$^{+2}$ |

Total Ion Chromatogram

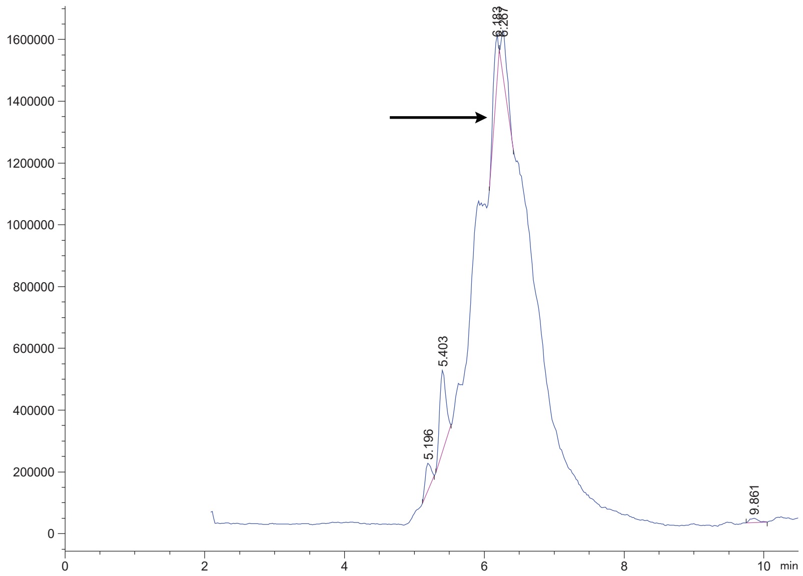

Variable Wavelength Detector: 220 nm

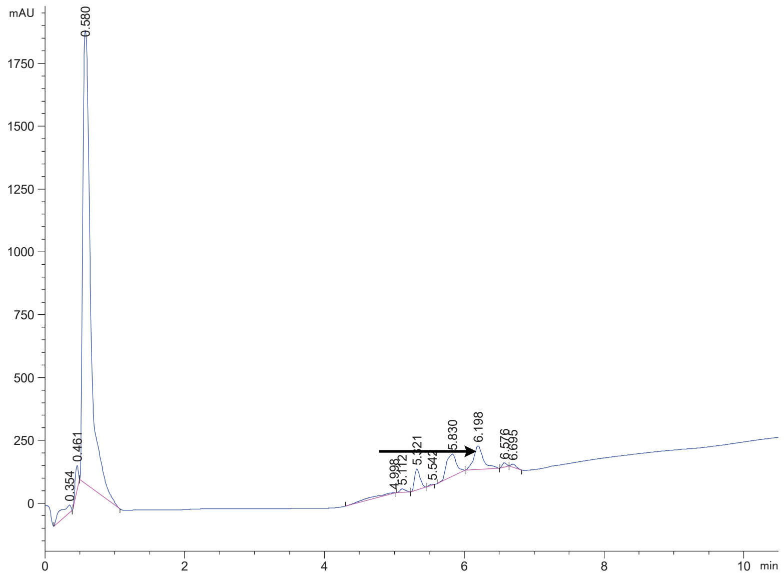

### FAP 25 crude

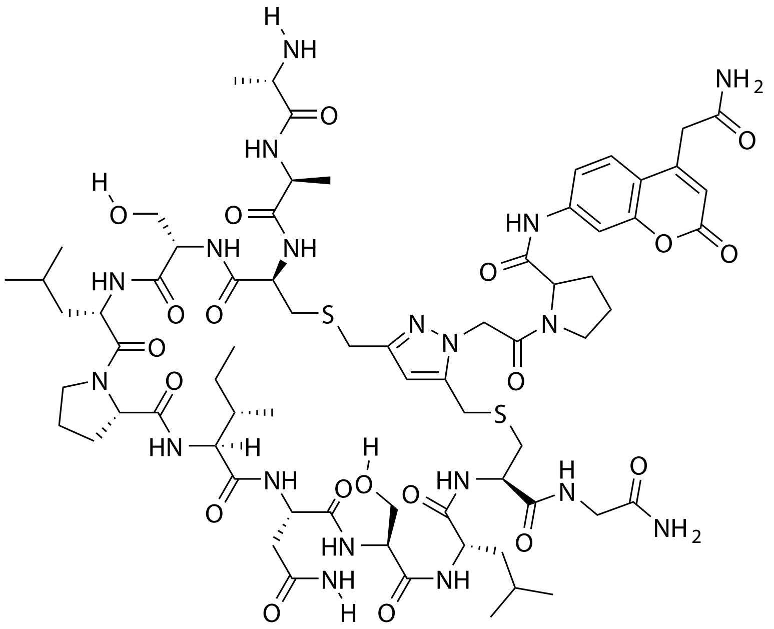


| Probe Name | Sequence | Expected Mass $[M+1]^{+1}$ or $[M+2]^{+2}$ | Found Mass |
|---|---|---|---|
| FAP 25 | AACSLPINSLCG | 1594.7102 | 1595.9 |

Total Ion Chromatogram

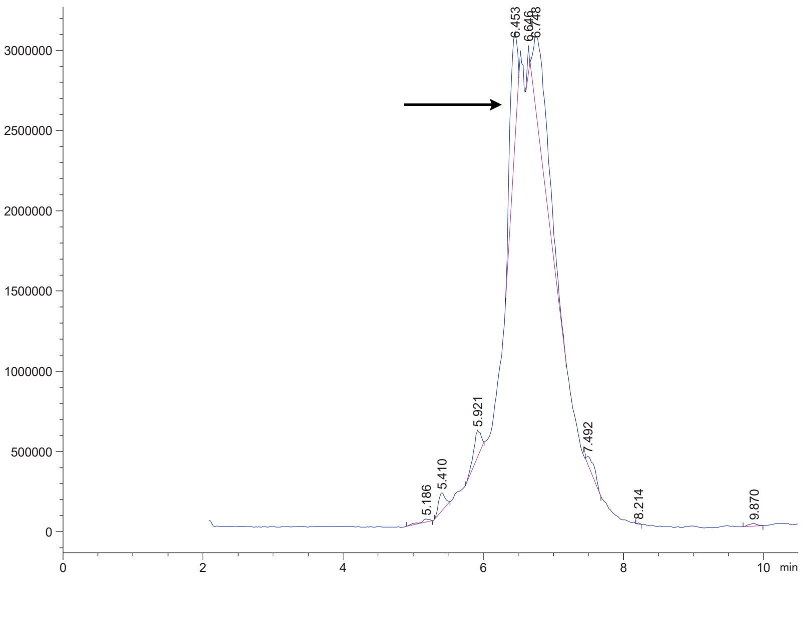


Variable Wavelength Detector: 220 nm

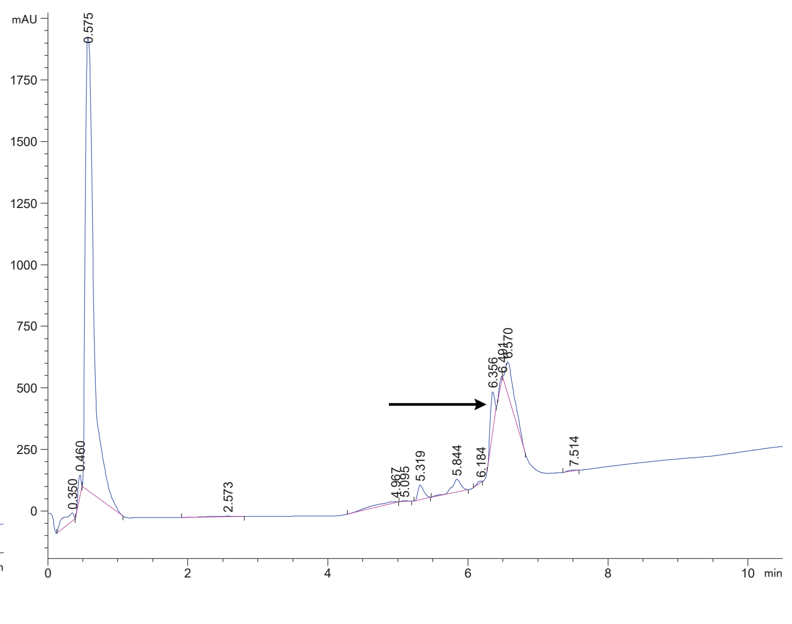

FAP 26 crude

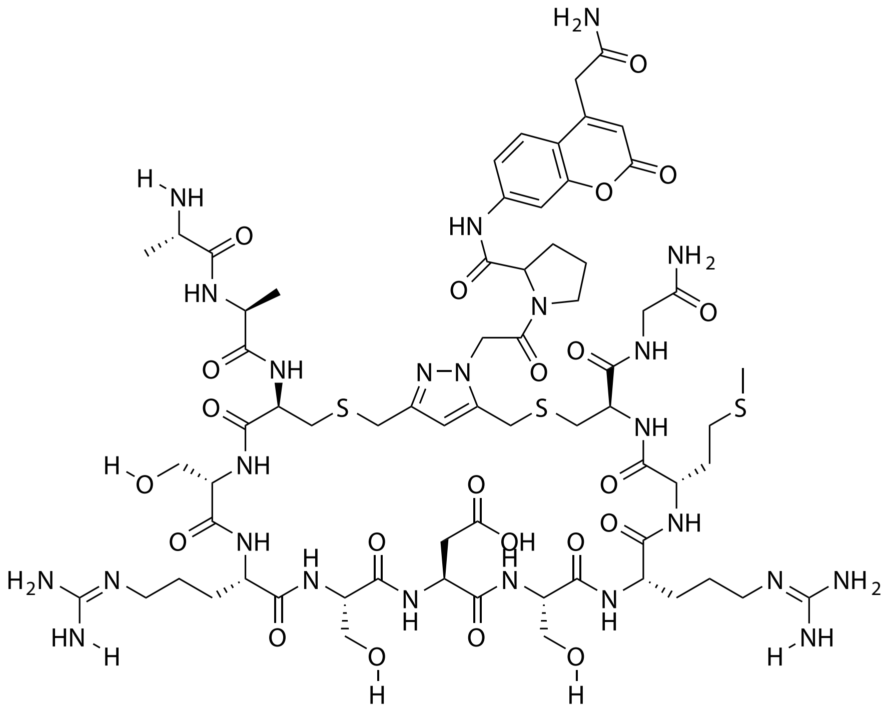

| **Probe Name** | **Sequence** | **Expected Mass $[M+1]^{+1}$ or $[M+2]^{+2}$** | **Found Mass** |
|---|---|---|---|
| FAP 26 | AACSRSDSRMCG | 845.3359 $[M+2]^{+2}$ | 845.2 $[M+2]^{+2}$ |

Total Ion Chromatogram

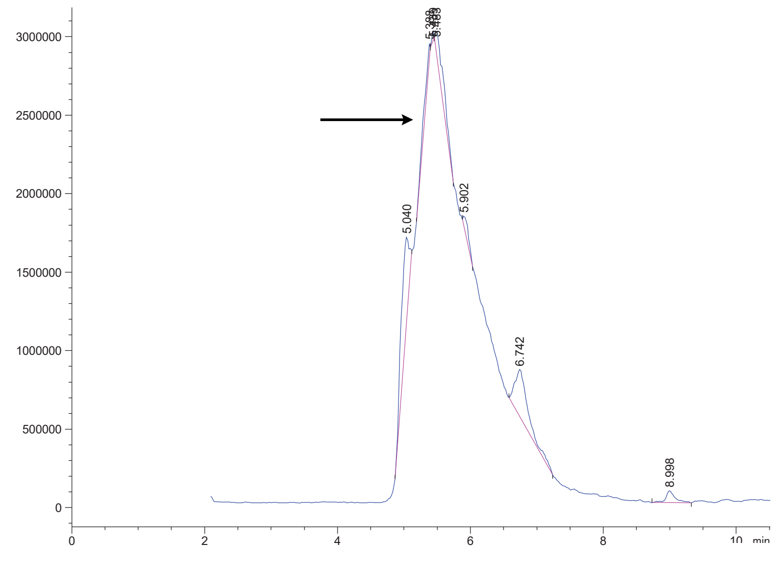


Variable Wavelength Detector: 220 nm

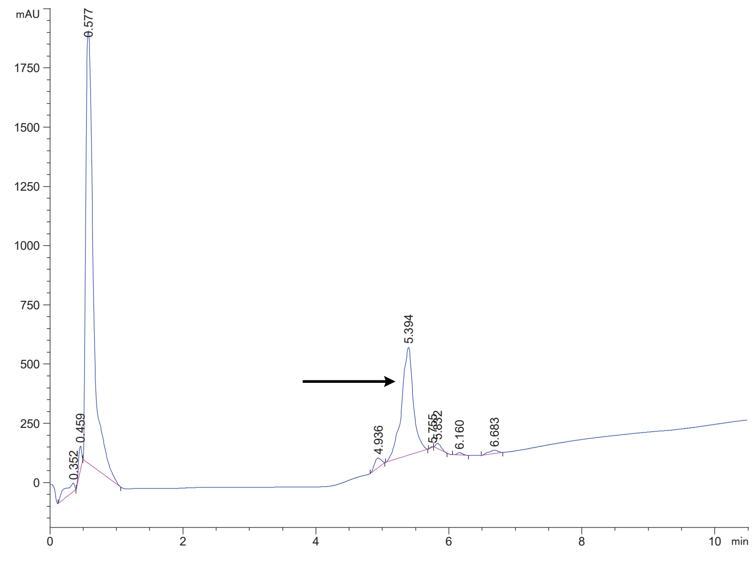

FAP 27 crude

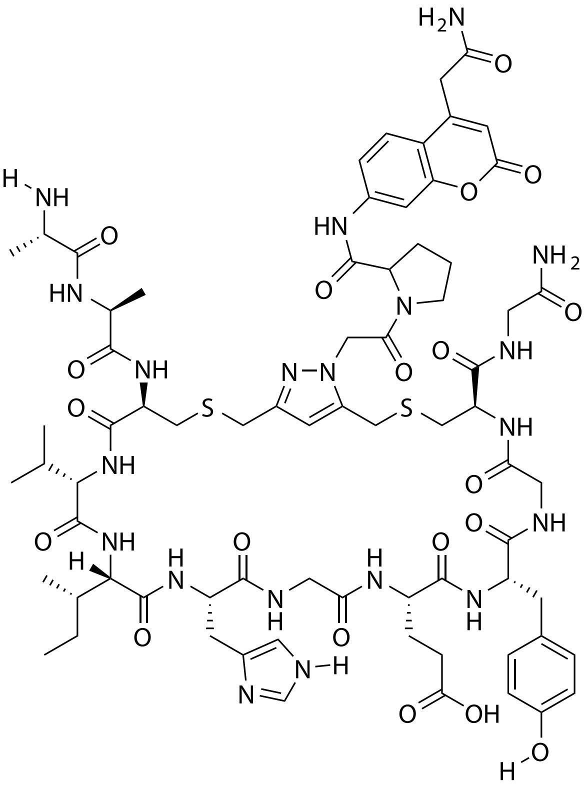

| Probe Name | Sequence | Expected Mass [M+1]$^{+1}$ or [M+2]$^{+2}$ | Found Mass |
|---|---|---|---|
| FAP 27 | AACVIHGEYGCG | 1625.6585 | 1625.9 |

Total Ion Chromatogram

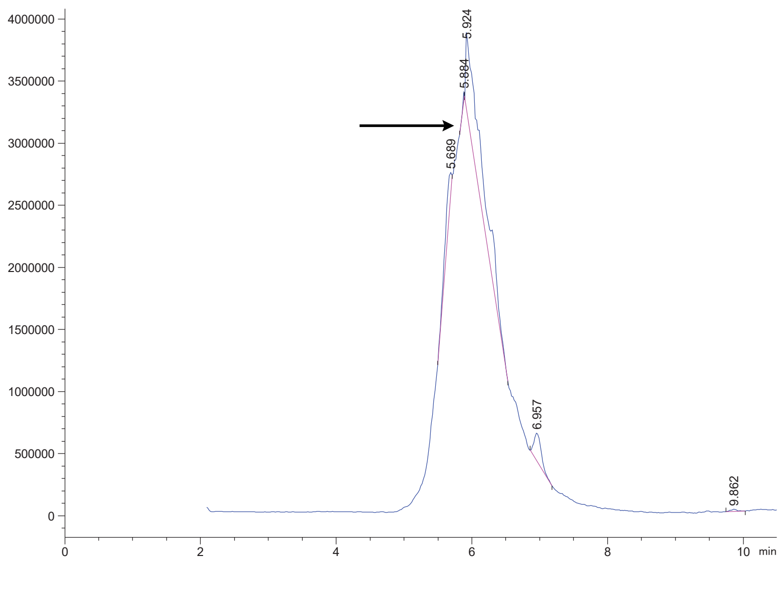

Variable Wavelength Detector: 220 nm

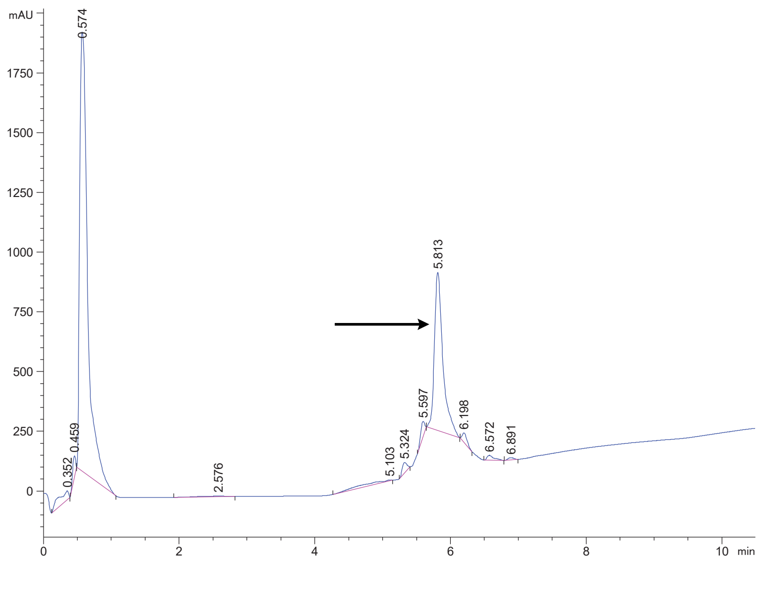

FAP 28 crude

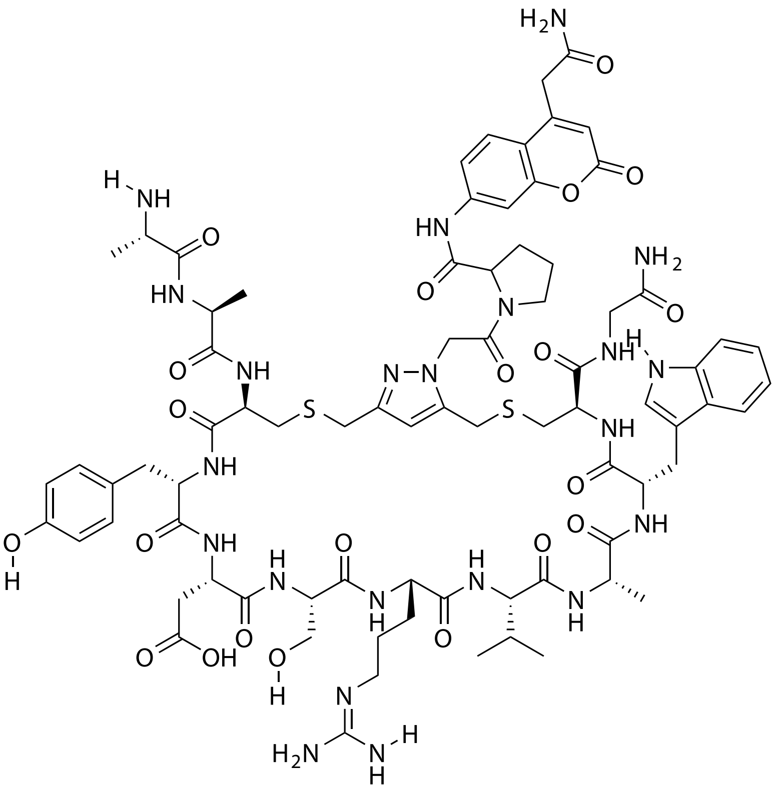

| Probe Name | Sequence | Expected Mass [M+1]$^{+1}$ or [M+2]$^{+2}$ | Found Mass |
|---|---|---|---|
| FAP 28 | AACYDSRVAWCG | 874.3572 [M+2]$^{+2}$ | 874.2 [M+2]$^{+2}$ |

Total Ion Chromatogram

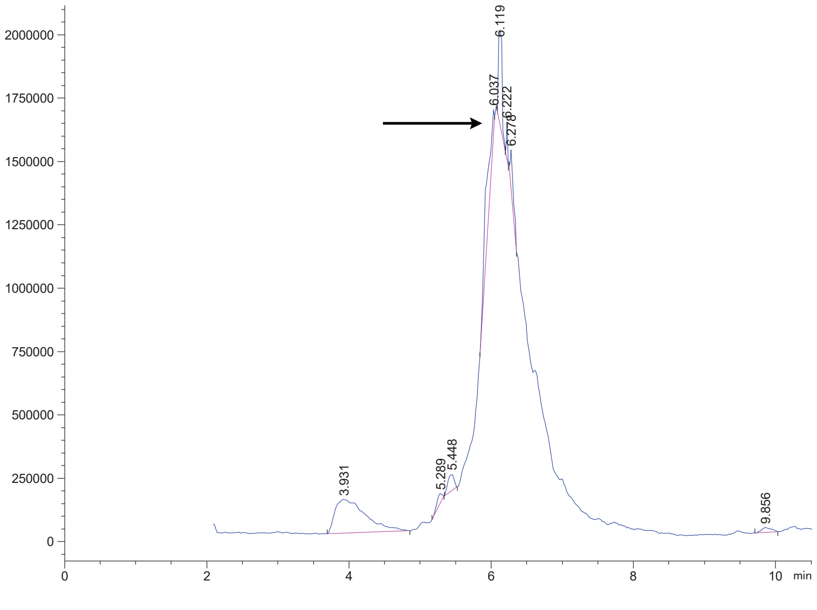


Variable Wavelength Detector: 220 nm

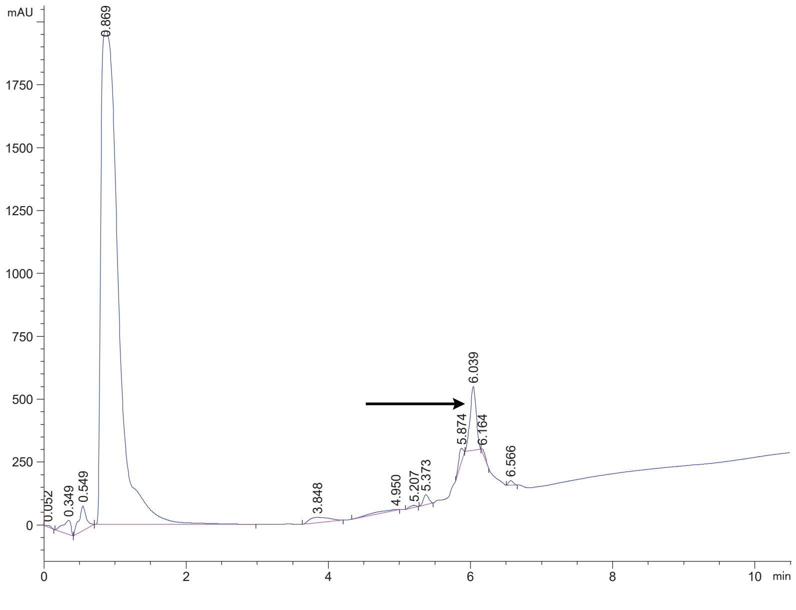

FAP 29 crude

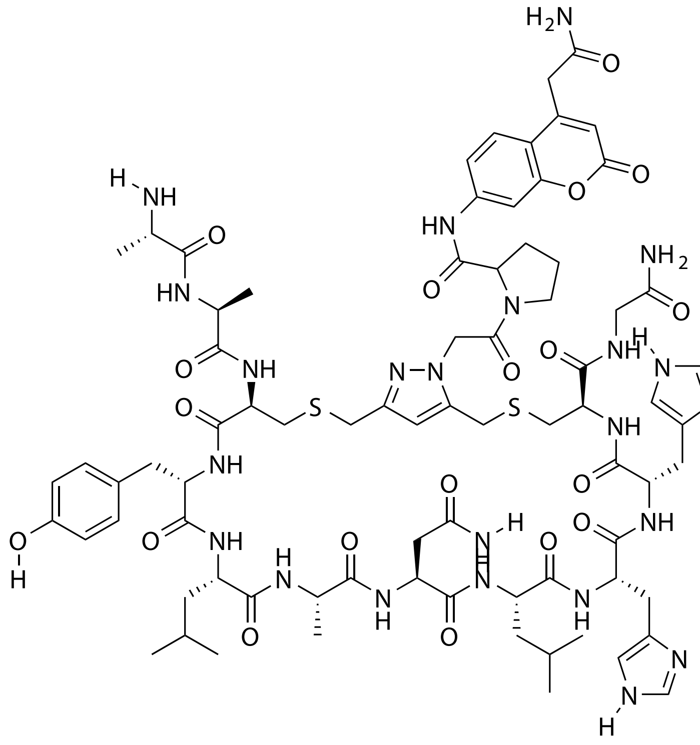

| Probe Name | Sequence | Expected Mass [M+1]$^{+1}$ or [M+2]$^{+2}$ | Found Mass |
|---|---|---|---|
| FAP 29 | AACYLANLHHCG | 859.8677 [M+2]$^{+2}$ | 859.8 [M+2]$^{+2}$ |

Total Ion Chromatogram

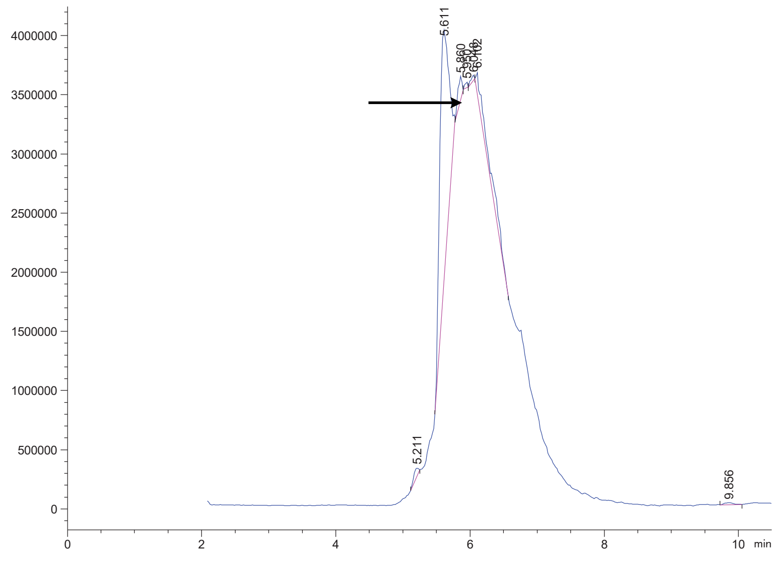

Variable Wavelength Detector: 220 nm

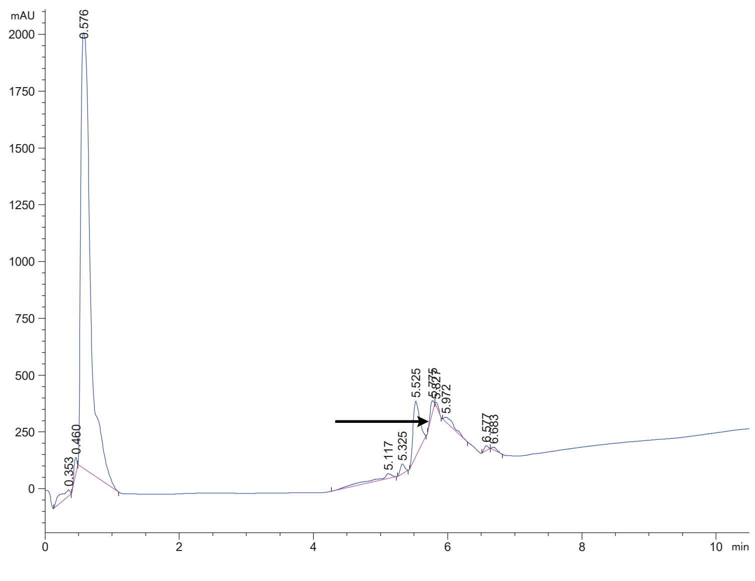

FAP 30 crude

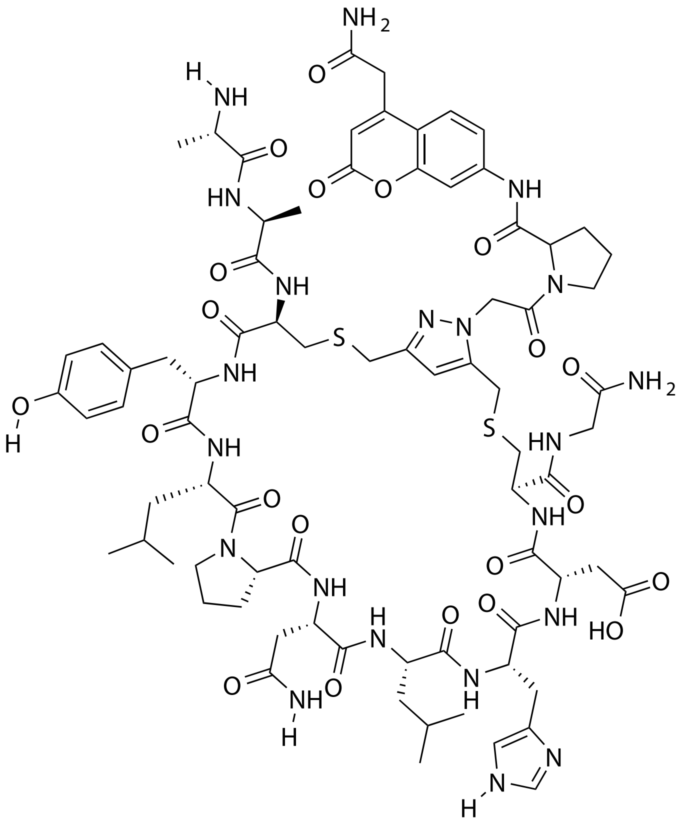

| Probe Name | Sequence | Expected Mass $[M+1]^{+1}$ or $[M+2]^{+2}$ | Found Mass |
|---|---|---|---|
| FAP 30 | AACYLPNLHDCG | 1722.7112 | 1722.9 |

Total Ion Chromatogram

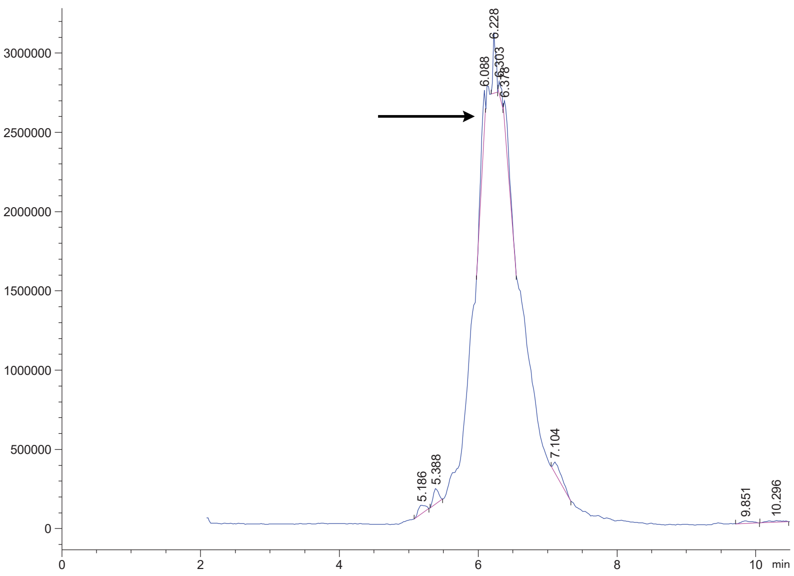


Variable Wavelength Detector: 220 nm

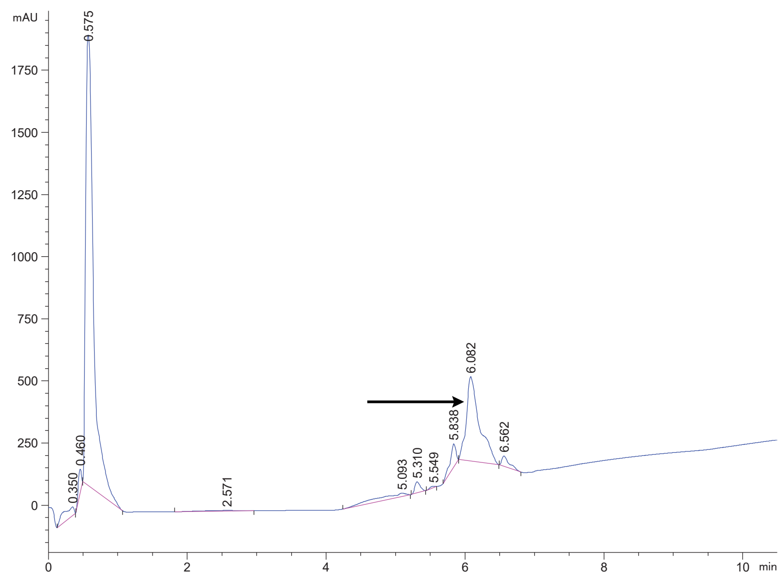

FAP 31 crude

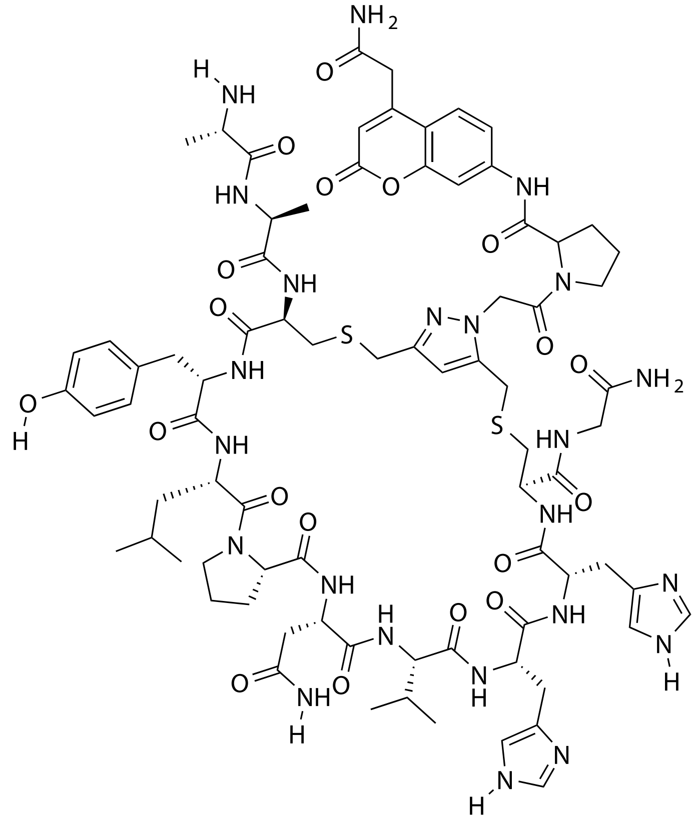

| Probe Name | Sequence | Expected Mass $[M+1]^{+1}$ or $[M+2]^{+2}$ | Found Mass |
|---|---|---|---|
| FAP 31 | AACYLPNVHHCG | 865.8677 $[M+2]^{+2}$ | 865.8 $[M+2]^{+2}$ |

Total Ion Chromatogram

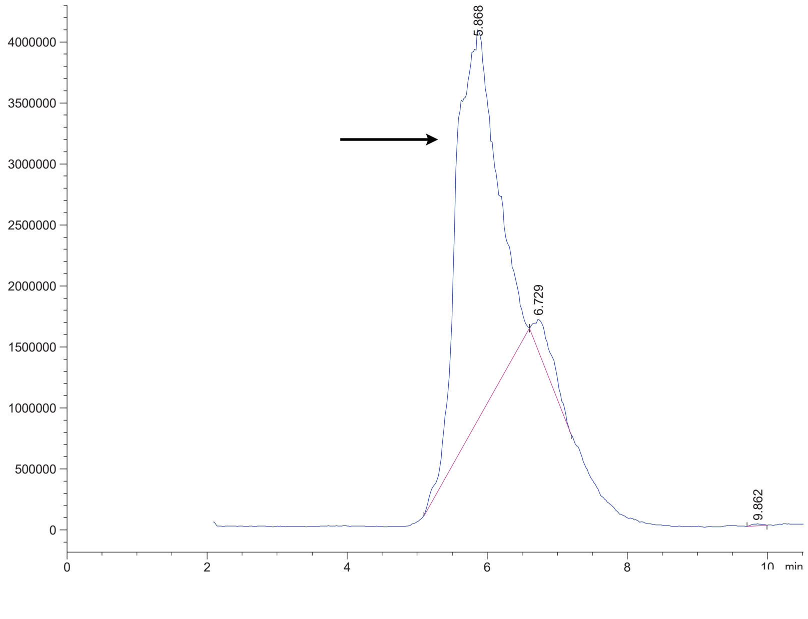

Variable Wavelength Detector: 220 nm

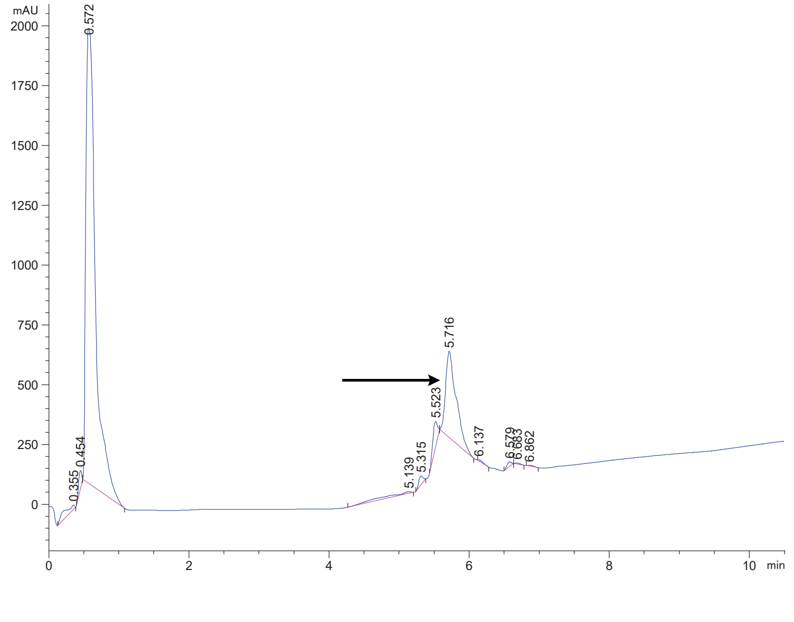

FAP 32 crude

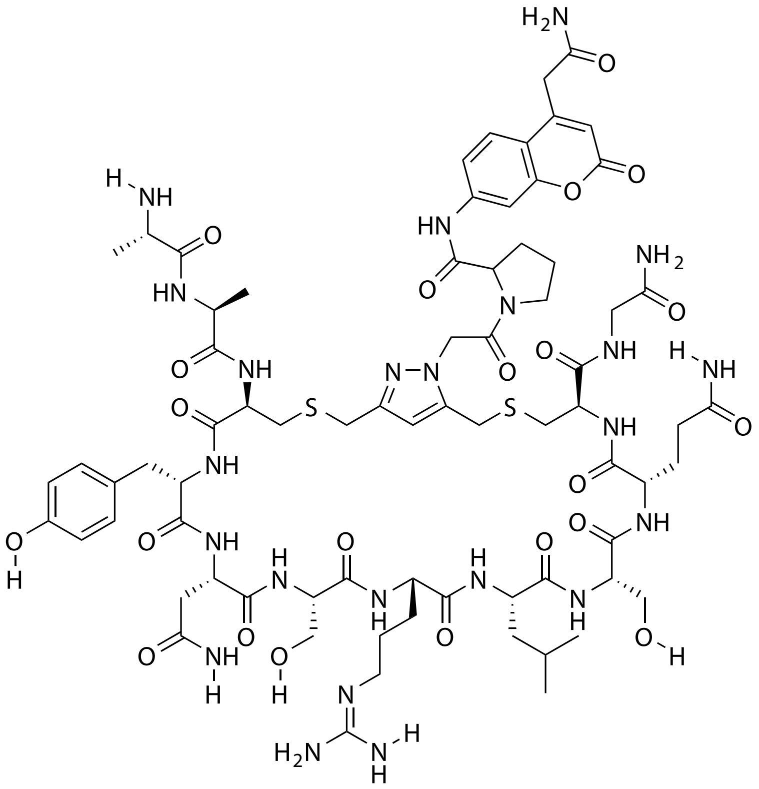

| Probe Name | Sequence | Expected Mass $[M+1]^{+1}$ or $[M+2]^{+2}$ | Found Mass |
|---|---|---|---|
| FAP 32 | AACYNSRLSQCG | 1718.7123 | 1717.9 |

Total Ion Chromatogram

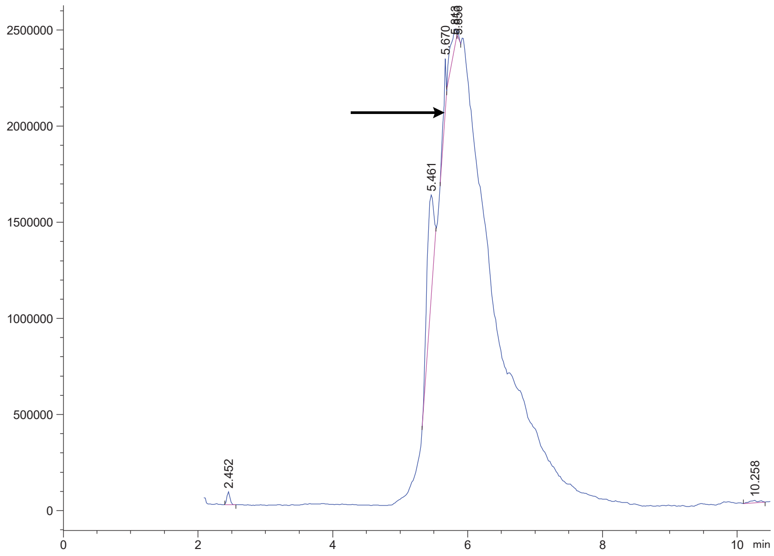

Variable Wavelength Detector: 220 nm

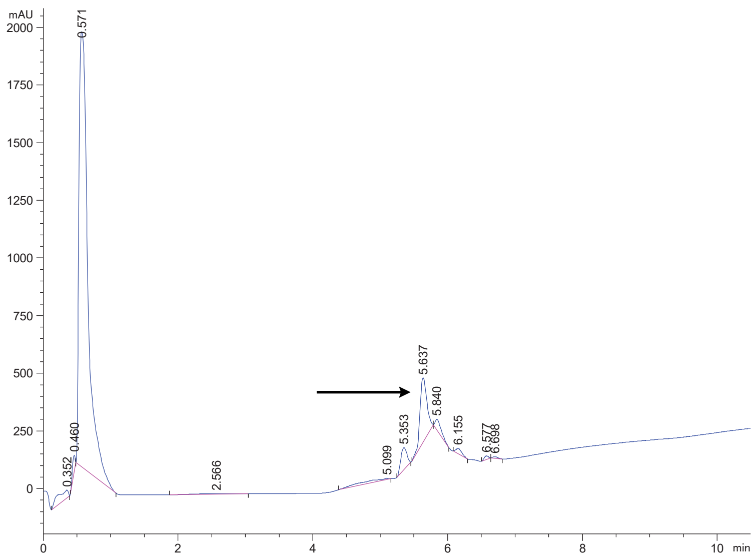

FAP 33 crude

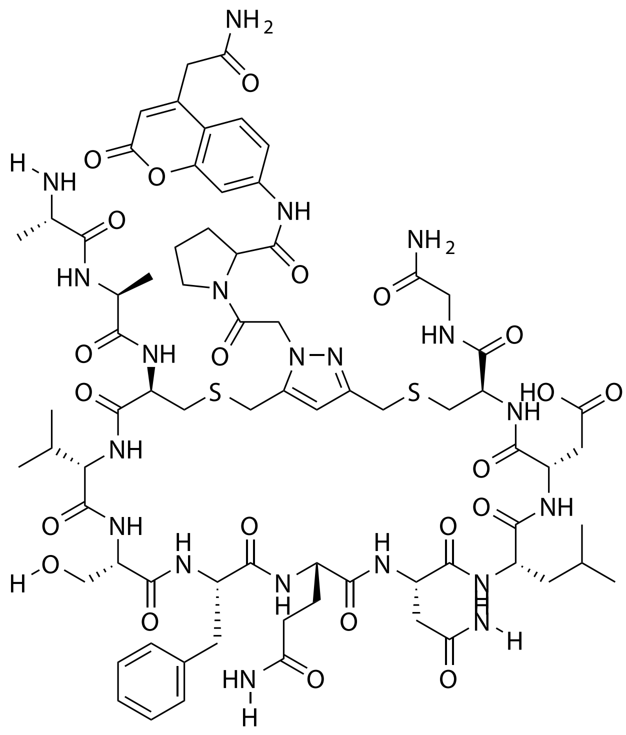


| Probe Name | Sequence | Expected Mass $[M+1]^{+1}$ or $[M+2]^{+2}$ | Found Mass |
|---|---|---|---|
| FAP 33 | AACVSFQNLDCG | 837.3437 | 837.2 $[M+2]^{+2}$ |

Total Ion Chromatogram

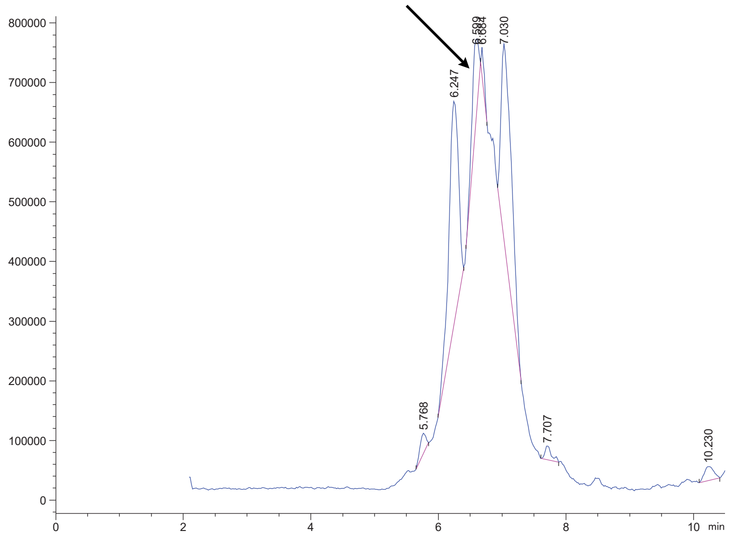


Variable Wavelength Detector: 220 nm

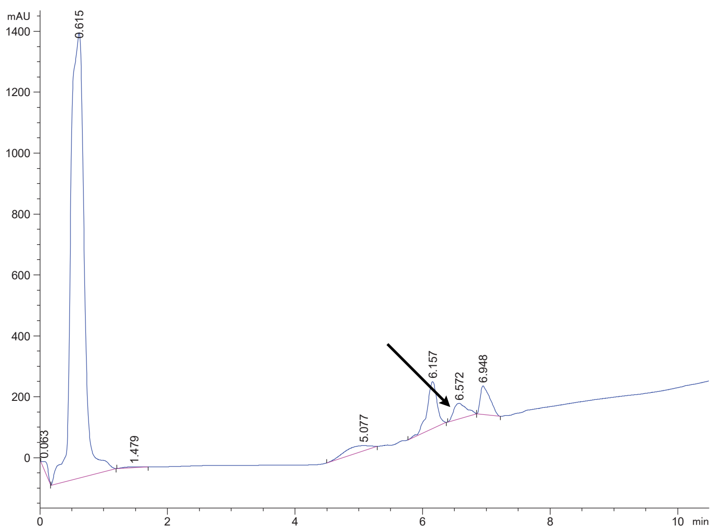

### Purified Molecules

Bio-orthogonal Linker

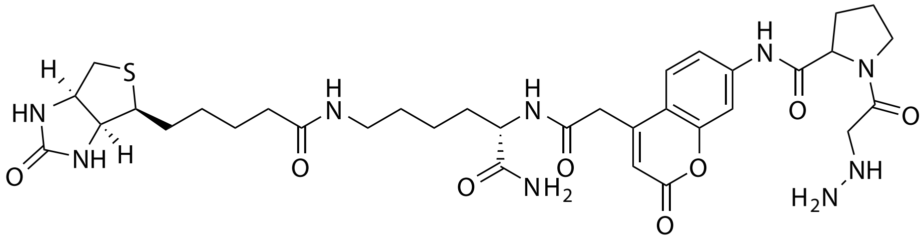


| Probe Name | Sequence | Expected Mass $[M+1]^{+1}$ or $[M+2]^{+2}$ | Found Mass |
|---|---|---|---|
| Bio-orthogonal linker | --- | 742.3302 | 742.5 |

Total Ion Chromatogram

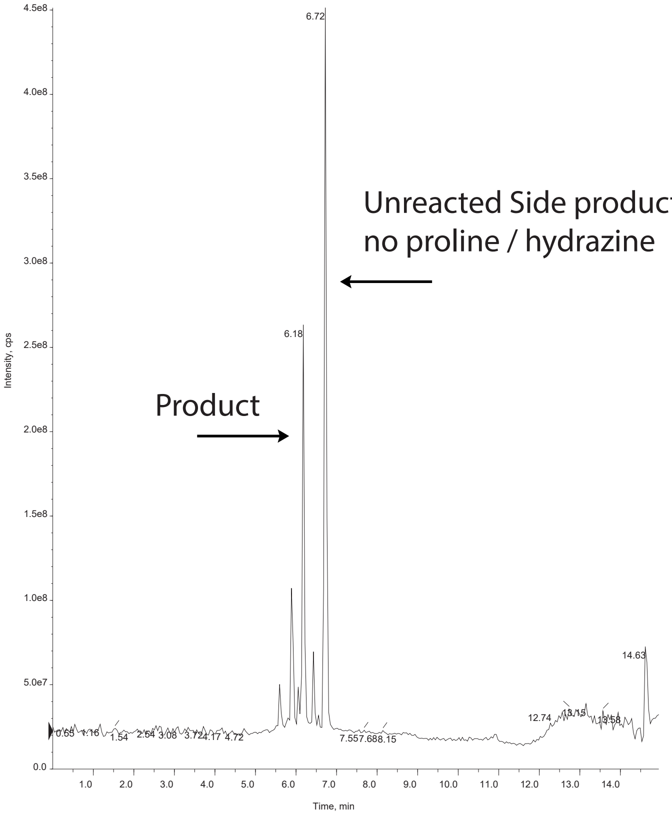


Variable Wavelength Detector: 220 nm

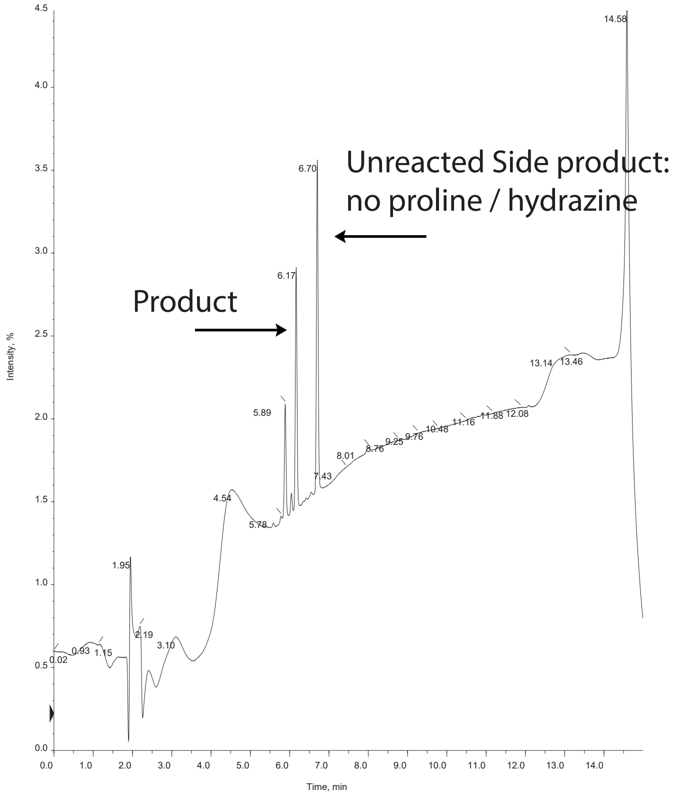

ACC Linker

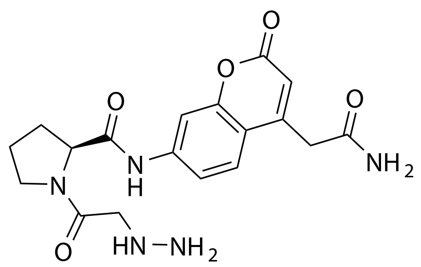


| Probe Name | Sequence | Expected Mass $[M+1]^{+1}$ or $[M+2]^{+2}$ | Found Mass |
|---|---|---|---|
| Bio-orthogonal linker | --- | 388.1576 | 388.0 |

Total Ion Chromatogram

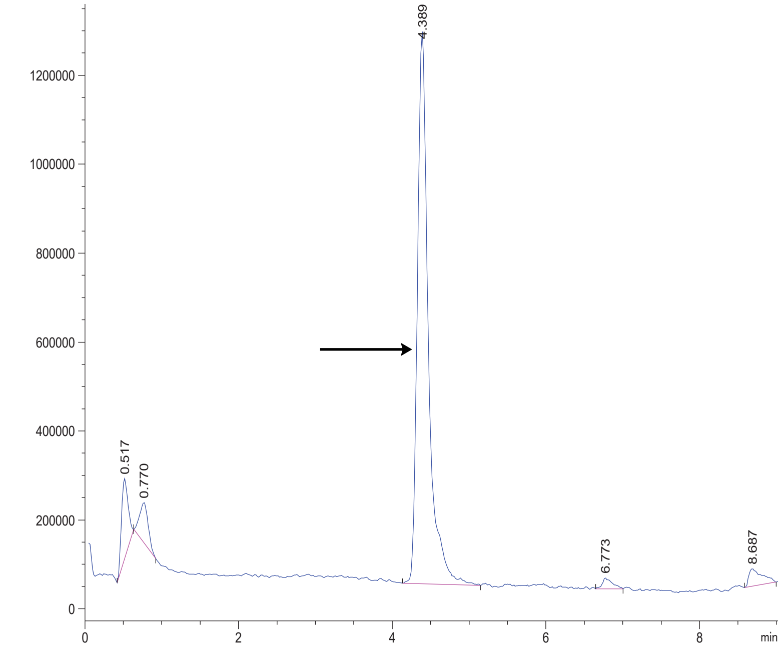


Variable Wavelength Detector: 220 nm

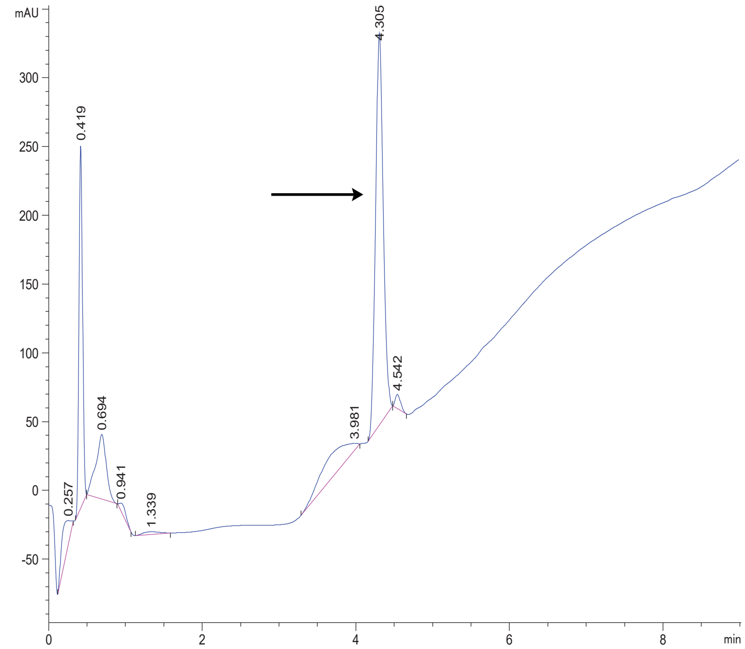

DPP4 1

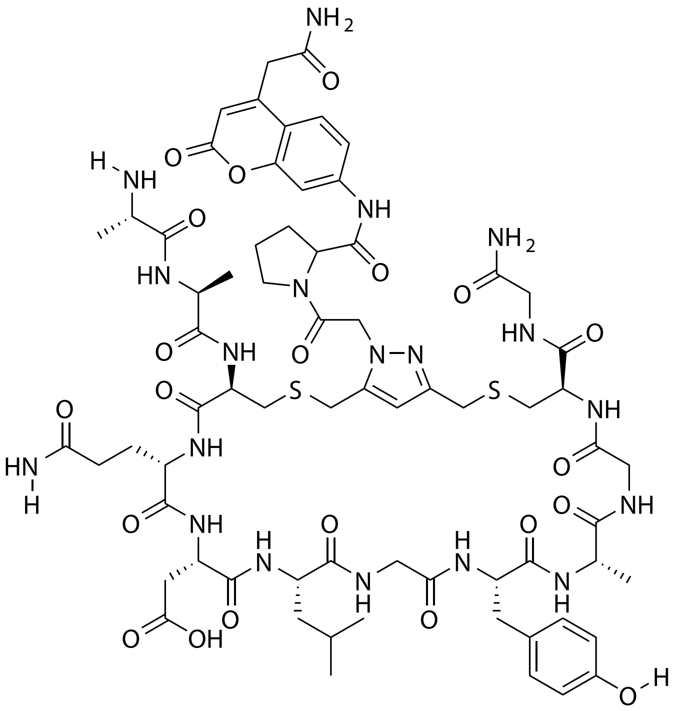


| Probe Name | Sequence | Expected Mass $[M+1]^{+1}$ or $[M+2]^{+2}$ | Found Mass |
|---|---|---|---|
| DPP4 1 | AACQDLGYAGCG | 1574.6112 | 1574.9 |

Total Ion Chromatogram

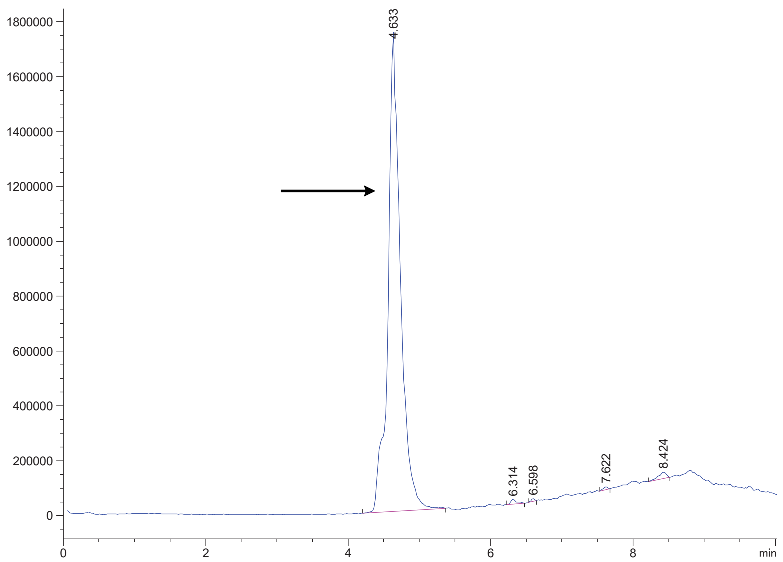


Variable Wavelength Detector: 220 nm

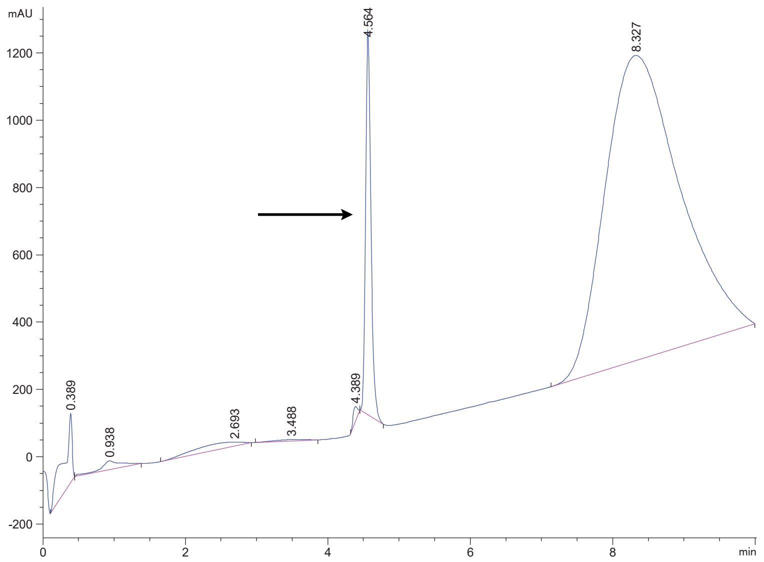

DPP4 7

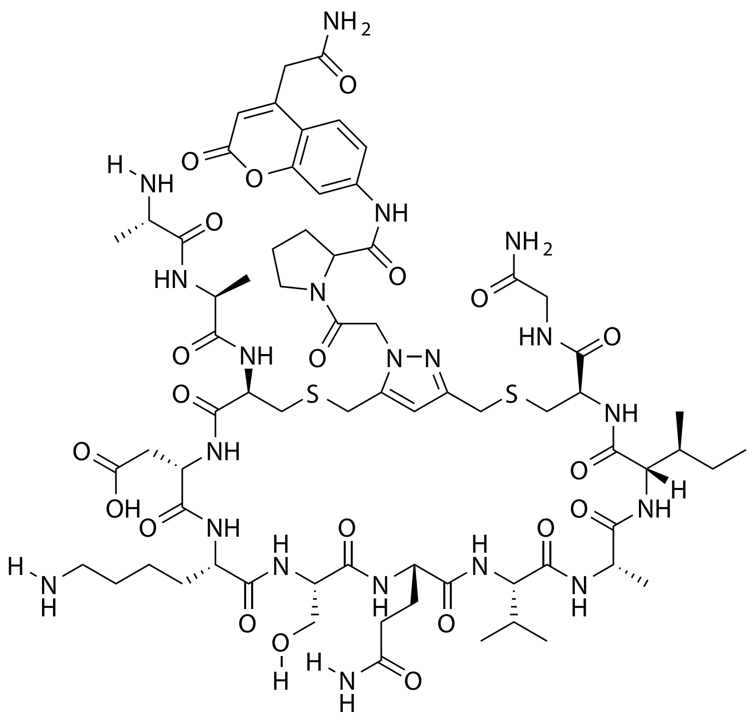

| Probe Name | Sequence | Expected Mass $[M+1]^{+1}$ or $[M+2]^{+2}$ | Found Mass |
|---|---|---|---|
| DPP4 7 | AACDKSQVAICG | 806.3540 $[M+2]^{+2}$ | 806.3 $[M+2]^{+2}$ |

Total Ion Chromatogram

Variable Wavelength Detector: 220 nm

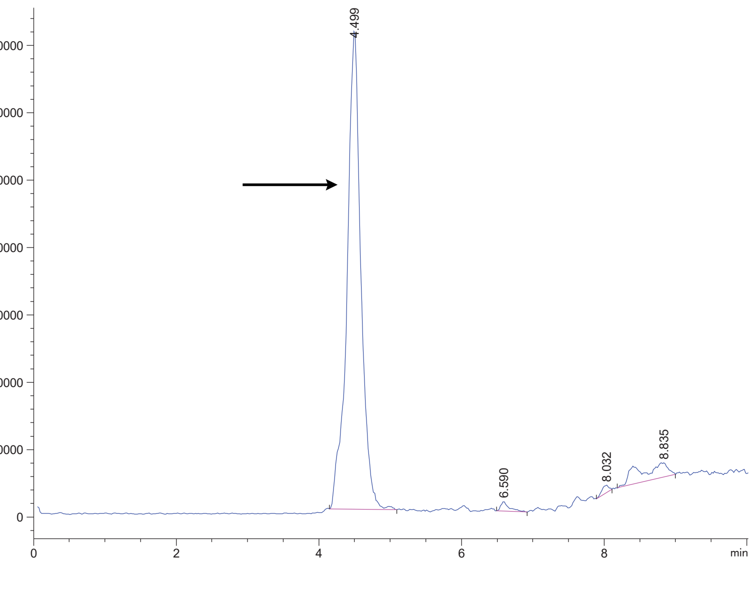


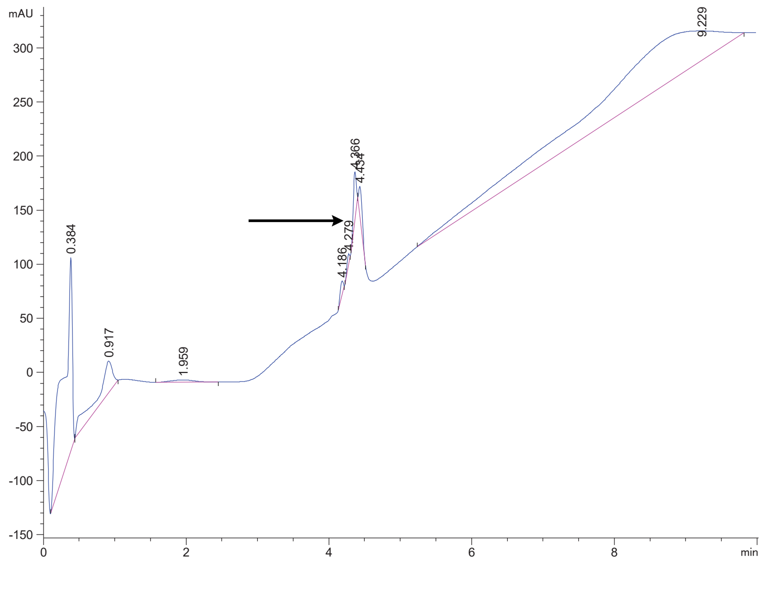

DPP4 10 AAs

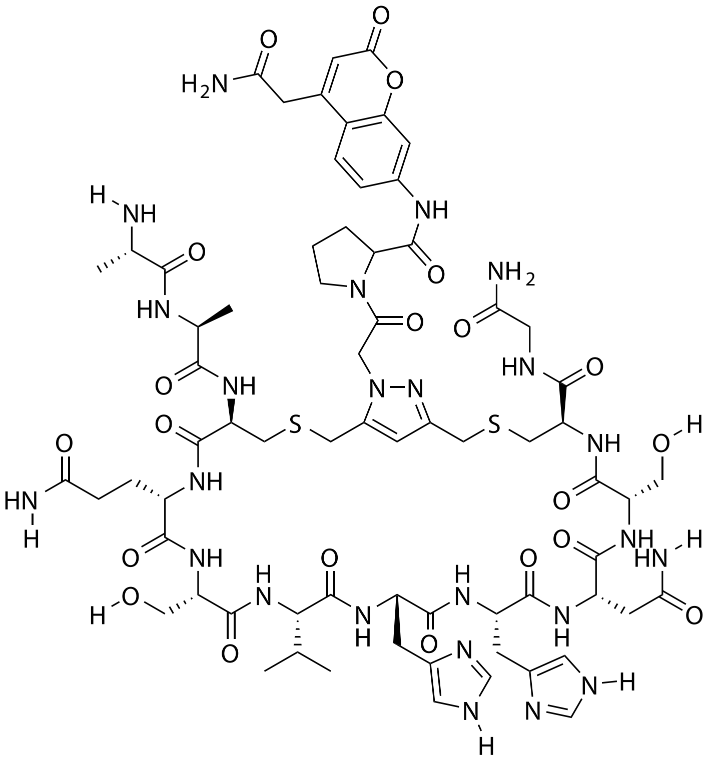

| Probe Name | Sequence | Expected Mass $[M+1]^{+1}$ or $[M+2]^{+2}$ | Found Mass |
|---|---|---|---|
| DPP4 10 AAs | AACQSVHHNSCG | 1659.6500 | 1660.2 |

Total Ion Chromatogram

Variable Wavelength Detector: 220 nm

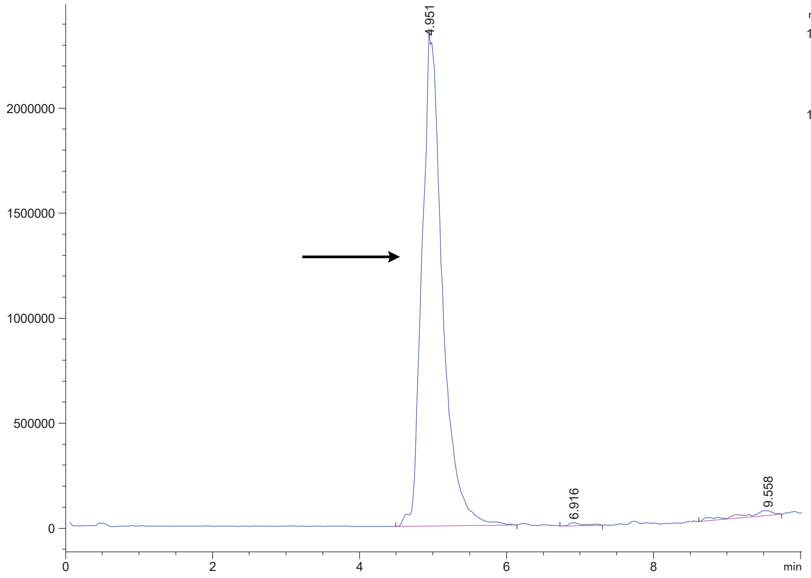


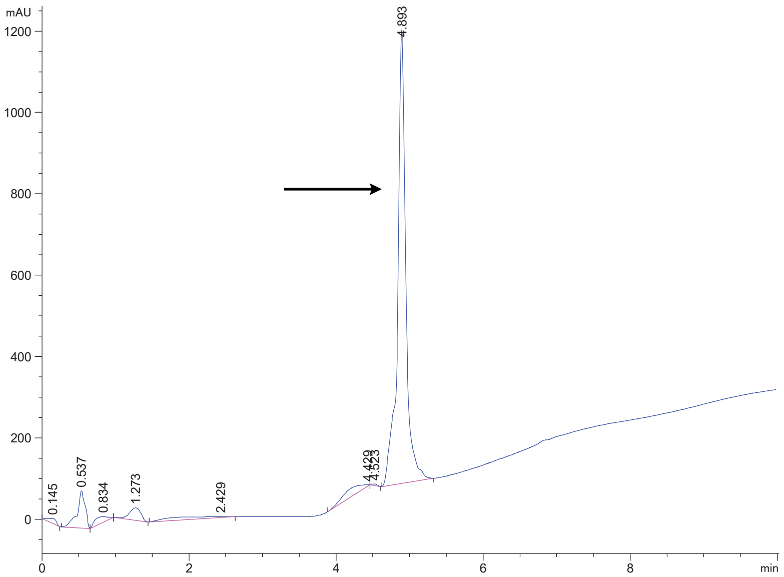

DPP4 10

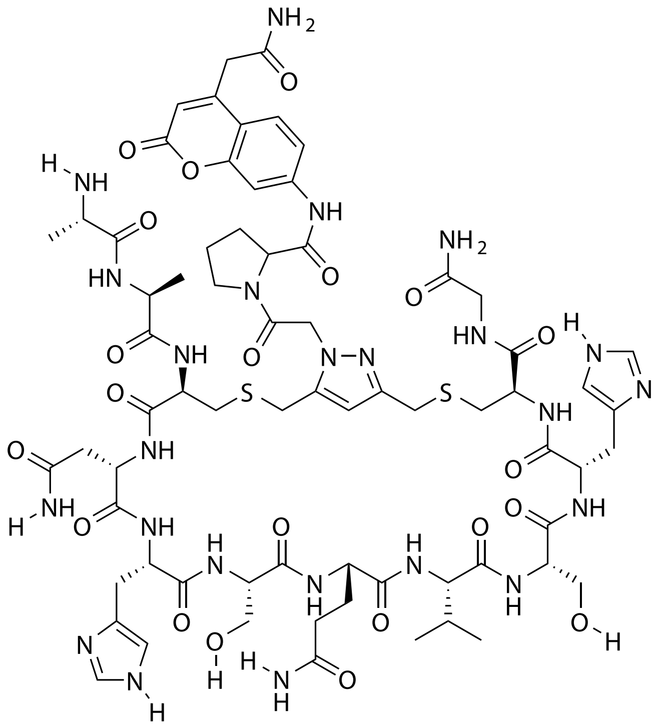

| **Probe Name** | **Sequence** | **Expected Mass $[M+1]^{+1}$ or $[M+2]^{+2}$** | **Found Mass** |
|---|---|---|---|
| DPP4 10 | AACNHSQVSHCG | 1659.6500 | 1659.9 |

Total Ion Chromatogram

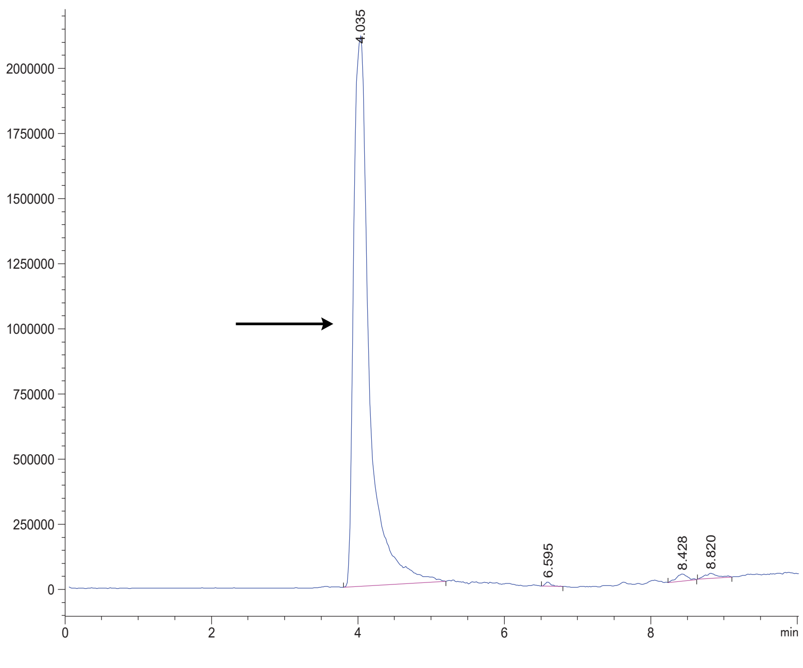


Variable Wavelength Detector: 220 nm

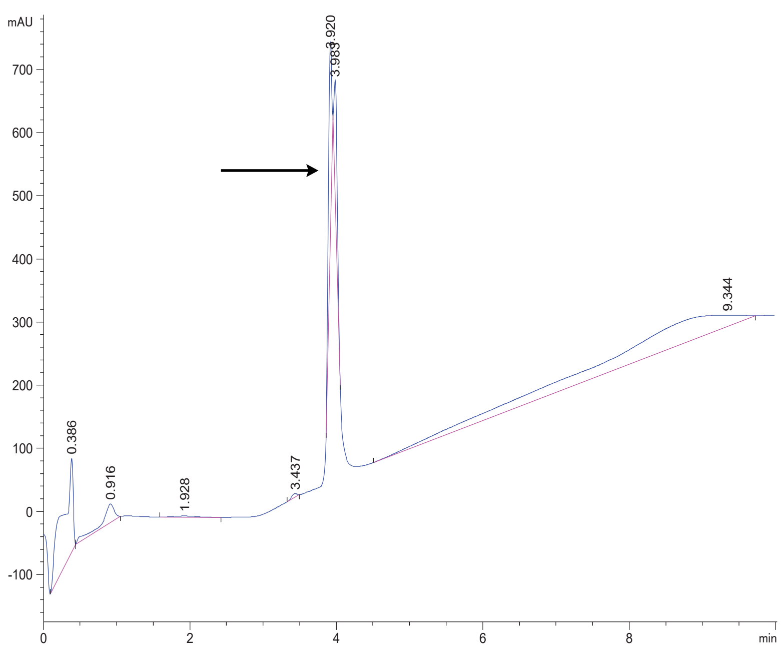

DPP4 10 AXs

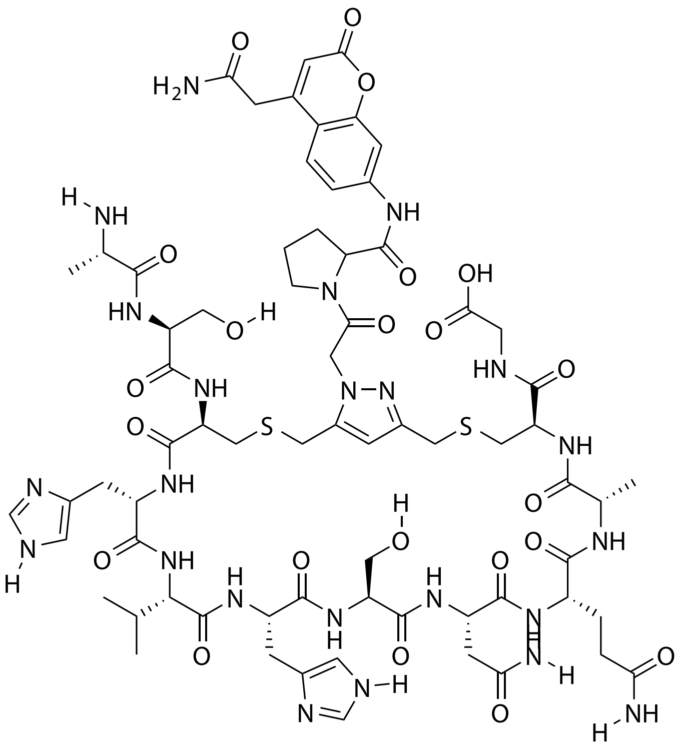

| Probe Name | Sequence | Expected Mass $[M+1]^{+1}$ or $[M+2]^{+2}$ | Found Mass |
| --- | --- | --- | --- |
| DPP4 10 AXs | ASCHVHSNQACG | 1659.6500 | 1660.3 |

Total Ion Chromatogram

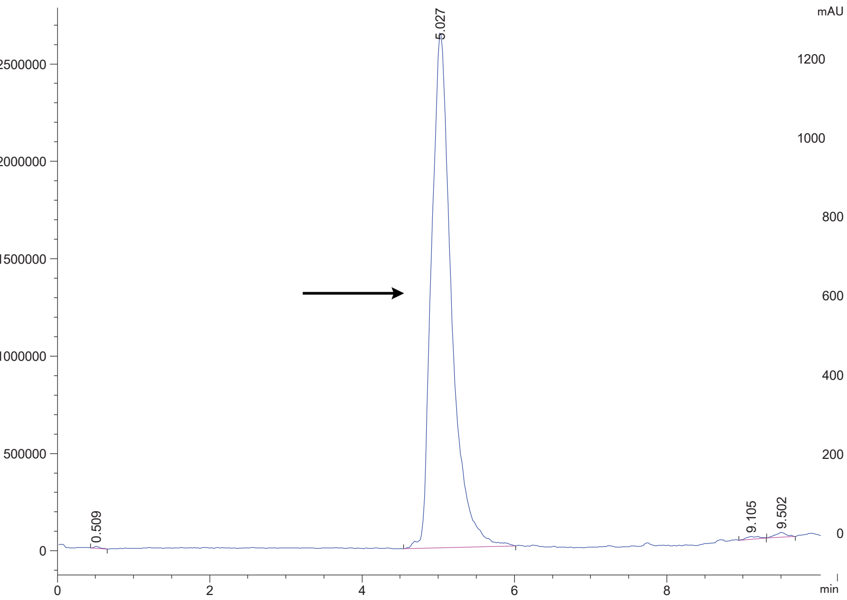


Variable Wavelength Detector: 220 nm

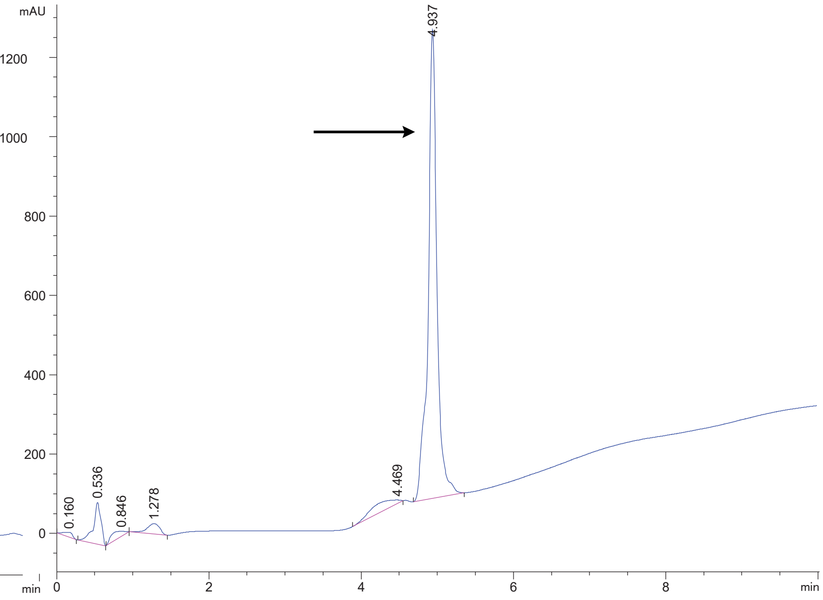

DPP4 10s

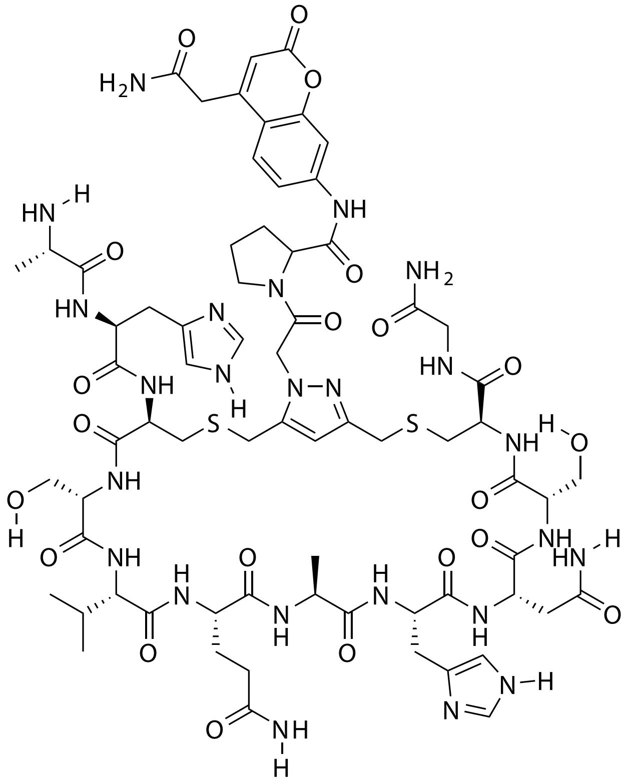

| Probe Name | Sequence | Expected Mass $[M+1]^{+1}$ or $[M+2]^{+2}$ | Found Mass |
|---|---|---|---|
| DPP4 10s | AHCSVQAHNSCG | 1659.6500 | 1659.9 |

Total Ion Chromatogram

Variable Wavelength Detector: 220 nm

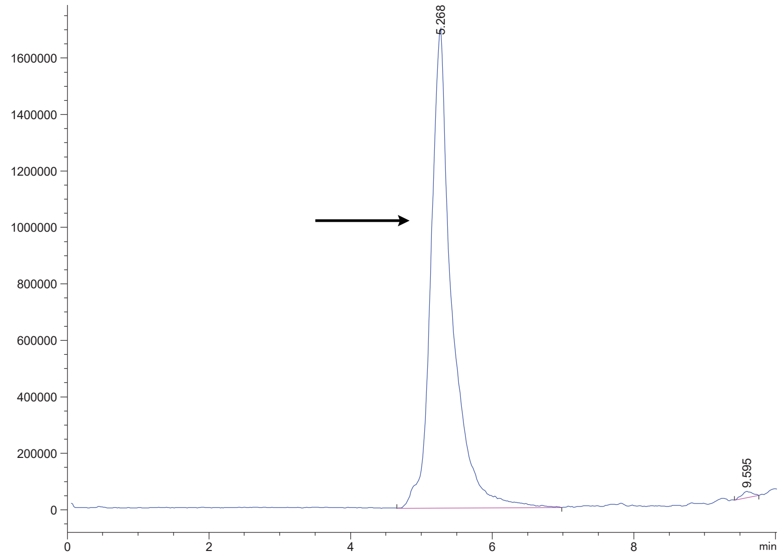


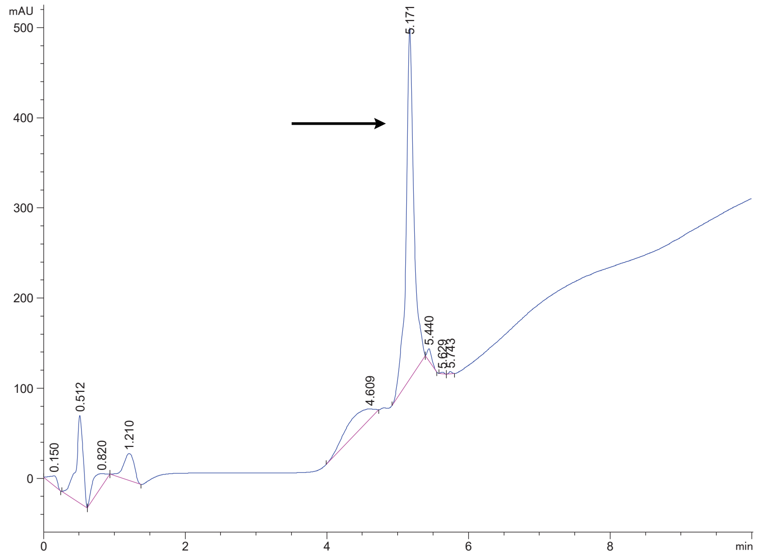

DPP4 10s.1

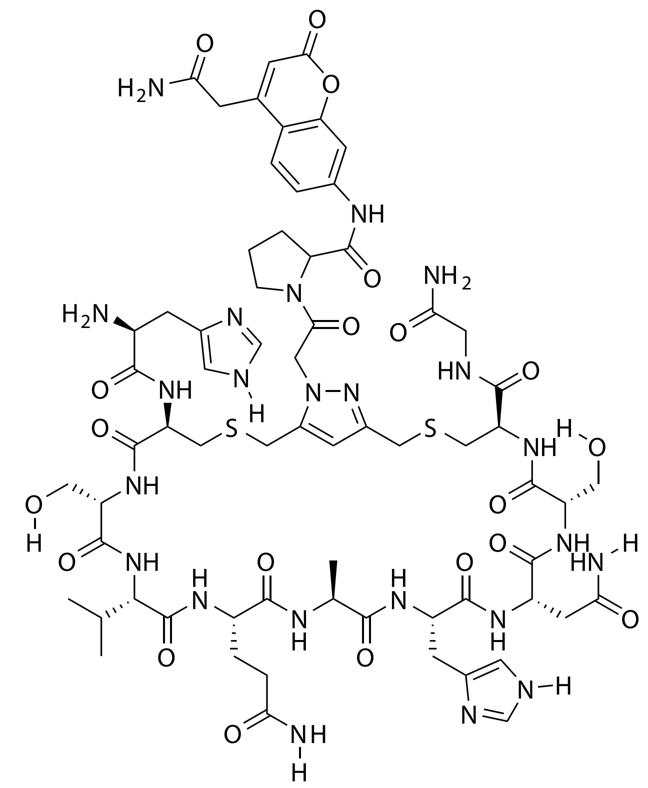

| Probe Name | Sequence | Expected Mass $[M+1]^{+1}$ or $[M+2]^{+2}$ | Found Mass |
|---|---|---|---|
| DPP4 10s.1 | HCSVQAHNSCG | 1588.6129 | 1588.8 |

Total Ion Chromatogram

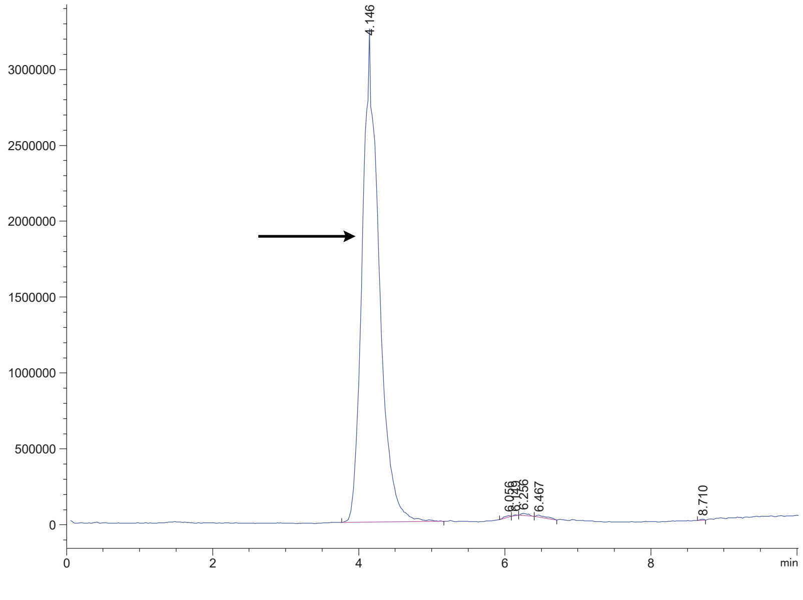

Variable Wavelength Detector: 220 nm

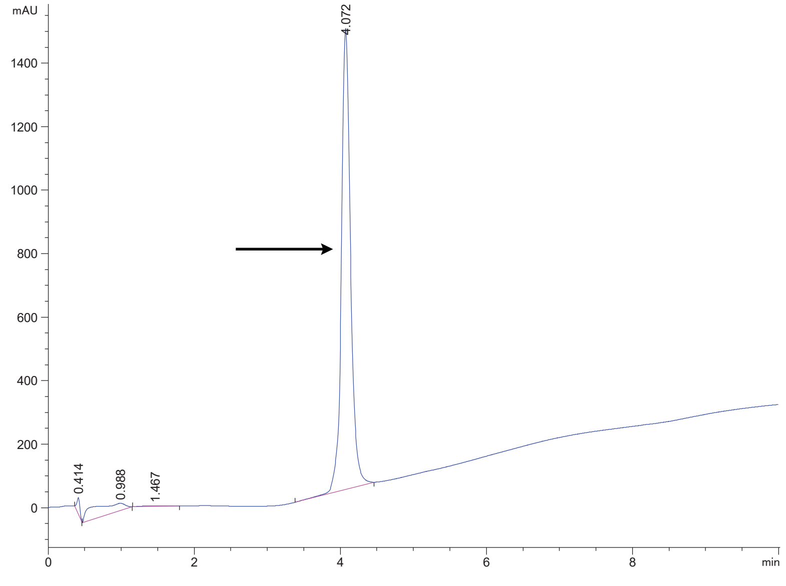

DPP4 10s.2

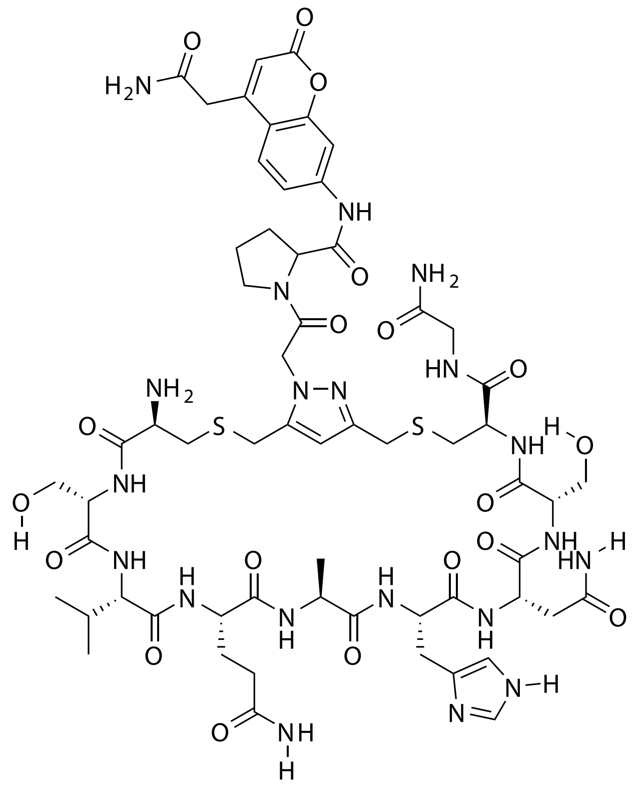

| Probe Name | Sequence | Expected Mass $[M+1]^{+1}$ or $[M+2]^{+2}$ | Found Mass |
|---|---|---|---|
| DPP4 10s.2 | CSVQAHNSCG | 1451.5540 | 1451.8 |

Total Ion Chromatogram

Variable Wavelength Detector: 220 nm

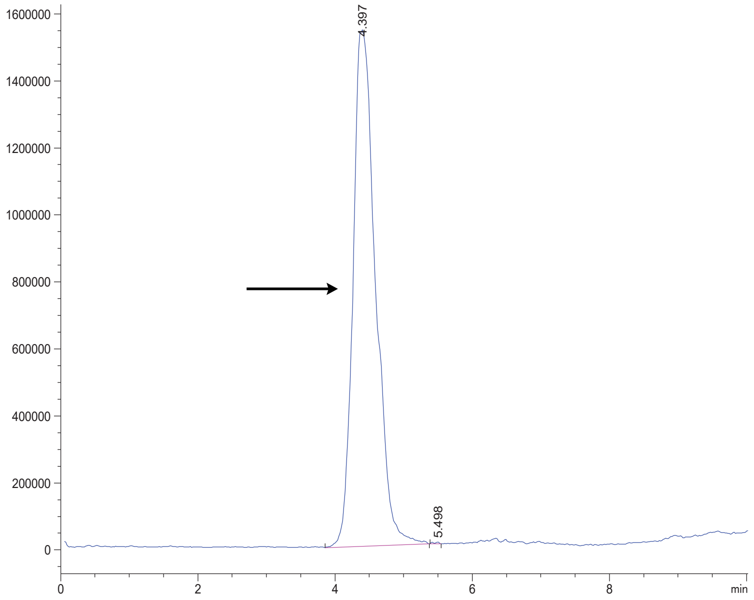


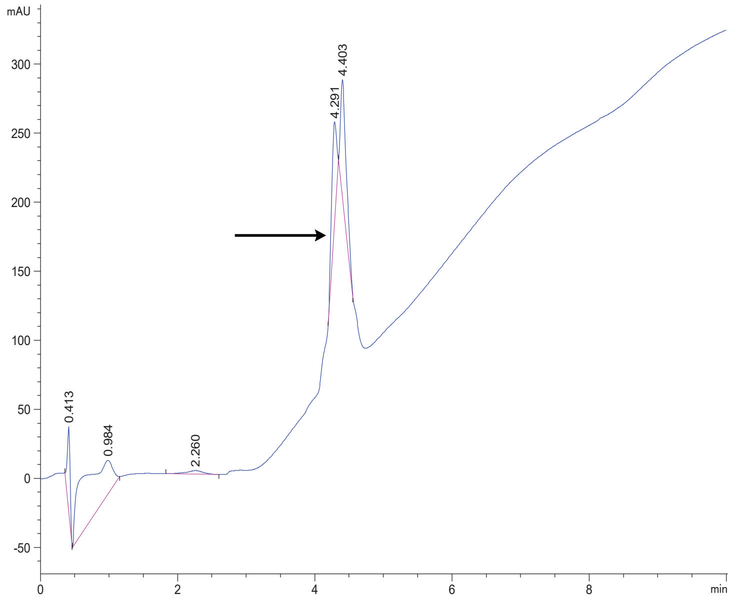

DPP4 10s.3

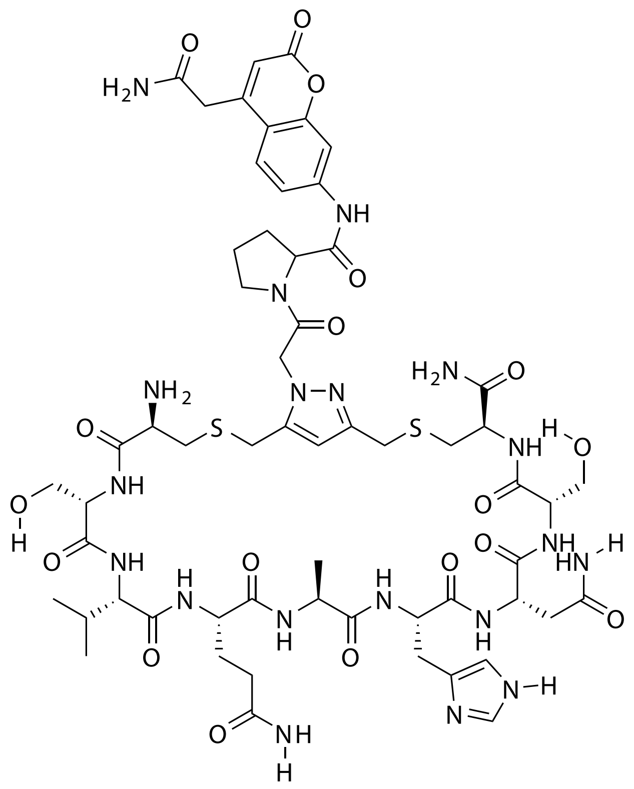

| Probe Name | Sequence | Expected Mass $[M+1]^{+1}$ or $[M+2]^{+2}$ | Found Mass |
|---|---|---|---|
| DPP4 10s.3 | CSVQAHNSC | 1394.5325 | 1394.9 |

Total Ion Chromatogram

Variable Wavelength Detector: 220 nm

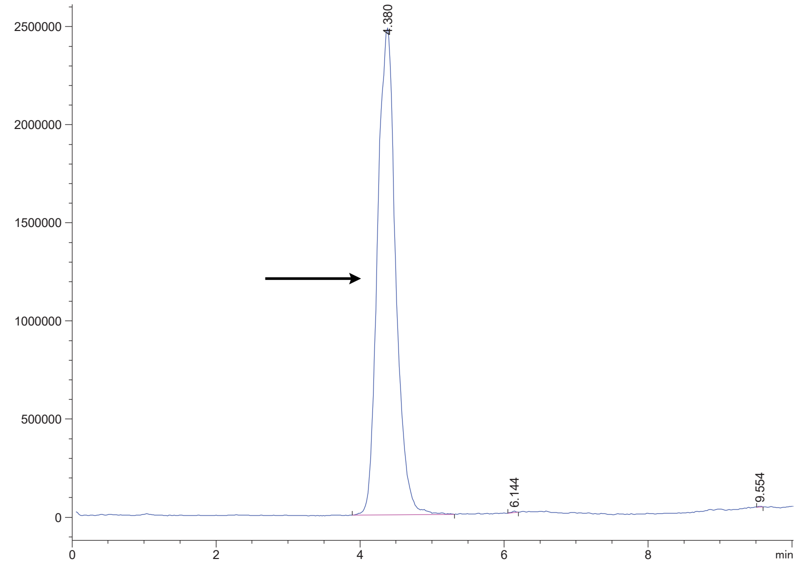


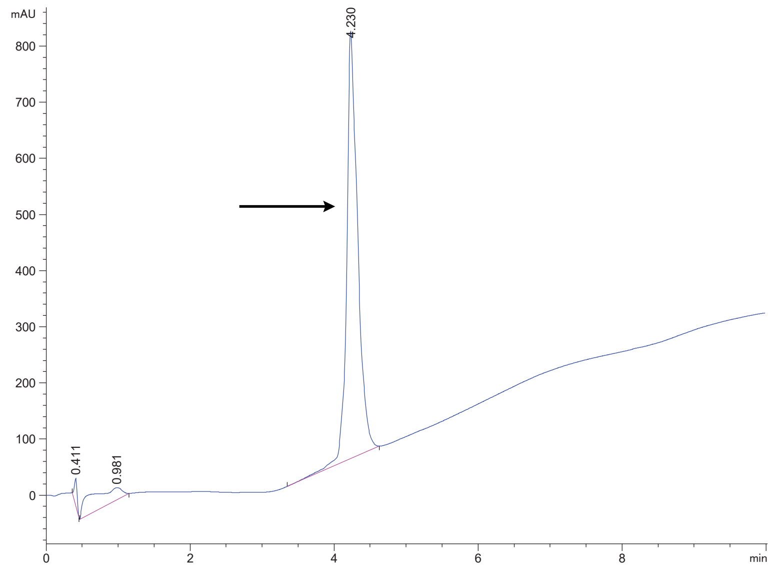

DPP4 10s.4

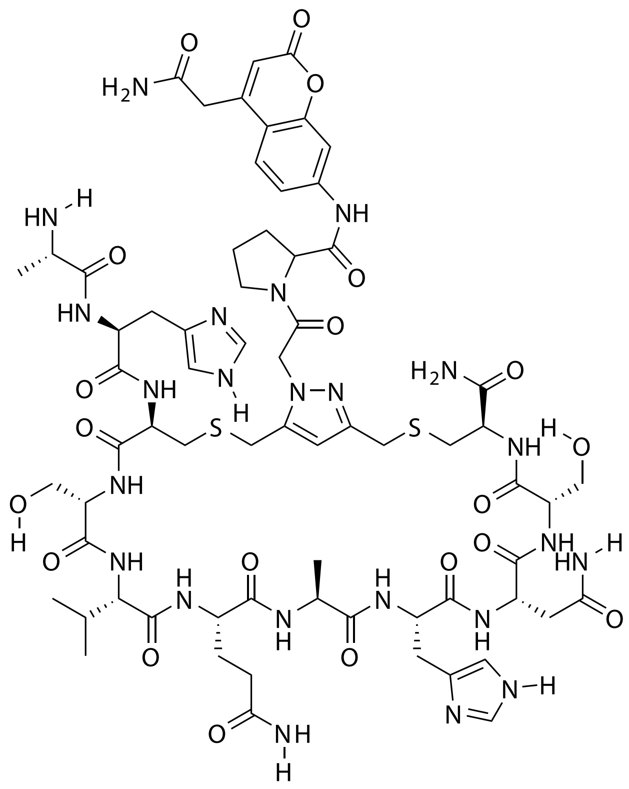

| Probe Name | Sequence | Expected Mass $[M+1]^{+1}$ or $[M+2]^{+2}$ | Found Mass |
|---|---|---|---|
| DPP4 10s.4 | AHCSVQAHNSC | 1602.6286 | 1602.9 |

Total Ion Chromatogram

Variable Wavelength Detector: 220 nm

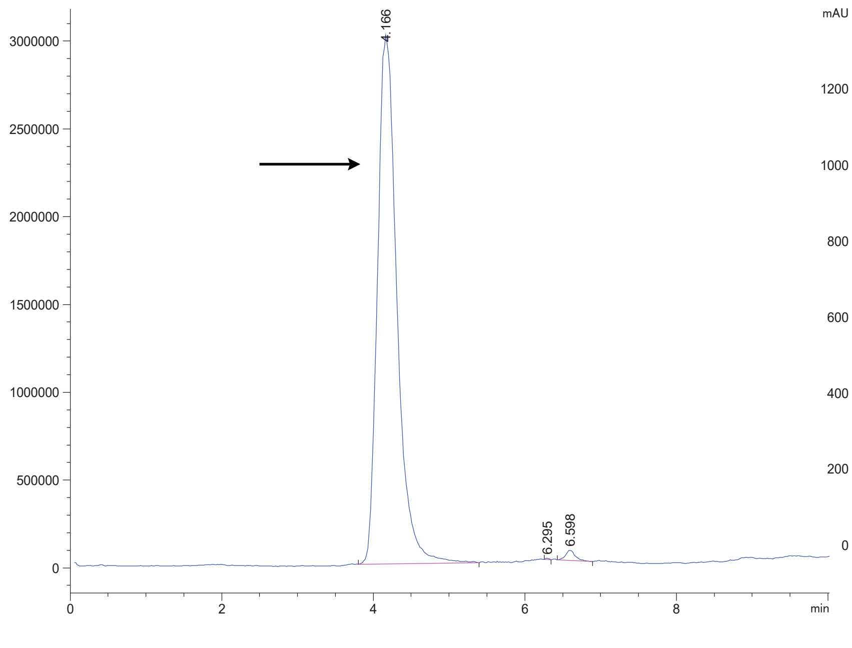


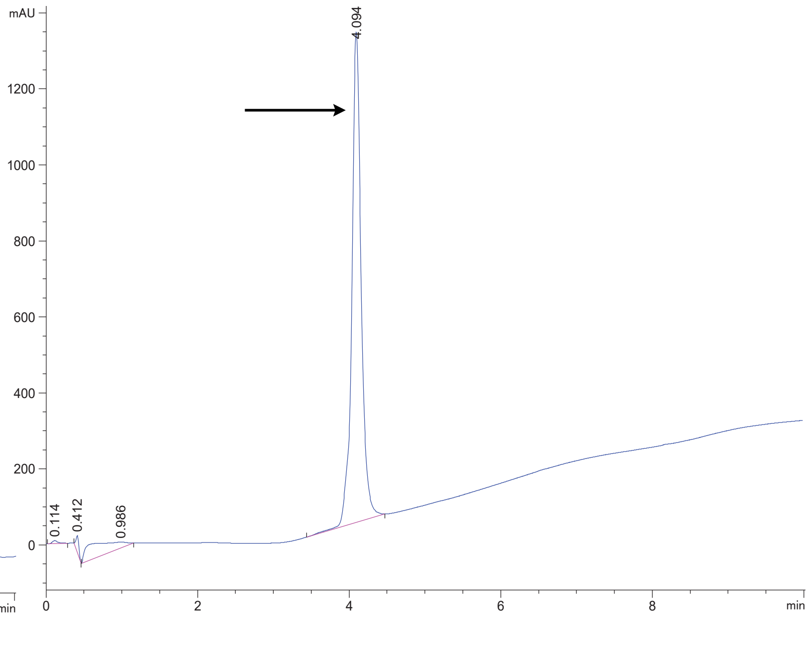

DPP4 10s.5

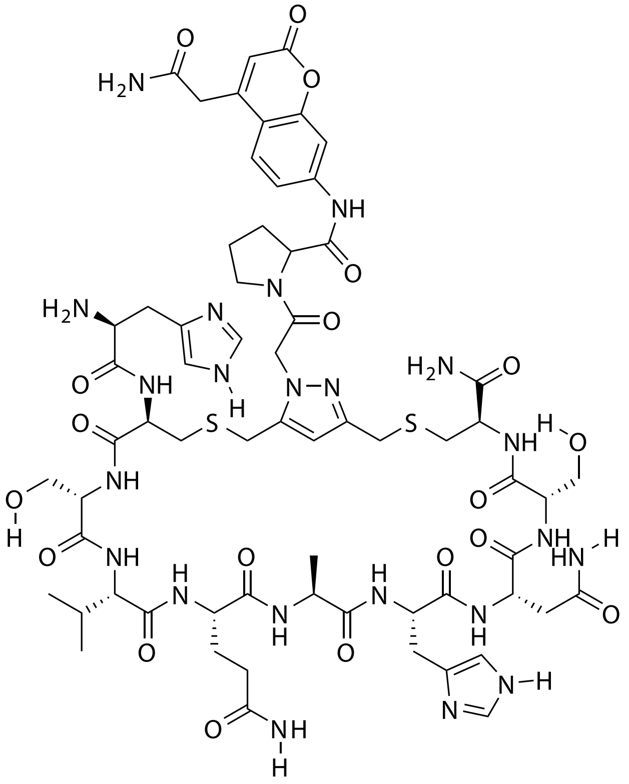

| Probe Name | Sequence | Expected Mass $[M+1]^{+1}$ or $[M+2]^{+2}$ | Found Mass |
|---|---|---|---|
| DPP4 10s.5 | HCSVQAHNSC | 1531.5915 | 1531.8 |

Total Ion Chromatogram

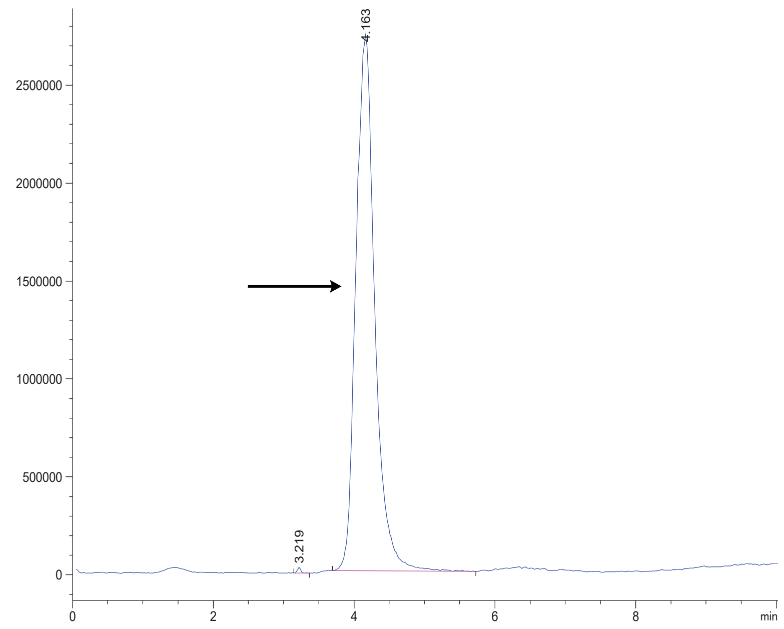


Variable Wavelength Detector: 220 nm

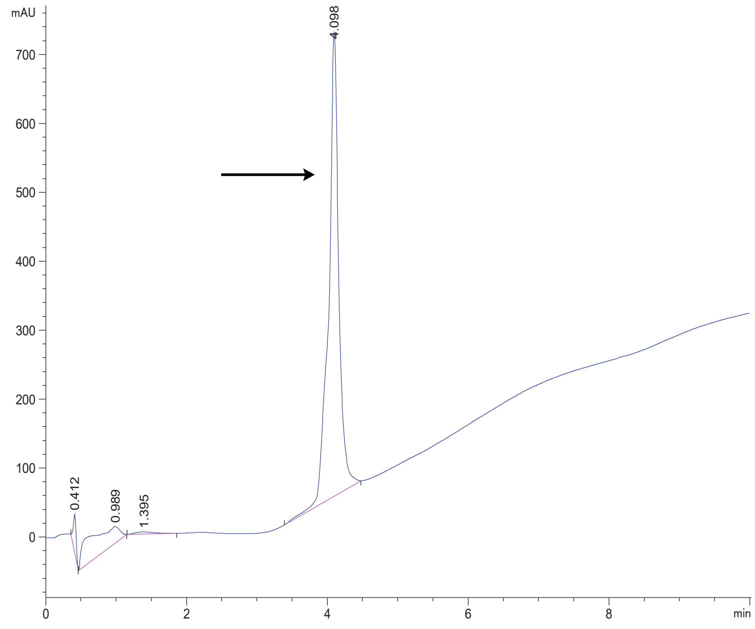

DPP4 11

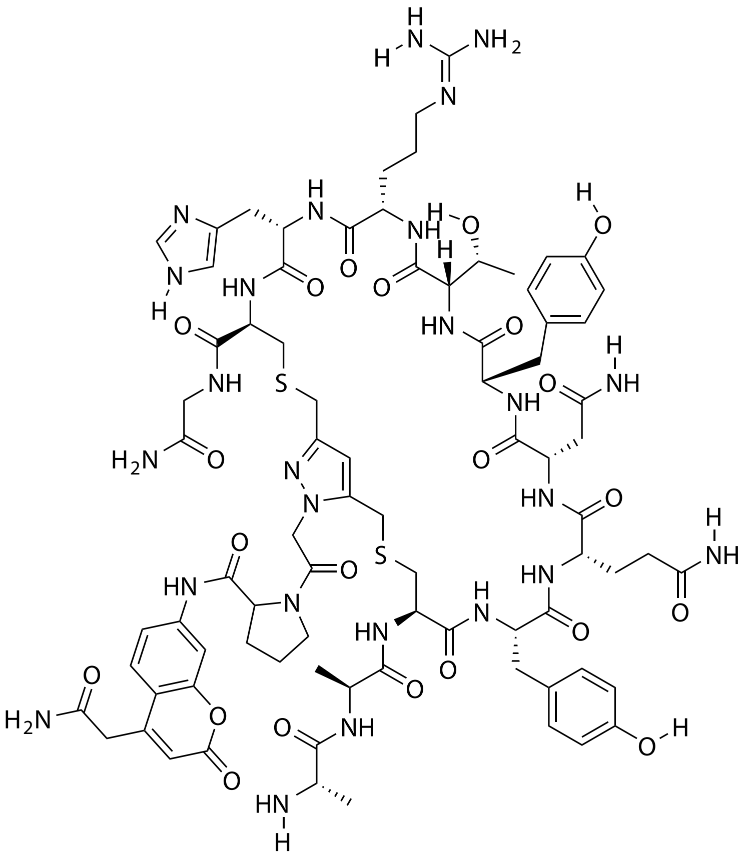

| Probe Name | Sequence | Expected Mass $[M+1]^{+1}$ or $[M+2]^{+2}$ | Found Mass |
|---|---|---|---|
| DPP4 11 | AACYQNYTRHCG | 1832.7341 $[M+2]^{+2}$ | 916.7 $[M+2]^{+2}$ |

Total Ion Chromatogram

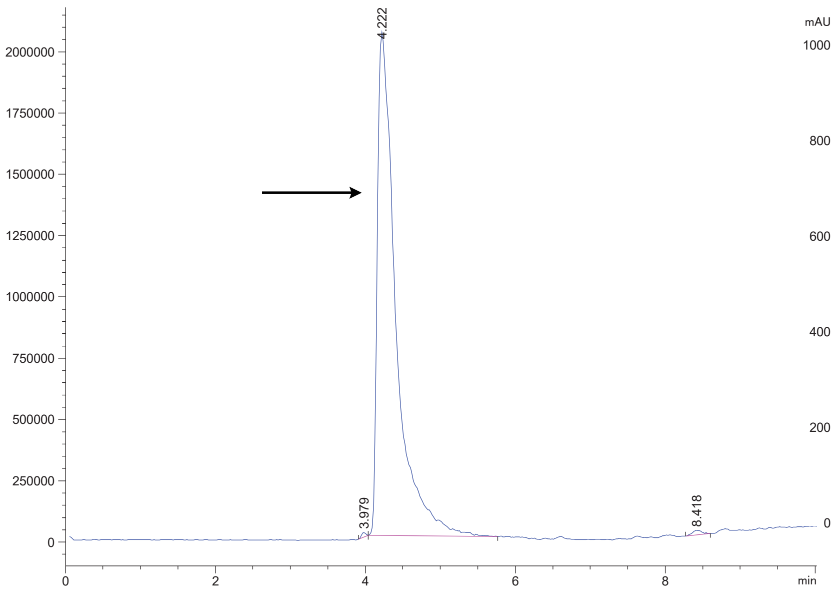


Variable Wavelength Detector: 220 nm

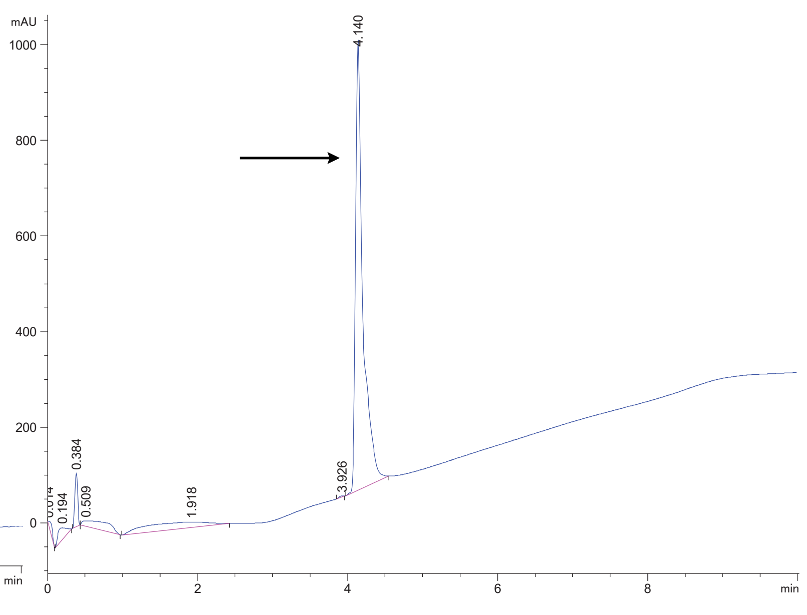

DPP4 13

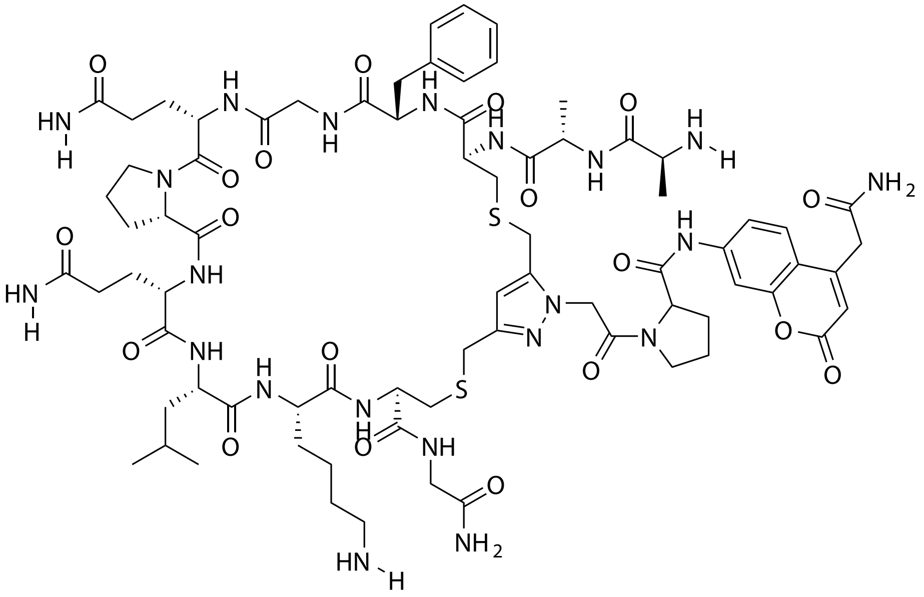

| Probe Name | Sequence | Expected Mass $[M+1]^{+1}$ or $[M+2]^{+2}$ | Found Mass |
|---|---|---|---|
| DPP4 13 | AACFGQPQLKCG | 1668.7371 | 1669.0 |

Total Ion Chromatogram

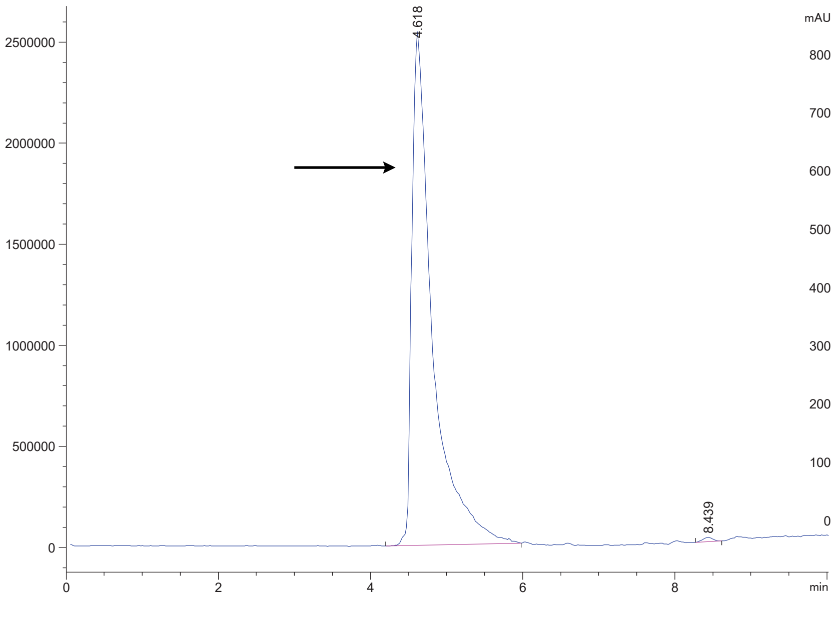

Variable Wavelength Detector: 220 nm

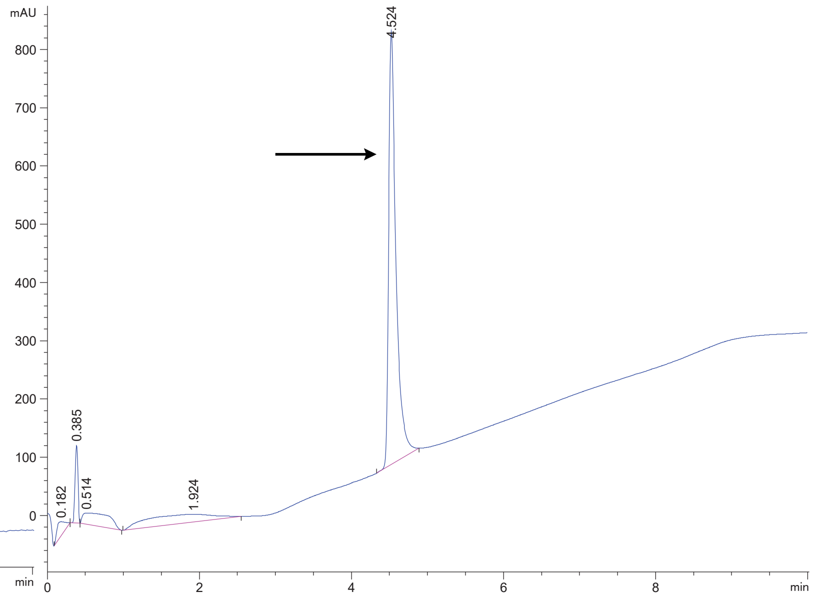

### DPP4 15

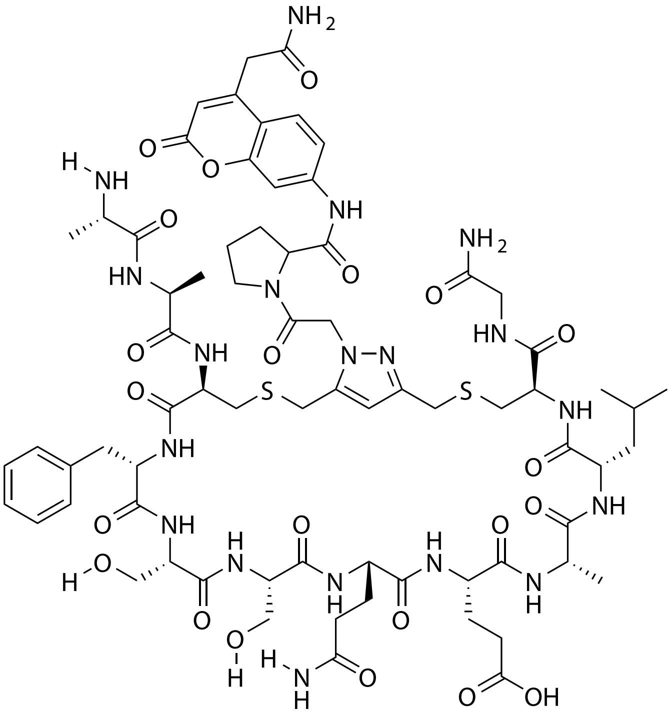

| Probe Name | Sequence | Expected Mass [M+1]$^{+1}$ or [M+2]$^{+2}$ | Found Mass |
|---|---|---|---|
| DPP4 15 | AACFSSQEALCG | 1632.6531 | 1632.9 |

Total Ion Chromatogram

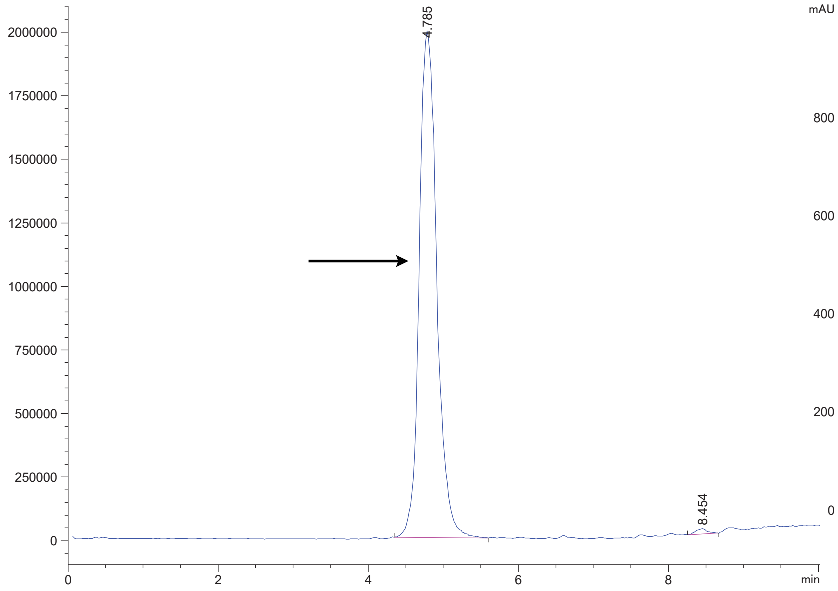


Variable Wavelength Detector: 220 nm

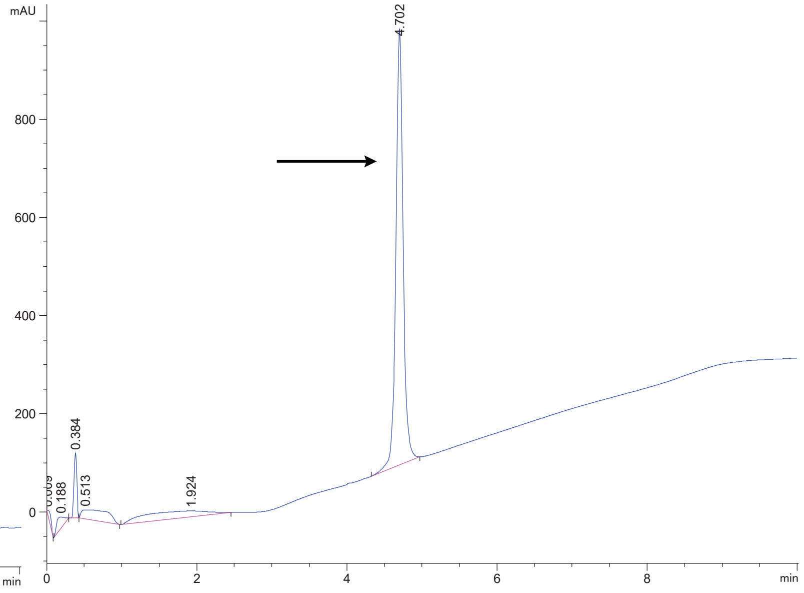

DPP4 18

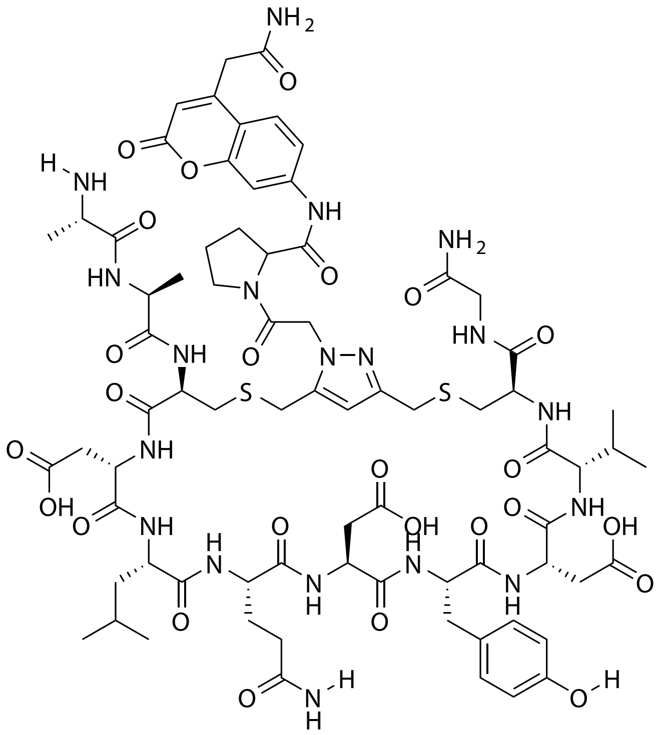

| Probe Name | Sequence | Expected Mass $[M+1]^{+1}$ or $[M+2]^{+2}$ | Found Mass |
|---|---|---|---|
| DPP4 18 | AACDLQDYDVCG | 1718.6534 | 1718.9 |

Total Ion Chromatogram

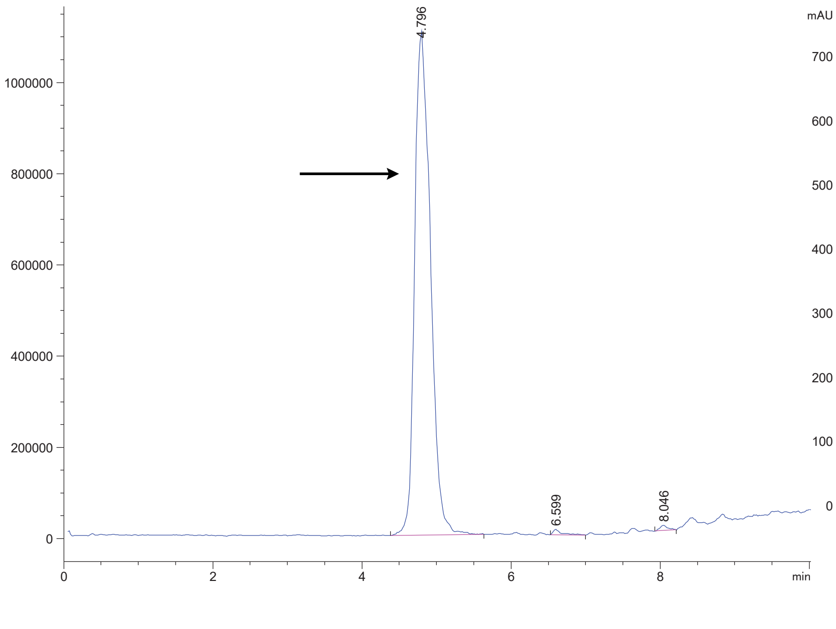

Variable Wavelength Detector: 220 nm

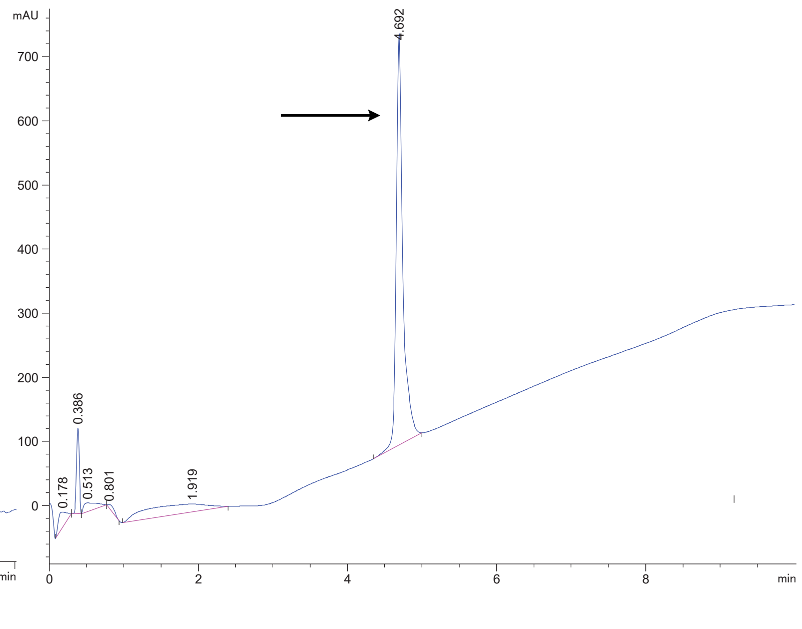

### DPP4 31

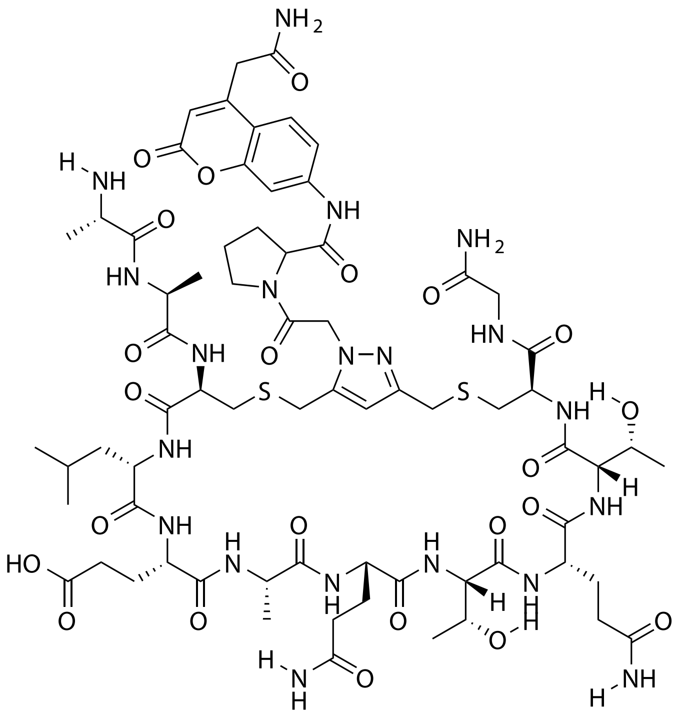

| **Probe Name** | **Sequence** | **Expected Mass $[M+1]^{+1}$ or $[M+2]^{+2}$** | **Found Mass** |
|---|---|---|---|
| DPP4 31 | AACLEAQTQTCG | 1641.6745 | 1642.0 |

Total Ion Chromatogram

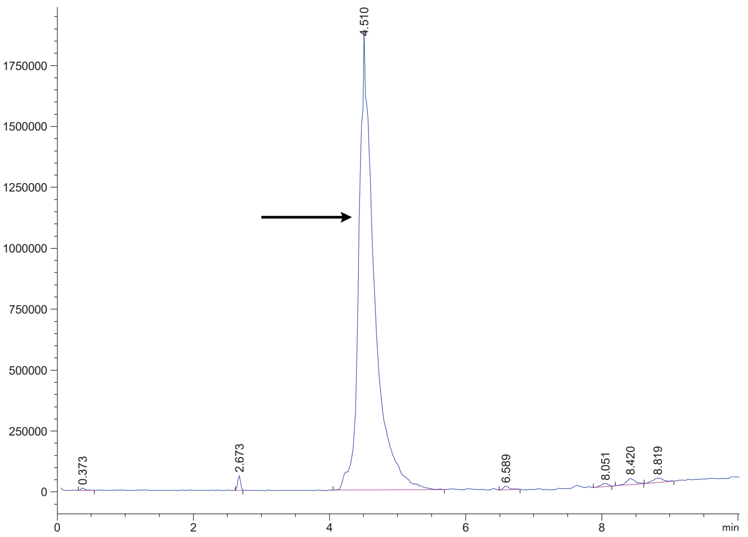


Variable Wavelength Detector: 220 nm

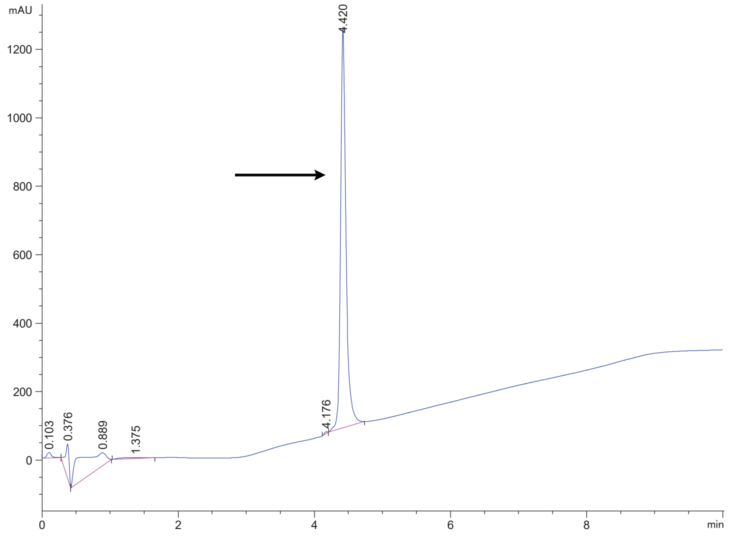

FAP 2

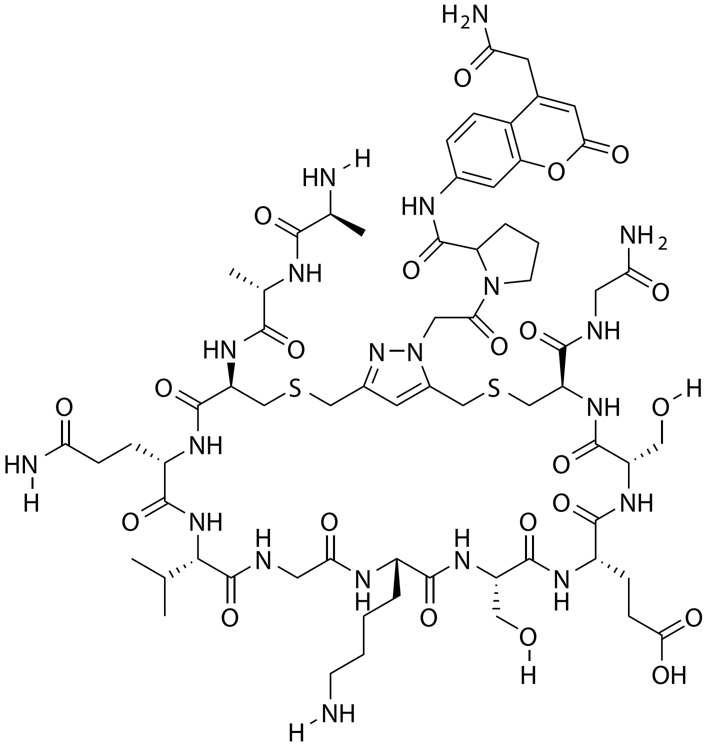

| Probe Name | Sequence | Expected Mass $[M+1]^{+1}$ or $[M+2]^{+2}$ | Found Mass |
|---|---|---|---|
| FAP 2 | AACQVGKSESCG | 1585.6483 | 1585.9 |

Total Ion Chromatogram

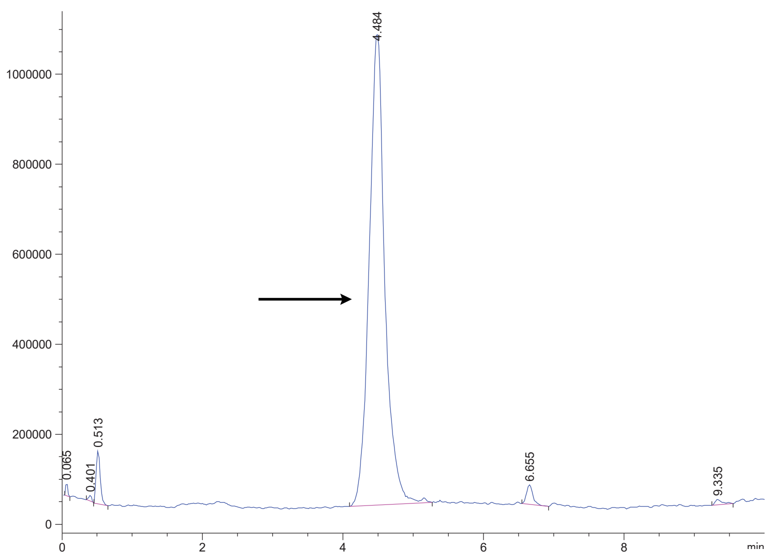

Variable Wavelength Detector: 220 nm

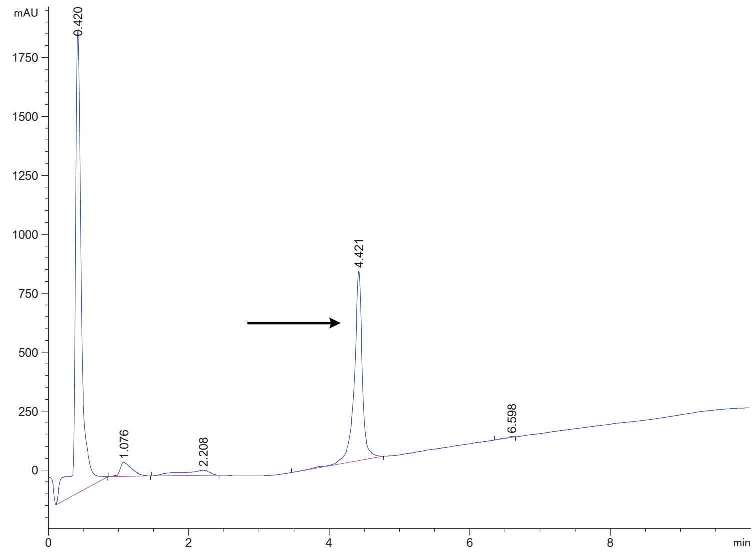

FAP 7

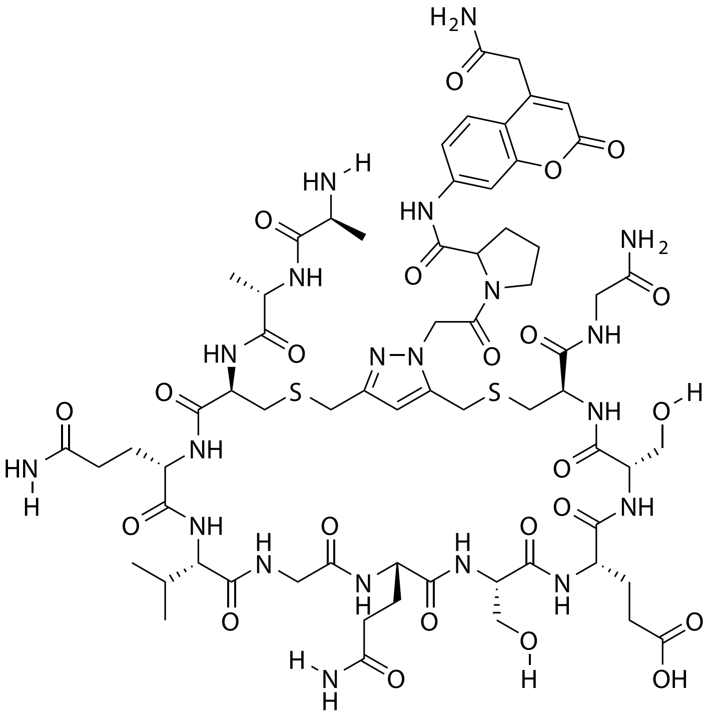

| Probe Name | Sequence | Expected Mass $[M+1]^{+1}$ or $[M+2]^{+2}$ | Found Mass |
|---|---|---|---|
| FAP 7 | AACQVGQSESCG | 1585.6119 | 1585.9 |

Total Ion Chromatogram

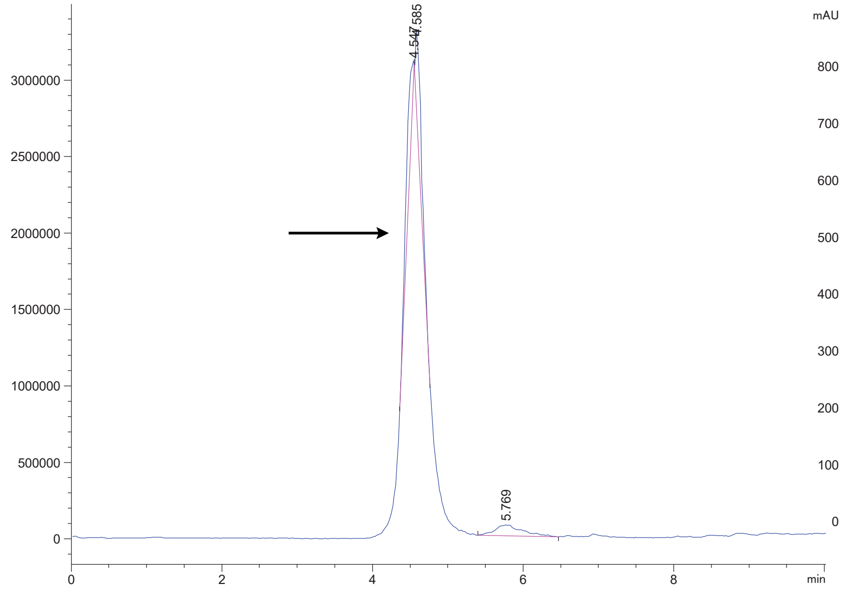


Variable Wavelength Detector: 220 nm

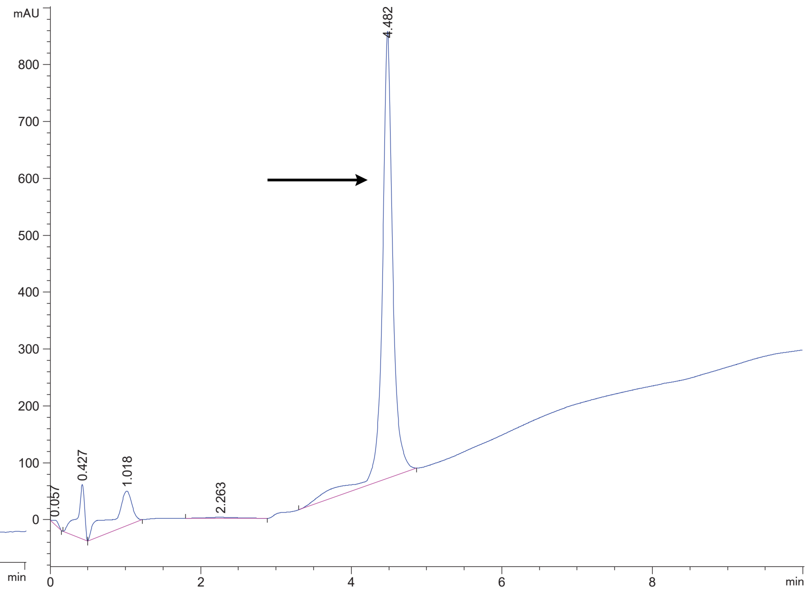

FAP 11

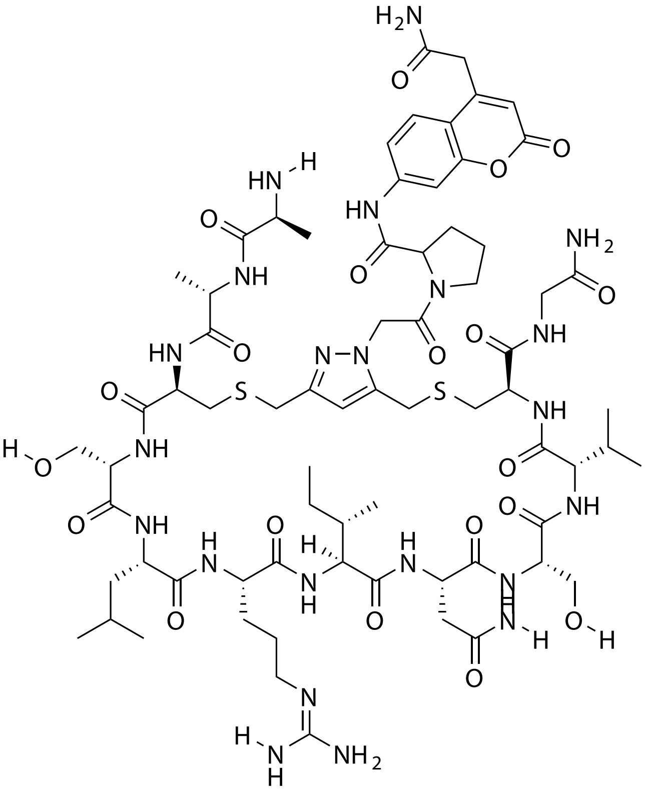

| Probe Name | Sequence | Expected Mass $[M+1]^{+1}$ or $[M+2]^{+2}$ | Found Mass |
|---|---|---|---|
| FAP 11 | AACSLRINSVCG | 1639.7429 | 1639.0 |

Total Ion Chromatogram

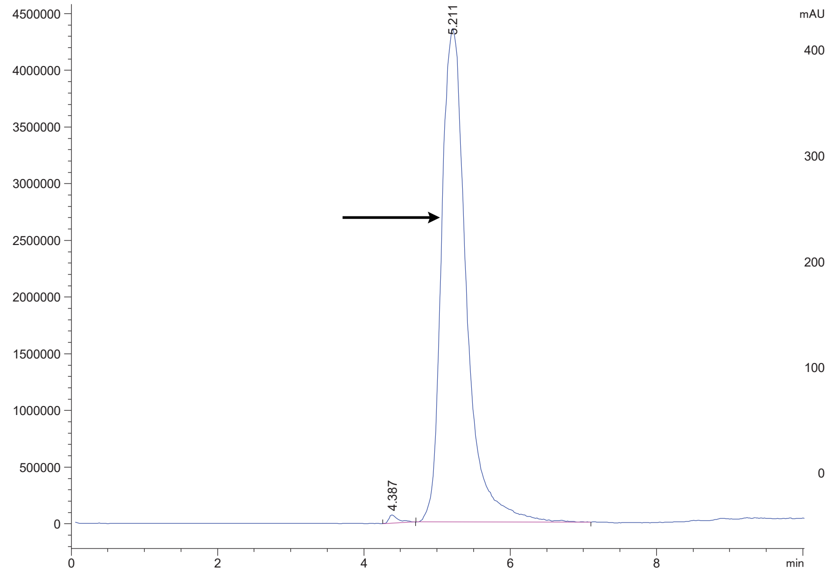


Variable Wavelength Detector: 220 nm

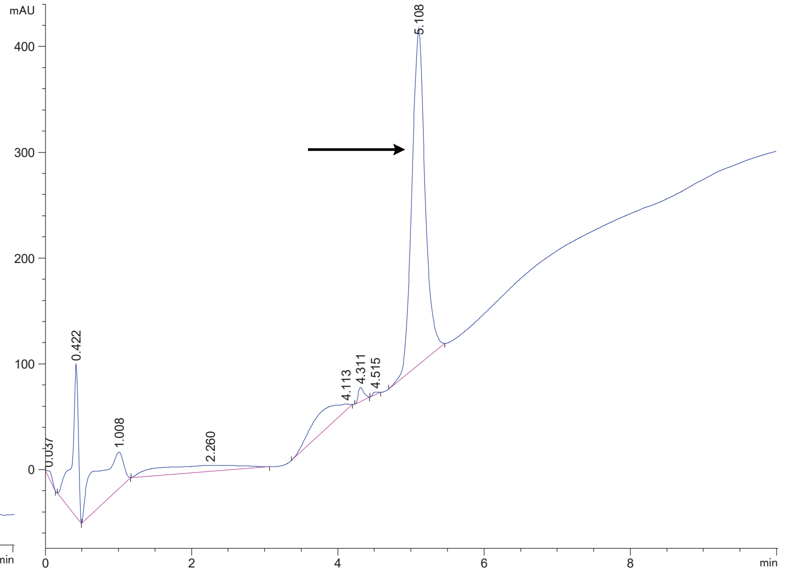

FAP 12

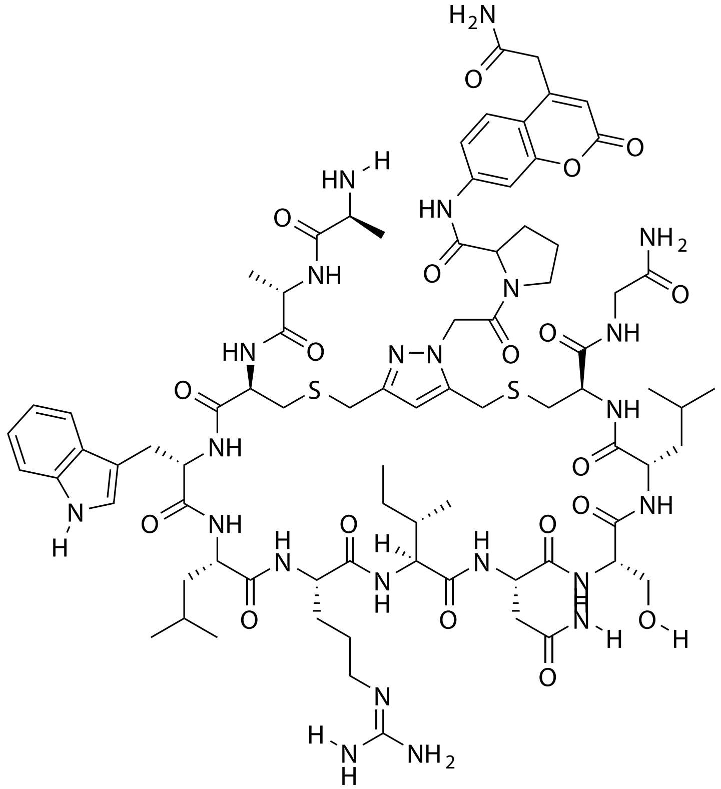

| Probe Name | Sequence | Expected Mass $[M+1]^{+1}$ or $[M+2]^{+2}$ | Found Mass |
|---|---|---|---|
| FAP 12 | AACWLRINSLCG | 876.9068 $[M+2]^{+2}$ | 876.4 $[M+2]^{+2}$ |

Total Ion Chromatogram

Variable Wavelength Detector: 220 nm

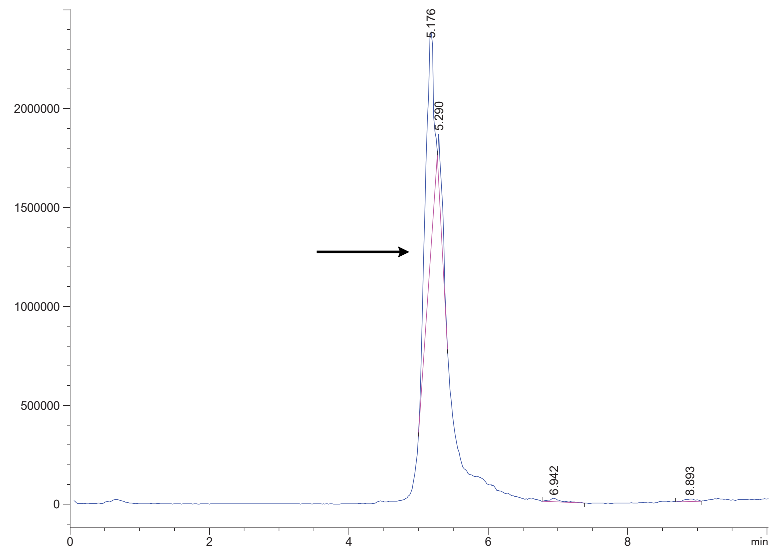


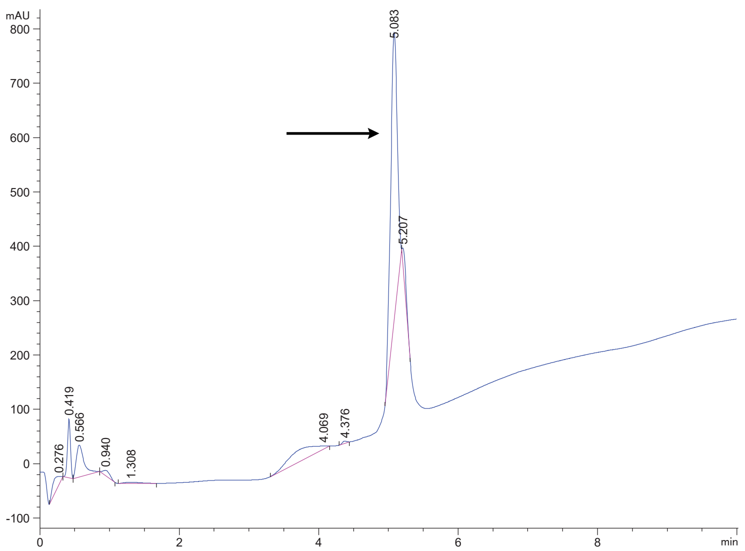

FAP 13

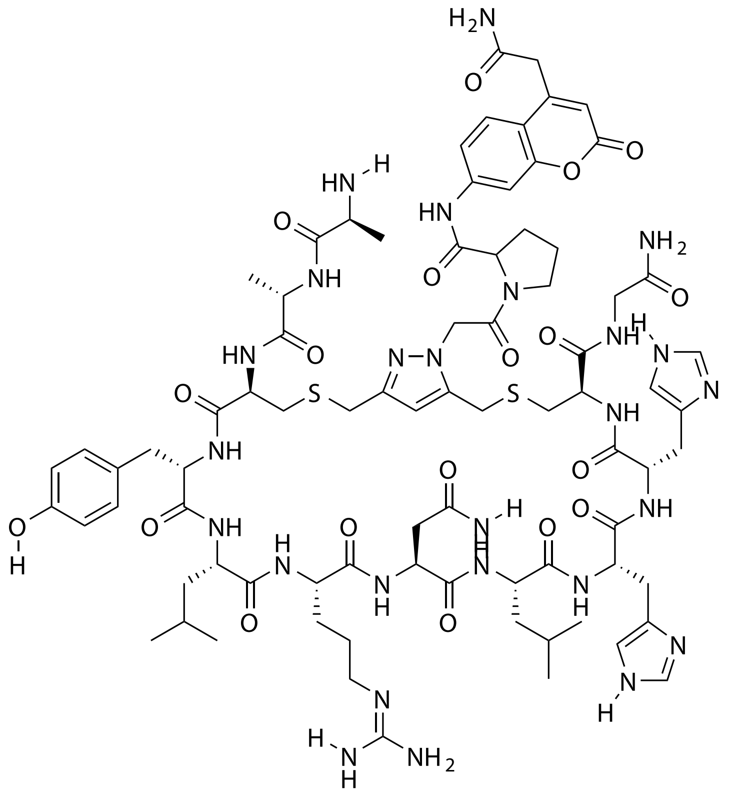

| Probe Name | Sequence | Expected Mass $[M+1]^{+1}$ or $[M+2]^{+2}$ | Found Mass |
|---|---|---|---|
| FAP 13 | AACYLRNLHHCG | 902.3997 $[M+2]^{+2}$ | 902.2 $[M+2]^{+2}$ |

Total Ion Chromatogram

Variable Wavelength Detector: 220 nm

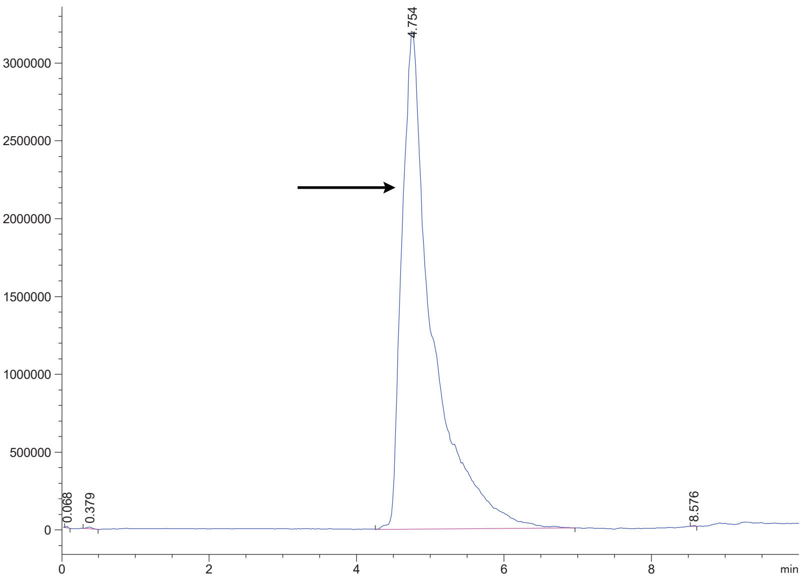

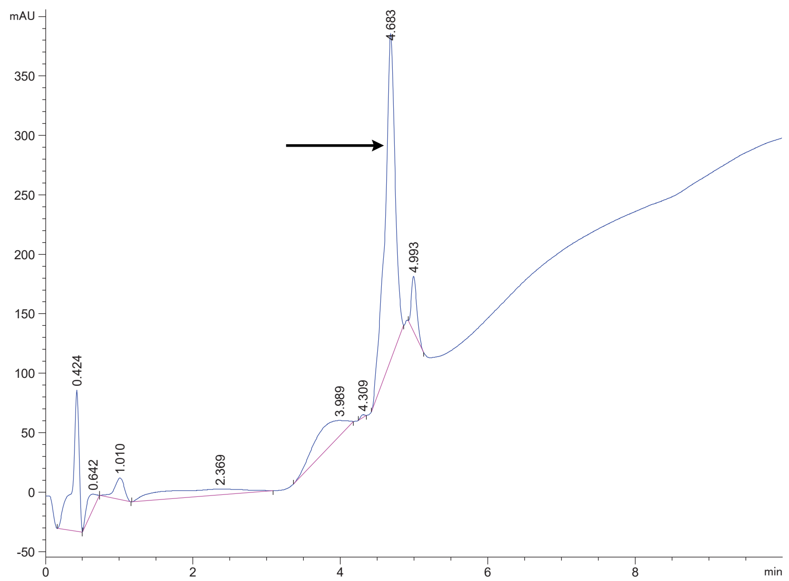

FAP 15

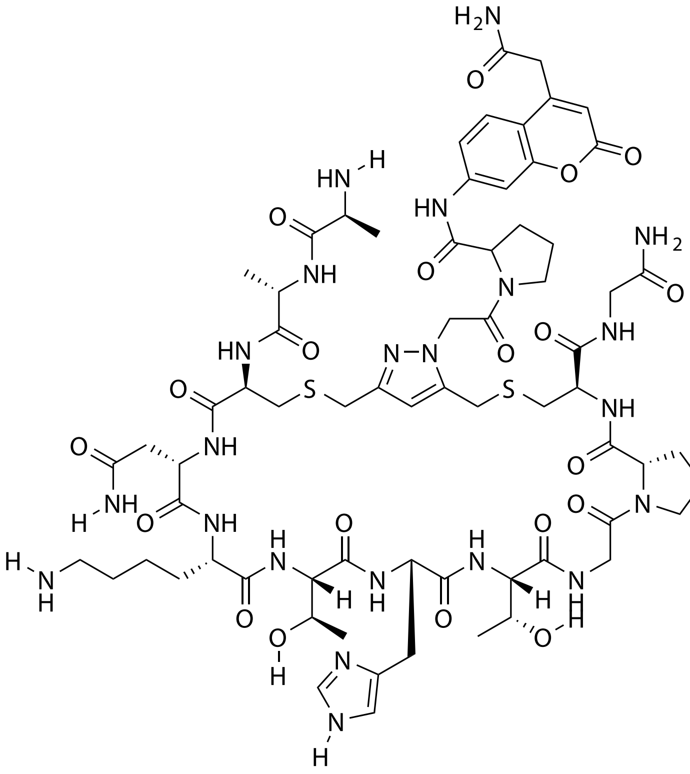

| Probe Name | Sequence | Expected Mass $[M+1]^{+1}$ or $[M+2]^{+2}$ | Found Mass |
|---|---|---|---|
| FAP 15 | AACNKTHTGPCG | 1605.6646 | 1604.9 |

Total Ion Chromatogram

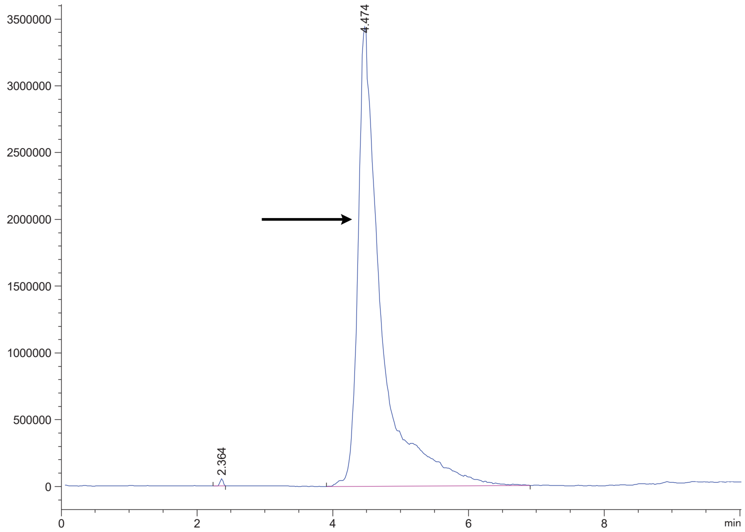


Variable Wavelength Detector: 220 nm

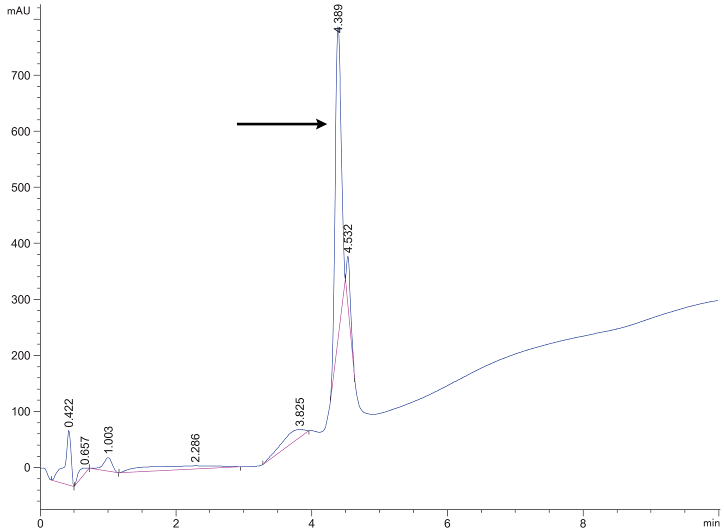

FAP 18

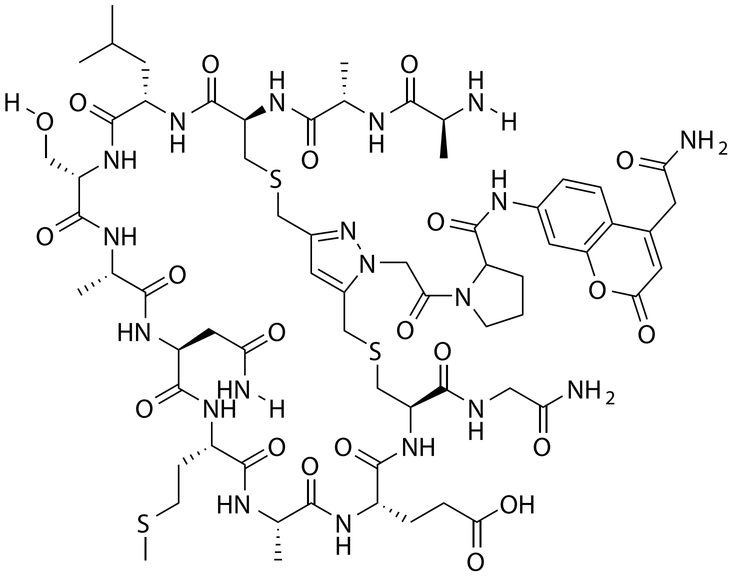

| Probe Name | Sequence | Expected Mass $[M+1]^{+1}$ or $[M+2]^{+2}$ | Found Mass |
|---|---|---|---|
| FAP 18 | AACLSANMAECG | 1586.6146 | 1586.9 |

Total Ion Chromatogram

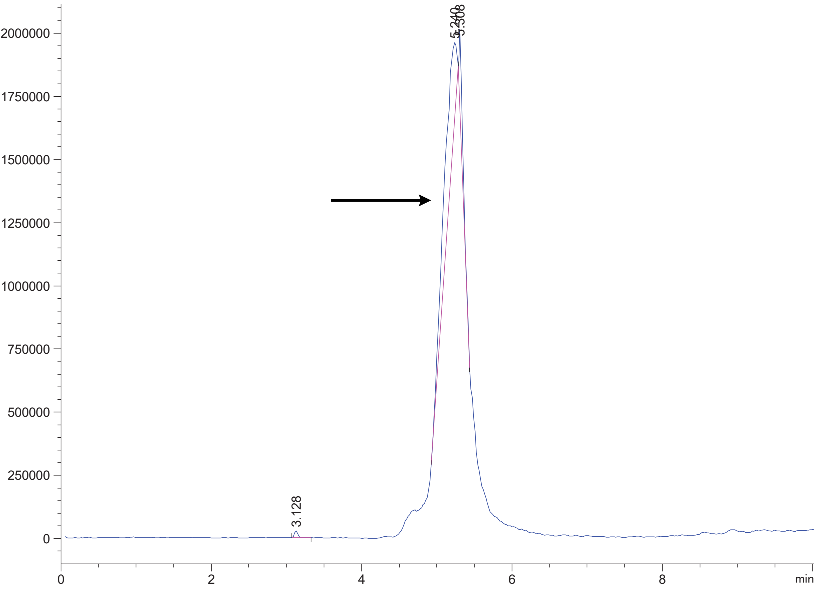


Variable Wavelength Detector: 220 nm

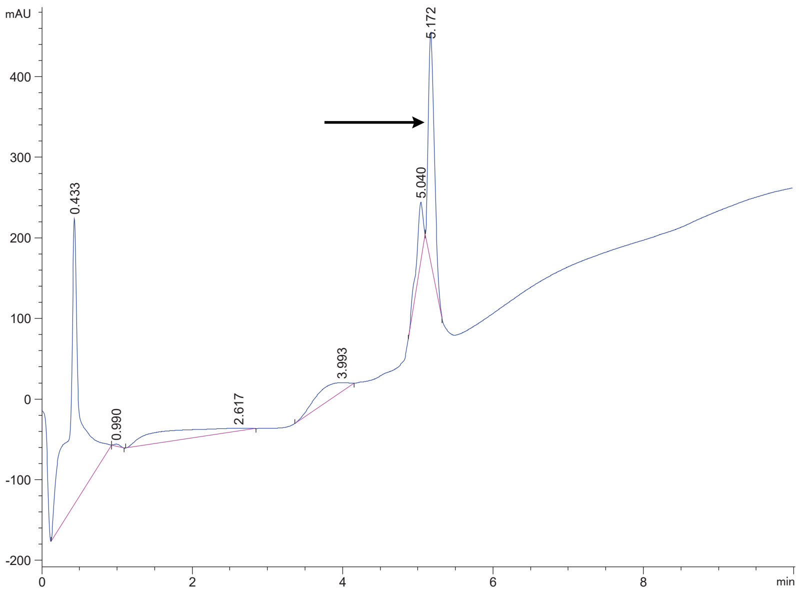

FAP 20

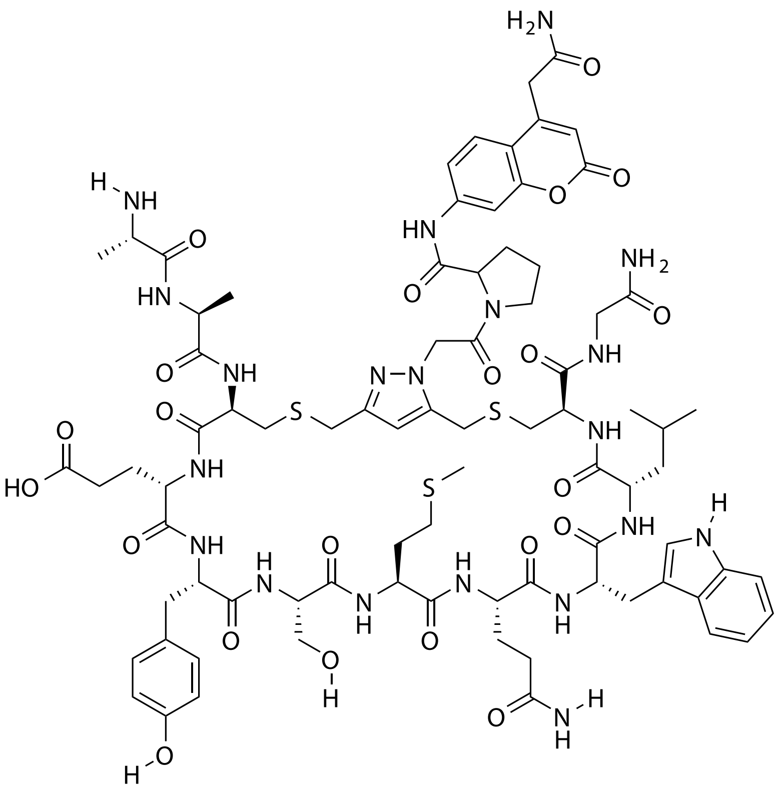

| Probe Name | Sequence | Expected Mass $[M+1]^{+1}$ or $[M+2]^{+2}$ | Found Mass |
|---|---|---|---|
| FAP 20 | AACEYSMQWLCG | 904.3533 $[M+2]^{+2}$ | 904.6 $[M+2]^{+2}$ |

Total Ion Chromatogram

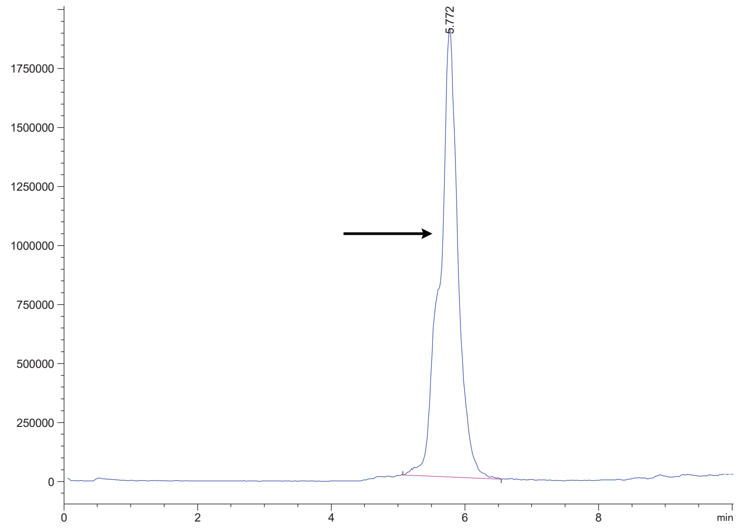


Variable Wavelength Detector: 220 nm

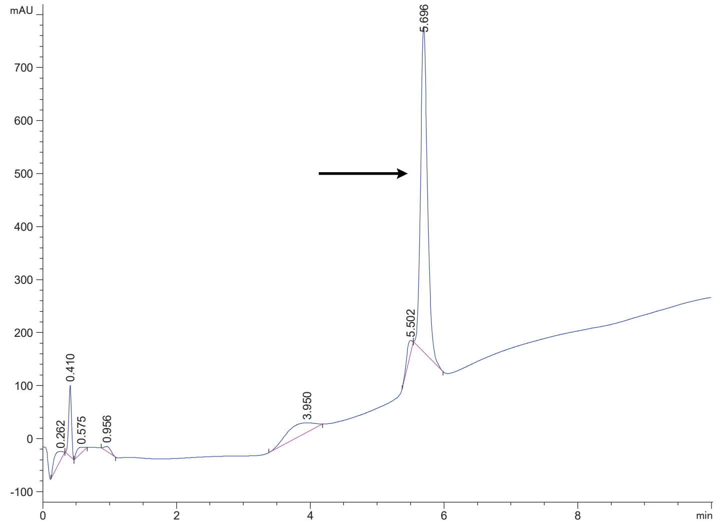

FAP 21

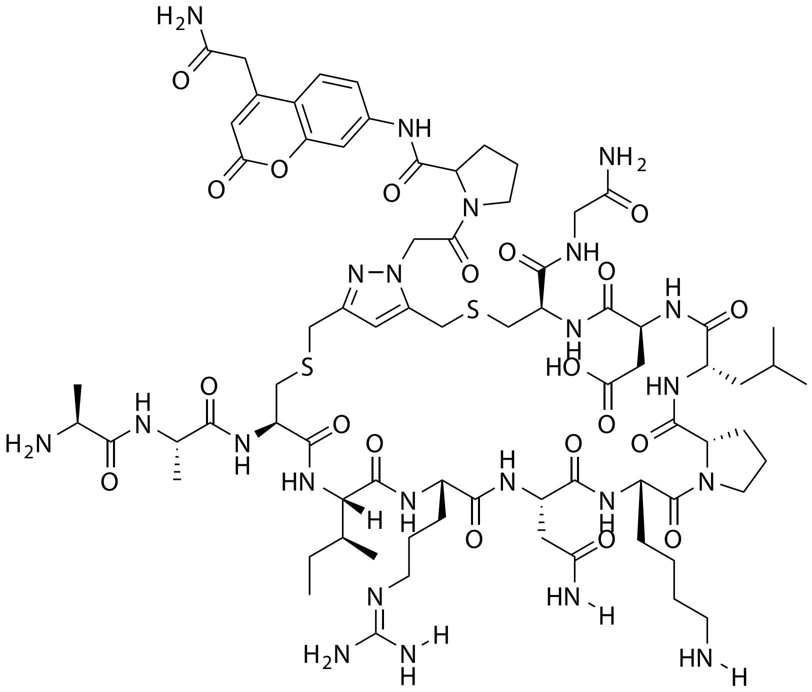


| Probe Name | Sequence | Expected Mass $[M+1]^{+1}$ or $[M+2]^{+2}$ | Found Mass |
|---|---|---|---|
| FAP 21 | AACIRNKPLDCG | 853.8965 $[M+2]^{+2}$ | 853.8 $[M+2]^{+2}$ |

Total Ion Chromatogram

Variable Wavelength Detector: 220 nm

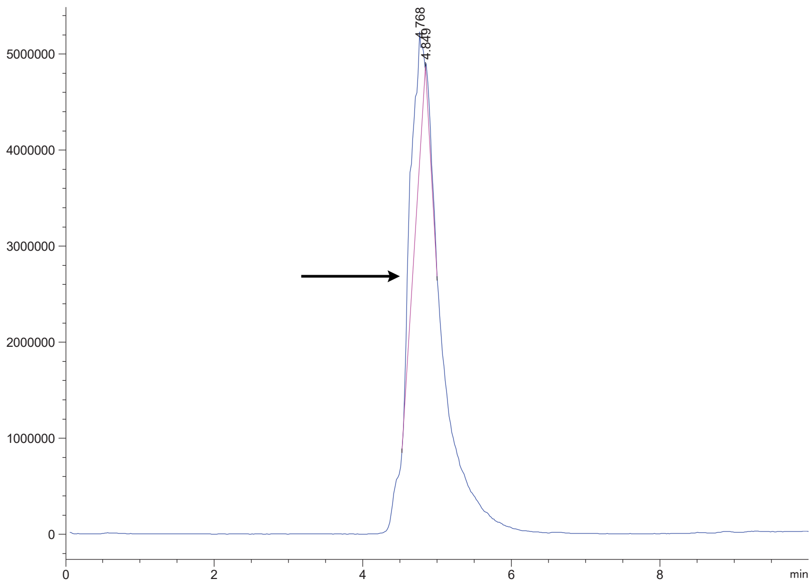


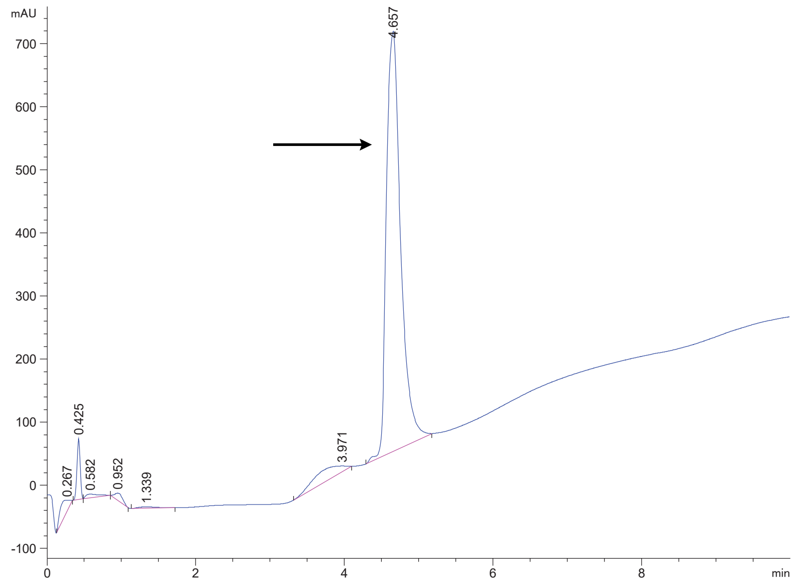

FAP 24

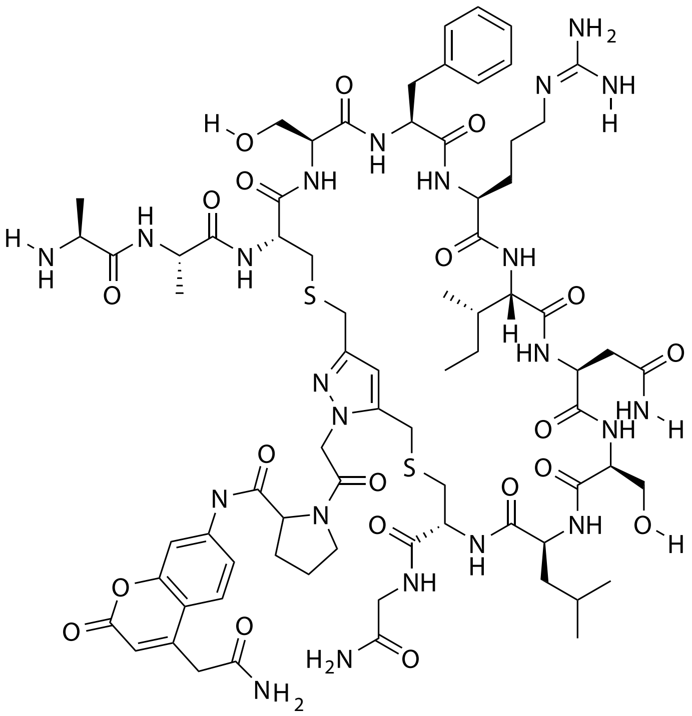

| Probe Name | Sequence | Expected Mass $[M+1]^{+1}$ or $[M+2]^{+2}$ | Found Mass |
|---|---|---|---|
| FAP 24 | AACSFRINSLCG | 844.3753 $[M+2]^{+2}$ | 844.2 $[M+2]^{+2}$ |

Total Ion Chromatogram

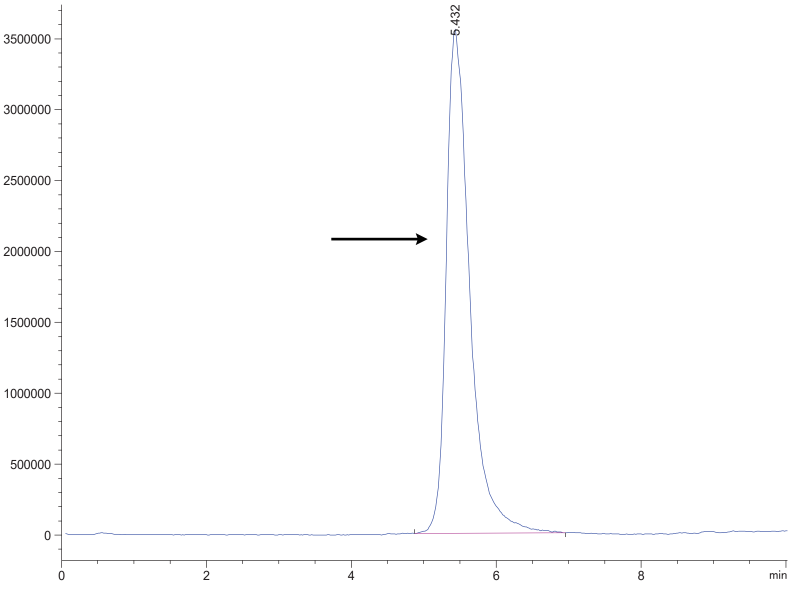


Variable Wavelength Detector: 220 nm

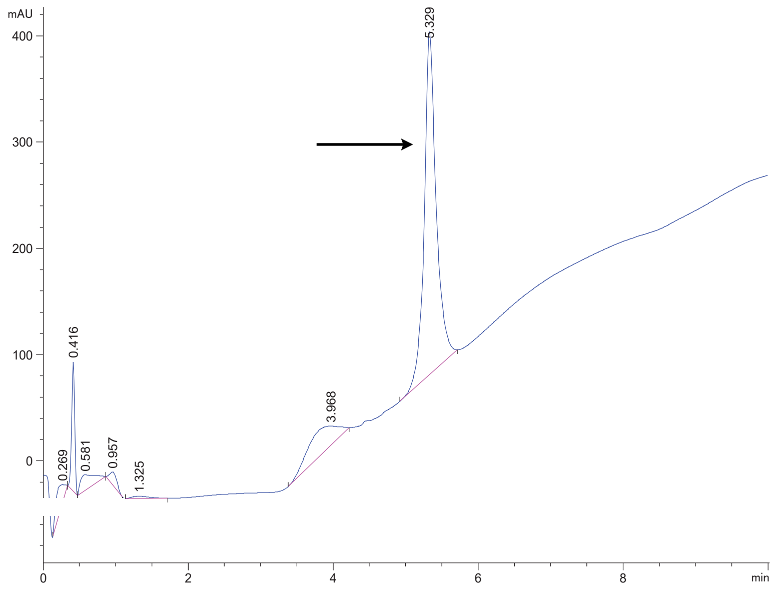

FAP 24.1

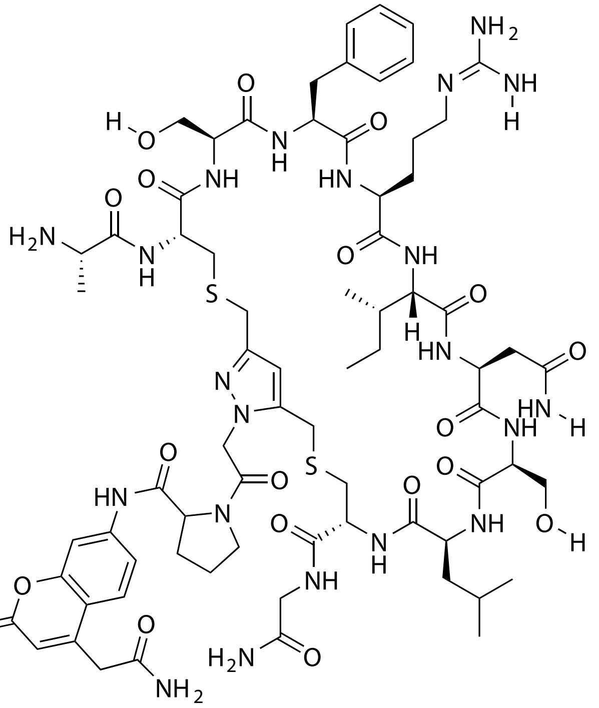

| Probe Name | Sequence | Expected Mass $[M+1]^{+1}$ or $[M+2]^{+2}$ | Found Mass |
|---|---|---|---|
| FAP 24.1 | ACSFRINSLCG | 1616.7058 | 1617.0 |

Total Ion Chromatogram

Variable Wavelength Detector: 220 nm

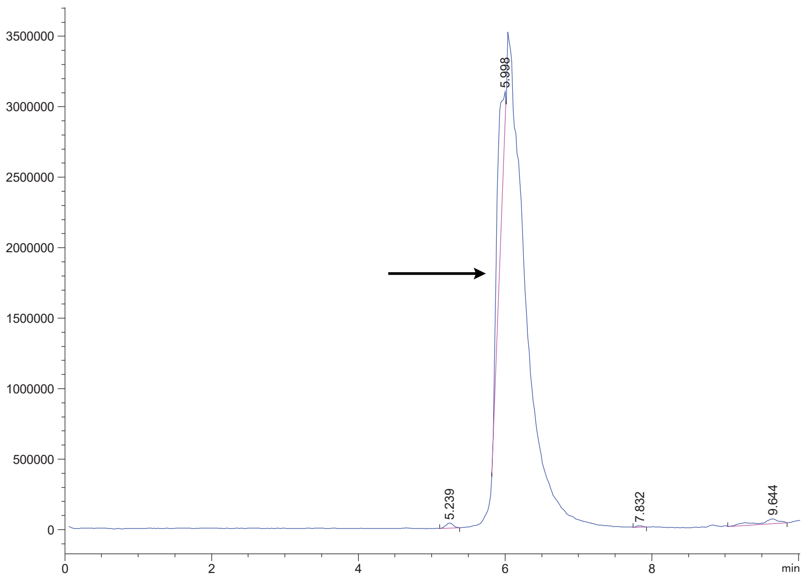


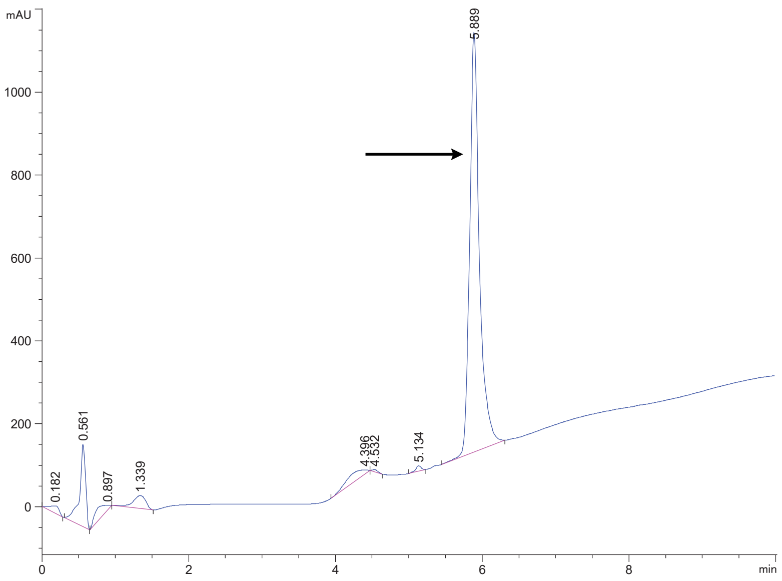

FAP 24.2

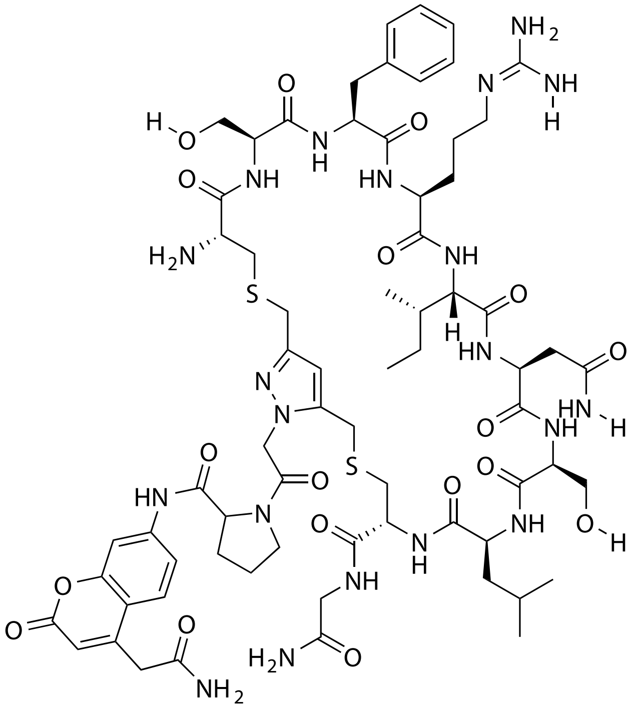

| **Probe Name** | **Sequence** | **Expected Mass $[M+1]^{+1}$ or $[M+2]^{+2}$** | **Found Mass** |
|---|---|---|---|
| FAP 24.2 | CSFRINSLCG | 1545.6686 | 1545.9 |

Total Ion Chromatogram

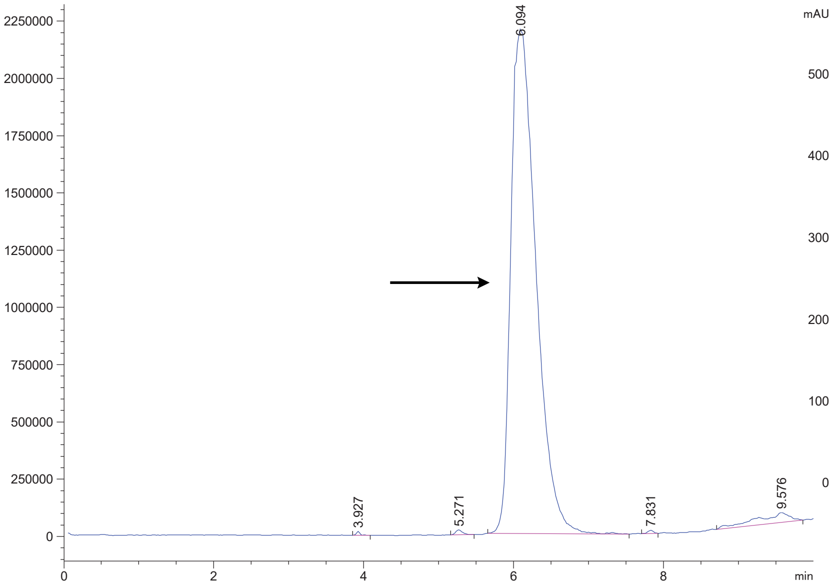


Variable Wavelength Detector: 220 nm

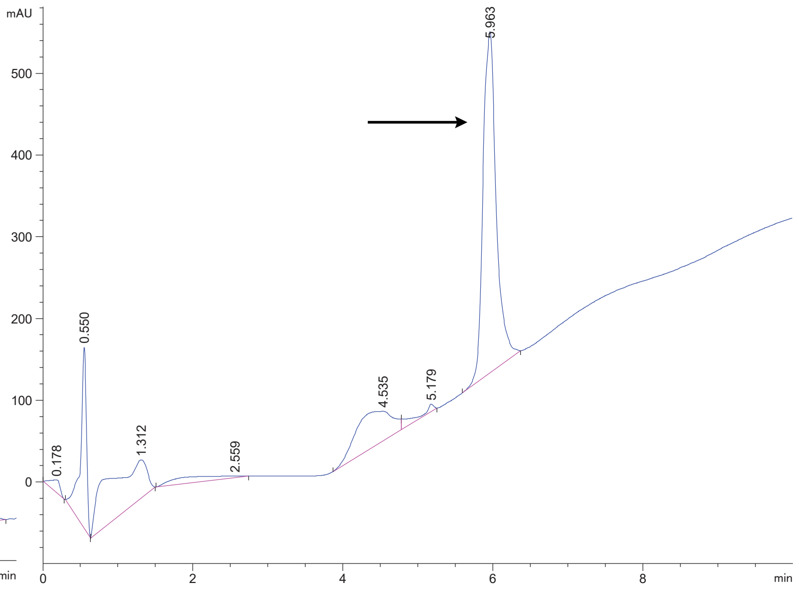

FAP 24.3

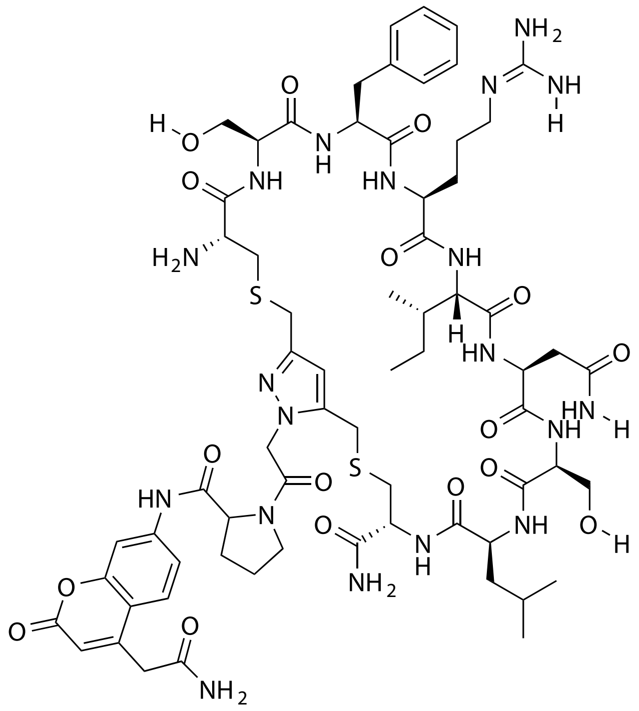

| **Probe Name** | **Sequence** | **Expected Mass $[M+1]^{+1}$ or $[M+2]^{+2}$** | **Found Mass** |
|---|---|---|---|
| FAP 24.3 | CSFRINSLC | 744.8275 $[M+2]^{+2}$ | 744.7 $[M+2]^{+2}$ |

Total Ion Chromatogram

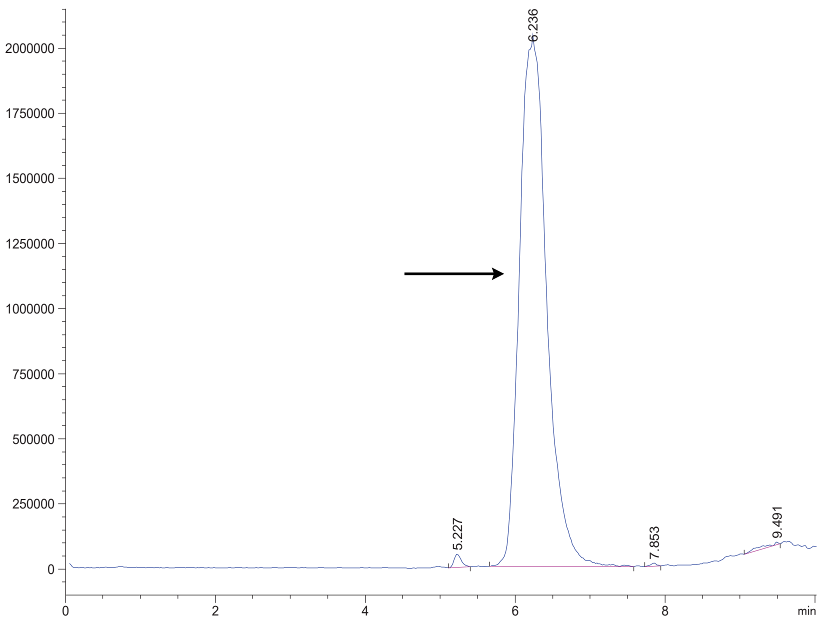


Variable Wavelength Detector: 220 nm

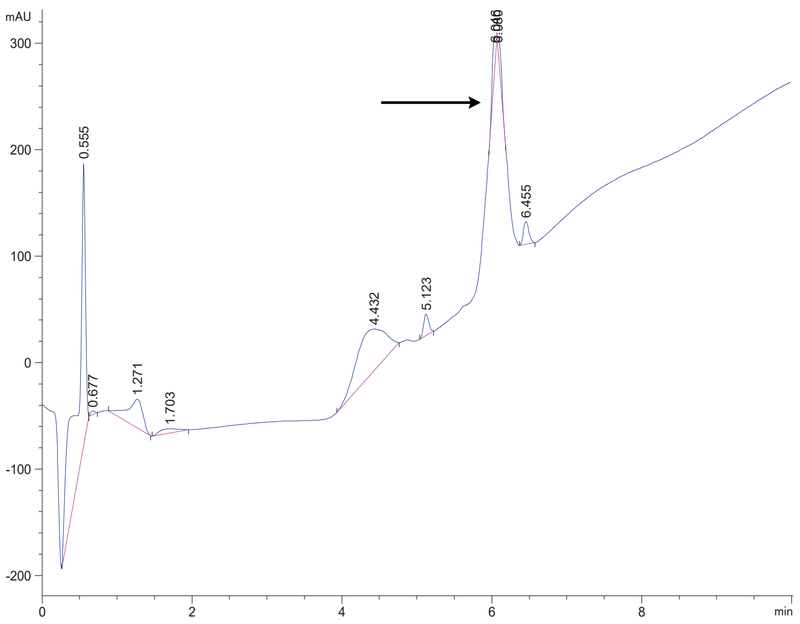

FAP 24.4

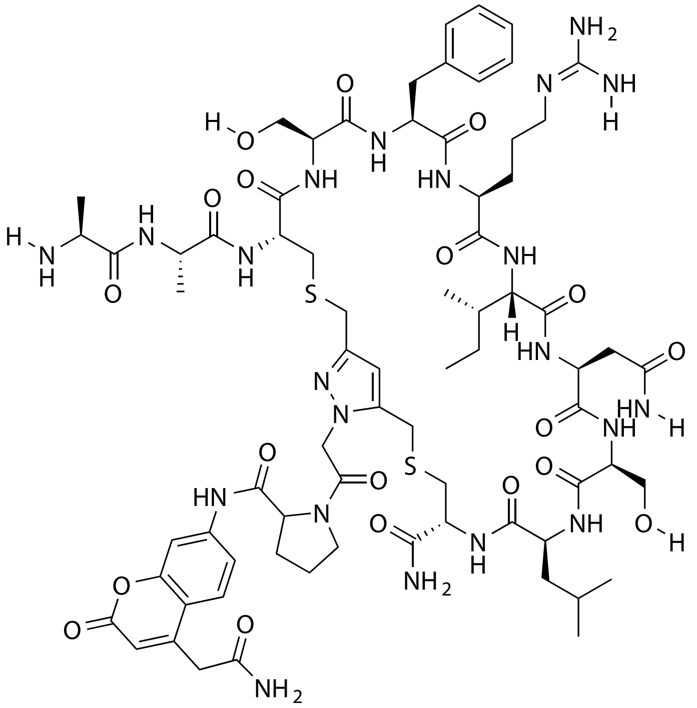


| Probe Name | Sequence | Expected Mass $[M+1]^{+1}$ or $[M+2]^{+2}$ | Found Mass |
|---|---|---|---|
| FAP 24.4 | AACSFRINSLC | 1630.7214 | 1631.0 |

Total Ion Chromatogram

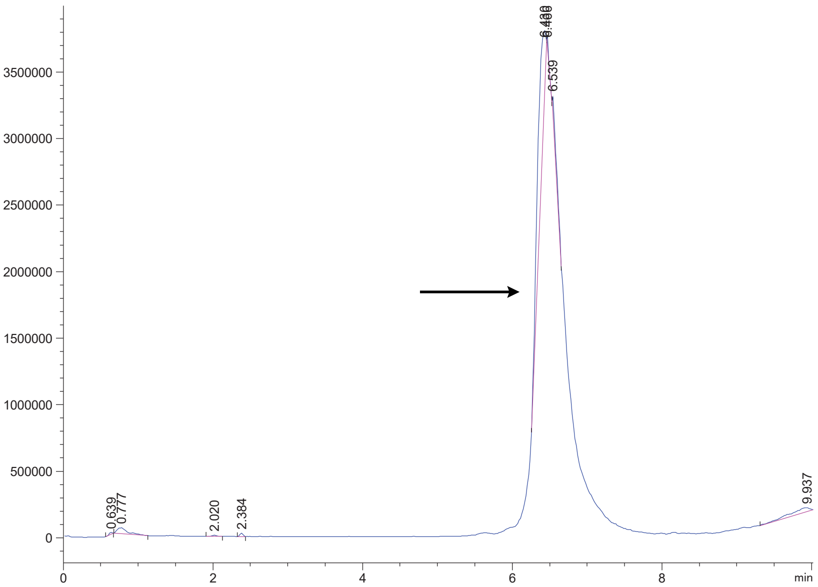


Variable Wavelength Detector: 220 nm

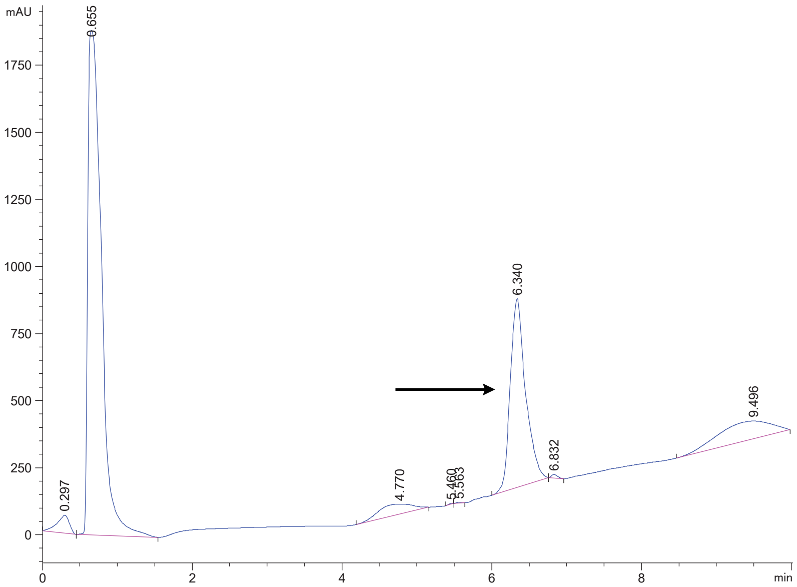

FAP 24.5

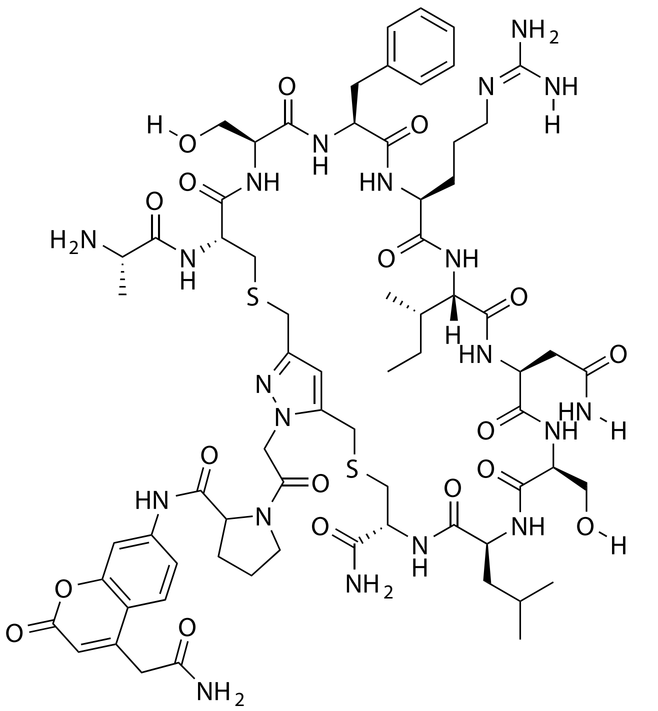

| Probe Name | Sequence | Expected Mass [M+1]$^{+1}$ or [M+2]$^{+2}$ | Found Mass |
|---|---|---|---|
| FAP 24.5 | ACSFRINSLC | 1559.6843 | 1559.9 |

Total Ion Chromatogram

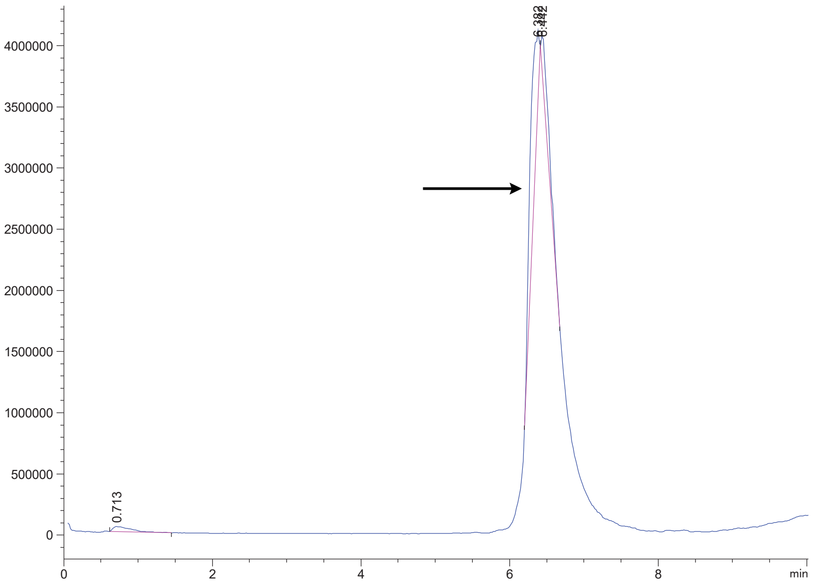

Variable Wavelength Detector: 220 nm

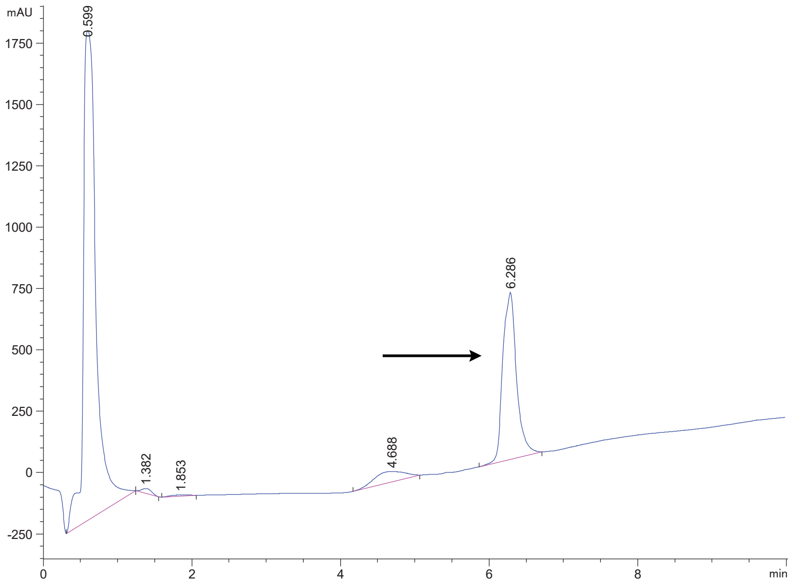

FAP 24s

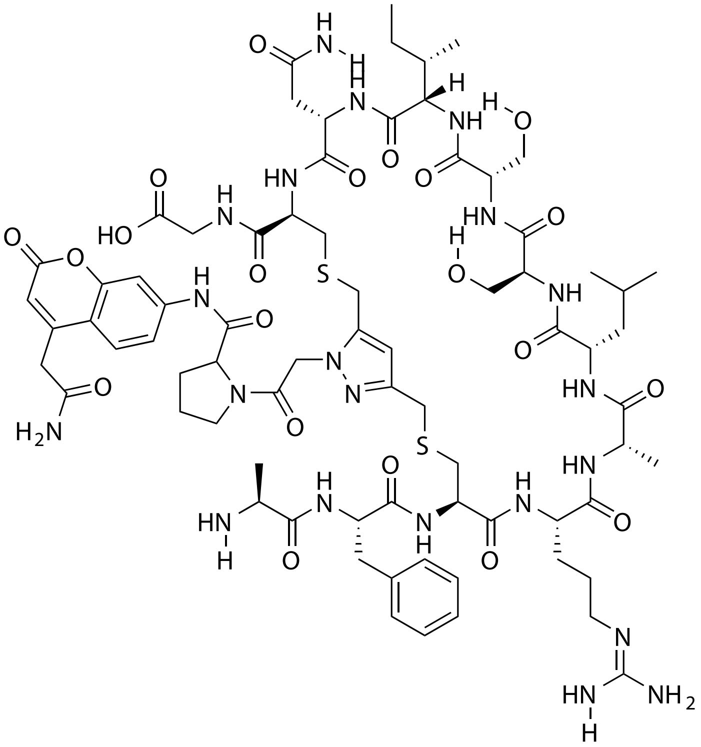

| Probe Name | Sequence | Expected Mass $[M+1]^{+1}$ or $[M+2]^{+2}$ | Found Mass |
|---|---|---|---|
| FAP 24s | AFCRALSSINCG | 1687.7429 | 1688.0 |

Total Ion Chromatogram

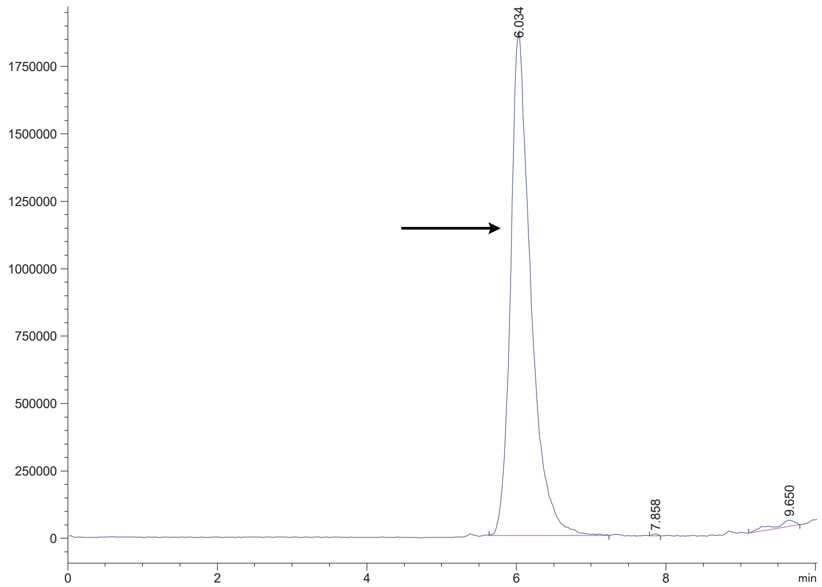


Variable Wavelength Detector: 220 nm

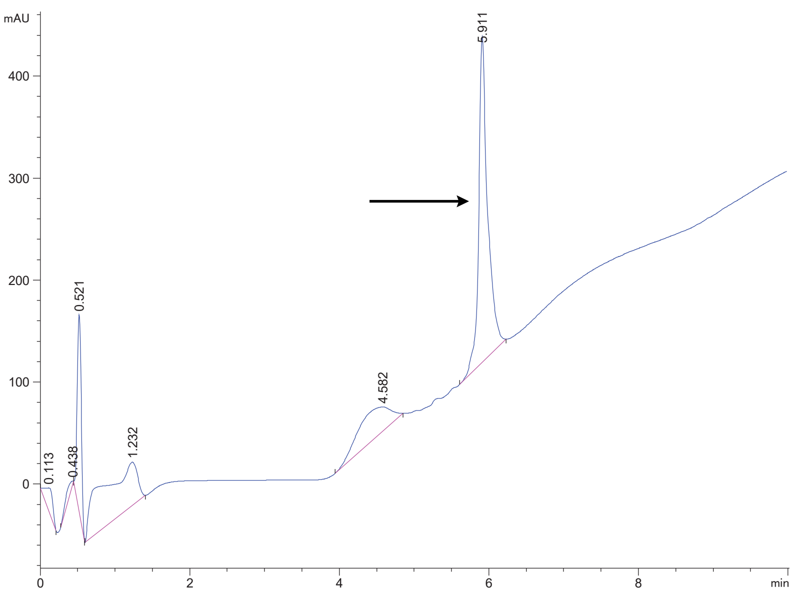

FAP 28

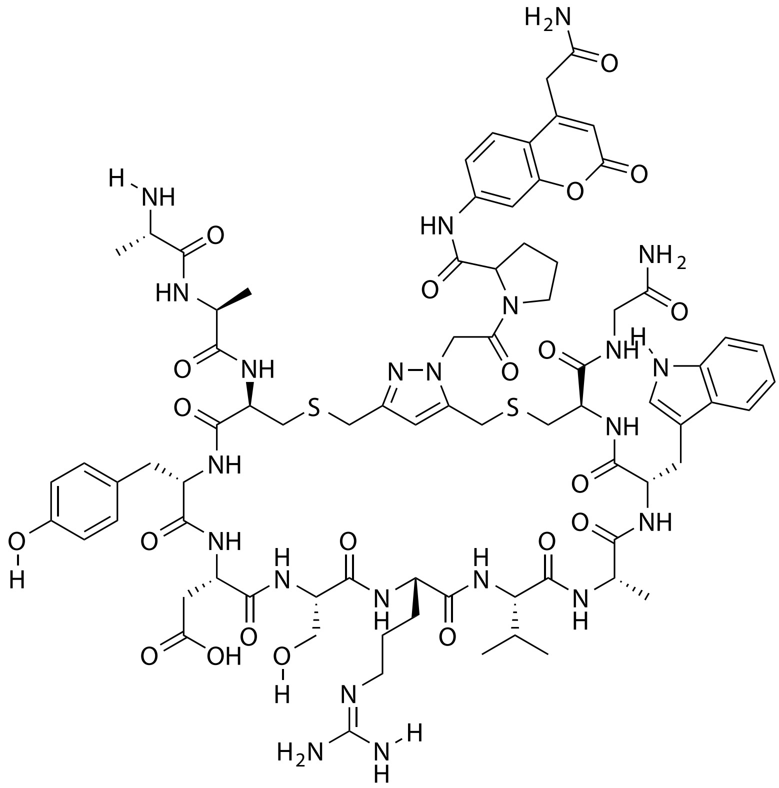

| Probe Name | Sequence | Expected Mass $[M+1]^{+1}$ or $[M+2]^{+2}$ | Found Mass |
|---|---|---|---|
| FAP 28 | AACYDSRVAWCG | 874.3572 $[M+2]^{+2}$ | 874.2 $[M+2]^{+2}$ |

Total Ion Chromatogram

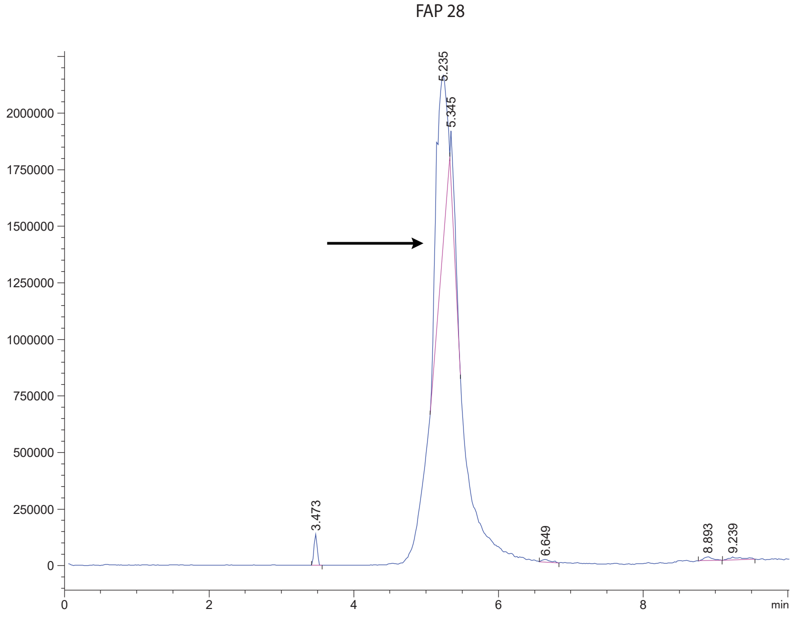


Variable Wavelength Detector: 220 nm

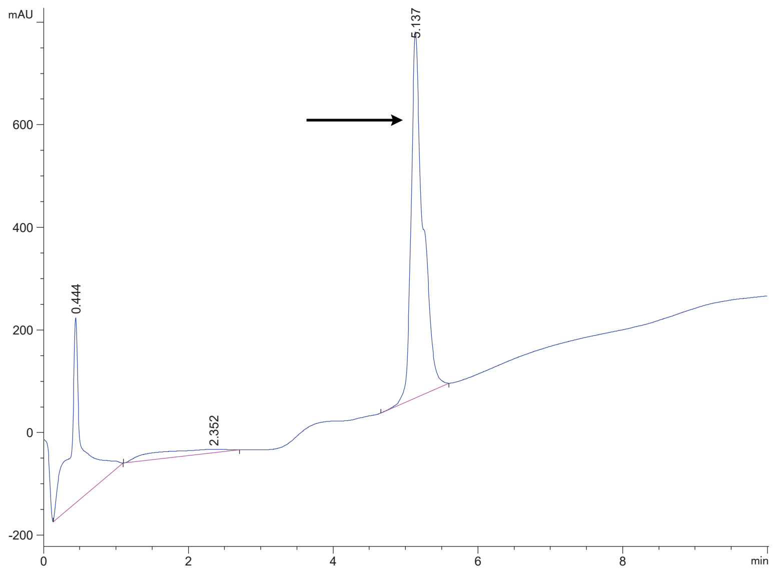

FAP 32

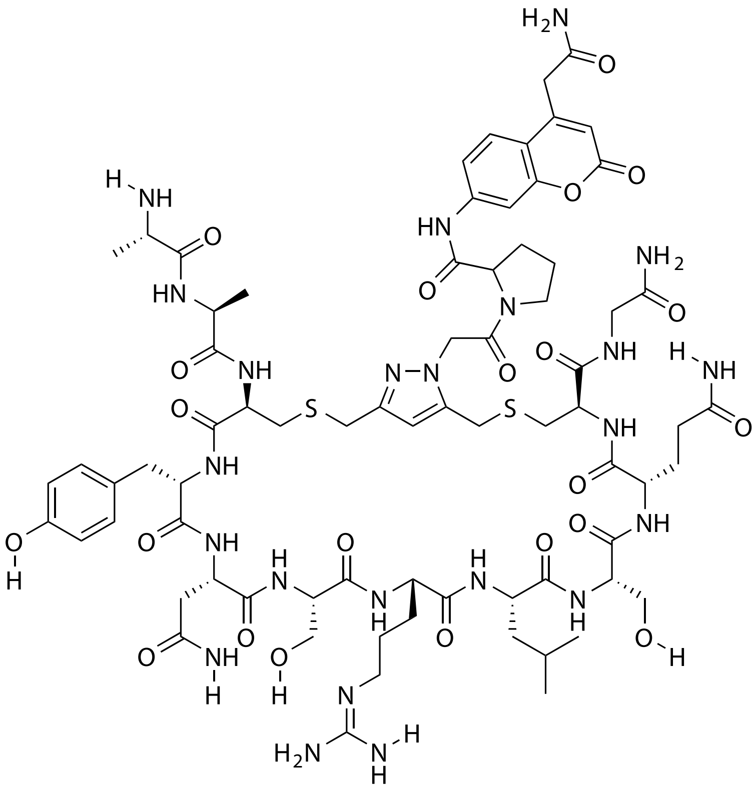

| Probe Name | Sequence | Expected Mass $[M+1]^{+1}$ or $[M+2]^{+2}$ | Found Mass |
|---|---|---|---|
| FAP 32 | AACYNSRLSQCG | 1718.7123 | 1717.9 |

Total Ion Chromatogram

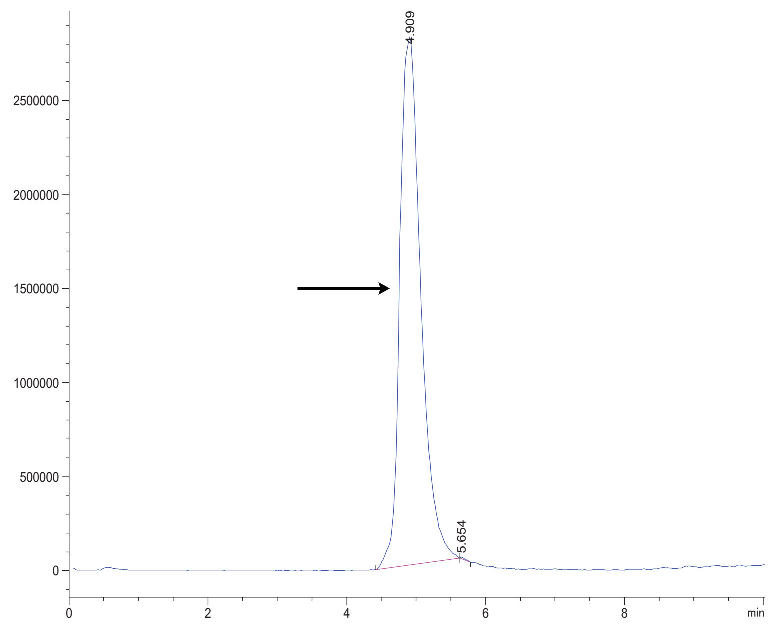

Variable Wavelength Detector: 220 nm

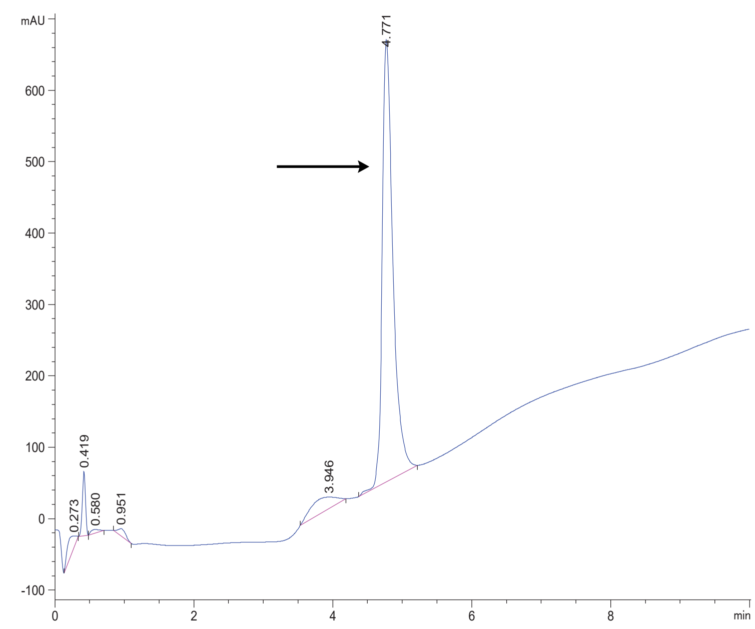

FAP 33

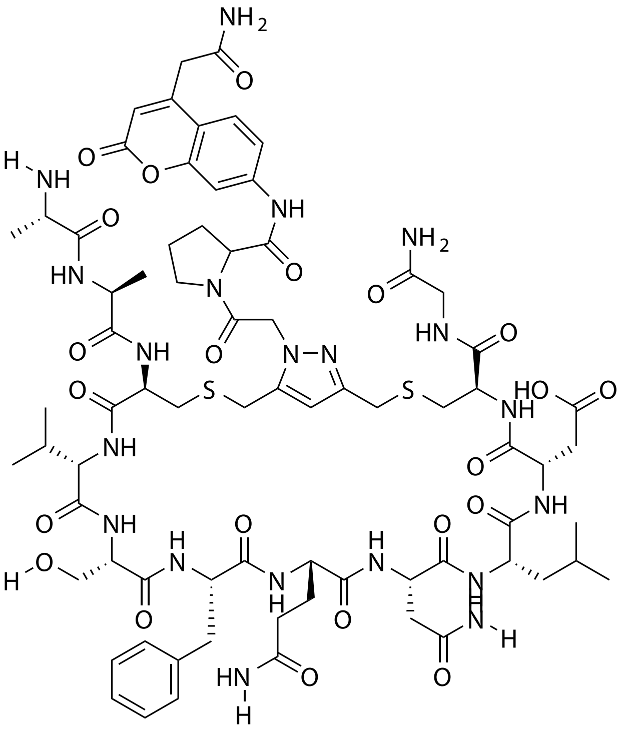

| Probe Name | Sequence | Expected Mass $[M+1]^{+1}$ or $[M+2]^{+2}$ | Found Mass |
|---|---|---|---|
| FAP 33 | AACVSFQNLDCG | 837.3437 | 837.2 $[M+2]^{+2}$ |

Total Ion Chromatogram

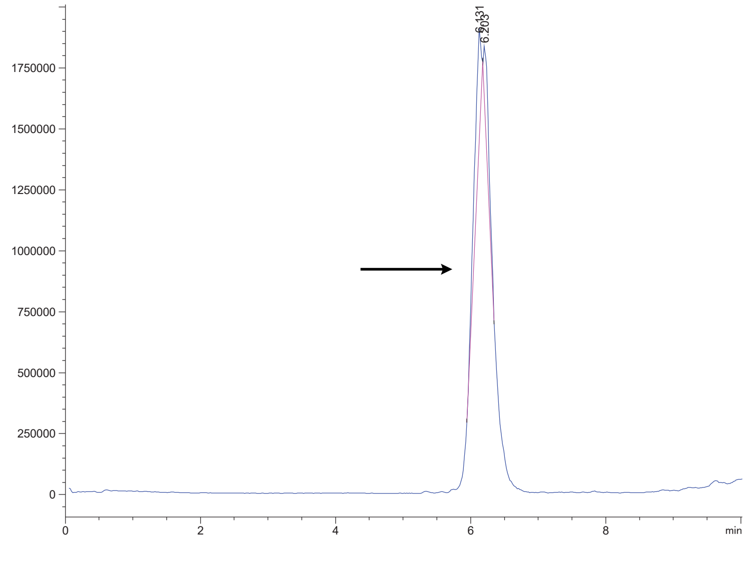

Variable Wavelength Detector: 220 nm

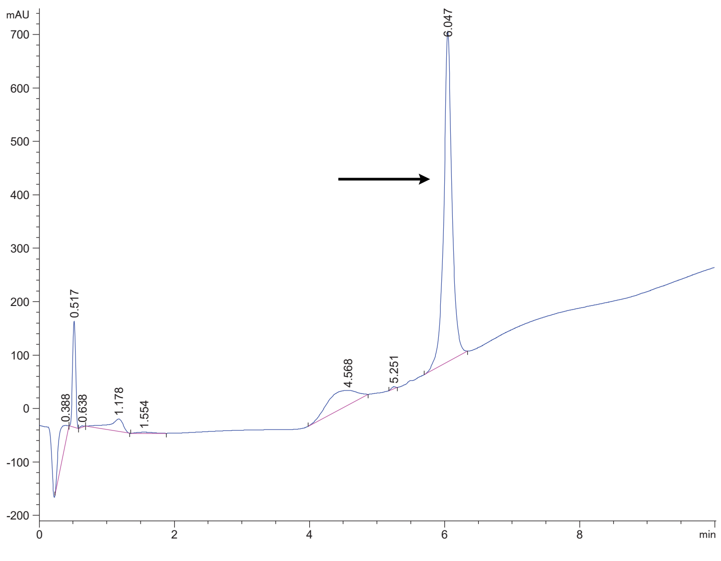

FAP 33.1

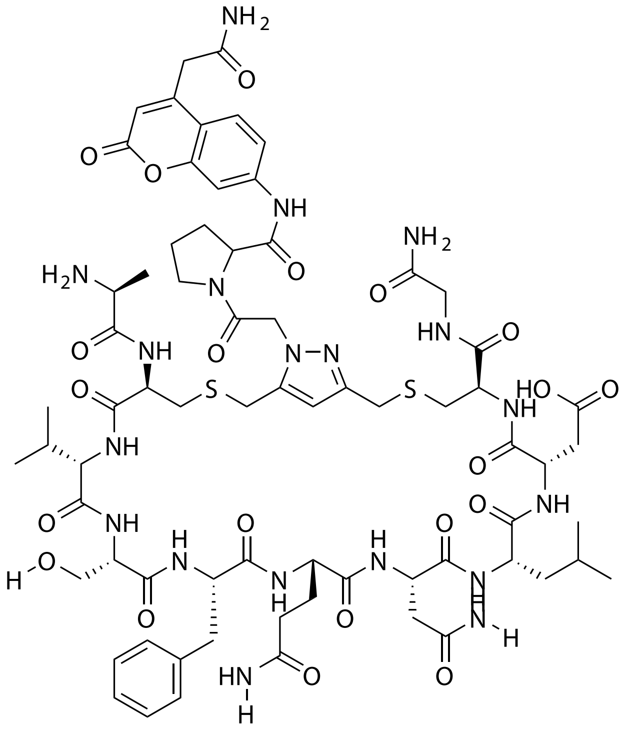


| Probe Name | Sequence | Expected Mass $[M+1]^{+1}$ or $[M+2]^{+2}$ | Found Mass |
|---|---|---|---|
| FAP 33.1 | ACVSFQNLDCG | 1602.6425 | 1602.9 |

Total Ion Chromatogram

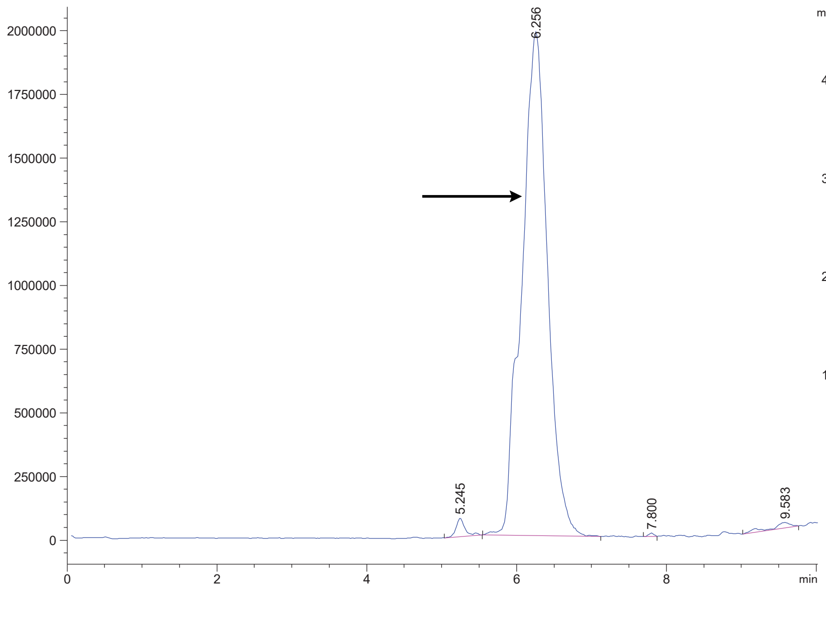


Variable Wavelength Detector: 220 nm

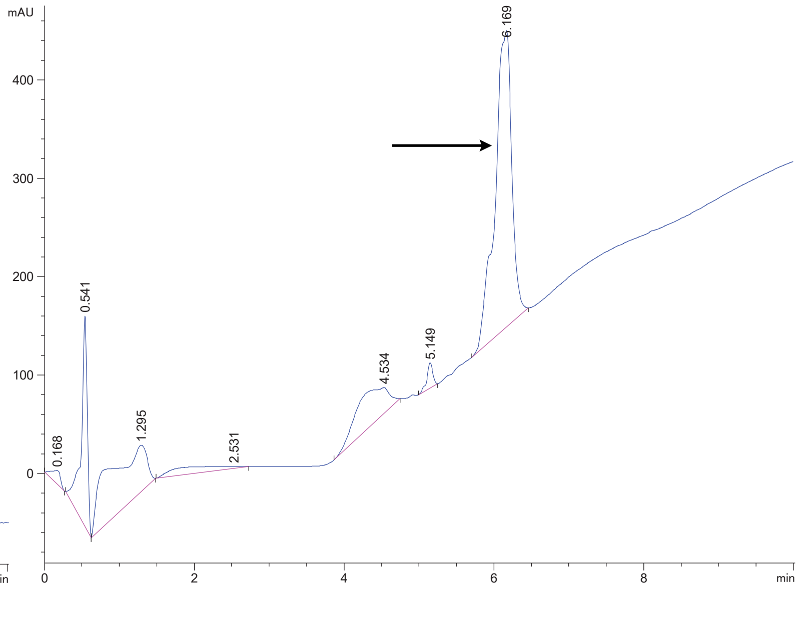

FAP 33.2

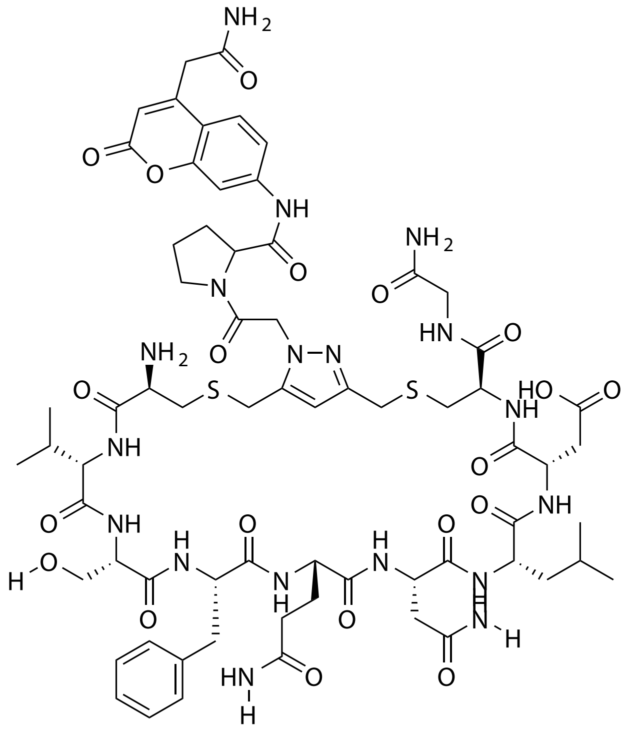

| Probe Name | Sequence | Expected Mass $[M+1]^{+1}$ or $[M+2]^{+2}$ | Found Mass |
|---|---|---|---|
| FAP 33.2 | CVSFQNLDCG | 1531.6054 | 1531.9 |

Total Ion Chromatogram

Variable Wavelength Detector: 220 nm

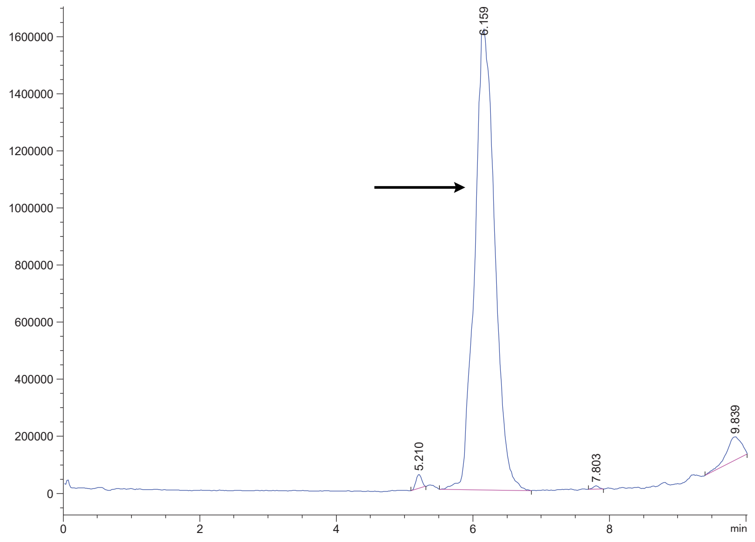


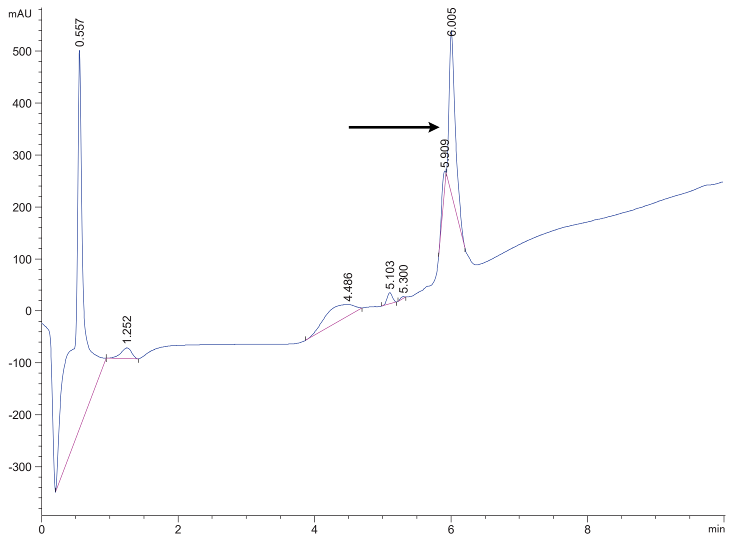

### FAP 33.2A1

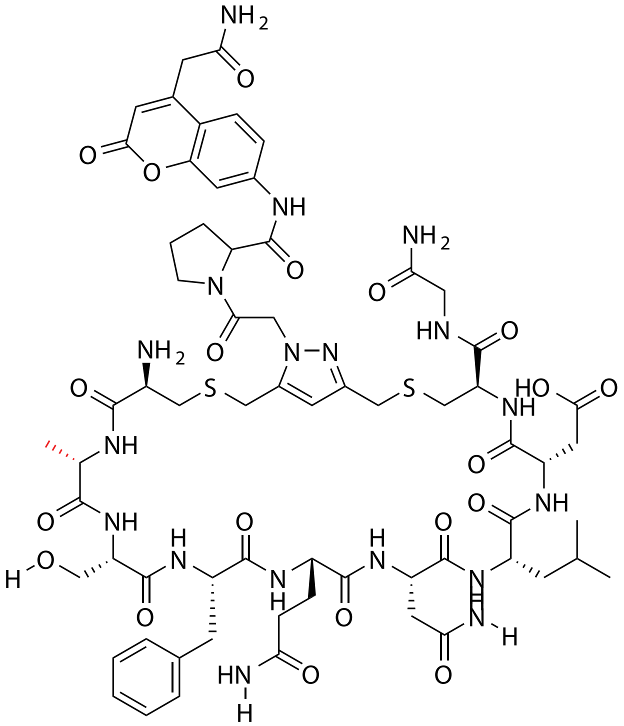


| Probe Name | Sequence | Expected Mass $[M+1]^{+1}$ or $[M+2]^{+2}$ | Found Mass |
|---|---|---|---|
| FAP 33.2 A1 | CASFQNLDCG | 1503.5741 | 1503.8 |

Total Ion Chromatogram

Variable Wavelength Detector: 220 nm

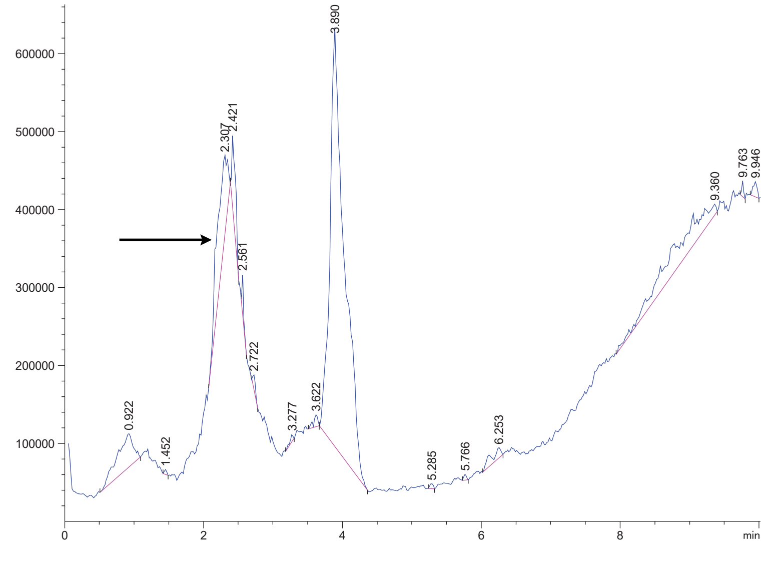


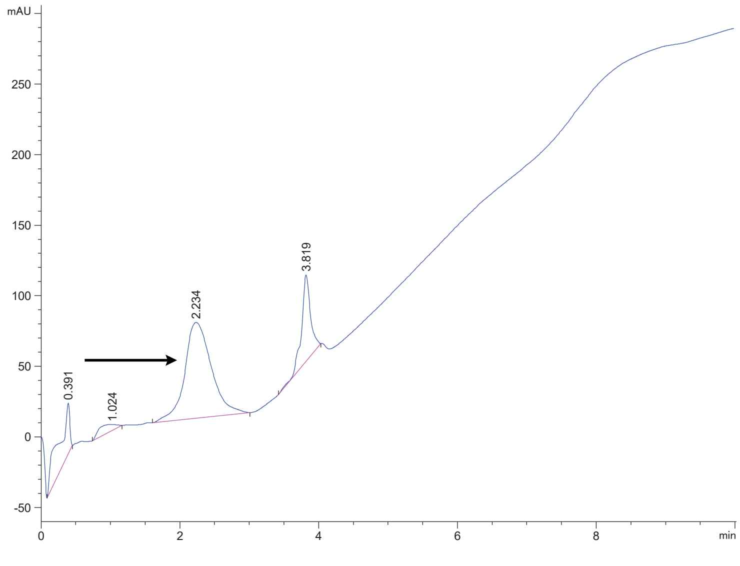

FAP 33.2A2

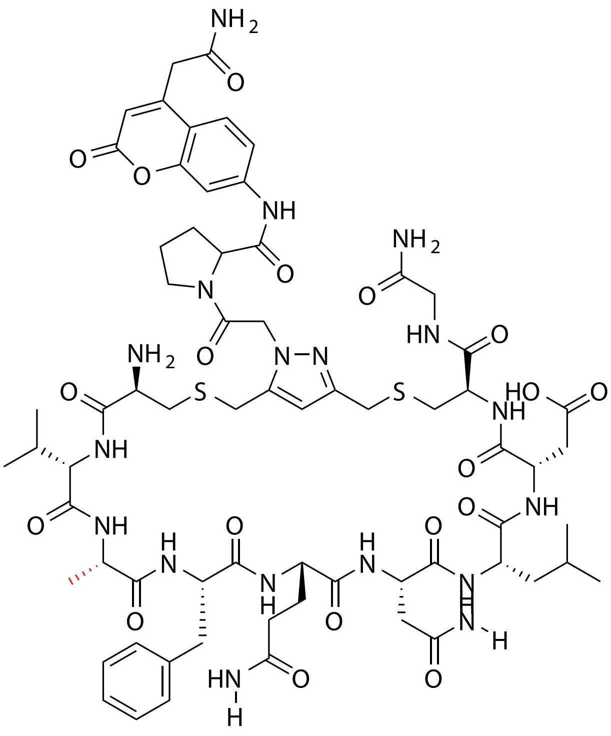

| Probe Name | Sequence | Expected Mass $[M+1]^{+1}$ or $[M+2]^{+2}$ | Found Mass |
|---|---|---|---|
| FAP 33.2 A2 | CVAFQNLDCG | 1515.6105 | 1515.9 |

Total Ion Chromatogram

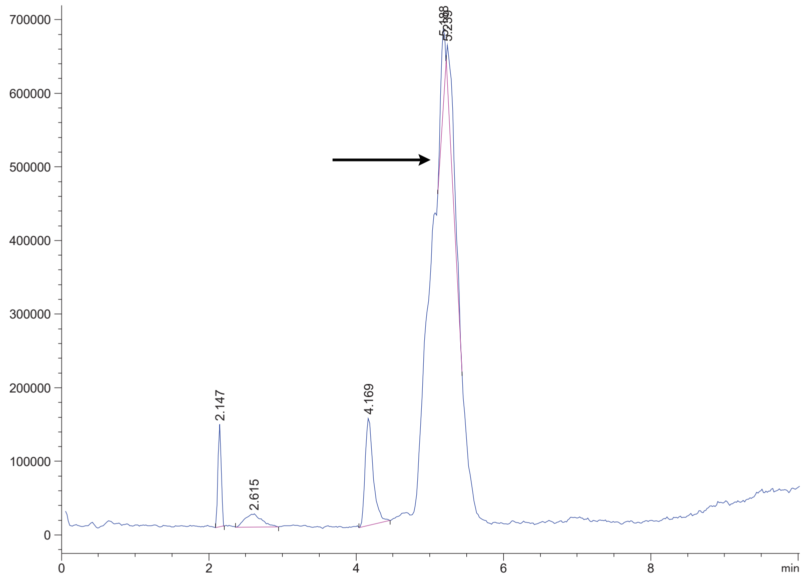


Variable Wavelength Detector: 220 nm

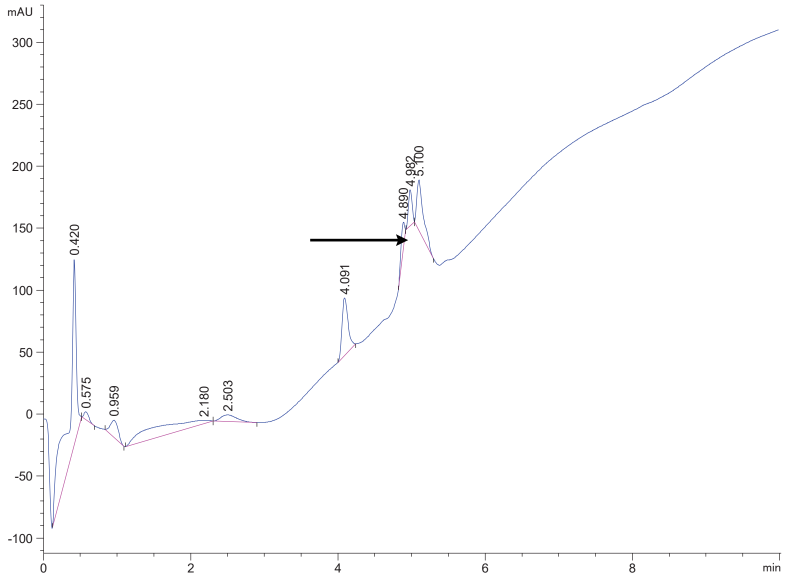

FAP 33.2A3

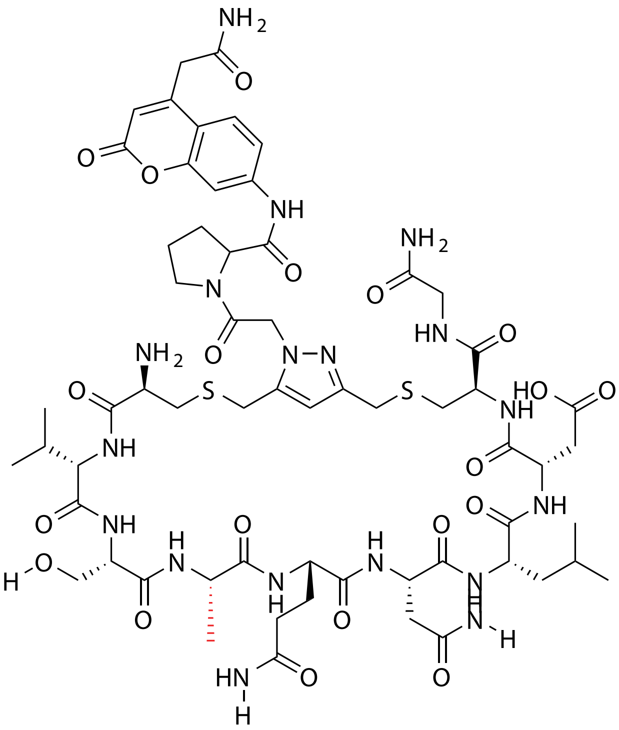

| Probe Name | Sequence | Expected Mass [M+1]$^{+1}$ or [M+2]$^{+2}$ | Found Mass |
|---|---|---|---|
| FAP 33.2 A3 | CVSAQNLDCG | 1455.5741 | 1455.8 |

Total Ion Chromatogram

Variable Wavelength Detector: 220 nm

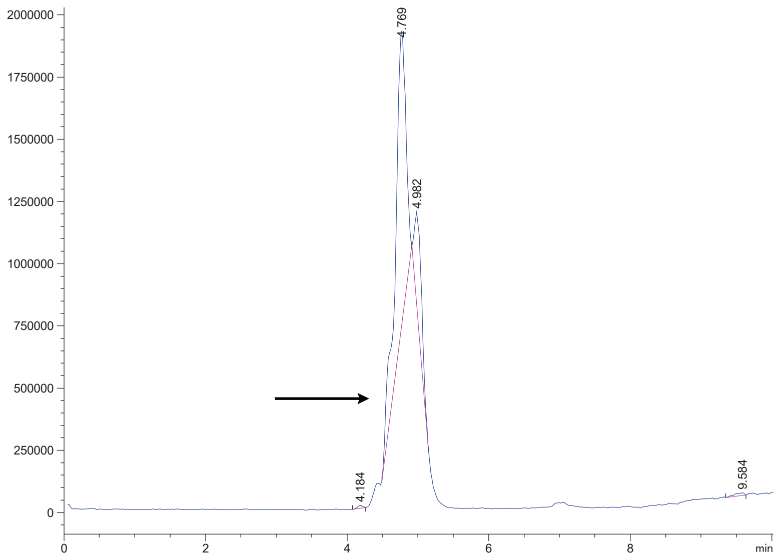


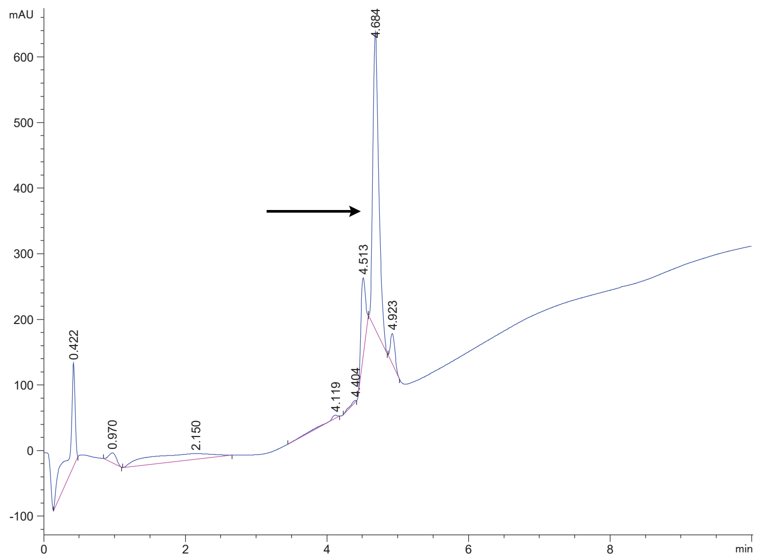

FAP 33.2A4

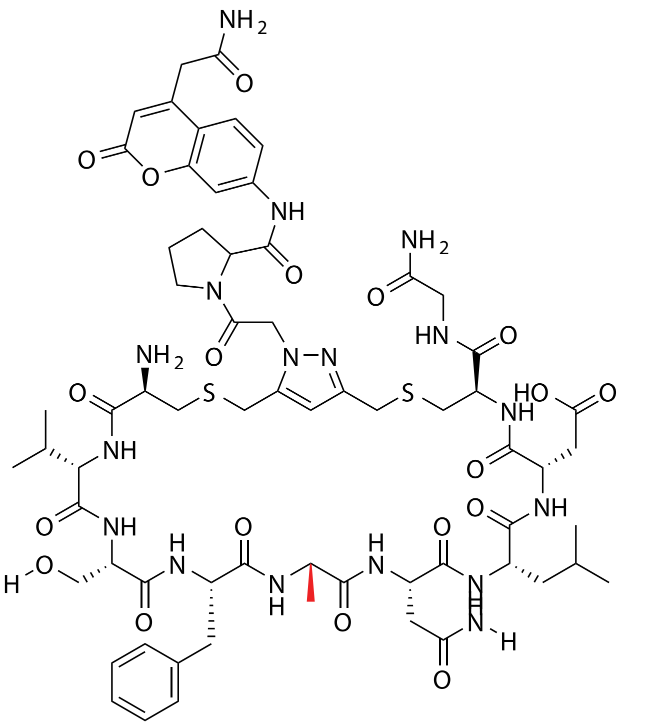

| Probe Name | Sequence | Expected Mass $[M+1]^{+1}$ or $[M+2]^{+2}$ | Found Mass |
|---|---|---|---|
| FAP 33.2 A4 | CVSFANLDCG | 1474.5839 | 1474.9 |

Total Ion Chromatogram

Variable Wavelength Detector: 220 nm

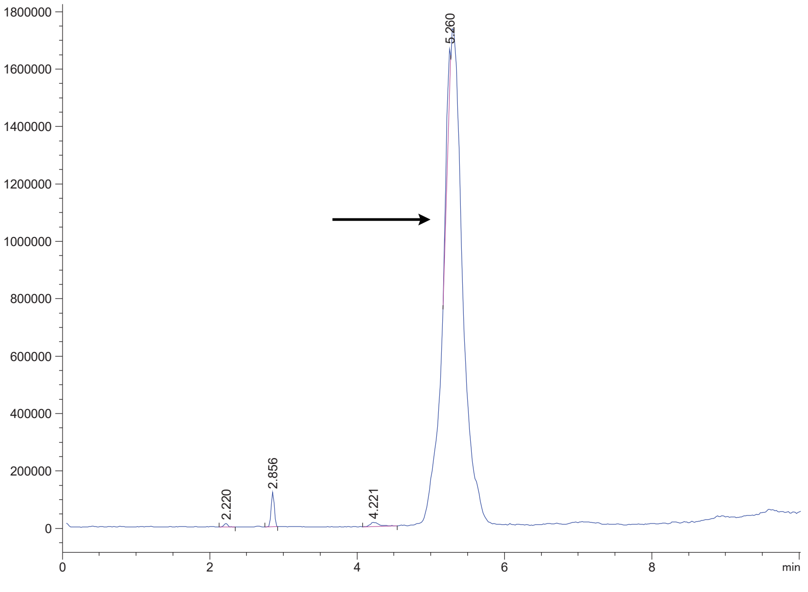


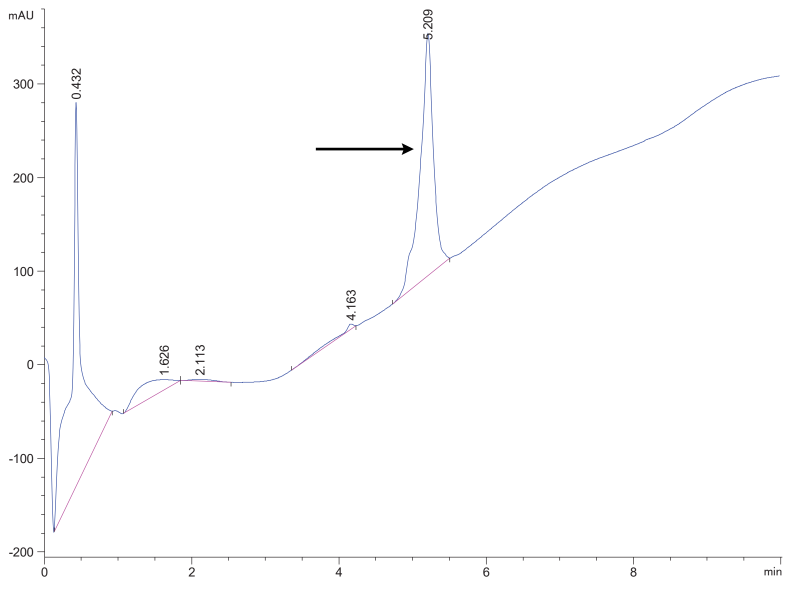

FAP 33.2A5

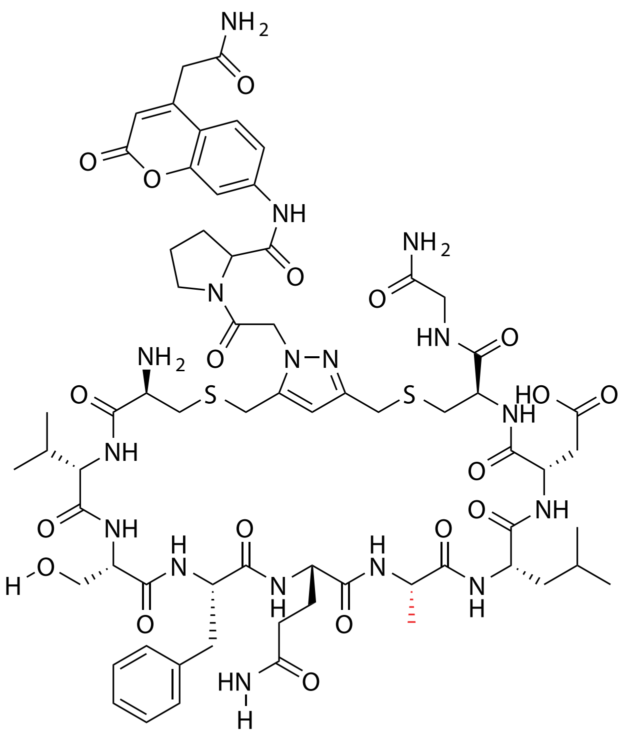


| Probe Name | Sequence | Expected Mass $[M+1]^{+1}$ or $[M+2]^{+2}$ | Found Mass |
|---|---|---|---|
| FAP 33.2 A5 | CVSFQALDCG | 1488.5996 | 1488.9 |

Total Ion Chromatogram

Variable Wavelength Detector: 220 nm

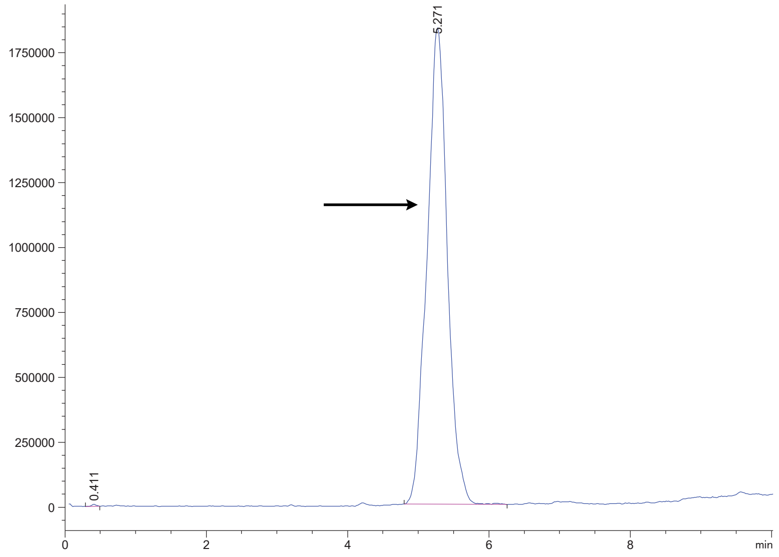


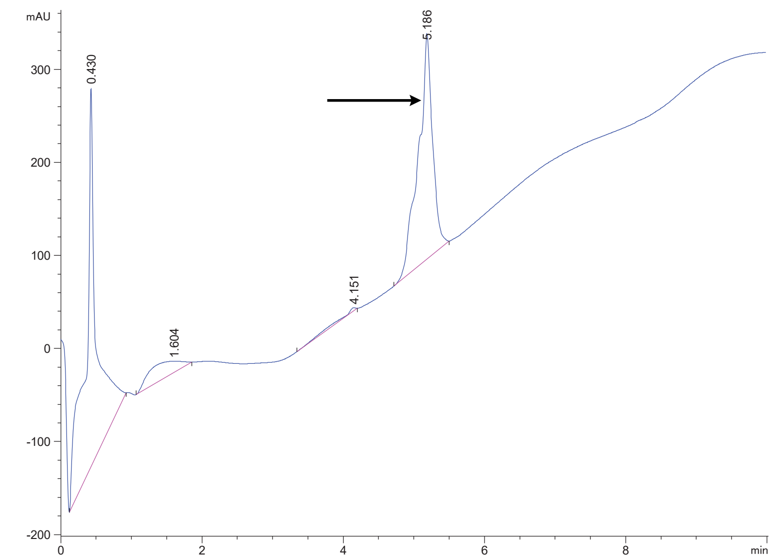

FAP 33.2A6

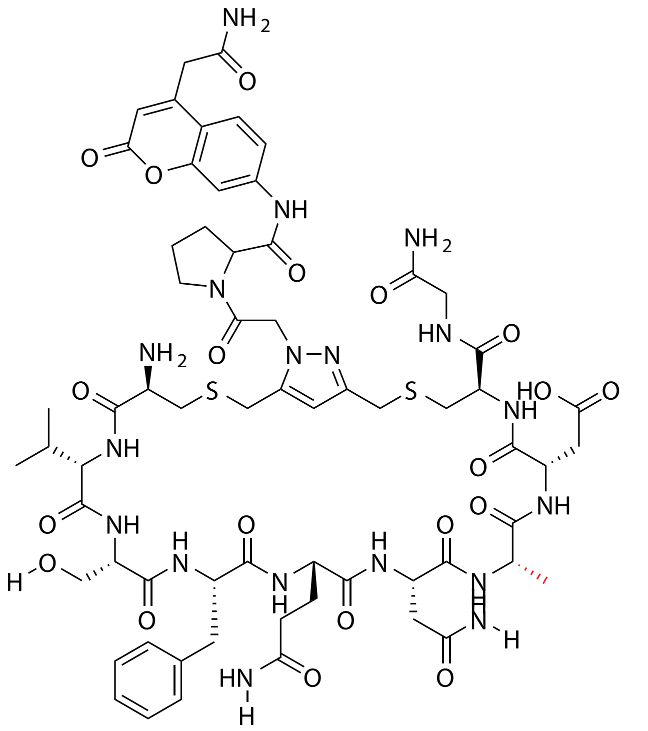


| Probe Name | Sequence | Expected Mass $[M+1]^{+1}$ or $[M+2]^{+2}$ | Found Mass |
|---|---|---|---|
| FAP 33.2 A6 | CVSFQNADCG | 1489.5584 | 1489.8 |

Total Ion Chromatogram

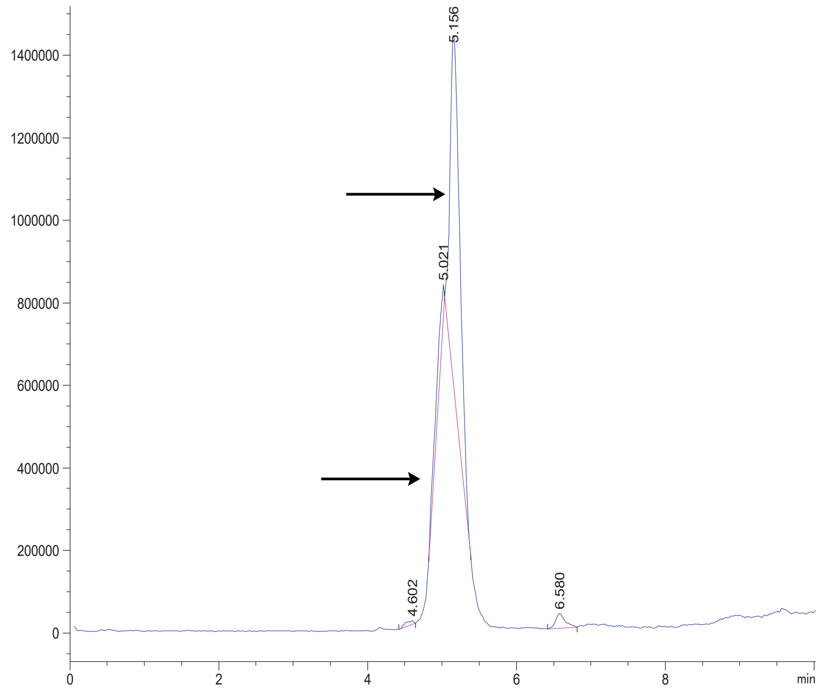


Variable Wavelength Detector: 220 nm

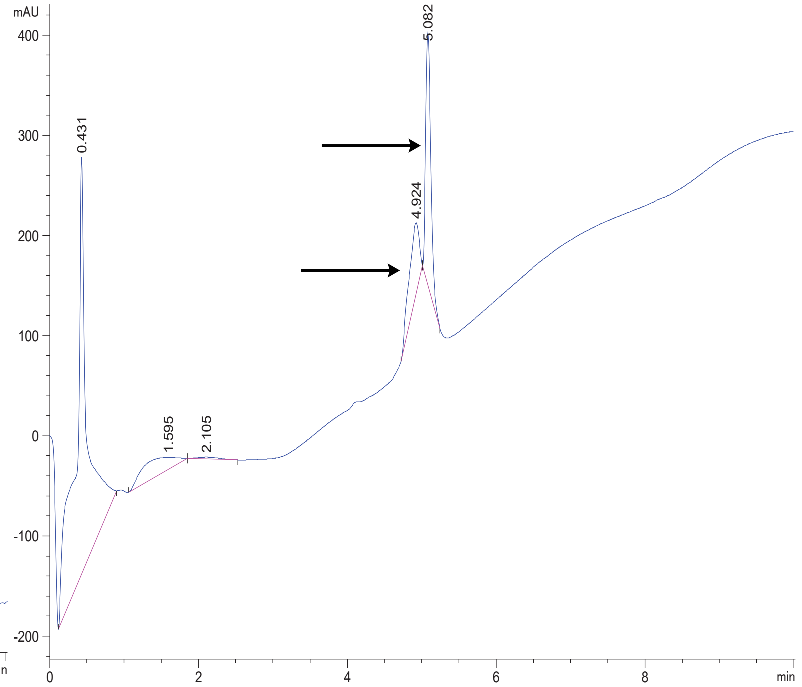

FAP 33.2A7

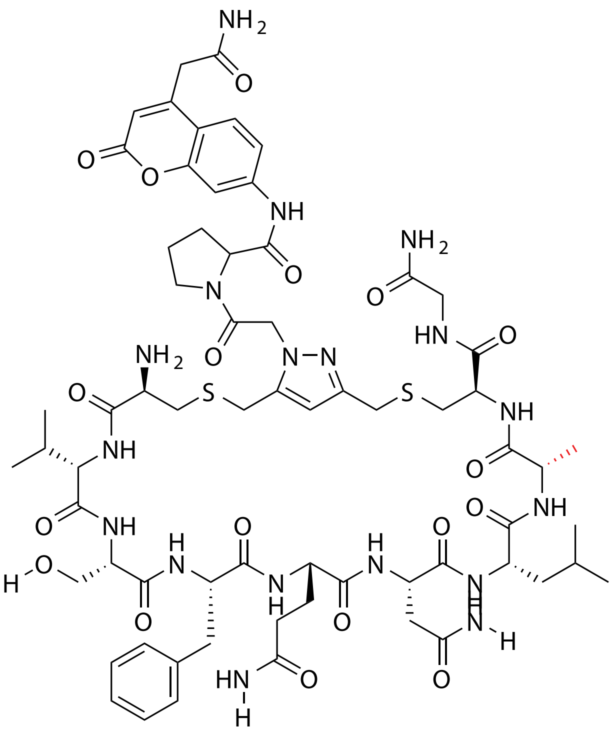

| Probe Name | Sequence | Expected Mass $[M+1]^{+1}$ or $[M+2]^{+2}$ | Found Mass |
|---|---|---|---|
| FAP 33.2 A7 | CVSFQNLACG | 1487.6155 | 1487.9 |

Total Ion Chromatogram

Variable Wavelength Detector: 220 nm

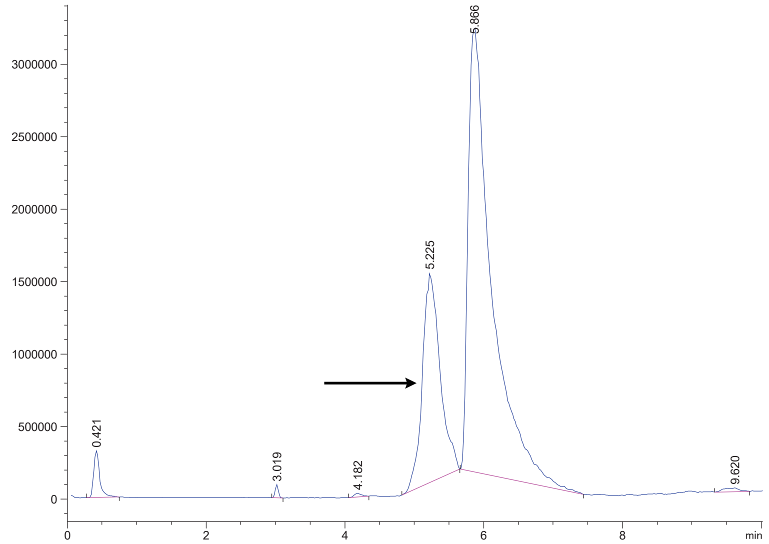


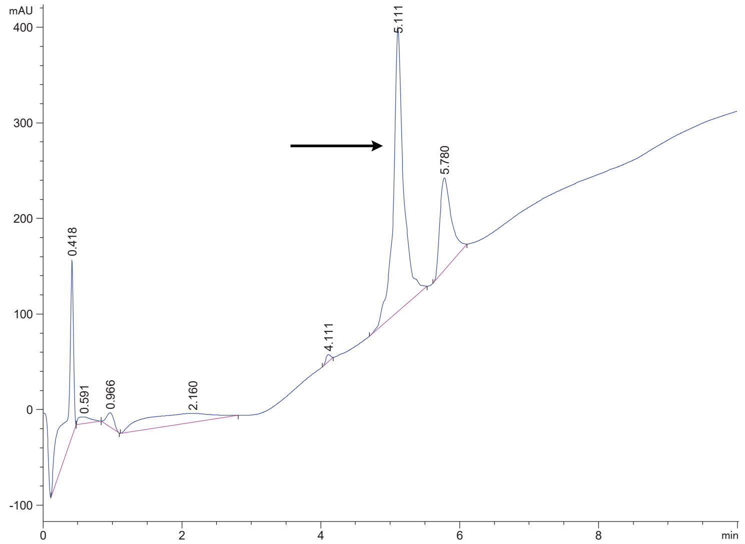

FAP 33.3

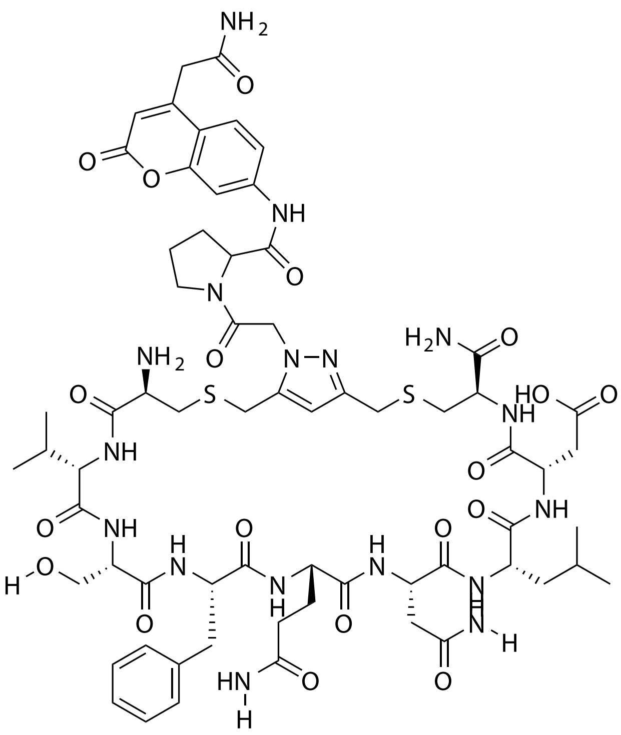

| Probe Name | Sequence | Expected Mass $[M+1]^{+1}$ or $[M+2]^{+2}$ | Found Mass |
|---|---|---|---|
| FAP 33.3 | CVSFQNLDC | 1474.5839 | 1474.9 |

Total Ion Chromatogram

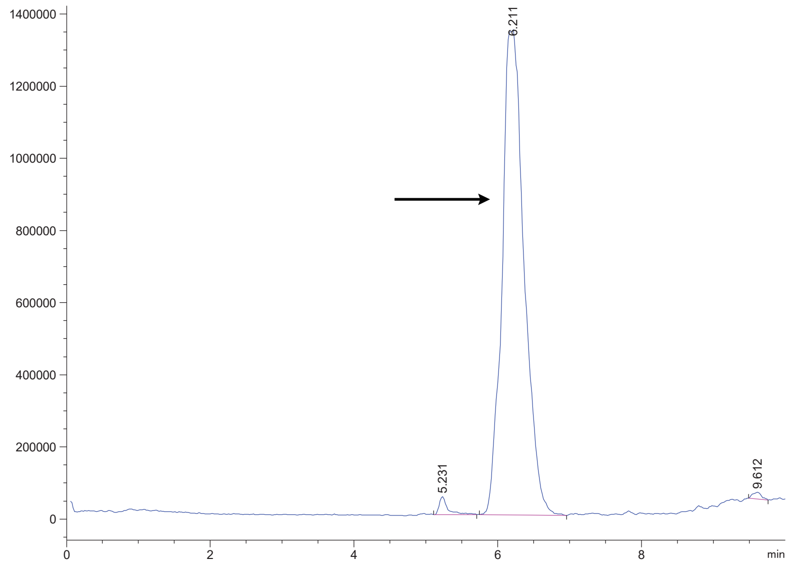

Variable Wavelength Detector: 220 nm

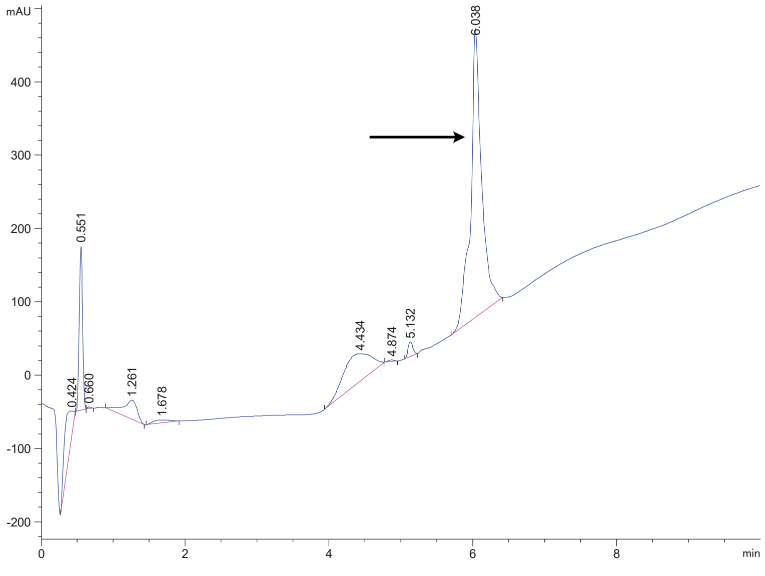

### FAP 33.5

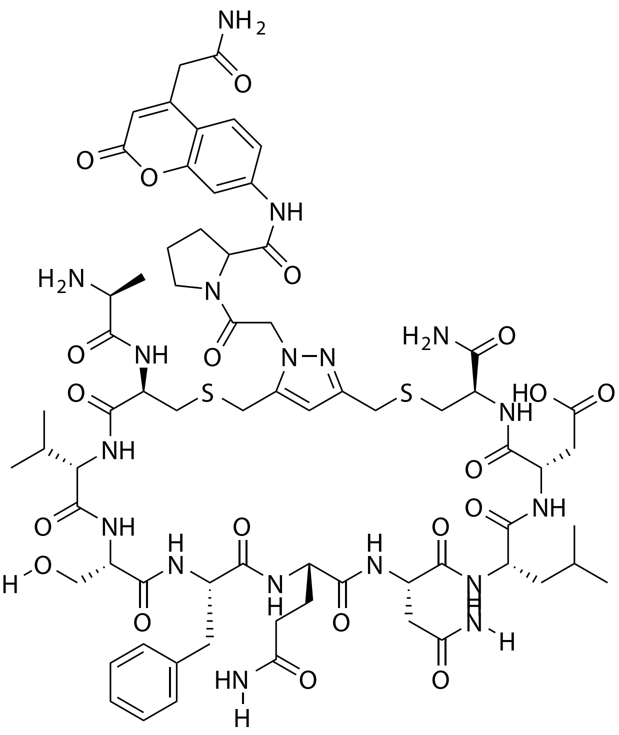

| Probe Name | Sequence | Expected Mass $[M+1]^{+1}$ or $[M+2]^{+2}$ | Found Mass |
|---|---|---|---|
| FAP 33.5 | ACVSFQNLDC | 1545.6210 | 1545.9 |

Total Ion Chromatogram

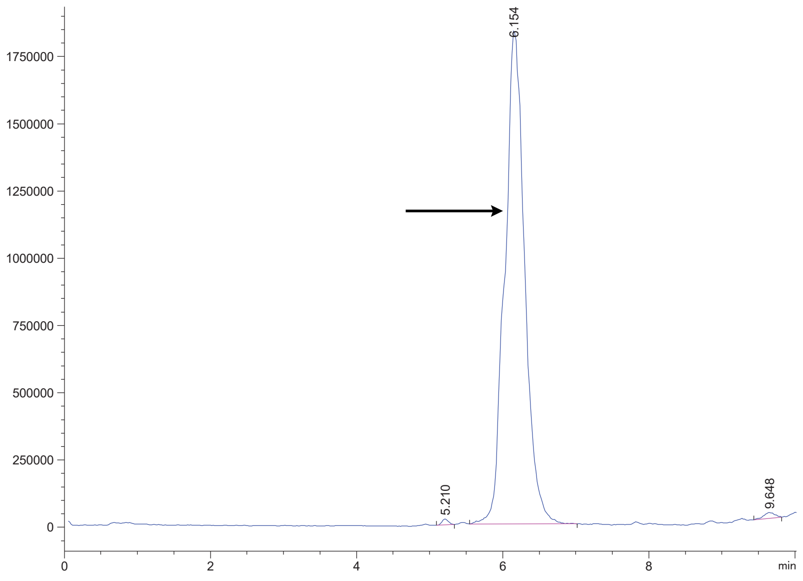


Variable Wavelength Detector: 220 nm

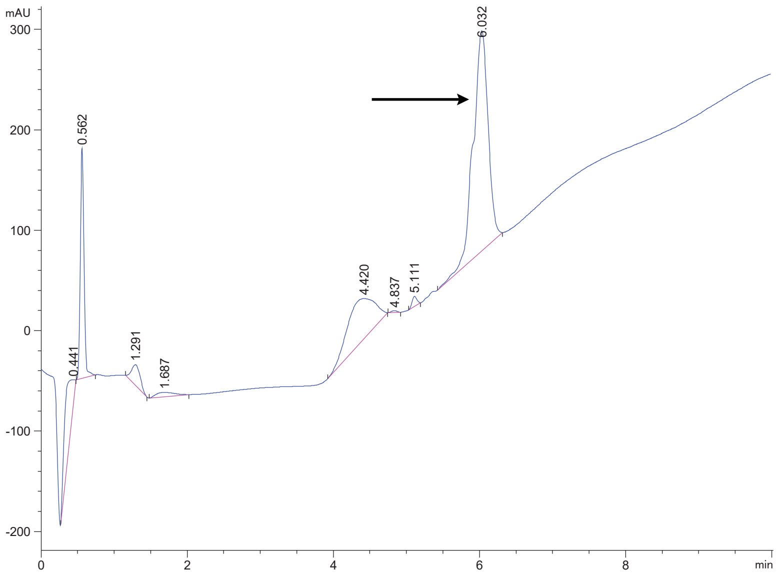

FAP 33s

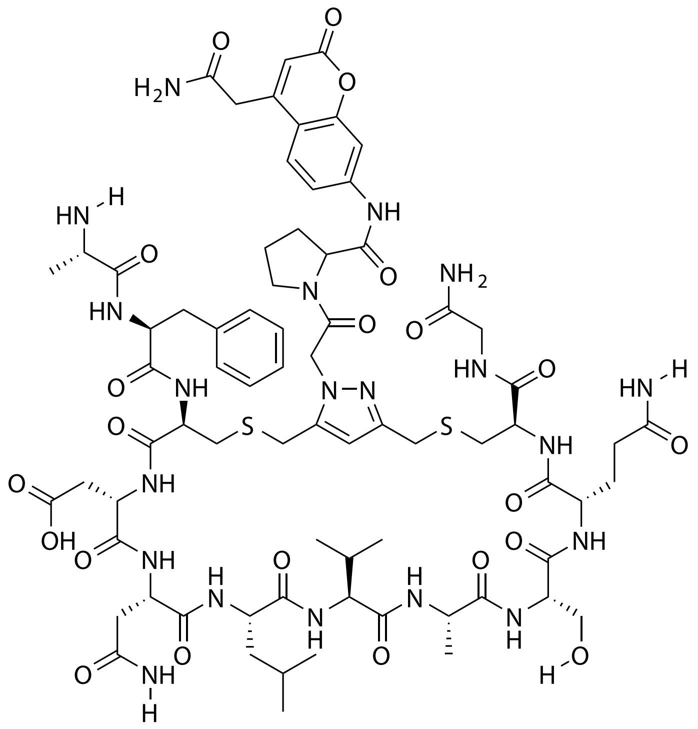


| Probe Name | Sequence | Expected Mass $[M+1]^{+1}$ or $[M+2]^{+2}$ | Found Mass |
|---|---|---|---|
| FAP 33s | AFCDNLVASQCG | 1673.6796 | 1673.9 |

Total Ion Chromatogram

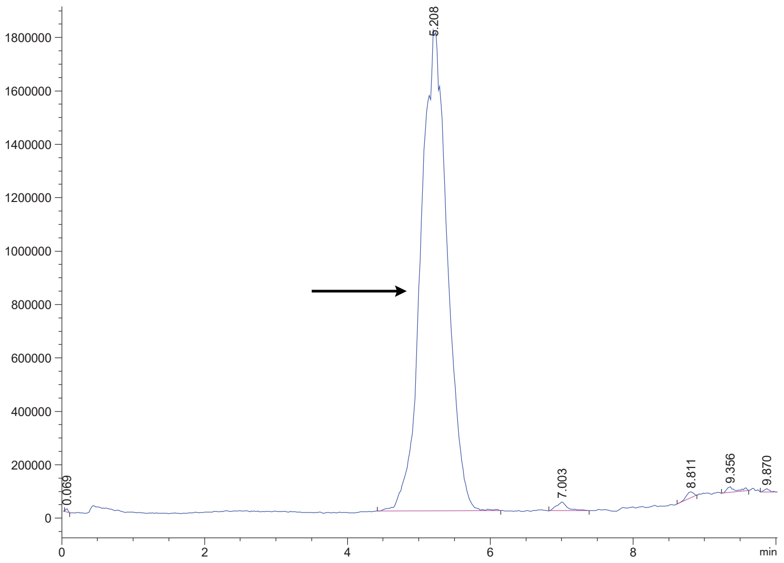


Variable Wavelength Detector: 220 nm

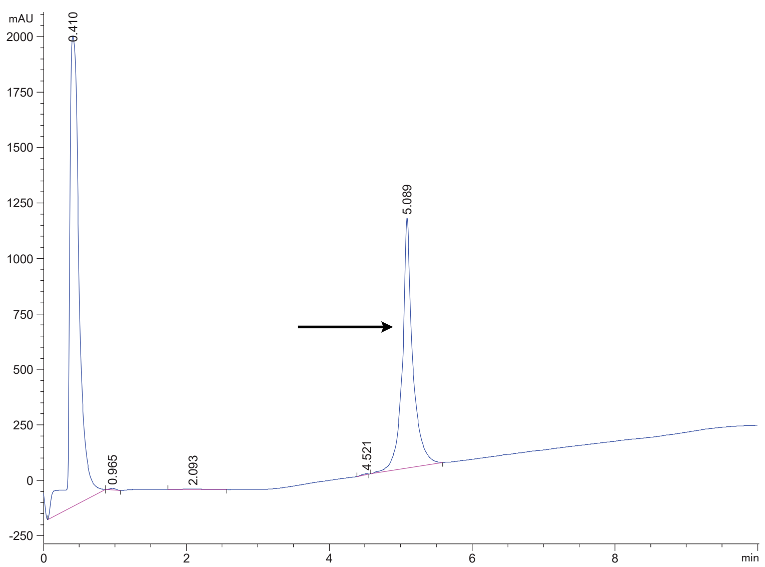

### FAP 33.2 (1)

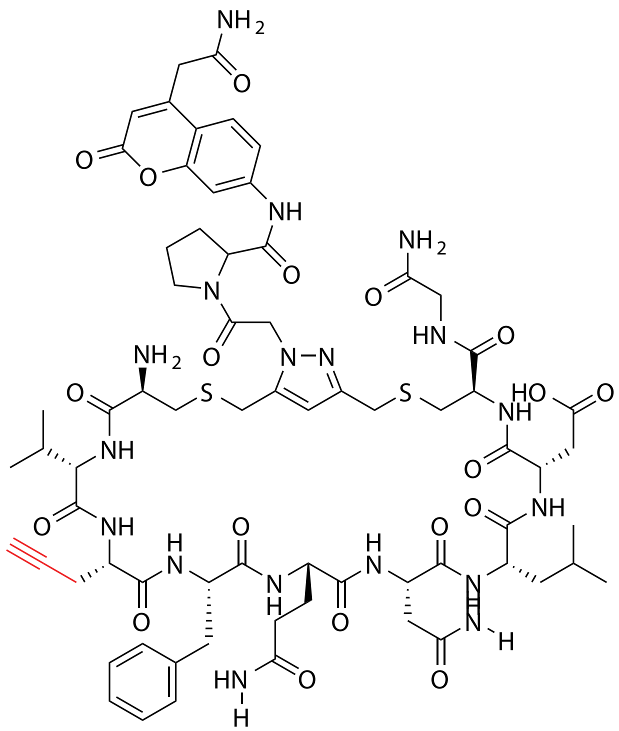


| Probe Name | Sequence X= Unnatural | Unnatural Amino Acid Number (X) | Expected Mass [M+1]+1 | Found Mass |
|---|---|---|---|---|
| FAP 33.2 (1) | CV**X**FQNLDCG | 1 | 1539.6105 | 1539.9 |

Total Ion Chromatogram

Variable Wavelength Detector: 220 nm

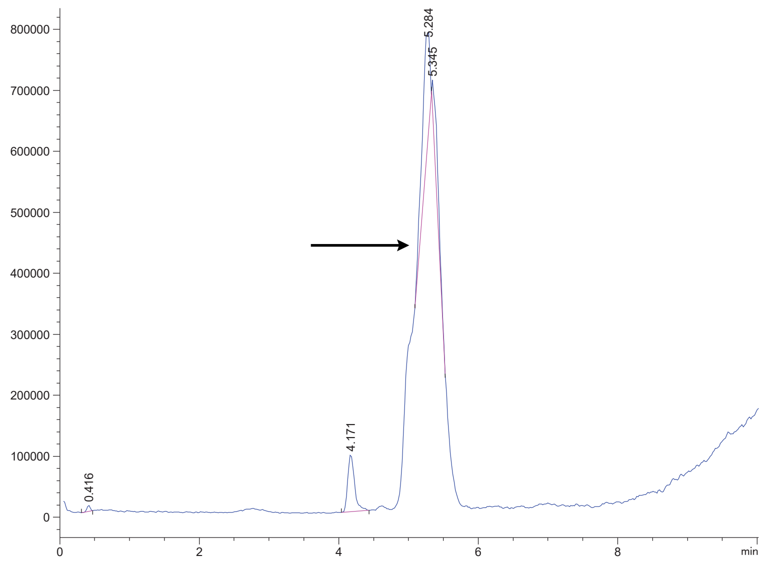


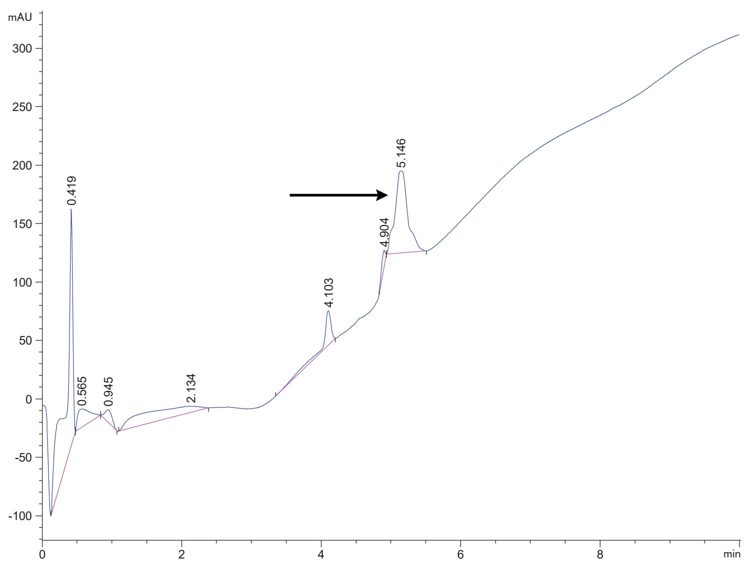

FAP 33.2 (3)

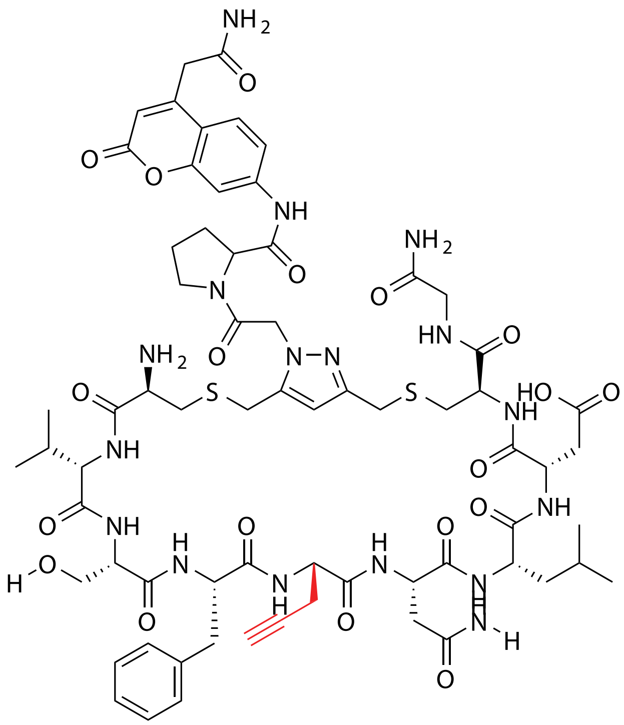

| Probe Name | Sequence X= Unnatural | Unnatural Amino Acid Number (X) | Expected Mass [M+1]+1 | Found Mass |
|---|---|---|---|---|
| FAP 33.2 (3) | CVSF**X**NLDCG | 3 | 1498.5839 | 1498.9 |

Total Ion Chromatogram

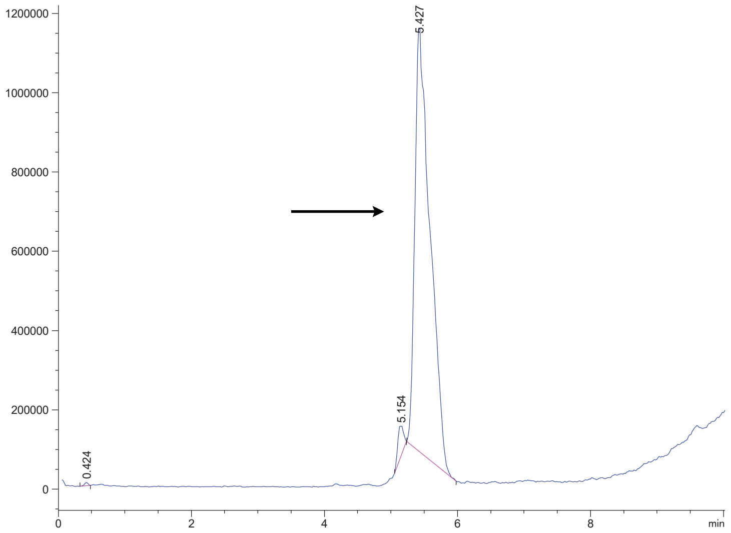

Variable Wavelength Detector: 220 nm

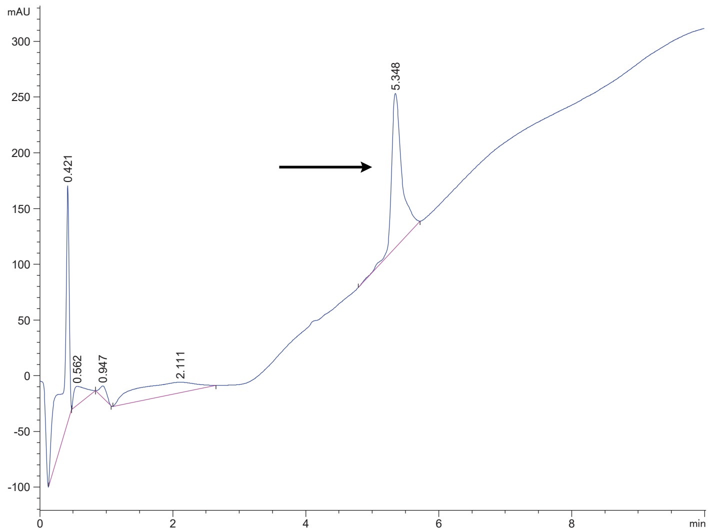

FAP 33.2 (4)

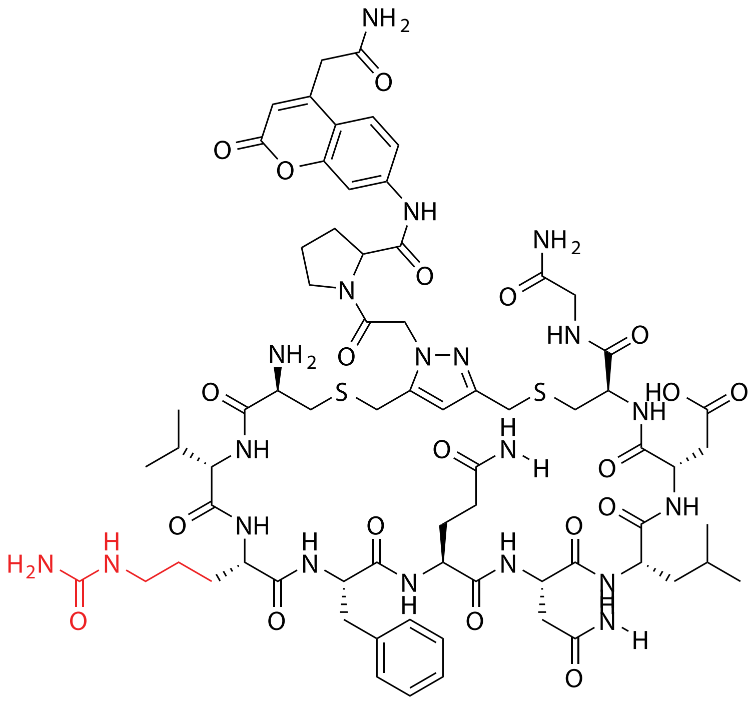

| Probe Name | Sequence<br>X= Unnatural | Unnatural Amino Acid Number (X) | Expected Mass $[M+1]^{+1}$ | Found Mass |
|---|---|---|---|---|
| FAP 33.2 (4) | CV**X**FQNLDCG | 4 | 1601.6585 | 1601.9 |

Total Ion Chromatogram

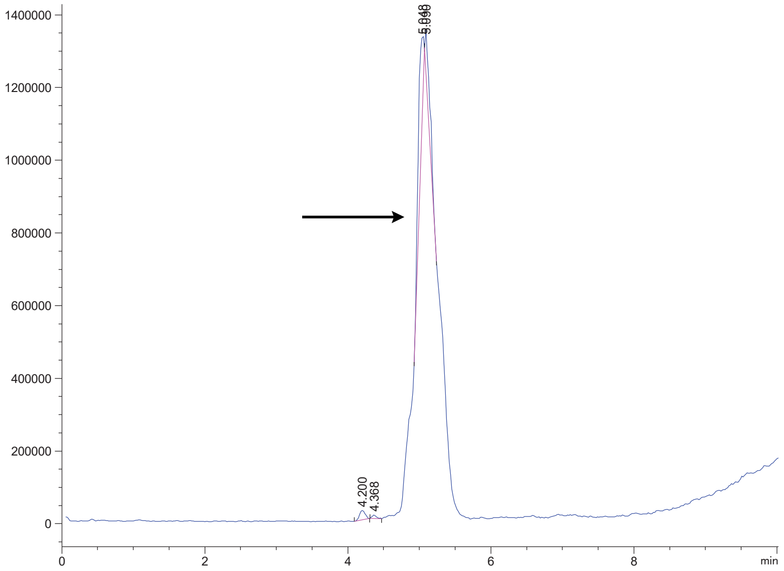


Variable Wavelength Detector: 220 nm

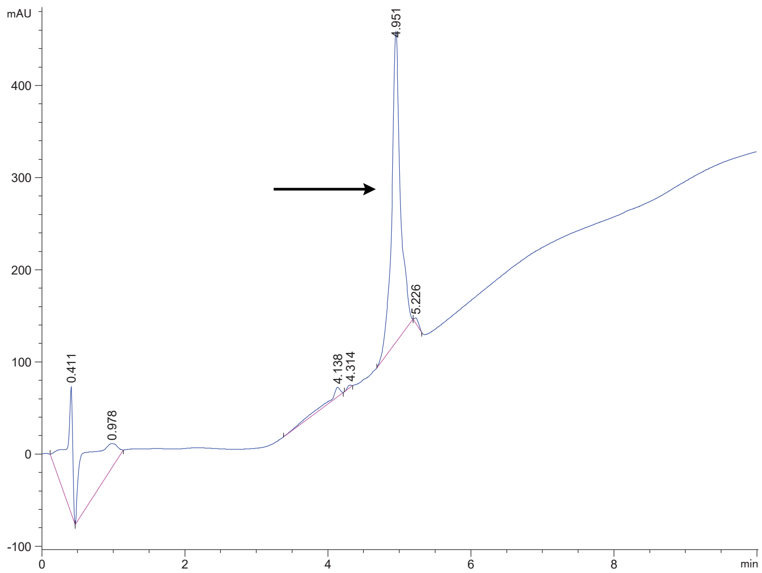

FAP 33.2 (7)

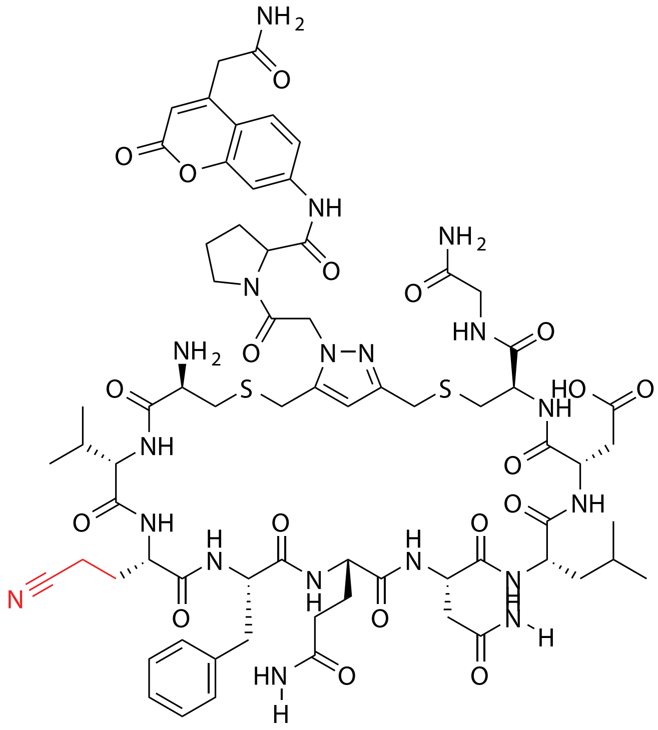

| Probe Name | Sequence X= Unnatural | Unnatural Amino Acid Number (X) | Expected Mass $[M+1]^{+1}$ | Found Mass |
|---|---|---|---|---|
| FAP 33.2 (7) | CV**X**FQNLDCG | 7 | 1554.6214 | 1555.0 |

Total Ion Chromatogram

Variable Wavelength Detector: 220 nm

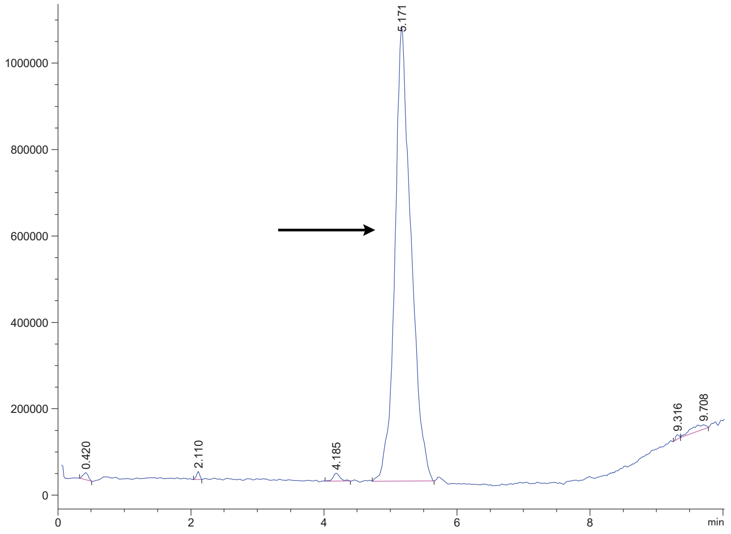


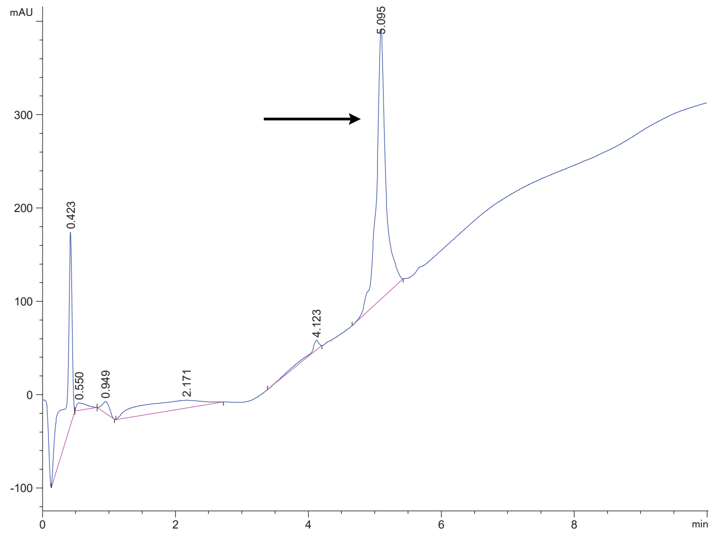

FAP 33.2 (8)

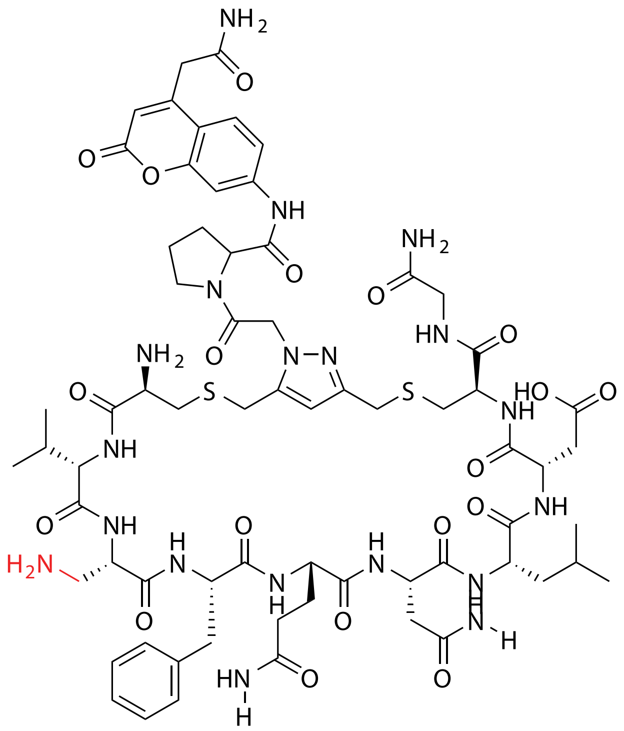

| Probe Name | Sequence X= Unnatural | Unnatural Amino Acid Number (X) | Expected Mass $[M+1]^{+1}$ | Found Mass |
|---|---|---|---|---|
| FAP 33.2 (8) | CV**X**FQNLDCG | 8 (Dap) | 1530.6214 | 1530.9 |

Total Ion Chromatogram

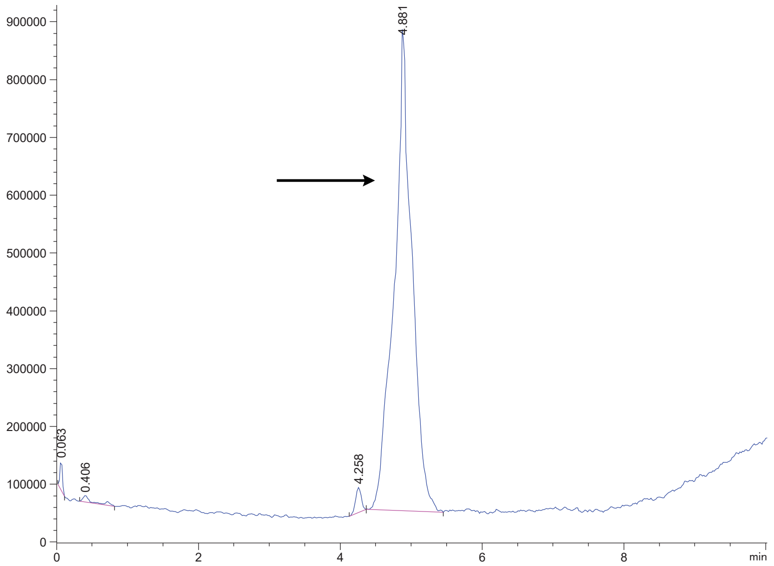


Variable Wavelength Detector: 220 nm

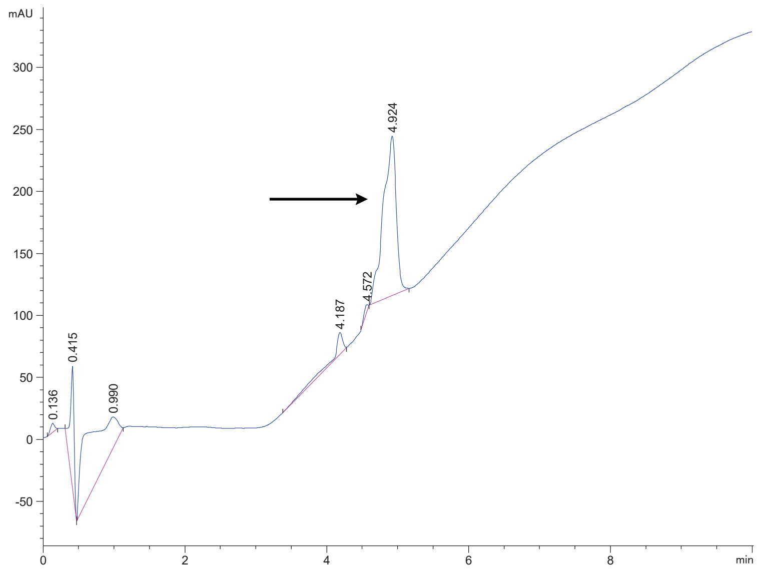

FAP 33.2 (11)

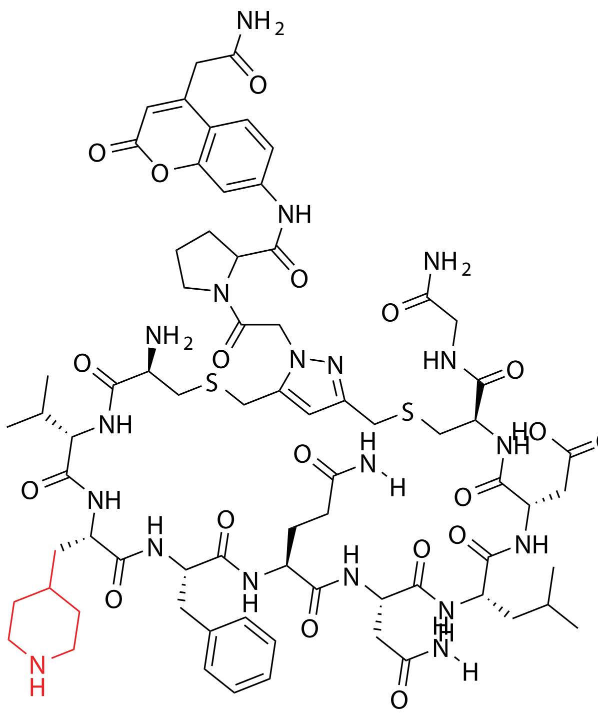


| Probe Name | Sequence<br>X= Unnatural | Unnatural Amino Acid Number (X) | Expected Mass $[M+1]^{+1}$ | Found Mass |
|---|---|---|---|---|
| FAP 33.2 (11) | CV**X**FQNLDCG | 11 | 1598.6840 | 1599.0 |

Total Ion Chromatogram

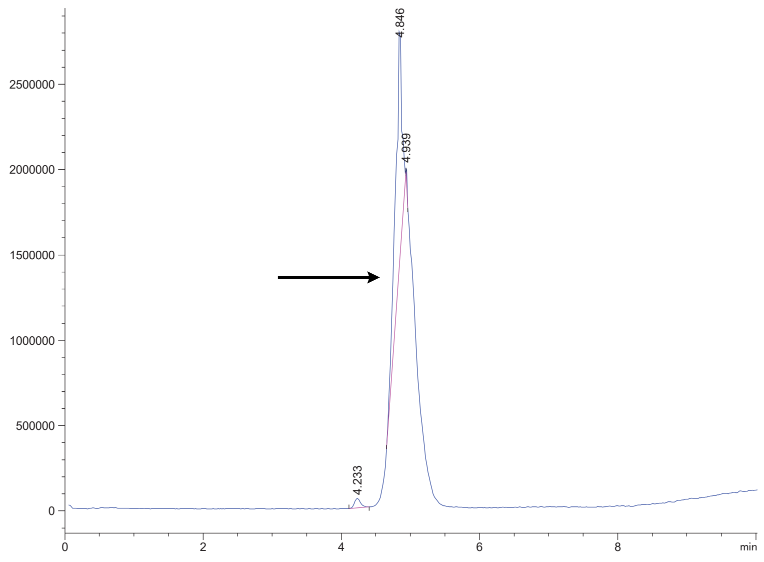


Variable Wavelength Detector: 220 nm

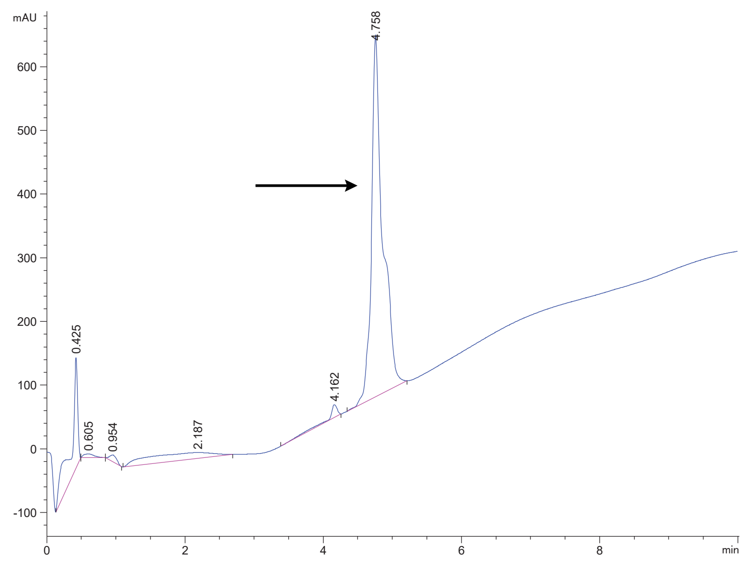

FAP 33.2 (12)

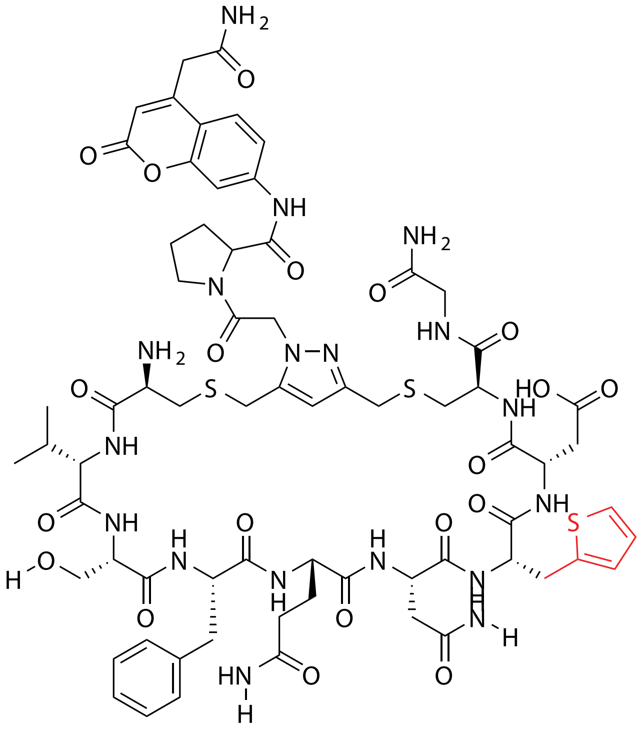

| Probe Name | Sequence X= Unnatural | Unnatural Amino Acid Number (X) | Expected Mass [M+1]$^{+1}$ | Found Mass |
|---|---|---|---|---|
| FAP 33.2 (12) | CVSFQN**X**DCG | 12 | 1571.5461 | 1571.8 |

Total Ion Chromatogram

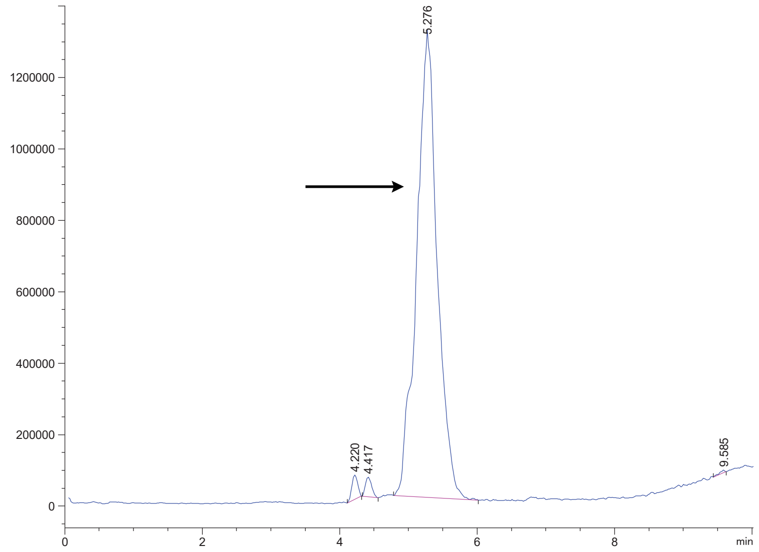


Variable Wavelength Detector: 220 nm

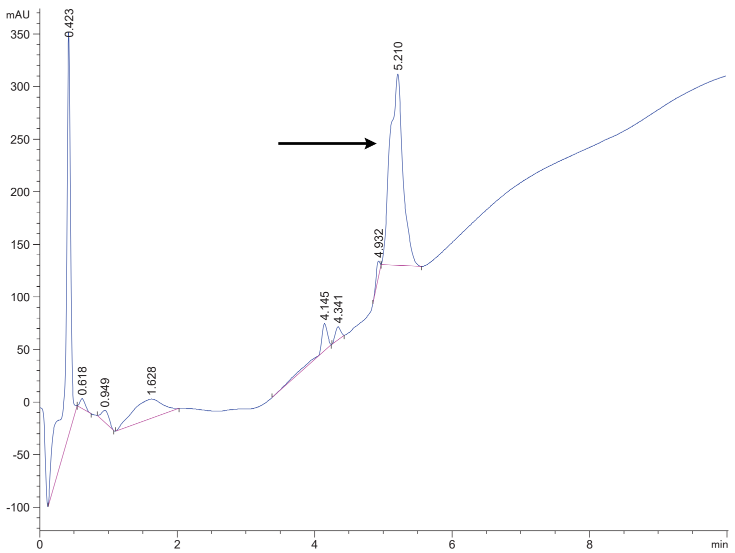

FAP 33.2 (13)

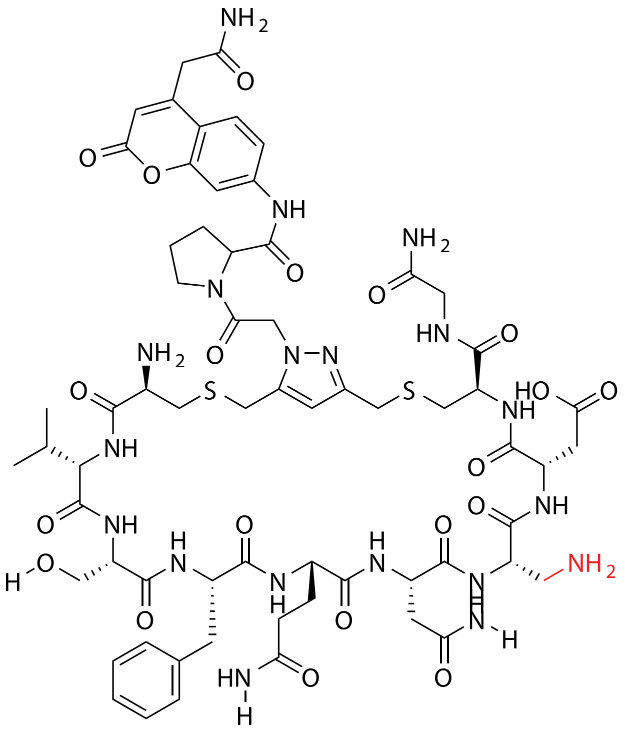

| Probe Name | Sequence X= Unnatural | Unnatural Amino Acid Number (X) | Expected Mass $[M+1]^{+1}$ | Found Mass |
|---|---|---|---|---|
| FAP 33.2 (13) | CVSFQN**X**DCG | 13 (Dap) | 1504.5693 | 1504.9 |

Total Ion Chromatogram

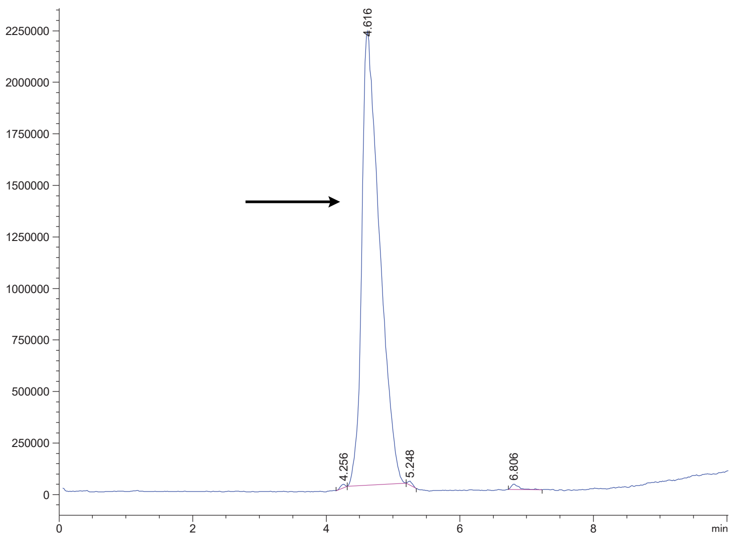


Variable Wavelength Detector: 220 nm

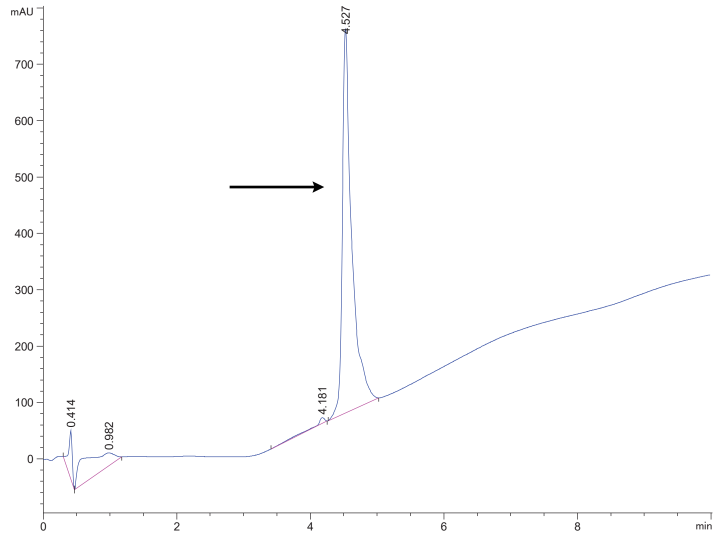

FAP 33.2 (14)

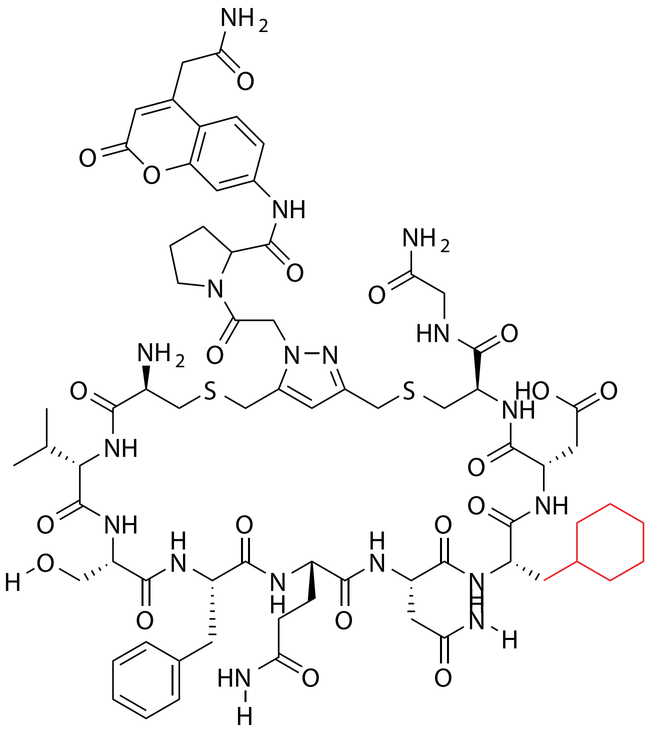

| Probe Name | Sequence X= Unnatural | Unnatural Amino Acid Number (X) | Expected Mass $[M+1]^{+1}$ | Found Mass |
|---|---|---|---|---|
| FAP 33.2 (14) | CVSFQN**X**DCG | 14 | 1571.6367 | 1571.9 |

Total Ion Chromatogram

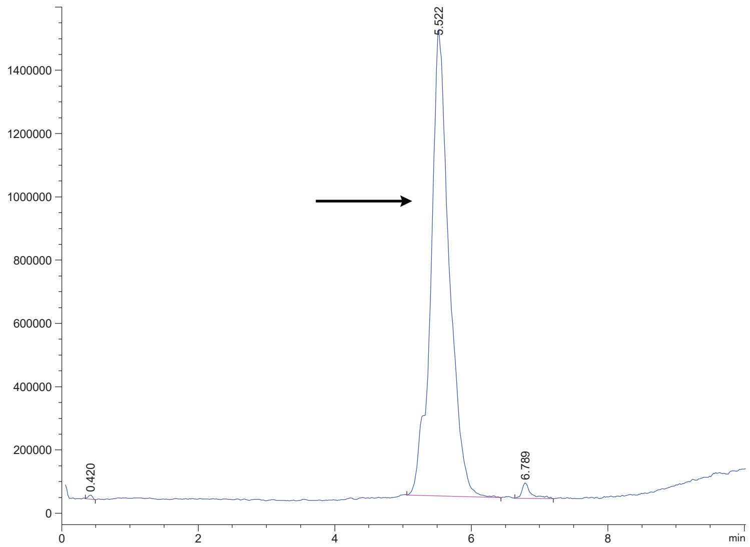


Variable Wavelength Detector: 220 nm

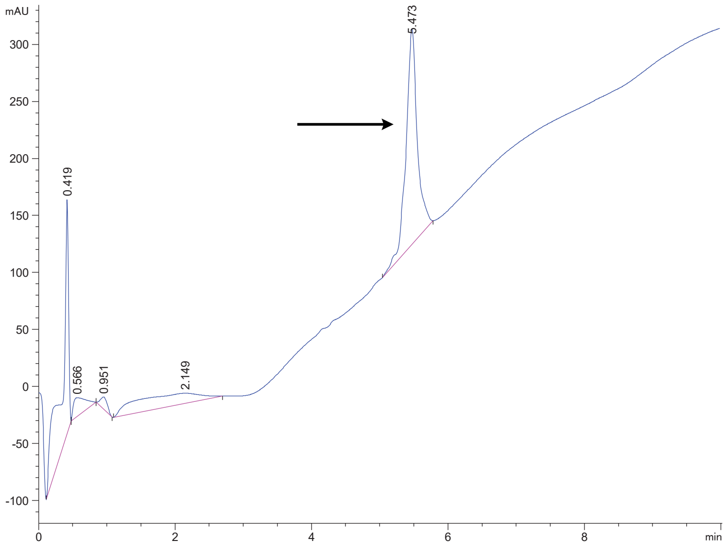

FAP 33.2 (15)

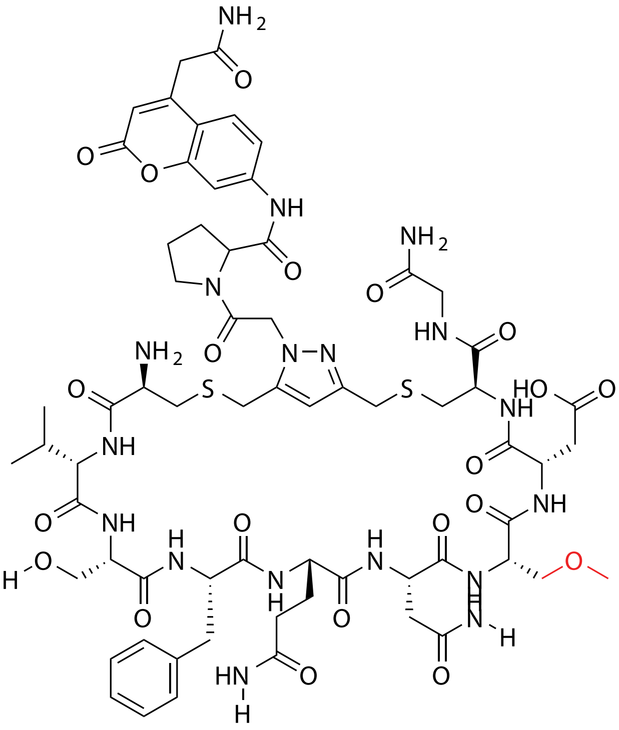


| Probe Name | Sequence X= Unnatural | Unnatural Amino Acid Number (X) | Expected Mass [M+1]$^{+1}$ | Found Mass |
|---|---|---|---|---|
| FAP 33.2 (15) | CVSFQN**X**DCG | 15 | 1519.5690 | 1519.8 |

Total Ion Chromatogram

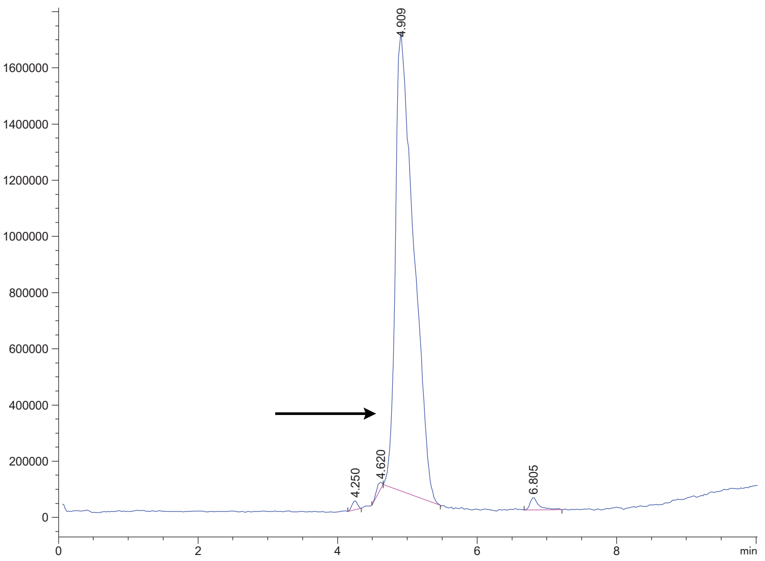


Variable Wavelength Detector: 220 nm

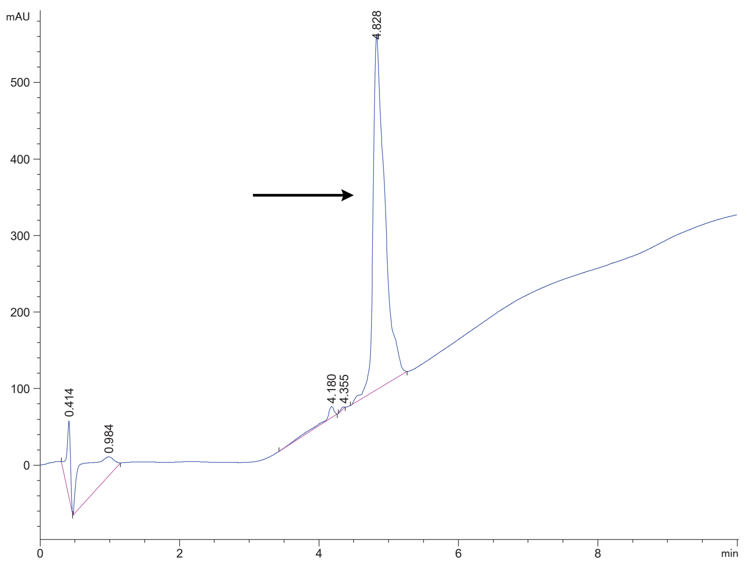

FAP 33.2 (16)

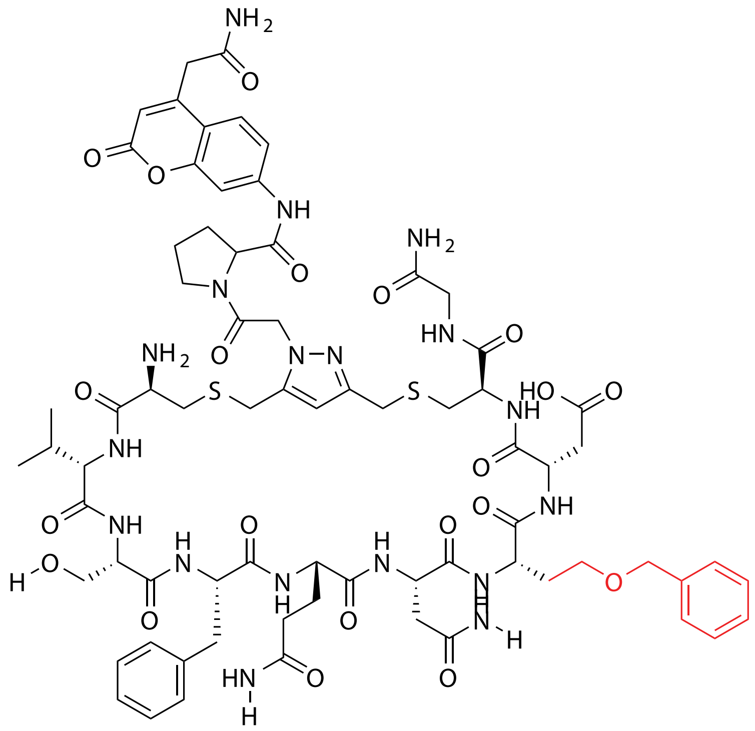

| Probe Name | Sequence X= Unnatural | Unnatural Amino Acid Number (X) | Expected Mass $[M+1]^{+1}$ | Found Mass |
|---|---|---|---|---|
| FAP 33.2 (16) | CVSFQN**X**DCG | 16 | 1609.6159 | 1609.9 |

Total Ion Chromatogram

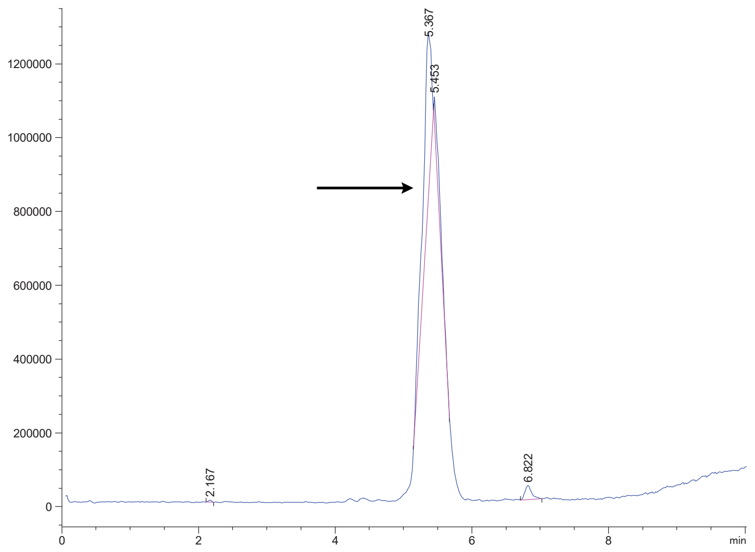


Variable Wavelength Detector: 220 nm

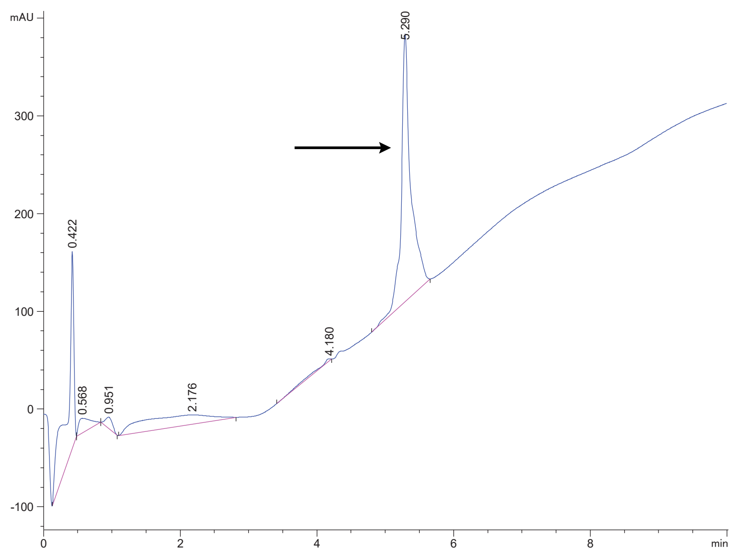

FAP 33.2 (17)

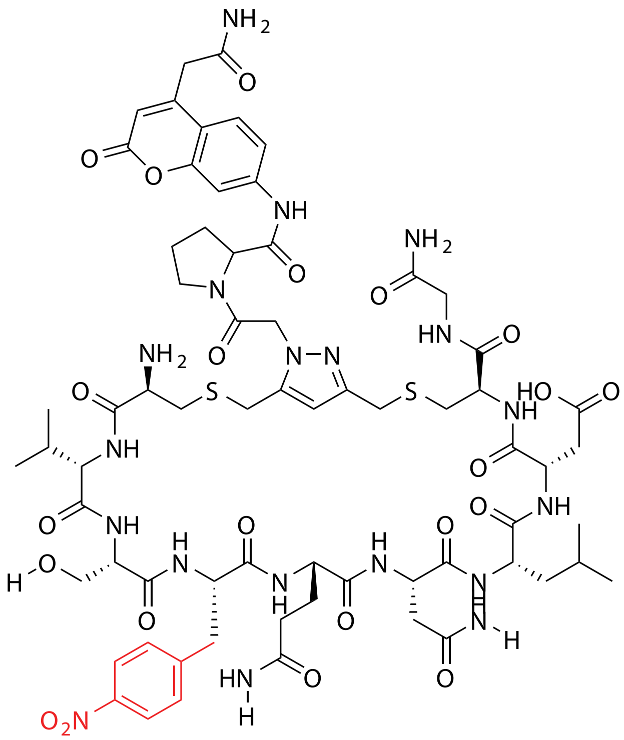


| Probe Name | Sequence X= Unnatural | Unnatural Amino Acid Number (X) | Expected Mass $[M+1]^{+1}$ | Found Mass |
|---|---|---|---|---|
| FAP 33.2 (17) | CVS**X**QNLDCG | 17 | 1576.5905 | 1576.9 |

Total Ion Chromatogram

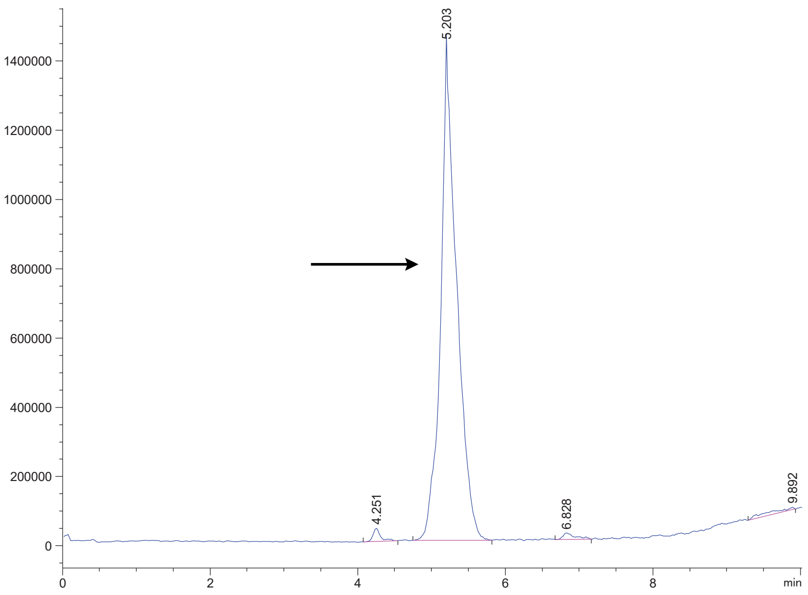


Variable Wavelength Detector: 220 nm

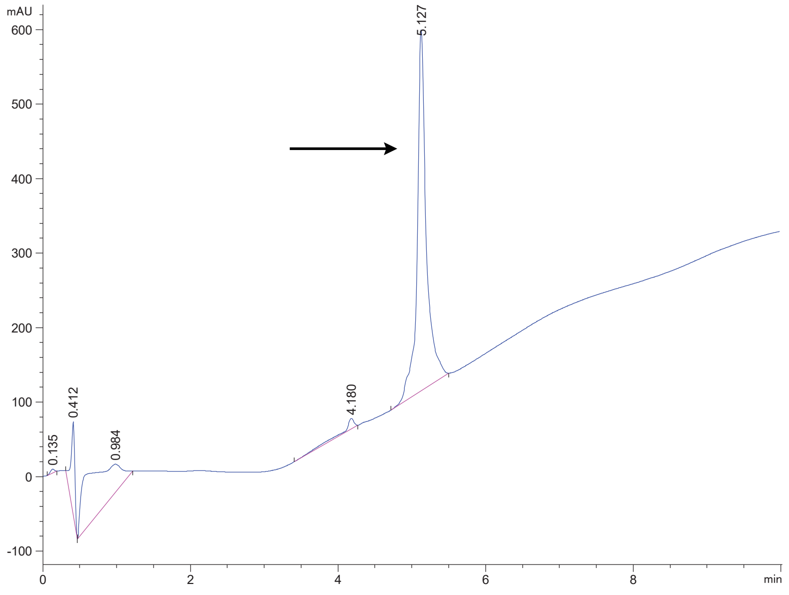

FAP 33.2 (18)

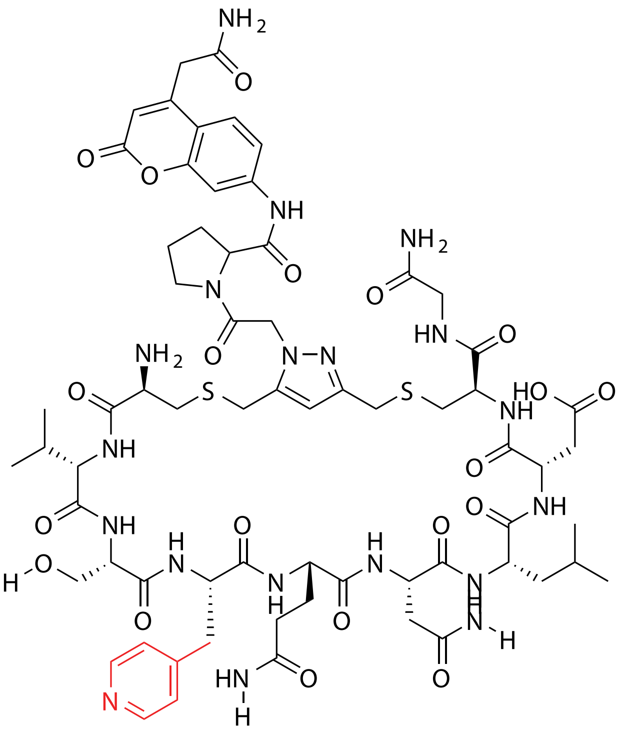

| Probe Name | Sequence X= Unnatural | Unnatural Amino Acid Number (X) | Expected Mass $[M+1]^{+1}$ | Found Mass |
|---|---|---|---|---|
| FAP 33.2 (18) | CVS**X**QNLDCG | 18 | 1532.6006 | 1532.9 |

Total Ion Chromatogram

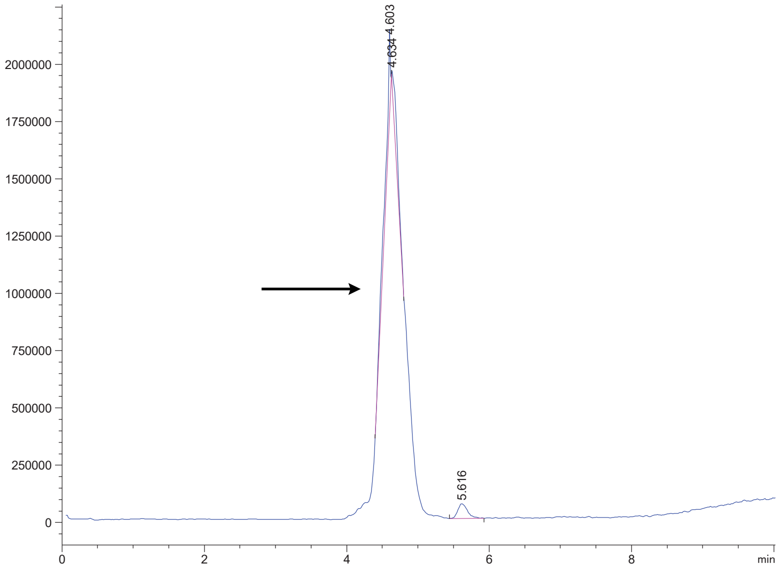

Variable Wavelength Detector: 220 nm

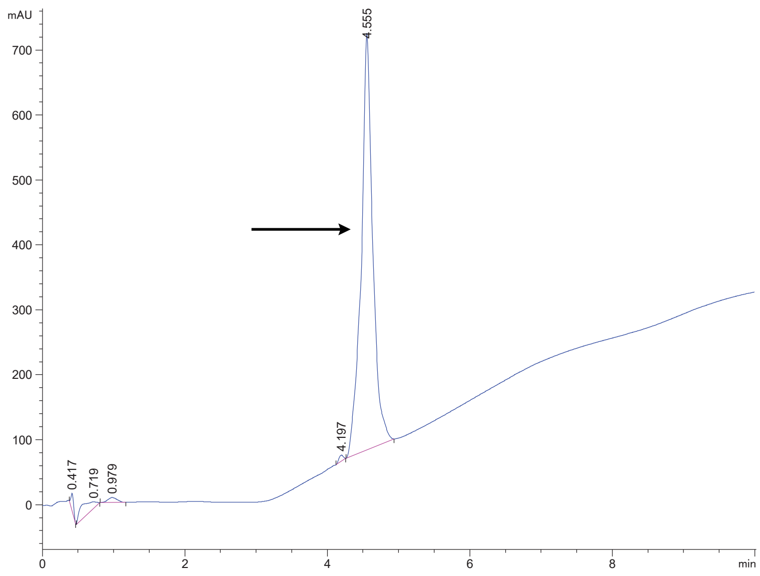

FAP 33.2 (19)

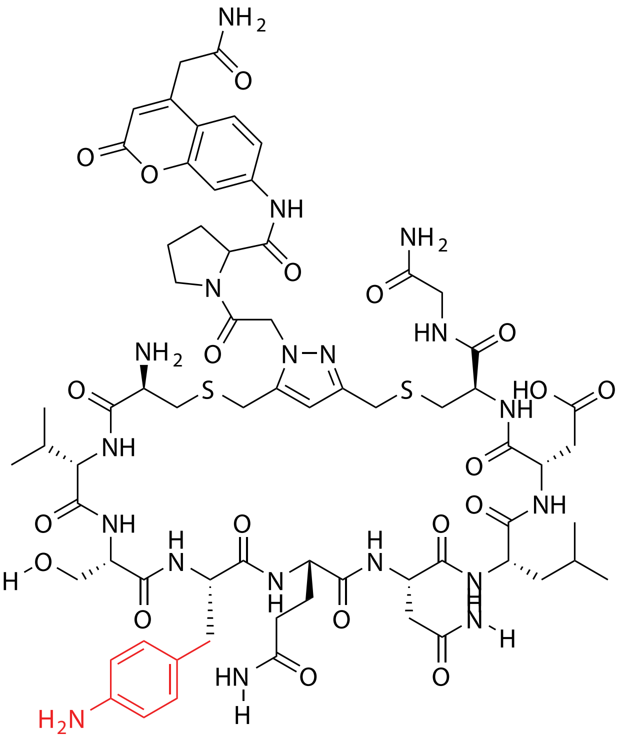

| **Probe Name** | **Sequence X= Unnatural** | **Unnatural Amino Acid Number (X)** | **Expected Mass [M+1]$^{+1}$** | **Found Mass** |
|---|---|---|---|---|
| FAP 33.2 (19) | CVS**X**QNLDCG | 19 | 1546.6163 | 1546.9 |

Total Ion Chromatogram

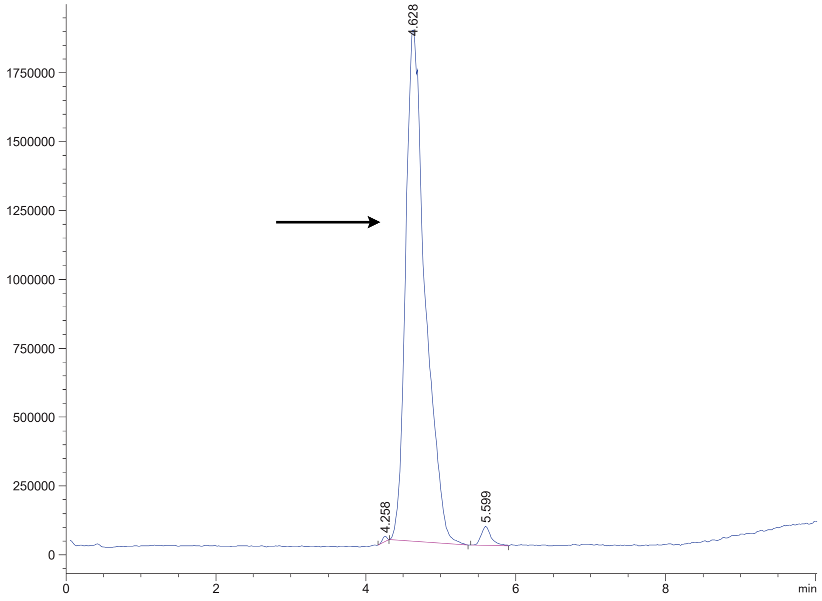


Variable Wavelength Detector: 220 nm

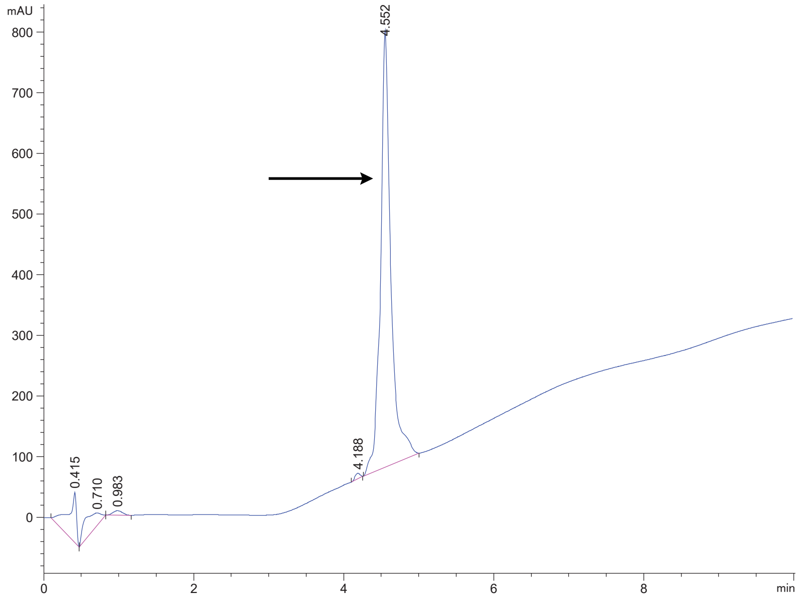

FAP 33.2 (20)

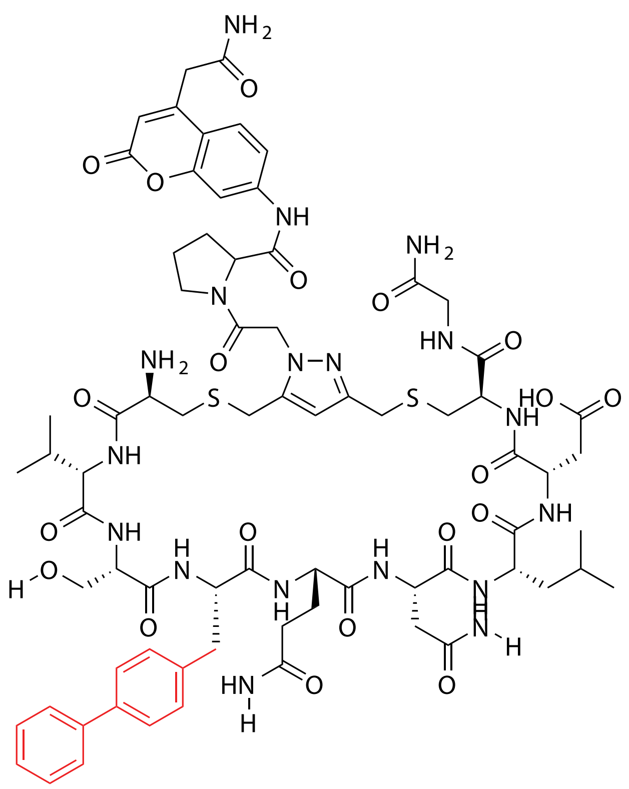

| Probe Name | Sequence X= Unnatural | Unnatural Amino Acid Number (X) | Expected Mass [M+1]$^{+1}$ | Found Mass |
|---|---|---|---|---|
| FAP 33.2 (20) | CVS**X**QNLDCG | 20 | 1607.6367 | 1607.9 |

Total Ion Chromatogram

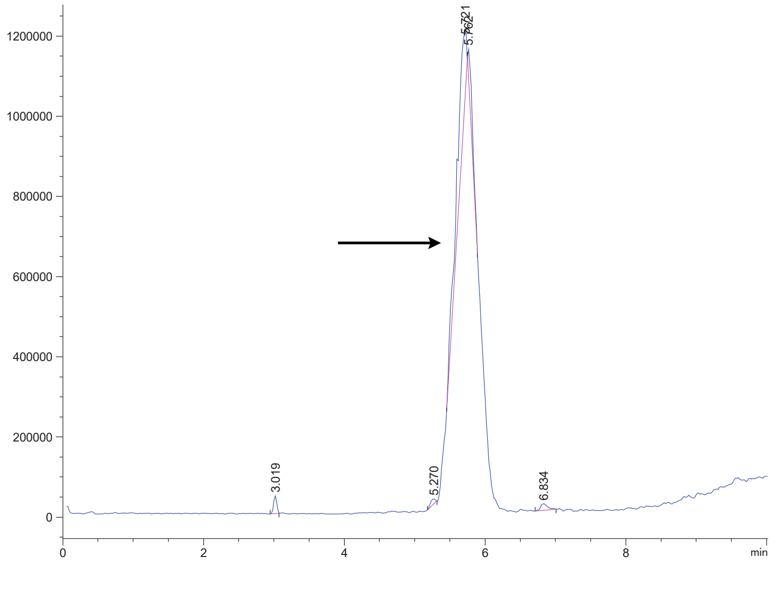

Variable Wavelength Detector: 220 nm

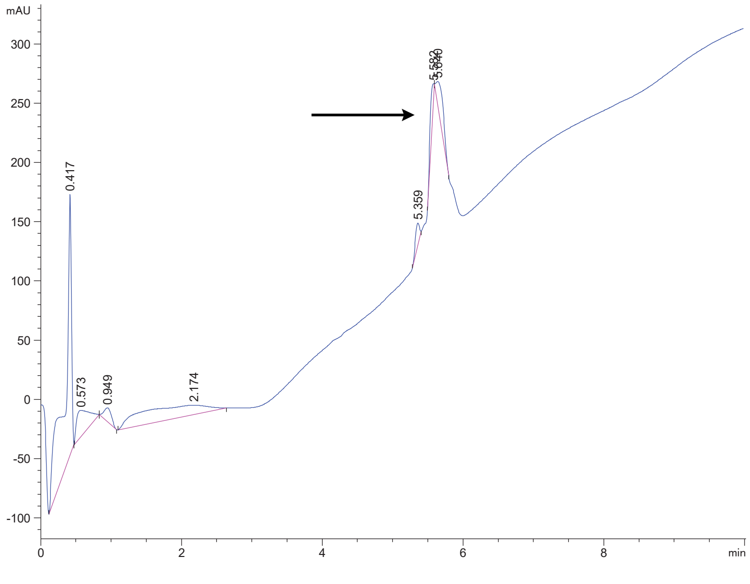

FAP 33.2 (21)

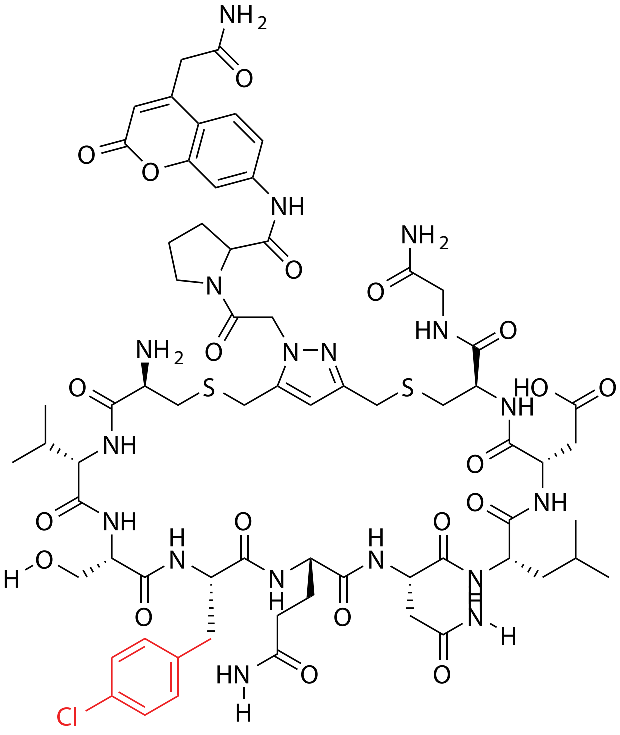

| Probe Name | Sequence X= Unnatural | Unnatural Amino Acid Number (X) | Expected Mass $[M+1]^{+1}$ | Found Mass |
|---|---|---|---|---|
| FAP 33.2 (21) | CVS**X**QNLDCG | 21 | 1565.5664 | 1565.8 |

Total Ion Chromatogram

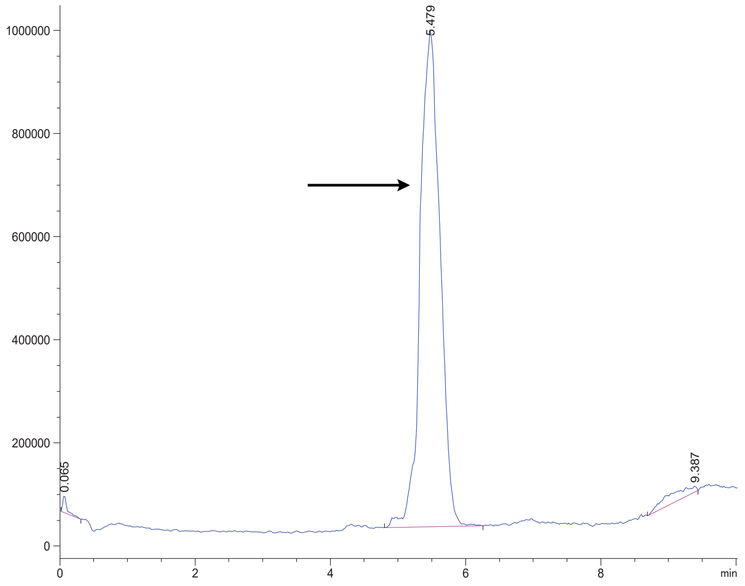


Variable Wavelength Detector: 220 nm

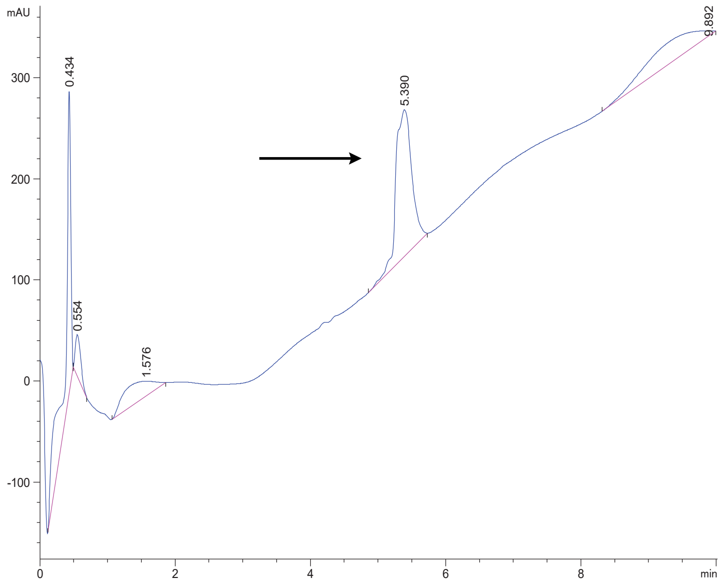

FAP 33.2 (22)

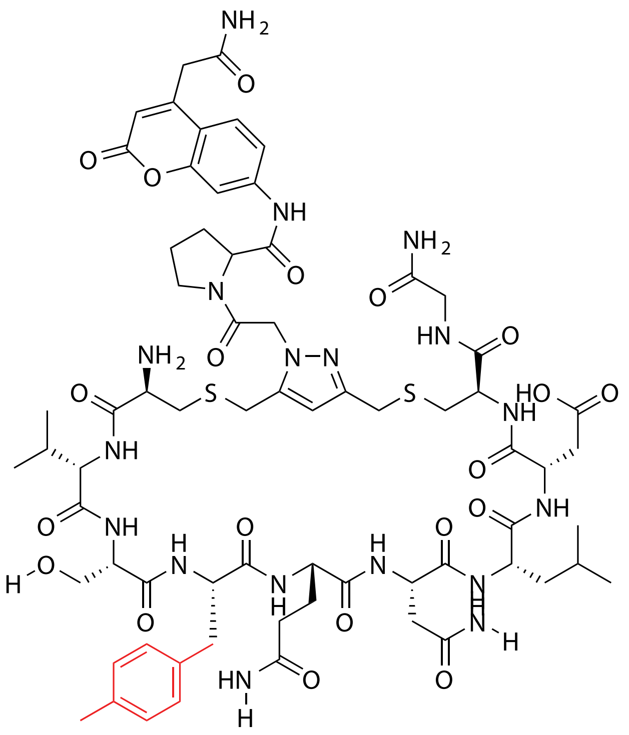

| Probe Name | Sequence X= Unnatural | Unnatural Amino Acid Number (X) | Expected Mass $[M+1]^{+1}$ | Found Mass |
|---|---|---|---|---|
| FAP 33.2 (22) | CVS**X**QNLDCG | 22 | 1545.6210 | 1545.9 |

Total Ion Chromatogram

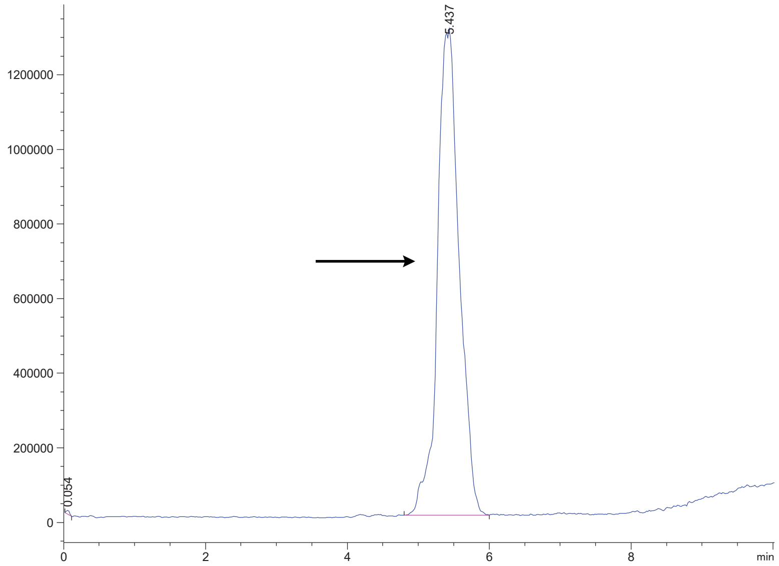


Variable Wavelength Detector: 220 nm

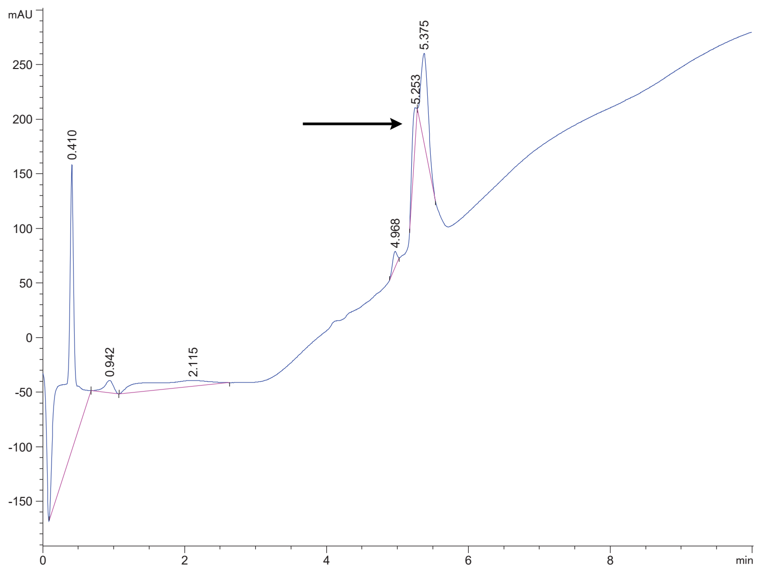

FAP 33.2 (23)

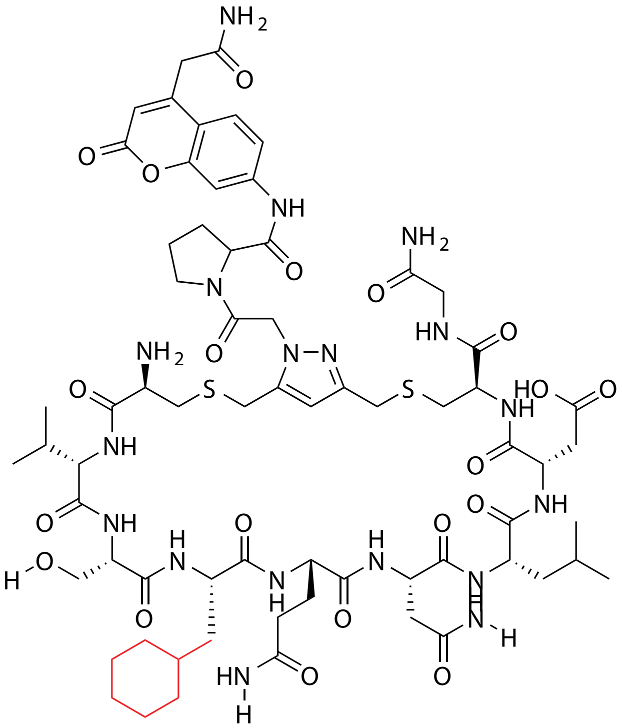


| Probe Name | Sequence X= Unnatural | Unnatural Amino Acid Number (X) | Expected Mass $[M+1]^{+1}$ | Found Mass |
|---|---|---|---|---|
| FAP 33.2 (23) | CVS**X**QNLDCG | 23 | 1537.6523 | 1537.9 |

Total Ion Chromatogram

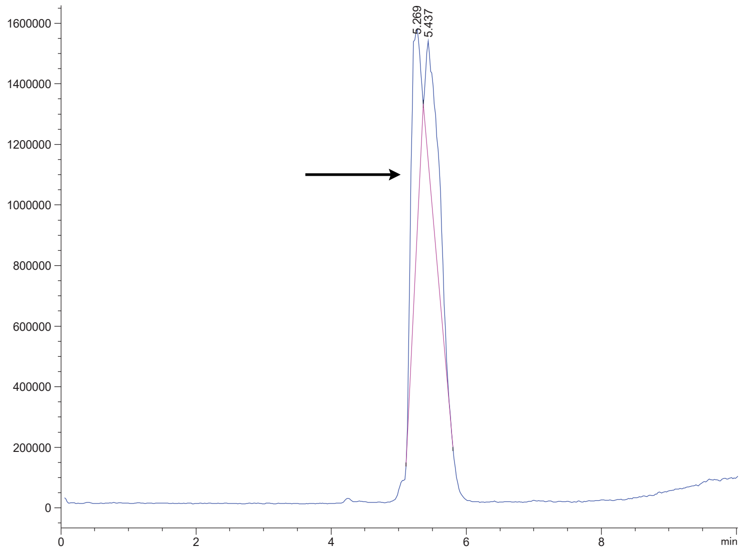


Variable Wavelength Detector: 220 nm

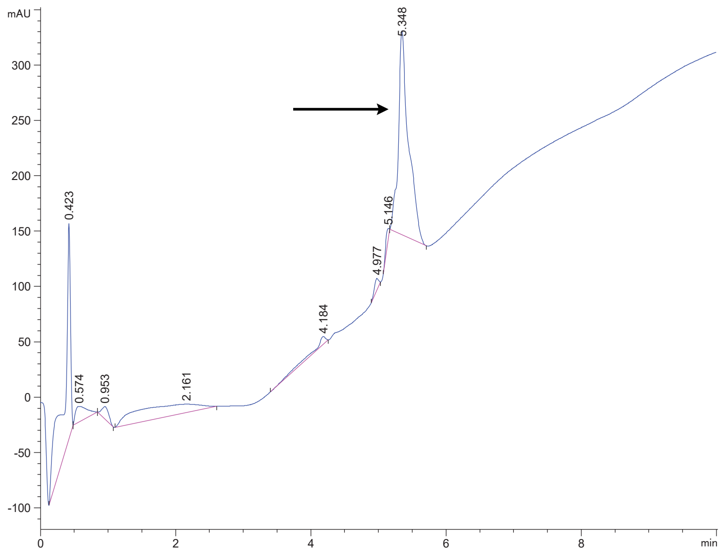

FAP 33.2 (24)

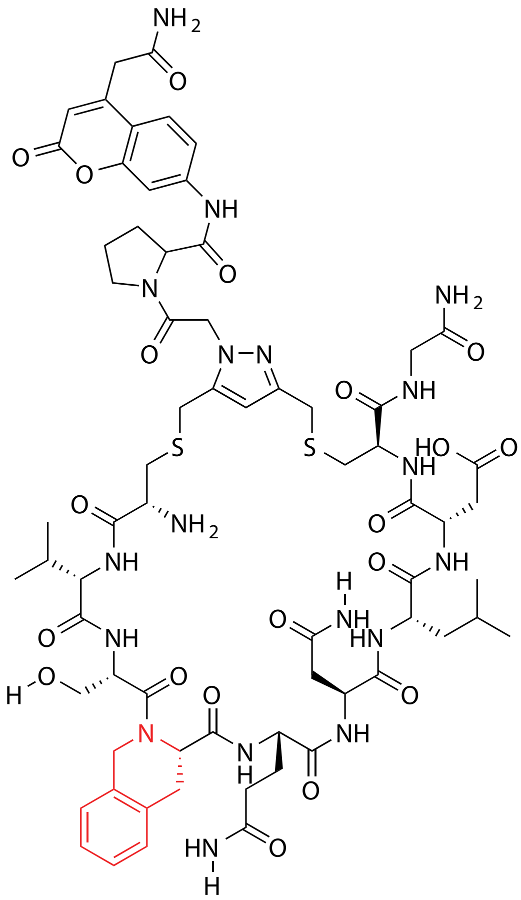

| Probe Name | Sequence X= Unnatural | Unnatural Amino Acid Number (X) | Expected Mass $[M+1]^{+1}$ | Found Mass |
|---|---|---|---|---|
| FAP 33.2 (24) | CVS**X**QNLDCG | 24 | 1543.6054 | 1543.9 |

Total Ion Chromatogram

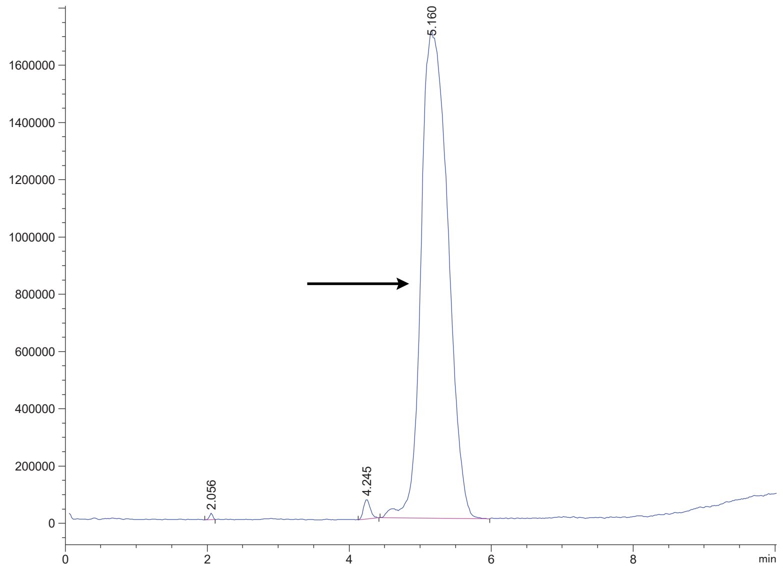


Variable Wavelength Detector: 220 nm

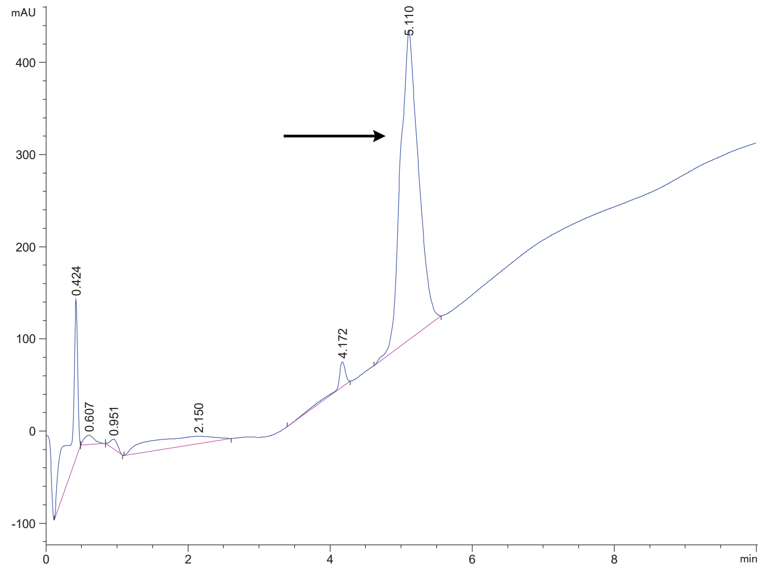

FAP 33.2alk

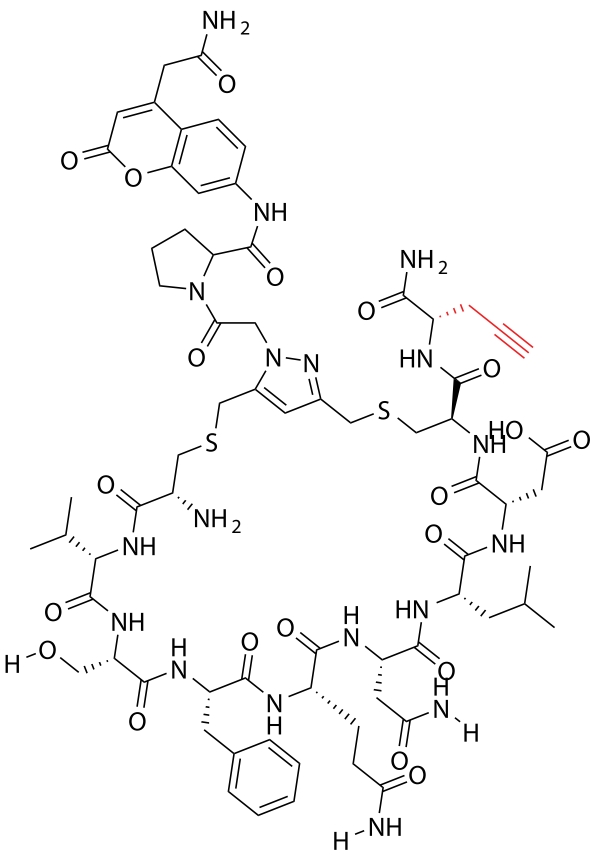

| Probe Name | Sequence | Expected Mass $[M+1]^{+1}$ or $[M+2]^{+2}$ | Found Mass |
|---|---|---|---|
| FAP 33.2alk | CVSFQNLDCalk | 1569.6210 | 1569.9 |

Total Ion Chromatogram

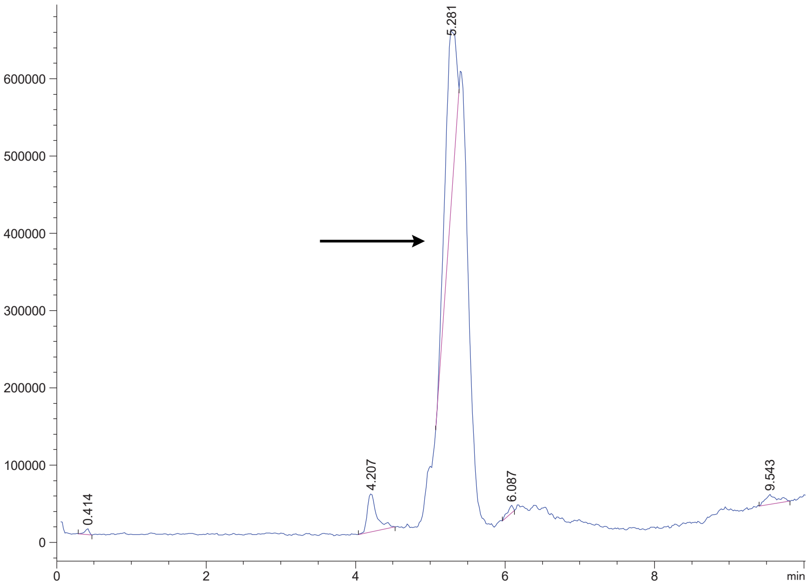


Variable Wavelength Detector: 220 nm

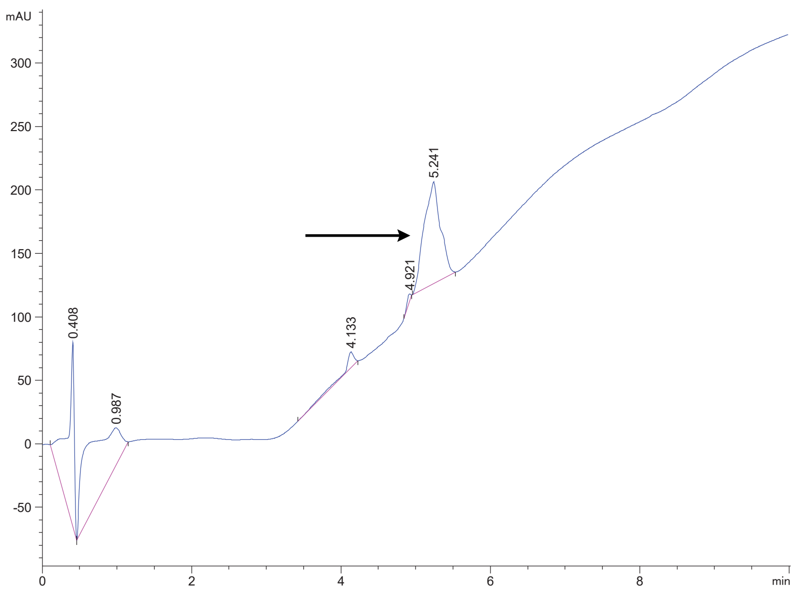

FAP 33.2ac

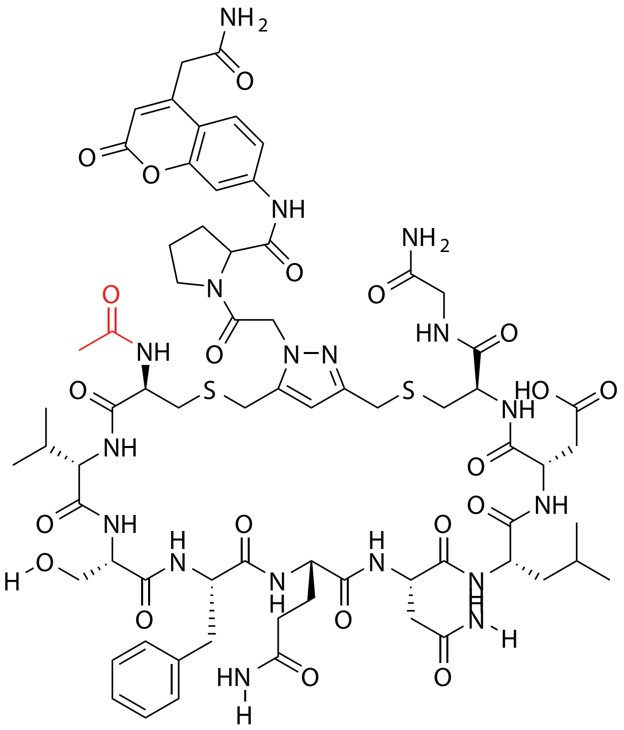

| Probe Name | Sequence | Expected Mass [M+1]$^{+1}$ or [M+2]$^{+2}$ | Found Mass |
|---|---|---|---|
| FAP 33.2ac | Acetyl-CVSFQNLACG | 1573.6159 | 1573.8 |

Total Ion Chromatogram

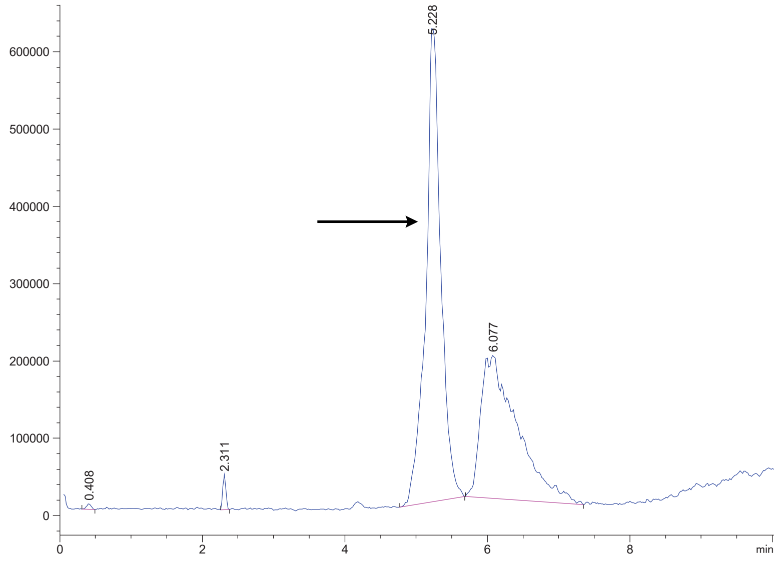


Variable Wavelength Detector: 220 nm

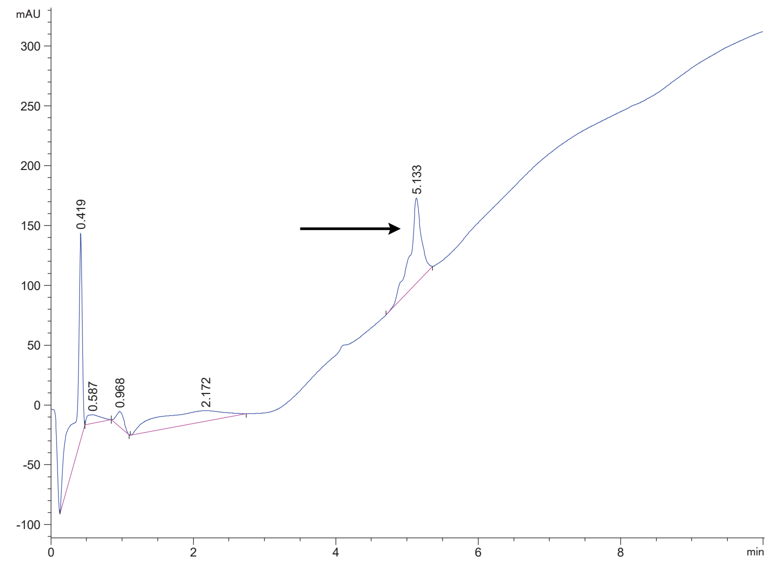

FAP 33.2 (2)

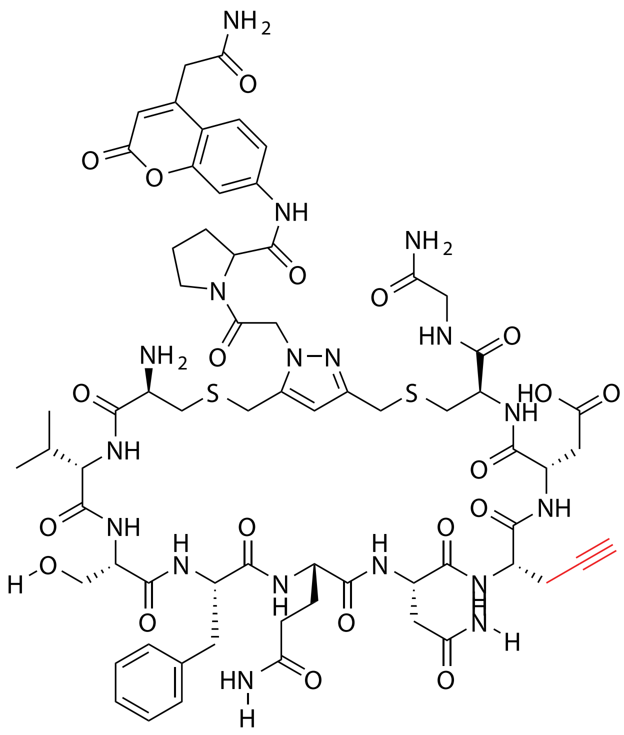

| Probe Name | Sequence X= Unnatural | Unnatural Amino Acid Number (X) | Expected Mass [M+1]$^{+1}$ | Found Mass |
|---|---|---|---|---|
| FAP 33.2 (2) | CVSFQN**X**DCG | 2 | 1513.5584 | 1513.8 |

Total Ion Chromatogram

Variable Wavelength Detector: 220 nm

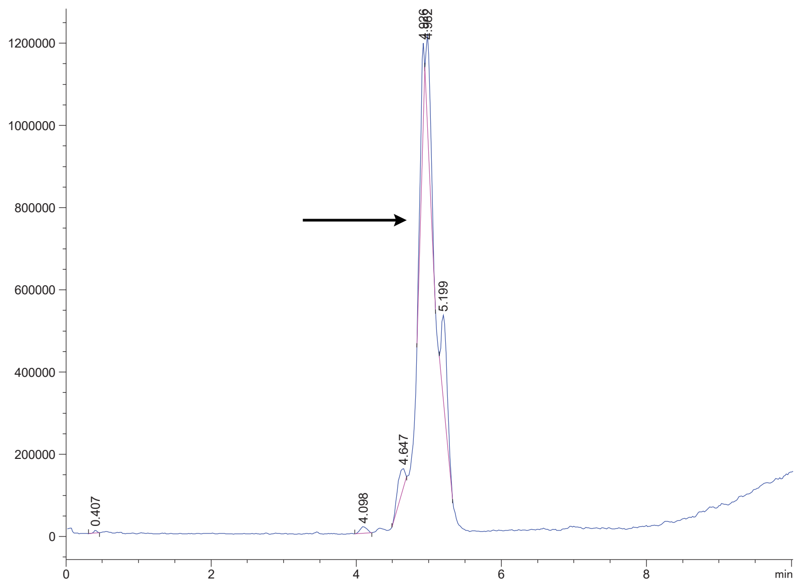

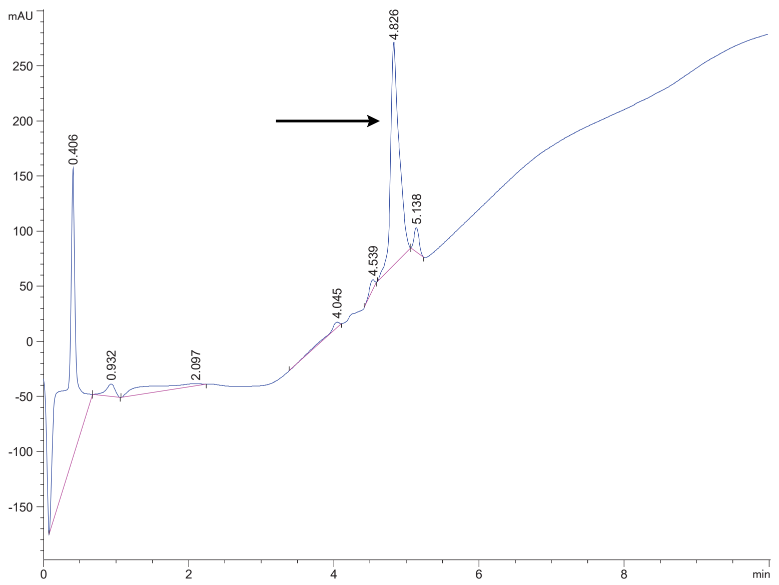
